# Decomposed Linear Dynamical Systems (dLDS) models reveal instantaneous, context-dependent dynamic connectivity in *C. elegans*

**DOI:** 10.1101/2024.05.31.596903

**Authors:** Eva Yezerets, Noga Mudrik, Adam S. Charles

## Abstract

Mounting empirical evidence indicates that neural “tuning” can be highly variable within an individual across time and especially across individuals. Furthermore, modulatory effects can change the *relationship* between neurons in the brain as a function of behavioral or other conditions, meaning that the changes in activity (the derivative) may be as important as the activity itself. However, current computational models fail to capture the nonstationarity and variability of neural coding, preventing the quantitative evaluation of these effects, especially during individuals’ adaptation to their environments. Here we present a novel way to study the effects of adaptation in one of the most well-studied organisms, *C. elegans*, leveraging recent advances in dynamical systems modeling, specifically decomposed Linear Dynamical Systems (dLDS).Our approach enables the discovery of multiple parallel neural processes on different timescales using a low-dimensional set of linear operators that can be recombined in different ratios. Our model identifies “dynamic connectivity,” describing patterns of dynamic neural interactions in time. We use these patterns to identify instantaneous, contextually-dependent, hierarchical roles of neurons; discover the underlying variability of neural representations even under seemingly discrete behaviors; and learn a single aligned latent space underlying multiple individual worms’ activity. By analyzing individual worms and neurons, we found evidence that 1) changes in interneuron connectivity mediate efficient task-switching and 2) changes in sensory neuron connectivity show a mechanism of adaptation.

## 1. Introduction

Neural systems must continuously adapt to the ever-changing and multifacted environments that animals navigate. Such modulation of activity has been observed across species and occurs on timescales from milliseconds to minutes. Understanding how systems modulate their internal dynamics to adapt their computation and match novel experiences is thus critical in learning how the brain flexibly and robustly drives behaviors. However, models of time-varying activity-based population-level interactions remain limited, in part due to the complexity of estimating ever-changing interaction terms in large-scale data [1; 2]. *C. elegans*, whose small size and transparency have enabled scientists to capture its “whole-brain” activity and create a wealth of literature on individual neurons’ functions, thus offers a unique opportunity to study the modulation of neural coding.

Using only 302 neurons, *C. elegans* can pirouette, navigate mazes, adapt to stimuli, and even learn associations [1; 3; 4; 5; 6]. These sensorimotor and cognitive processes all occur in parallel with other homeostatic functions. *C. elegans* is also an incredibly well-mapped organism. The anatomical connectome and proposed functions for almost all of its 302 neurons are available [7; 8; 9; 10; 11]. While the time-averaged neural activity required to execute its complex repertoire of behaviors appears as a stable connectome, the anatomical scaffold alone is insufficient to understand the mechanisms that *C. elegans* implements. Analysis over time within one animal [12] or between animals [12; 13; 14] reveals variability in *C. elegans* population neural coding. This is not noise; this variability is *required* for adaptation and reflects worm individuality in behavioral tendencies.

Recent work has revealed that the diversity of signaling mechanisms and timescales in *C. elegans* may be the reason why the stationary, i.e., time-averaged, anatomical connectome doesn’t tell the full story. Randi *et al.* 2023 [11] showed a number of discrepancies between the functional connections recorded from pairs of neurons and the activity predicted based on the anatomical path lengths between them [11]. Moreover, the connectome of long-distance chemical messengers (e.g., neuropeptides) is being mapped [15; 16], and its concordance with the anatomical connectome appears limited, or at least complex [1; 17]. For one, *C. elegans* neurons can communicate both with and without traditional action potentials—consider the nociceptor ASH, which can generate graded responses to stimuli. Action potentials in the traditional sense are not required for signaling from ASH to AVA, a key interneuron [18]. Instead, synaptic communication is predominantly based on graded potentials. For another, pairs of cells can communicate with each other on multiple timescales, from milliseconds across gap junctions, to seconds for action potentials, to minutes across gap junctions again, with neuromodulators concurrently playing a role on the timescale of seconds to minutes. For example, these overlapping timescales of signaling are present in the defecation-related pacemaker signal [19; 20].

It is particularly difficult to map function onto the *C. elegans* anatomical connectome because direct, synaptic signaling and long-distance chemical messengers may operate on similar timescales to each other in this species [11]. Therefore, either a multi-hop connection directly across neurons or a neuropeptide could reasonably implement a seconds-long functional response that is relevant to a seconds-timescale behavior. Moreover, multiple processes happen on overlapping timescales in *C. elegans*, including homeostatic processes such as managing energy levels and attention [21], and even its most clearly annotated neural pathways are used in slightly different ratios throughout behavioral motifs [22]. All of these factors contribute to the challenge of determining each neuron’s function in *C. elegans*, even if those functions *were* stable and universal across worms and conditions. However, *C. elegans* also generates *nonstationary* cognitive processes and behaviors. In fact, Atanas *et al.* 2023 [23] showed that 20 to 30% of *C. elegans* neurons changed their behavior encodings between the first and second halves of a single recording period, either via gain modulation, changes in tuning, or simply changing which behaviors they encoded. These encodings are also known to be context-dependent, e.g., the AIA neuron’s sensory-context-dependent role in roaming vs. dwelling behavior depending on food in the environment [24]; BAG’s behavioral-context-dependent role in self-produced (reafferent) oxygen sensing [25]; RIM’s context-dependent role in mechanosensation during turns vs. other behaviors [26]; AWC’s context-dependent role in deciding whether the worm will be attracted to an odor, or will avoid it [27]; and context-dependent differences in nervous system-wide sensory-interneuron-motor interactions during male-hermaphrodite *C. elegans* mating [28]. The mechanisms governing these internal neural states seem to be largely neuromodulatory [29], as opposed to purely synaptic, thus requiring neural population-level analysis. In particular, the context-dependent roles of interneurons are especially difficult to pin down — existing models do not have the granularity to describe their complex and changing roles in neural encoding.

Moreover, the species has demonstrated associative and non-associative learning (e.g., habituation, sensitization) [30; 31] via chemosensation, mechanosensation, and proprioception, with short-term memory shown to persist over minutes [4] and long-term memory over hours [6] to up to a day [32; 33; 34]. Because learning can occur over the course of a minutes-long trial in this species, stationary functional connectomes averaged across trials are meaningful for understanding the general functions of certain networks of neurons, but they are insufficient for understanding how neurons create behavioral changes over the seconds-to-minutes timescale.

Finally, even without nonstationarity, behaviors that appear stereotyped are not completely so; each worm’s neural dynamics are unique [13; 14]. For example, some worms simply tend to roam more than other worms do [13], and worms can also develop these differences throughout development [12].

To address these gaps in mechanistic understanding, multiple groups have characterized aspects of *C. elegans*’ stationary functional connectome [11; 15]. For example, recent work has described functional connectivity modes via principal component analysis (PCA) [23], and has used topologically motivated analyses [35] to conclude that the redundancy and flexibility of *C. elegans* neural circuitry means that only perturbing certain hub neurons has a significant impact on function. Moreover, another recent PCA-based linear time-invariant low-dimensional manifold model of *C. elegans* [36] showed that behavior-related brain states have wide-reaching effects across the *C. elegans* nervous system, creating global brain states that hierarchically gate neuron activity and allow for distributed coding. These examples show the mechanistic insights about *C. elegans* neural coding that can be derived time-averaged models.

However, to track *time-varying* relationships between cognitive states or processes both within and across individuals, we need dynamical systems models. Such models reinterpret high-dimensional time series data, such as neural population activity, as noisy manifestations of underlying, lower-dimensional processes that evolve within certain constraints in the state space, e.g., on a manifold. The manifold can be shared across individuals, representing a common functional landscape of cognitive processes. The evolution between latent states can be described by trajectories, i.e., dynamics. With such a model, it is possible to represent latent functional states at each time point and map those back to the neural activity patterns for each subject to examine dynamic connectivity. To date, time-varying models of *C. elegans* neural dynamics have primarily employed switching linear dynamical systems (SLDS) [37; 38; 39]. Such models assign a single, discrete state to each time point or time window, but the states are nonstationary, unlike in dynamic mode decomposition, for example [40]. SLDS models are locally linear and hierarchical, with continuous latent states being controlled by discrete overall neural population states. These have shown strong concordance between model-inferred neural states and experimentally-labelled behavioral states. However, such models are limited by the assumption that the entire neural population is experiencing the same overall state, and that these overall states transition abruptly from one to the other.

A non-switching, nonlinear model of *C. elegans* also found whole-worm neural dynamics reflecting two primary behavior states, forward and reverse crawling, as two loops on a low-dimensional manifold [14]. This model successfully predicts neural activity and behavior across individuals. However, the authors acknowledged that the worms had a high degree of individuality, both in terms of which neurons were identified in each recording and in terms of their behavior and utilization of neurons to execute similar behaviors. Because the gross behaviors could be decoded from almost any individual neuron with some fidelity, it was possible to build a shared dynamics model across worms. Moreover, this model is nonlinear, which makes its components and their interactions less interpretable than a linear model. Thus, there remains a need for linear models that do not abstract away from worm individuality in order to help researchers understand the mechanisms of how individual worms adjust their neural activity and connectivity in order to adapt to their environments, or even to learn.

Not all dynamical systems models capture time-varying dynamics. A linear time-invariant model [41] demonstrated the value of anatomical-connectome-constrained models in recapitulating the expected effects of optogenetically stimulating one neuron at a time and recording from other neurons. This paper claimed that while multi-hop, indirect, extrasynaptic connections do exist in the worm, the effects of single neurons on the system are well-modeled using what we know about *C. elegans* synaptic connections. However, this paper’s definitions of function, causality, and anatomical connectivity are time-averaged. While interpretable due to linearity and the anatomical-connectome-based restrictions on the weights, these constraints prevent the model from accounting for adaptation and learning, which also relies on extrasynaptic signaling.

We address these gaps in available modeling frameworks with our recently developed decomposed linear dynamical systems (dLDS) model (Fig. 1; more details in Section 2.1 and Methods Section 6.1). dLDS allows for locally linear, nonstationary dynamics that can be active simultaneously in different continuous ratios, and that can transition continuously without such hierarchical state constraints [42]. The coefficients that describe the time-varying utilization of these dynamics can show differences between instances of a behavior within a trial while still containing decodable information about behavior-related latent neural processes. The dynamics operators (DOs) themselves describe relationships between latent neural processes that can be mapped back to functional relationships between neurons. Furthermore, these operators can be linearly recombined in different ratios, as regulated by a set of coefficients that capture when different dynamics turn on and to what extent, enabling the modeling of gradual transitions in neural dynamic connectivity throughout behavior.

**Figure 1:**
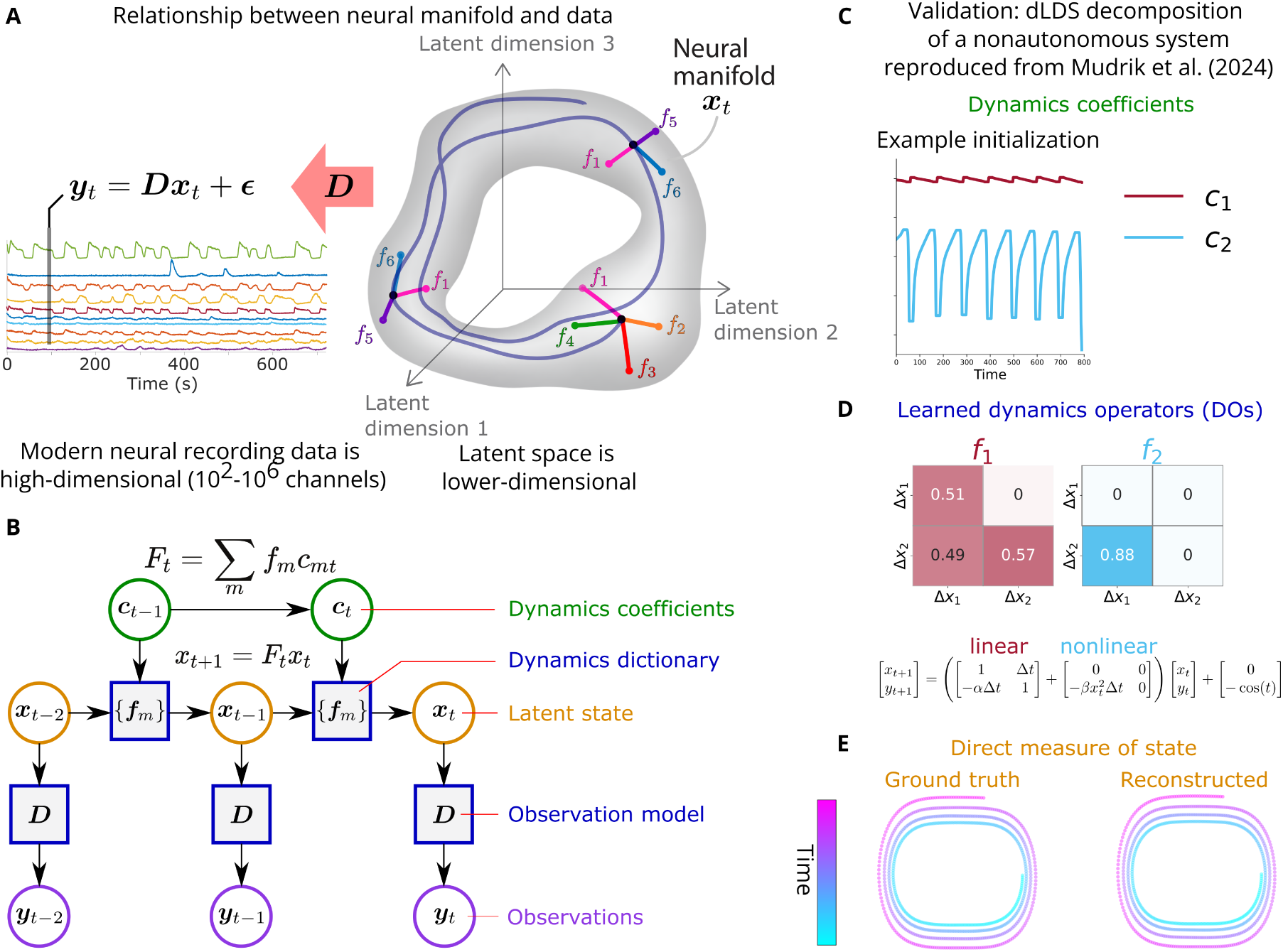
Modeling latent manifolds from neuroimaging data. **A:** Recorded neural activity ***y*** relates to latent neural states on the manifold ***x*** via observation model (matrix) ***D***. Neural dynamics (trajectories) on the manifold operate in the tangent space. These linear operators can be summed linearly. The manifold depicted here is an illustration; we do not constrain the model to a particular manifold, nor do we define a particular manifold in the process of learning the dLDS model. **B:** Graphical model depicting the relationships between data ***y*** and model parameters from time point to time point (dynamics coefficients ***c***, DO dictionary of matrices ***f***, latent states ***x***, and observation model ***D***). Dynamics coefficients ***c*** describe DO use over time. **C-E:** Example of a dLDS decomposition of a nonautonomous dynamical system - the Duffing oscillator, reproduced from [42]. dLDS can recover the activity of a simple nonautonomous system (**E**) using dynamics coefficients (**C**) dynamics operators (**D**) that represent its linear and nonlinear components.

In addition, we present an across-worm aligned^1^ extension of dLDS. Worms may not only exhibit individuality in their behavior and neural encodings of behavior and internal states, but also have slightly different recorded neurons, trial durations, experimental conditions, and frequencies, durations, and qualities of performed behaviors. In order to identify shared vs. individual mechanisms of neural signaling across worms, it is necessary to have a common framework or space of operators for comparing them.

Using dLDS models learned for individual *C. elegans* (per-worm) models and the across-worm aligned model extension we develop here, we discover the dynamic (i.e., nonstationary, anatomy-unconstrained functional) connectivity maps from learned DOs. These maps represent dynamic connections beyond anatomy or one-to-one direct stimulation-induced stationary functional connections and reveal key nonstationary properties of neurons in *C. elegans*. For one, the dynamic connectivity maps reveal how stereotyped discrete behavior has a neural basis that, while patterned, is also variable across repetitions of the same behavior and across individuals. We further show that our model’s decomposition of the dynamics underlying these maps disentangles concurrent processes within a trial, such as adaptation to varying aversive oxygen levels while performing a repetitive behavior.

This variability in latent neural dynamics underlying discrete behavioral states can highlight functional networks related to latent neural processes that can be otherwise difficult to observe, isolate, and test. These difficulties arise because such circuits can be the result of internal processes separate from the measured behavior and environmental variables, or because they reflect those variables indirectly, e.g., in the form of intermediary computations bridging sensing and behavior. In other words, even when activity levels of a neuron look similar across behaviors, the dynamic connectivity of those neurons with the rest of the network differs, indicating new roles for neurons with and without known functions. We are thus able to characterize the dynamic roles of interneurons such as AVFR, RIS, and RIGL, which show evidence of different dynamic encodings when modulated strongly by sensory vs. motor information, as well as the sensory BAG neurons, the motor neuron RMED, and the interneuron/motor neuron SABD, which strongly reflect both behavior and stimulation state and the interactions between their levels. These results complement and extend the extensive study of stationary properties of neurons in *C. elegans*, bringing a new analysis of their nonstationary properties.

## 2. Results

### 2.1 dLDS model architecture

Our analysis builds on the recently developed decomposed linear dynamical systems (dLDS) formulation [42], which is a dynamical systems model whose learned parameters reflect latent, concurrent neural processes through a generative model. In dLDS, the observed neural fluorescence ***y****_t_* at time *t* is modeled by latent states ***x****_t_* through a linear generative model, i.e.,

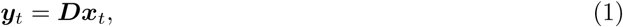

where ***D*** describes to what extent each latent dimension represents each recorded trace. Specifically, each column of ***D*** contains a consistantly co-active group of neurons representative of one latent dimension (a single entry of ***x****_t_*). In dLDS, we assume a constant latent state representation ***D*** for each individual worm. dLDS then models the brain dynamics as a dynamical system over the latent state ***x****_t_*.

Two key aspects drive the form of the dynamical systems model in dLDS: the need to capture concurrent processes, and the need to capture time-varying changes to the network interations. The former aims to model the fact that independent systems, such as sensory and motor networks, can be independently recruited. The latter aims to model adaptation via neuromodulation as an implicit change in the network behavior. To capture these effects, dLDS models the trajectory of the latent state as a time-varying linear dynamical system, i.e.,

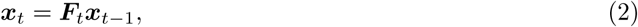

where the time varying linear system ***F****_t_* is modeled as a linear combination of stereotyped interconnection patterns that drive the full dynamics. Specifically,

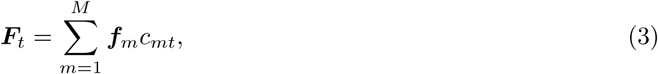

where the set of *M* dynamics operators {***f****_m_*}*_m_*_=1_*_,…,M_* are system properties (i.e., patterns of interconnection between neurons that are reused) and the coefficients *c_mt_*represent when different interconnections drive the neural dynamics at each time *t* (i.e., when *c_mt_* = 0) and how much influence they have. To induce greater interpretability and independence in terms of the use of different interconnectivity patterns over time, we further assume that the *c_mt_*coefficients are mostly zero at each time-step, i.e., the temporally local dynamics are most heavily driven by only few connectivity patterns. We refer to the set of {***f****_m_*}*_m_*_=1_*_,…,M_* operators as dynamics operators (DOs) and *c_mt_* = 0 as dynamics coefficients.

In other words, the vocabulary of the dLDS model consists of DOs, and the grammar that regulates how they work together to generate latent neural states consists of the dynamics coefficients. Then, we use ***D*** to translate from the language of latent states to the language of observable neural activity.

We fit the DOs in a data-driven way through an expectation maximization algorithm that approximates the Maximum Likelihood of the data given the DOs {***f****_m_*}*_m_*_=1_*_,…,M_* and the latent state representation ***D***. The learning procedure strongly resembles an iterative dictionary learning optimization which iterates between updating estimates for the unknown variables ***x****_t_* and ***c****_t_*, and updating the estimates for the model parameters {***f****_m_*}*_m_*_=1_*_,…,M_* and ***D*** (for more details, see Section 6.1). Experimental model reconstruction performance and parameters are explained in detail in Tables 1 and 2.

**Table 1:** dLDS performance in terms of *R*^2^.

| Model type | Protocol | Worm number | $R^2$ |
| --- | --- | --- | --- |
| Per-worm | Stim | 1 | 0.86 |
|  |  | 2 | 0.78 |
|  |  | 3 | 0.83 |
|  |  | 4 | 0.79 |
|  |  | 5 | 0.79 |
|  |  | 6 | 0.81 |
|  |  | 7 | 0.7 |
| Aligned | Stim | 1 | 0.87 |
|  |  | 2 | 0.84 |
|  |  | 3 | 0.85 |
|  |  | 4 | 0.82 |
|  |  | 5 | 0.75 |
|  |  | 6 | 0.8 |
|  |  | 7 | 0.83 |
| Per-worm | NoStim | 1 | 0.85 |
|  |  | 2 | 0.86 |
|  |  | 3 | 0.81 |
|  |  | 4 | 0.85 |
|  |  | 5 | 0.77 |

**Table 2:**
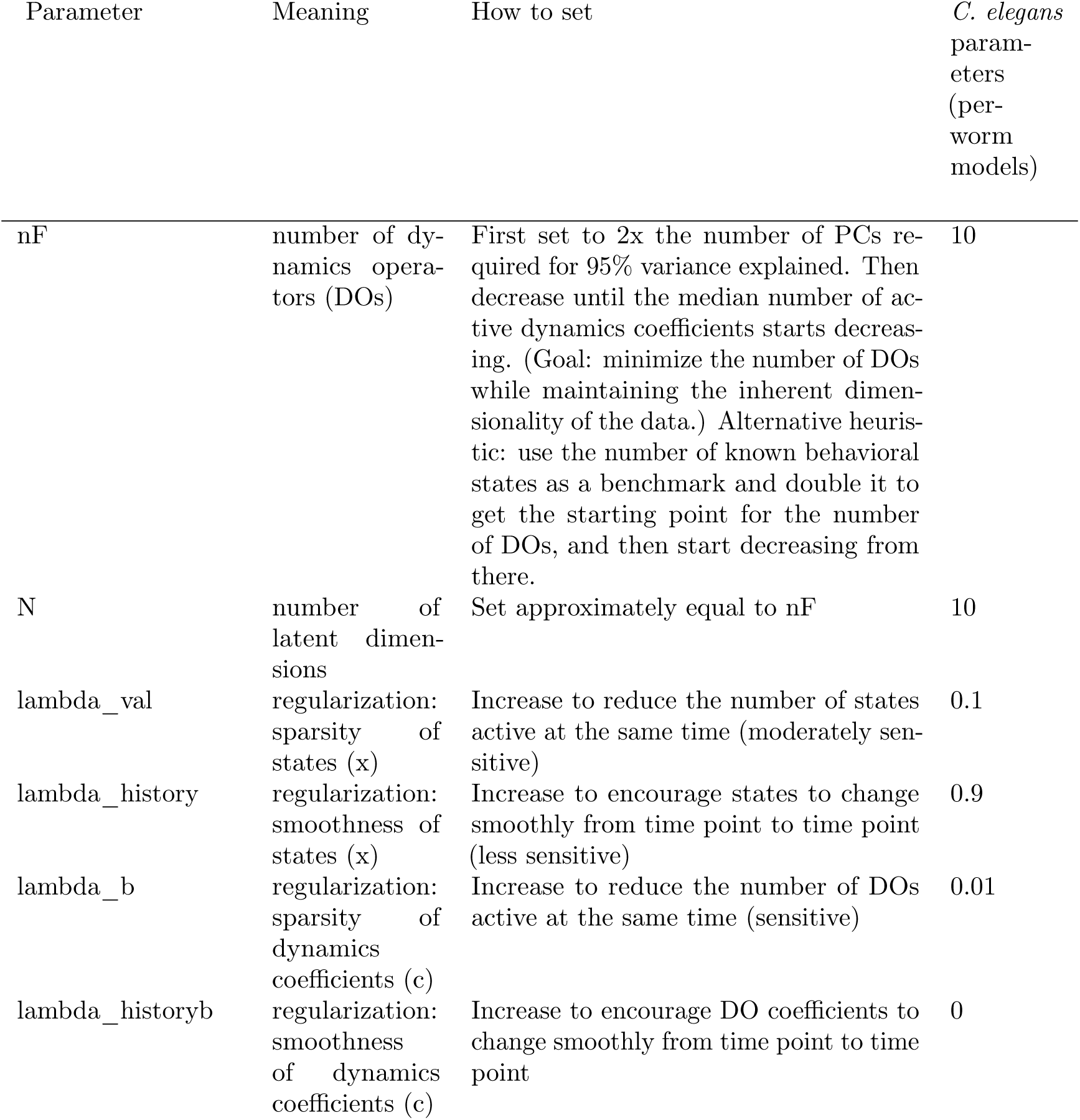
Meaning of key dLDS parameters, heuristics for setting them, and settings used for *C. elegans* models.

### *Validation:* dLDS characterizes the dynamics of nonautonomous dynamical systems

To validate dLDS, we demonstrate its ability to capture nonlinear behavior on a simulated dynamical system modulated by an external driving force. Specifically, we tested dLDS on the Duffing oscillator, a classic example of a nonlinear, nonautonomous dynamical system. The Duffing oscillator is characterized by the second-order differential equation,

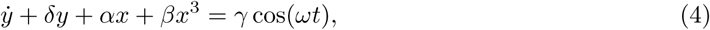

which generates a dynamical system that can be both damped and driven. In this version of the equation, 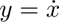, which leads to a discrete-time matrix version of the Duffing equation as

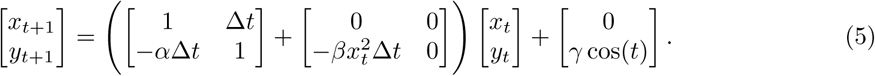

Our prior work [42] demonstrated that, across initializations, dLDS can recover a Duffing oscillator’s behavior and coefficients. Specifically, the dynamics operators learned by dLDS separated out the linear (DO 1) and nonlinear (DO 2) components of the discrete-time version of the Duffing equation (e.g., an example reproduced from [42] for *γ* = −1 is shown in Fig. 1) Thus, dLDS recovers a meaningful decomposition of this nonautonomous dynamical system. Additional demonstrations of dLDS on Lorenz attractors, Fitz Hugh Nagumo models, and other examples can be found in Mudrik *et al.* 2024 [42].

### 2.2 Assessment and validation of dLDS fits to *C. elegans* dynamics

We applied the dLDS model to *C. elegans* calcium imaging data from Kato *et al.* 2015 [3]. This dataset was collected from 12 worms (Table 3). Five of the worms experienced hyperoxia for 18 minutes (NoStim trials) to spontaneously elicit a pirouetting behavior, i.e., a sequence of forward crawling, turns, and reverse crawling. Seven of the worms were exposed to hyperoxia (21% oxygen) for the first half of the experiment, and alternating bouts of hypo- and hyperoxia (4% vs. 21% oxygen) for the second half in order to entrain the pirouettes to the oxygen levels (Stim trials). Approximately 100 calcium traces, imaged at 3 Hz, were identified from the imaging data per worm. Of these traces, only approximately 40 were labeled as originating from one or more known neurons, some with multiple neurons per trace and with some neurons appearing as labels for multiple traces. Moreover, once the protocol was established, the calcium imaging recordings were actually performed in immobilized worms.

**Table 3:** *C. elegans* data characteristics.

| Condition | Worm | Duration<br>(s) | Time<br>points | Recorded<br>traces | Labelled with one<br>or more neurons | Oxygen level<br>switch times (s) | Behavior<br>states |
| --- | --- | --- | --- | --- | --- | --- | --- |
| Stim | 1 | 720 | 2198 | 107 | 45 | every 30 s<br>starting at<br>360 until<br>the end | 4 |
|  | 2 |  | 2017 | 122 | 44 |  |  |
|  | 3 |  | 2197 | 124 | 40 |  |  |
|  | 4 |  | 2018 | 134 | 55 |  |  |
|  | 5 |  | 2201 | 123 | 48 |  |  |
|  | 6 |  | 2017 | 151 | 44 |  |  |
|  | 7 |  | 2019 | 146 | 51 |  |  |
| NoStim | 1 | 1080 | 3137 | 109 | 34 | N/A | 8 |
|  | 2 |  | 3134 | 107 | 44 |  |  |
|  | 3 |  | 3059 | 131 | 33 |  |  |
|  | 4 |  | 3311 | 126 | 48 |  |  |
|  | 5 |  | 3021 | 129 | 49 |  |  |

Despite this limitation of immobilization [22], [3] also provided expert labels for the discrete fictive pirouetting behavioral states by inferring their behavior from the activity of key neurons. These were coarse-grained for the Stim experiments (1: ‘FWD’ forward crawling, 2: ‘REV’ reverse crawling, 3: ‘REVSUS’ sustained reverse crawling, 4: ‘TURN’ post reversal turn) and fine-grained for the NoStim experiments (1: ‘FWD’ forward crawling, 2: ‘SLOW’ forward slowing, 3: ‘DT’ dorsal post reversal turn, 4: ‘VT’ ventral post reversal turn, 5: ‘REV1’ reverse crawling, 6: ‘REV2’ reverse crawling, 7: ‘REVSUS’ sustained reverse crawling, ‘NOSTATE’ - ambiguous). We then applied dLDS to each individual worm’s normalized (dF/F) traces.

We fit the dLDS model to each worm’s neural activity, using 10 dynamical operators and 10 latent states (see Section 6.1). We first validate that dLDS learns meaningful representations of the *C. elegans* dynamics by comparing the temporal and spatial statistics to known properties.

We can make inferences about the features of systems that generate that neural activity, and therefore the behavior, using the interpretable locally-linear properties (i.e., that the dynamics at each time point and the latent generative model are linear) of the dLDS model. All neurally-generated variables should be encoded in neural activity, representing a noisy version of discrete behavioral states. Therefore, the learned model parameters should reflect quantitative information such as speed of movement or modulation in response to sensory stimuli, as well as other background cognitive processes, e.g., hunger, wakefulness, etc. We thus test dLDS’s capabilities to capture these simultaneously active neural processes and variables by characterizing the dynamics of *C. elegans* neural activity.

The dLDS DO coefficients ***c*** reflect the utilization of DOs to generate latent neural states. Qualitatively, we observed that, for each worm, the same coefficients appeared in similar patterns (“motifs”) during the same *C. elegans* behaviors. These combinations of latent processes were discrete in that similar sets of DOs appeared every time. However, the DOs were also continuous because they appeared in slightly different ratios, which changed gradually over time or shifted in response to environmental variables such as stimulation protocols without switching completely. We also successfully decoded behavior states from the dynamics coefficients (see t-SNE and Isomap clustering by behavior states and linear discriminant analysis classification by behavior states in Figure 2, Supplementary Figures 1, 2, and 3).

**Figure 2:**
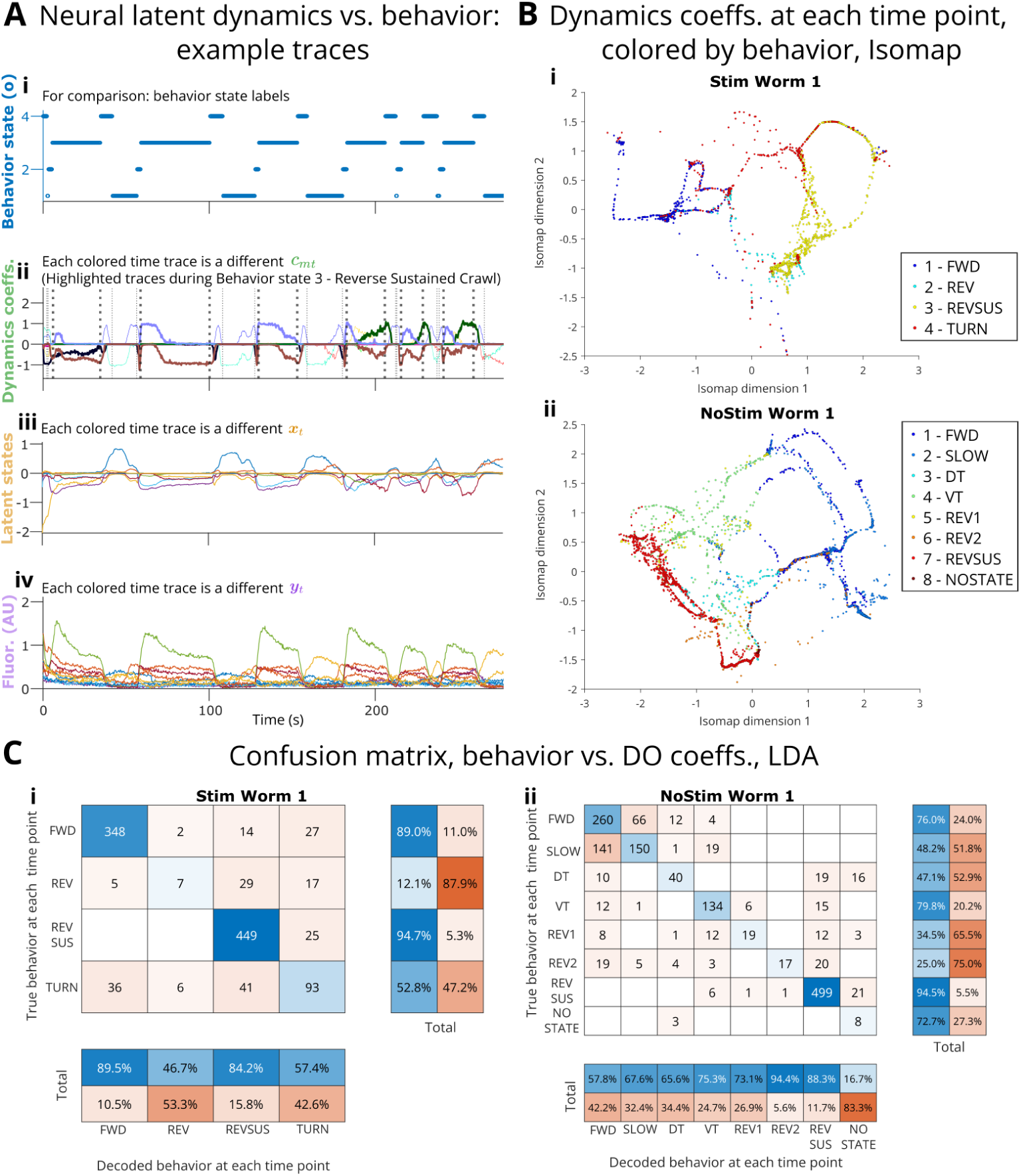
Model coefficients vs. behavior. **A:** Sample of time traces of behavior, dynamics (DO) coefficients, latent states, and fluorescence data. textbfB, C Behavior decoding from dynamics coefficients for Stim Worm 1 and NoStim Worm 1. Behavior labels: Stim 1 FWD forward crawling, 2 TURN post-reversal turn, 3 REV reverse crawling, 4 REVSUS sustained reverse crawling; NoStim 1 FWD forward crawling, 2 SLOW forward slowing, 3 DT dorsal post-reversal turn, 4 VT ventral post-reversal turn, 5 REV1 and 6 REV2 reverse crawling, 7 REVSUS reverse sustained crawling, 8 NOSTATE ambiguous. **B.i, B.ii:** Isomap finds a ring-like 2D embedding of the DO coefficients themselves, well clustered by behavior state (more examples in Supplementary Figs. 1, 2, 3). Each dot represents a time point in DO coefficient trace; see Section 6.1 and Table 3 for model parameter settings and counts. While biased toward the two behavior states with the highest base rates (1-FWD and 3-REVSUS in Stim, 1-FWD and 7-REVSUS in NoStim), linear discriminant analysis (LDA, **C.i, C.ii**) achieved high decoding performance on the DO coefficients.

To observe these behavior-related encodings in the DO coefficients ***c*** more quantitatively, we performed clustering analyses using t-distributed stochastic neighbor embedding (t-SNE) and Isomap, and classification vs. behavioral states using Support Vector Machine (SVM) models. Thus, from the dLDS DO utilization coefficients ***c*** and from their first derivative 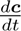, it was possible to decode the behavioral state labels inferred by Kato *et al.* 2015 [3] (Fig. 2 - Stim Worm 1 and NoStim Worm 1 shown in detail; Isomap for ***c*** only for all others, Supplementary Figs. 1, 2, 3)^2^. By clustering and coloring by behavioral states, t-SNE appears to show a continuum from forward to backward crawling (DO coefficients’ first derivative 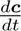), while Isomap embedding reveals a ring-like structure from state to state (with additional loops) in the coefficients and a spoke-like structure in the first derivative. Via linear discriminant analysis, it is also possible to classify at least the two major behavioral states from the DO coefficients alone (Fig. 2 - Stim Worm 1 and NoStim Worm 1). The observations above were similar when decoding from the latent states ***x***. In Sections 2.3, 2.4, and 6.3 and Supplementary Figures 4, 5, 6, and 7, we also investigated the behavioral and environmental context-dependent differences in DO utilization and what the DOs mean in the neural space.

### 2.3 *Novel capabilities:* dLDS identifies instantaneous dynamic connectivity between neurons

Having found evidence that the DO coefficients reflect behaviorally-relevant, continuous patterns in the neural activity, we then set out to quantify how the DOs themselves affect the neural activity and connectivity. Given that dLDS DOs transition smoothly (e.g., Fig. 3, Supplementary Figs. 8, 9, 10, 11, 12, 13), it is important to understand which neurons corresponding to those DOs are active and correlated at each time point in order to interpret the gradual effects of these DOs. This is even more important because these smooth transitions might reflect gradual changes in neural activity during or between behavioral states or other time-varying factors, such as oxygen levels. We created dynamic connectivity maps that can tie latent states and dynamics operators back to the neural space by simulating the effects of any linear combination of DOs at any time point (or the effects when applied to an impulse function of latent states). The dynamic connectivity maps essentially capture the correlational structure of an equivalent linear time-invariant system with the dynamics coefficients *c_mt_* frozen for all time (see Methods Section 6.4). By tracking how the dynamic connectivity maps change over time with the changes in *c_mt_*, we can track the dynamic networks active across all the recorded neurons.

**Figure 3:**
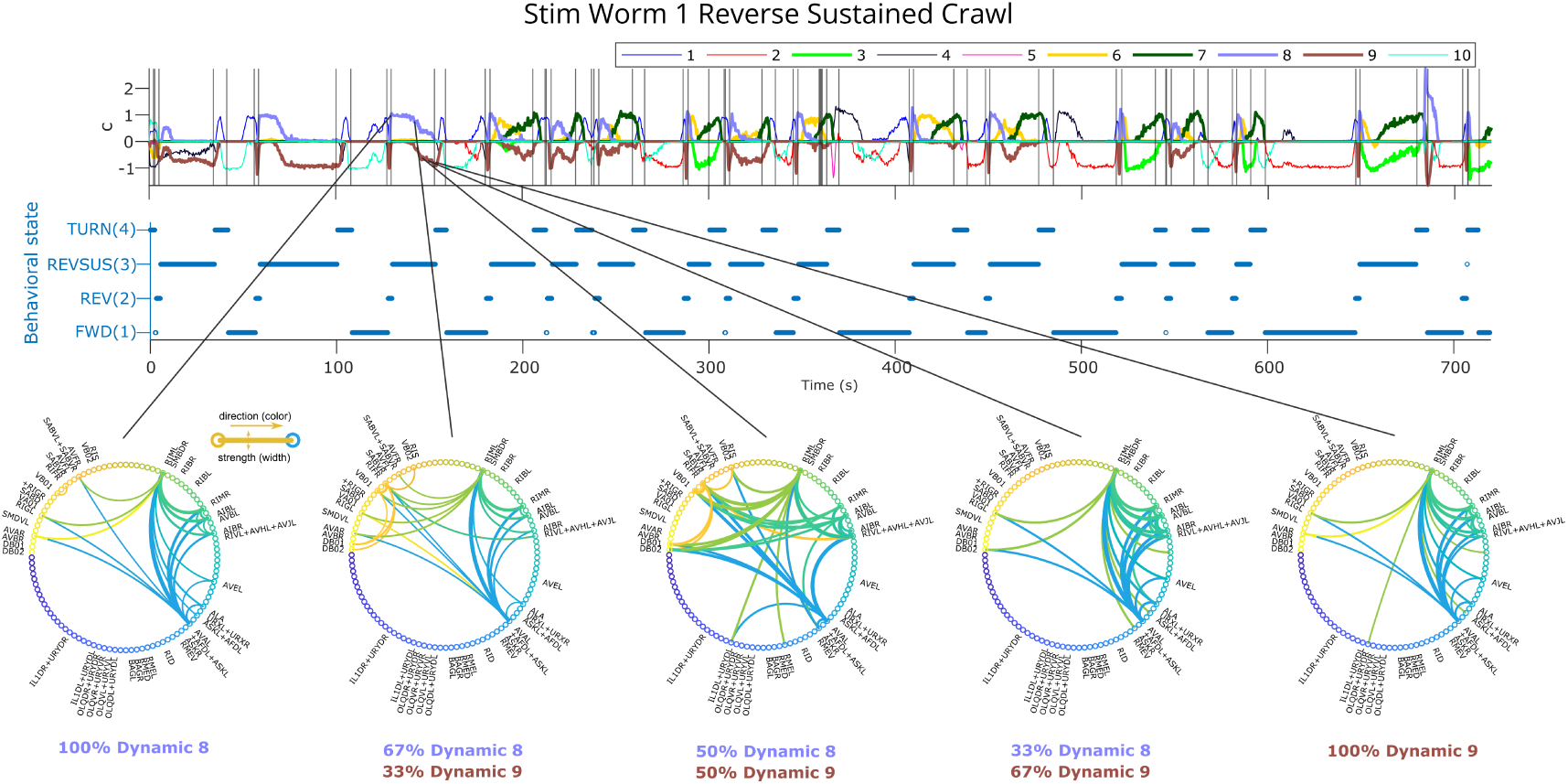
Continuous transition between DOs during behavior. DOs 8 and 9 in Stim Worm 1 are active during behavior state 3 (REVSUS). As DO 8 transitions to DO 9, connectivity between a core set of interneurons and the sensory neurons as well as motor neurons evolves in terms of which neurons are involved as well as in terms of strength and quantity. Here we show these two DOs linearly combined in different ratios, from 100% DO 8, to 67% DO 8 and 33% DO 9, all the way to 100% DO 9. We show the connection strengths (above a theshold) in the circle graphs in the right column, with a large percentage of neurons unlabelled.

As described in Section 2.2, the results abounded with occurrences of repeated motifs of simultaneously active DOs (often corresponding to behavior states) that appeared similarly but in slightly different ratios at different times throughout the trial and also gradually evolved during the course of a behavior state. In Figure 3, we show a generative example of the effects of different ratios of two DOs in combination. These two DOs appeared in combination during reverse sustained crawling in Stim Worm 1, but did not merely turn on when the reverse sustained crawling behavior started and turn off when it ended — their dynamics coefficients increased and decreased in magnitude gradually throughout the behavior. We see a small, gradual transition in which neurons participate most strongly as we move from one operator to another — for example, the SABV neurons and RIFR, cholinergic motor/interneurons [44], become less connected and less active. This suggests that while most of the circuit participating in this behavior stays constant throughout the behavior, gradual changes at the neuron interaction level might also occur throughout the course of the behavior. Such continuous changes to the latent dynamics throughout behavior states are perhaps needed to generate preparation to change to the next behavior or other internal processes.

### 2.4 *Biological findings:* Dynamics learned from dLDS help identify dynamic roles for interneurons and connectivity adaptation for sensory neurons

In order to study how different classes of neurons (sensory, motor, and interneurons) use dynamic encoding to generate behavior in a changing environment, we used dLDS to reconstruct individual neurons’ estimated activity and connectivity from time point to time point throughout the trial. Given the alternating hypoxia/hyperoxia oxygen stimulation protocol (second half of the Stim experiments), we investigated if and how some DO coefficients were different in the two halves of the Stim experiments, how the differently-utilized DOs might feature oxygen-sensing neurons in their corresponding dynamic connectivity maps, and how interneurons reflect behavior and stimulation.

We found that most of the neurons studied, in terms of activity and connectivity to and from the neuron, had a significant relationship with stimulation state, behavior state, and the interaction of the two. Thus, we observed that we could recover global behavior- or stimulation-related low-dimensional manifold states, similar to the global manifold gating effects reported by Fieseler *et al.* 2025 [36], i.e., there do appear to be some global brain states that strongly modulate the activity of a broad range of neurons. However, dLDS does not have to time-average across such brain states; we can observe instantaneous dynamics for each neuron and observe how its connectivity creates or responds to such brain states.

Digging deeper, each neuron class shows different patterns in the ways each uses its connectivity to fulfill its functions. First, connections to and from the oxygen-sensing BAG neurons (Fig. 4) were strongest during acute hypoxia and are fairly system-wide, which was to be expected given their known oxygen-sensing function. Importantly, these connections can adapt over time, using different neurons, not just less activity, as the worm habituates to hypoxia. Sensory neurons were also shown to be sensitive to motor information, which recent experimental work has shown is related to reafferent perception [25; 36].

**Figure 4:**
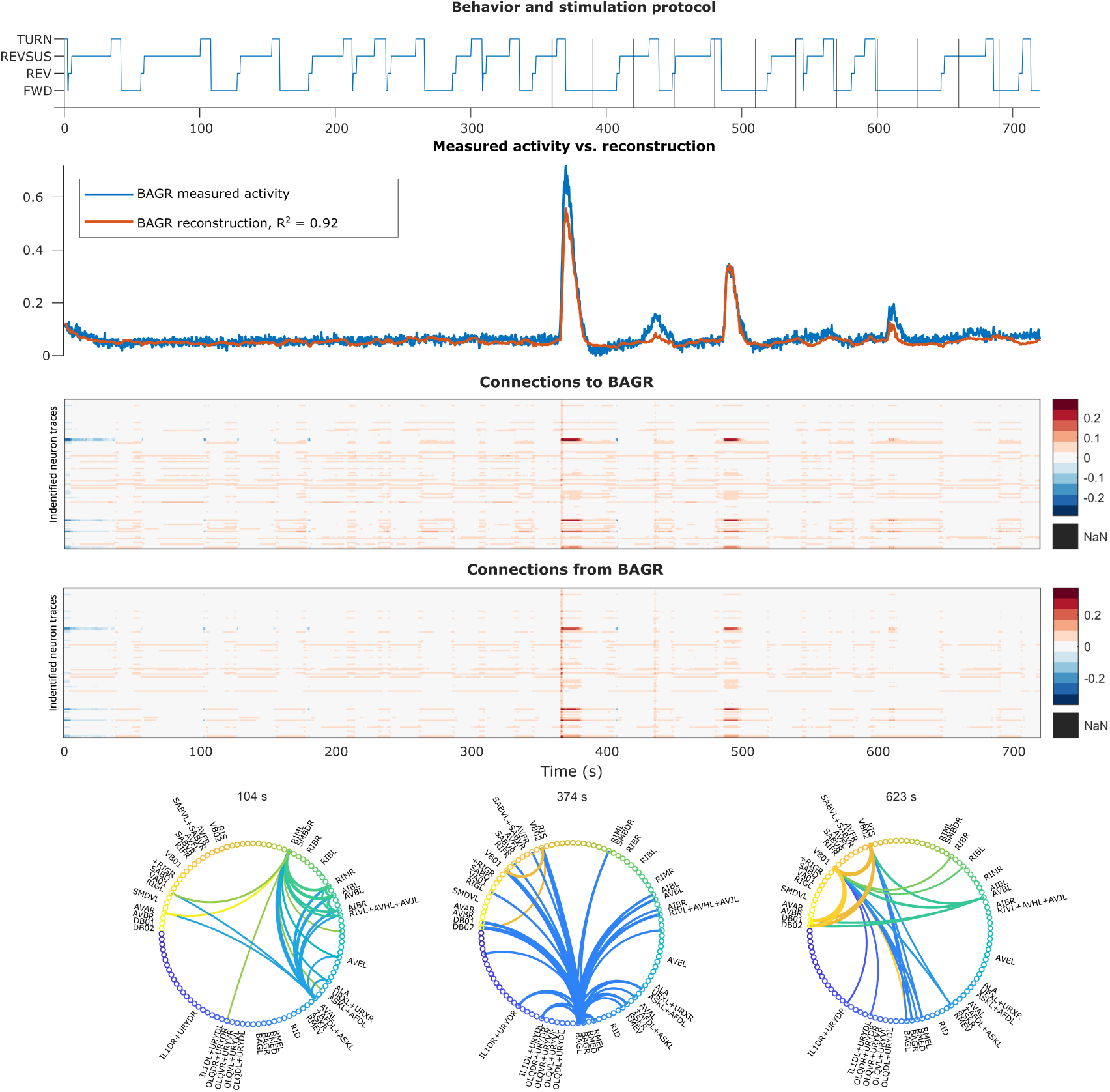
Reconstructed activity and dynamic connectivity to and from BAGR. **Top:** behavior state labels, with vertical lines indicating switches in the oxygen stimulation protocol. **Second from top:** measured BAGR activity vs. reconstructed BAGR activity (*y*^). **Third and fourth from top:** dynamic connectivity to vs. from BAGR at each time point. **Bottom row:** instantaneous dynamic connectivity maps at time points of strong BAGR connectivity before and during hypoxia. While BAGR sent out strong connections during the first bout of hypoxia, its connections shifted more strongly to motor/proprioceptive neurons in later bouts.

In comparison, connections to and from the RMED motor neuron (Fig. 5) were more stable throughout the experiment. However, these connections were also correlated both with bouts of pirouetting and with hypoxia. In fact, there were qualitatively different patterns and strengths of RMED inputs (“connectivity to”) depending on behavior state and hypoxia leading to RMED operating in two separate (positive vs. negative) regimes during most of the trial vs. during hypoxia. These context-dependent dynamics might even indicate a hierarchy between the standard pirouetting behavior and the hypoxia “alert” overriding it.

**Figure 5:**
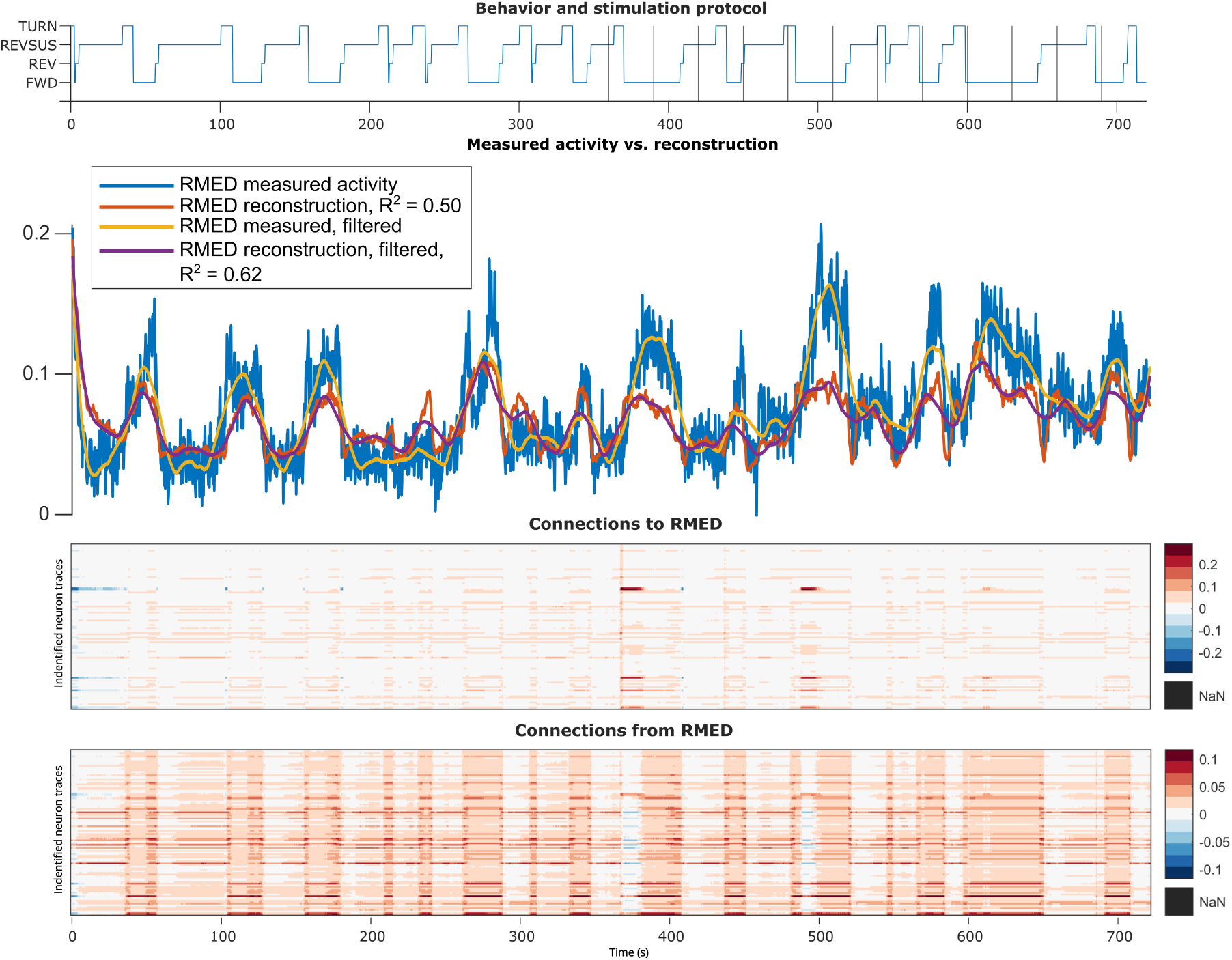
Stim Worm 1 RMED motor neuron reconstructed activity and connectivity.

Finally, we show an example interneuron, AVFR, whose function is less clear from the literature — unsurprisingly, it is more difficult to classify interneurons’ roles because they exist between sensory and motor neurons and seem to account for internal computation. While the noisy motor and interneurons were not as well-reconstructed as the BAG neurons, we saw a distinct pattern of behavior-dependent connectivity, interspersed with what seems to be an overriding hypoxia alert, as in the motor neuron. Importantly, the connectivity completely changed for short periods of time during hypoxia - the interneuron changed its connectivity, and thus its function, over time to relay information to different circuits, rather than always using the same mixed selectivity, irrespective of task variables. Thus, we can use dLDS dynamic connectivity to describe at each time step how even an interneuron’s role in its circuits is evolving.

#### 2.4.1 BAGL and BAGR neurons and their circuits

The BAG neurons are known to be involved in sensing increased carbon dioxide or decreased oxygen [44; 5; 45], making this pair of neurons a natural candidate for evaluating how sensory neurons can use their connectivity to adapt and habituate to changes in their environment. We put forward two possible hypotheses: all DOs might feature BAG relatively equally in order to make them all accommodate for the change in experimental conditions while creating the stereotyped behaviors, regardless of which half of the experiment they were in; or some DOs might feature BAG more strongly in a nonstationary way, possibly to account for hypoxia, separating out the underlying BAG-dependent circuits that are simultaneously active with the circuits that produce the stereotyped behaviors that persist throughout the experiment. This second hypothesis could not be explored with a switched linear dynamical system, where only one operator would represent each state at each time point.

What we found was even more nuanced and supports recent reports of the context-dependent function of BAG [25]. We observed that, while at the start of the alternating hypoxia/hyperoxia protocol (second half of the Stim experiments), there was a large spike in BAG activity, this was followed by smaller spikes. This led us to investigate whether the relationship between earlier and later BAG activity was merely a quantitative difference in *activity* of the same neurons, or whether habituation, and changes to *connectivity*, were leading to different responses to hypoxia over time. Using instantaneous dynamic connectivity, we quantitatively showed evidence of the latter. When hypoxia starts (around 360 s), BAG connectivity was strong to a large range of sensory, motor, and interneurons. Later in the trial, when hypoxia occurred again, connections from BAG were still present, but some of this sensory role shifted from external sensing (from BAG) to internal sensing (e.g., proprioceptive/motor neurons VB and DB).

Furthermore, based on the proportions of connectivity maps that most strongly featured the BAG neurons across the Stim worms, we found evidence that the BAG-featuring DOs were used in a nonstationary way throughout the experiments, potentially in relation to the alternating hypoxia/hyperoxia levels (Section 6.5, Table 4, Supplementary Figs. 14, 15, 16, 17, 18, 19, 20, 21, 22, 23, 24, 25, 26, 27, 28, 29, 30). We also observed that BAG-featuring DOs were only up to approximately half of the DOs active during the two major behavioral states (forward crawling and reverse sustained crawling), potentially with context-dependent bias toward one or the other major behavioral state. This indicates that there might be multiple interrelated but distinct circuits, some of which are BAG-related and some of which are more specific to generating the behaviors, that are active simultaneously, and that dLDS can separate them.

**Table 4:**
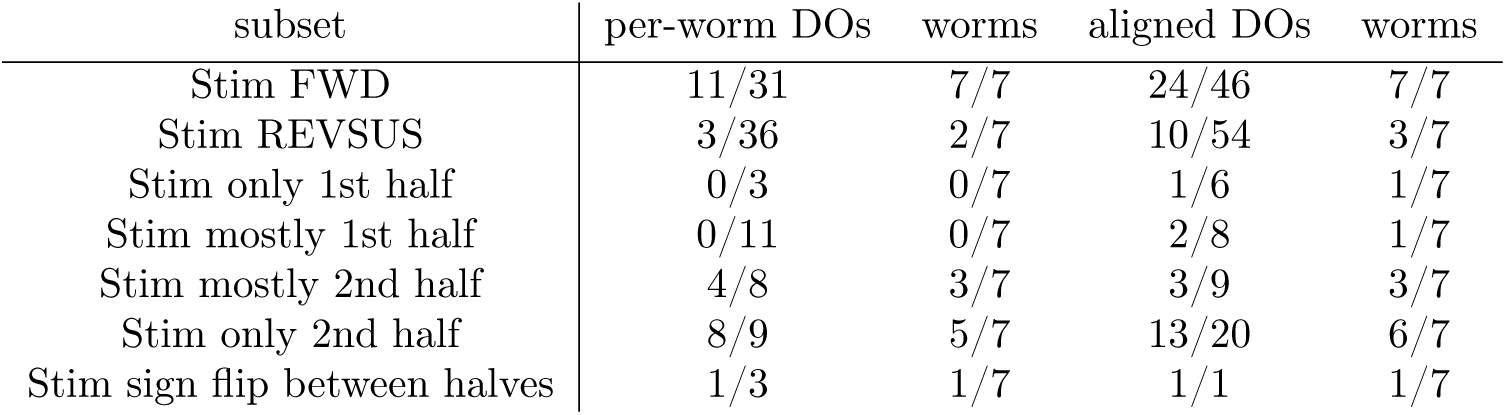
Tally of BAGR or BAGL presence in dynamic connectivity maps across conditions.

| subset | per-worm DOs | worms | aligned DOs | worms |
| --- | --- | --- | --- | --- |
| Stim FWD | 11/31 | 7/7 | 24/46 | 7/7 |
| Stim REVSUS | 3/36 | 2/7 | 10/54 | 3/7 |
| Stim only 1st half | 0/3 | 0/7 | 1/6 | 1/7 |
| Stim mostly 1st half | 0/11 | 0/7 | 2/8 | 1/7 |
| Stim mostly 2nd half | 4/8 | 3/7 | 3/9 | 3/7 |
| Stim only 2nd half | 8/9 | 5/7 | 13/20 | 6/7 |
| Stim sign flip between halves | 1/3 | 1/7 | 1/1 | 1/7 |

These results were supported by *χ*^2^, ANOVA, and MANOVA tests (Sections 5, 6, 7) on the reconstructed activity and connectivity, in a hierarchical way: at a low threshold for “active” or “connected”, the BAG neurons were more significantly influenced by behavior states, but at a high threshold, the stimulation protocol’s effects on their activity and connectivity were significant, which aligns with the visual patterns in Figure 4 and Supplementary Figure 31.

#### 2.4.2 RMED neuron

For comparison, we identified another forward crawling-biased neuron, RMED, a motor neuron with limited function documentation in the WormAtlas [44]. [46] proposed that the RME neurons regulate the amplitude of head-bending during locomotion. RMED-featuring DOs also showed some nonstationarity but were generally utilized in a more balanced way between halves of the experiment than the BAG neurons (Section 6.8, Table 8, Supplementary Figs. 14, 15, 16, 17, 18, 19, 20, 21, 22, 23, 24, 25, 26, 27, 28, 29, 30).

To further complicate the story, the *χ*^2^ tests of RMED only showed significant relationships between the connectivity to and from RMED and stimulation state or behavior, even though these are clear visually (Fig. 5 - Stim Worm 1 as an example), but they did show a consistently significant relationship between RMED reconstructed activity and both stimulation and behavior. As with BAGL and BAGR, the ANOVA results showed significant effects of behavior and usually significant effects of the stimulation setting. A possible mechanistic interpretation of the significant stimulation effects is that RMED appears to play a more direct role in sensorimotor circuits than otherwise previously indicated, and plays a different role during strong sensory signaling. Moreover, this degree of variability in terms of how RMED contributes to generating behavior could be another example of worm individuality, as shown in Figure 1D in [14].

#### 2.4.3 Other neurons for comparison: AVFR, RIGL, RIS, SABD

We selected four interneurons—AVFR, RIGL, RIS, and SABD—for further analysis based both on their limited function documentation in the Worm Atlas [44] and on the consistency with which they were uniquely and distinctly labelled across the 7 worms (AVFR in all 7, RIS in worms 1-6, RIGL in worms 2,3,4, and 6, and SABD in worm 1; See Fig. 6 and Supplementary Figs. 32, 33 for the activity and connectivity of the Stim Worm 1 neurons). Through an ANOVA analysis, we observed significant effects of oxygen stimulation on AVFR, RIS, and RIGL reconstructed activity in most of the worms (Table 6) but persistently significant effects on connectivity across all of the worms (Table 7).

**Figure 6:**
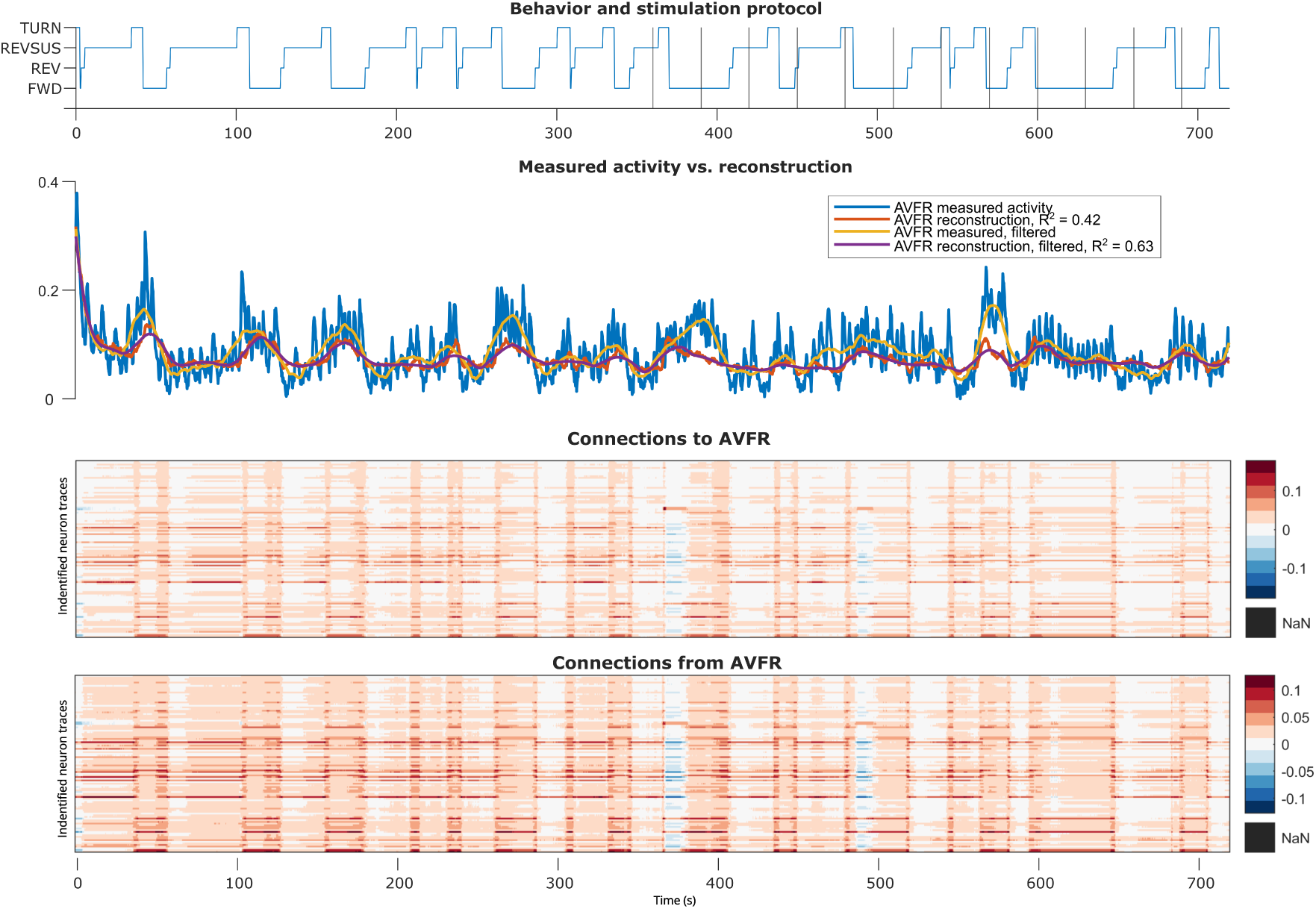
Stim Worm 1 AVFR interneuron reconstructed activity and connectivity.

**Table 5:** Two-sided *χ*^2^ tests for BAG and RMED activity and connectivity (inactive vs. active, defined by a threshold), behavior, and oxygen stimulation levels. Glossary: df: degrees of freedom, Q: chi squared test statistic, bhv: behavior, [neuron]connFrom: strongest connectivity value (absolute value) from that neuron, [neuron]connTo: strongest connectivity value to that neuron,[Neuron]yhat: reconstructed activity of that neuron, []3: data included only above threshold 3 standard deviations above median, []Pos: only positive connections included, []Neg: only negative connections included, FWD: forward crawling (value= 1) vs. all other behaviors, stimTime: first half vs. second half (value= 1), stimTimesDetailed: 21% oxygen vs. 4% (value= 1).

| Stim worm | variable 1 | variable 2 | p value | max level | df | Q |
| --- | --- | --- | --- | --- | --- | --- |
| 1 | bhv | AVFRconnFrom | 0.153422 | 3 | 3 | 5.264503 |
| 2 | bhv | AVFRconnFrom | 0.852213 | 1 | 3 | 0.788512 |
| 3 | bhv | AVFRconnFrom | 0.86671 | 1 | 3 | 0.727532 |
| 4 | bhv | AVFRconnFrom | 0.007016 | 3 | 3 | 12.1099 |
| 5 | bhv | AVFRconnFrom | 0.004201 | 3 | 3 | 13.21129 |
| 6 | bhv | AVFRconnFrom | 0.821141 | 3 | 3 | 0.917755 |
| 7 | bhv | AVFRconnFrom | 0.760397 | 3 | 3 | 1.169215 |
| 1 | FWD | AVFRconnFrom | 0.455954 | 2 | 1 | 0.555808 |
| 2 | FWD | AVFRconnFrom | 0.374551 | 1 | 1 | 0.788512 |
| 3 | FWD | AVFRconnFrom | 0.393684 | 1 | 1 | 0.727532 |
| 4 | FWD | AVFRconnFrom | 0.521763 | 2 | 1 | 0.410406 |
| 5 | FWD | AVFRconnFrom | 0.104907 | 2 | 1 | 2.629308 |
| 6 | FWD | AVFRconnFrom | 0.552443 | 2 | 1 | 0.352958 |
| 7 | FWD | AVFRconnFrom | 0.390819 | 2 | 1 | 0.736392 |
| 1 | Stim | AVFRconnFrom | 0.3172 | 2 | 1 | 1.000455 |
| 2 | Stim | AVFRconnFrom | 0.316951 | 2 | 1 | 1.001489 |
| 3 | Stim | AVFRconnFrom | 0.31698 | 2 | 1 | 1.001367 |
| 4 | Stim | AVFRconnFrom | 0.317191 | 2 | 1 | 1.000496 |
| 5 | Stim | AVFRconnFrom | 0.045206 | 2 | 1 | 4.010926 |
| 6 | Stim | AVFRconnFrom | 0.316951 | 2 | 1 | 1.001489 |
| 7 | Stim | AVFRconnFrom | 0.31743 | 1 | 1 | 0.999505 |
| 1 | StimDetailed | AVFRconnFrom | 0.563378 | 1 | 1 | 0.33389 |
| 2 | StimDetailed | AVFRconnFrom | 0.56322 | 1 | 1 | 0.33416 |
| 3 | StimDetailed | AVFRconnFrom | 0.56468 | 1 | 1 | 0.331666 |
| 4 | StimDetailed | AVFRconnFrom | 0.563349 | 1 | 1 | 0.333939 |
| 5 | StimDetailed | AVFRconnFrom | 0.247926 | 1 | 1 | 1.334952 |
| 6 | StimDetailed | AVFRconnFrom | 0.56322 | 1 | 1 | 0.33416 |
| 7 | StimDetailed | AVFRconnFrom | 0.563477 | 1 | 1 | 0.333719 |
| 1 | bhv | AVFRconnFrom3 | 0.002175 | 3 | 3 | 14.61688 |
| 2 | bhv | AVFRconnFrom3 | 1.53E-09 | 1 | 3 | 43.96989 |
| 3 | bhv | AVFRconnFrom3 | 6.80E-05 | 1 | 3 | 21.91259 |
| 4 | bhv | AVFRconnFrom3 | 0 | 3 | 3 | 233.665 |
| 5 | bhv | AVFRconnFrom3 | 5.67E-09 | 3 | 3 | 41.29154 |
| 6 | bhv | AVFRconnFrom3 | 0 | 3 | 3 | 164.9716 |
| 7 | bhv | AVFRconnFrom3 | 2.91E-09 | 3 | 3 | 42.6603 |
| 1 | FWD | AVFRconnFrom3 | 0.004727 | 2 | 1 | 7.981118 |
| 2 | FWD | AVFRconnFrom3 | 1.30E-09 | 1 | 1 | 36.81344 |
| 3 | FWD | AVFRconnFrom3 | 0.182261 | 1 | 1 | 1.779094 |
| 4 | FWD | AVFRconnFrom3 | 0 | 2 | 1 | 123.5084 |
| 5 | FWD | AVFRconnFrom3 | 0.004898 | 2 | 1 | 7.916752 |
| 6 | FWD | AVFRconnFrom3 | 0 | 2 | 1 | 84.89804 |
| 7 | FWD | AVFRconnFrom3 | 1.27E-07 | 2 | 1 | 27.90499 |
| 1 | Stim | AVFRconnFrom3 | 2.68E-13 | 1 | 1 | 53.43277 |
| 2 | Stim | AVFRconnFrom3 | 1.57E-13 | 2 | 1 | 54.48424 |
| 3 | Stim | AVFRconnFrom3 | 0.999273 | 2 | 1 | 8.30E-07 |
| 4 | Stim | AVFRconnFrom3 | 0 | 1 | 1 | 117.9731 |
| 5 | Stim | AVFRconnFrom3 | 0.000511 | 2 | 1 | 12.07675 |
| 6 | Stim | AVFRconnFrom3 | 0 | 1 | 1 | 235.9593 |
| 7 | Stim | AVFRconnFrom3 | 7.01E-12 | 2 | 1 | 47.02587 |
| 1 | StimDetailed | AVFRconnFrom3 | 1.19E-05 | 1 | 1 | 19.17793 |
| 2 | StimDetailed | AVFRconnFrom3 | 2.01E-05 | 1 | 1 | 18.17941 |
| 3 | StimDetailed | AVFRconnFrom3 | 0.996217 | 1 | 1 | 2.25E-05 |
| 4 | StimDetailed | AVFRconnFrom3 | 2.73E-05 | 1 | 1 | 17.59624 |
| 5 | StimDetailed | AVFRconnFrom3 | 0.044977 | 1 | 1 | 4.019492 |
| 6 | StimDetailed | AVFRconnFrom3 | 2.22E-16 | 1 | 1 | 67.06295 |
| 7 | StimDetailed | AVFRconnFrom3 | 7.42E-05 | 1 | 1 | 15.70119 |
| 1 | bhv | AVFRconnFromNeg | 0 | 1 | 3 | 99.65838 |
| 2 | bhv | AVFRconnFromNeg | NaN | 1 | 3 | NaN |
| 3 | bhv | AVFRconnFromNeg | NaN | 1 | 3 | NaN |
| 4 | bhv | AVFRconnFromNeg | NaN | 1 | 3 | NaN |
| 5 | bhv | AVFRconnFromNeg | 1.41E-07 | 4 | 3 | 34.70232 |
| 6 | bhv | AVFRconnFromNeg | NaN | 1 | 3 | NaN |
| 7 | bhv | AVFRconnFromNeg | NaN | 1 | 3 | NaN |
| 1 | FWD | AVFRconnFromNeg | 0 | 1 | 1 | 75.76626 |
| 2 | FWD | AVFRconnFromNeg | NaN | 1 | 1 | NaN |
| 3 | FWD | AVFRconnFromNeg | NaN | 1 | 1 | NaN |
| 4 | FWD | AVFRconnFromNeg | NaN | 1 | 1 | NaN |
| 5 | FWD | AVFRconnFromNeg | 0.007082 | 2 | 1 | 7.252064 |
| 6 | FWD | AVFRconnFromNeg | NaN | 1 | 1 | NaN |
| 7 | FWD | AVFRconnFromNeg | NaN | 1 | 1 | NaN |
| 1 | Stim | AVFRconnFromNeg | 1.54E-10 | 2 | 1 | 40.98106 |
| 2 | Stim | AVFRconnFromNeg | NaN | 1 | 1 | NaN |
| 3 | Stim | AVFRconnFromNeg | NaN | 1 | 1 | NaN |
| 4 | Stim | AVFRconnFromNeg | NaN | 1 | 1 | NaN |
| 5 | Stim | AVFRconnFromNeg | 2.02E-08 | 1 | 1 | 31.47144 |
| 6 | Stim | AVFRconnFromNeg | NaN | 1 | 1 | NaN |
| 7 | Stim | AVFRconnFromNeg | NaN | 1 | 1 | NaN |
| 5 | StimDetailed | AVFRconnFromNeg | 0.00121 | 1 | 1 | 10.4746 |
| 7 | StimDetailed | AVFRconnFromNeg | NaN | 1 | 1 | NaN |
| 1 | bhv | AVFRconnFromNeg3 | 8.88E-16 | 1 | 3 | 73.21871 |
| 2 | bhv | AVFRconnFromNeg3 | NaN | 1 | 3 | NaN |
| 3 | bhv | AVFRconnFromNeg3 | NaN | 1 | 3 | NaN |
| 4 | bhv | AVFRconnFromNeg3 | NaN | 1 | 3 | NaN |
| 5 | bhv | AVFRconnFromNeg3 | NaN | 1 | 3 | NaN |
| 6 | bhv | AVFRconnFromNeg3 | NaN | 1 | 3 | NaN |
| 7 | bhv | AVFRconnFromNeg3 | NaN | 1 | 3 | NaN |
| 1 | FWD | AVFRconnFromNeg3 | 1.67E-15 | 1 | 1 | 63.47413 |
| 2 | FWD | AVFRconnFromNeg3 | NaN | 1 | 1 | NaN |
| 3 | FWD | AVFRconnFromNeg3 | NaN | 1 | 1 | NaN |
| 4 | FWD | AVFRconnFromNeg3 | NaN | 1 | 1 | NaN |
| 5 | FWD | AVFRconnFromNeg3 | NaN | 1 | 1 | NaN |
| 6 | FWD | AVFRconnFromNeg3 | NaN | 1 | 1 | NaN |
| 7 | FWD | AVFRconnFromNeg3 | NaN | 1 | 1 | NaN |
| 1 | Stim | AVFRconnFromNeg3 | 1.62E-10 | 2 | 1 | 40.88402 |
| 2 | Stim | AVFRconnFromNeg3 | NaN | 1 | 1 | NaN |
| 3 | Stim | AVFRconnFromNeg3 | NaN | 1 | 1 | NaN |
| 4 | Stim | AVFRconnFromNeg3 | NaN | 1 | 1 | NaN |
| 5 | Stim | AVFRconnFromNeg3 | NaN | 1 | 1 | NaN |
| 6 | Stim | AVFRconnFromNeg3 | NaN | 1 | 1 | NaN |
| 7 | Stim | AVFRconnFromNeg3 | NaN | 1 | 1 | NaN |
| 5 | StimDetailed | AVFRconnFromNeg3 | NaN | 1 | 1 | NaN |
| 7 | StimDetailed | AVFRconnFromNeg3 | NaN | 1 | 1 | NaN |
| 1 | bhv | AVFRconnFromPos | 0.153422 | 3 | 3 | 5.264503 |
| 2 | bhv | AVFRconnFromPos | 0.852213 | 1 | 3 | 0.788512 |
| 3 | bhv | AVFRconnFromPos | 0.86671 | 1 | 3 | 0.727532 |
| 4 | bhv | AVFRconnFromPos | 0.007016 | 3 | 3 | 12.1099 |
| 5 | bhv | AVFRconnFromPos | 0.004201 | 3 | 3 | 13.21129 |
| 6 | bhv | AVFRconnFromPos | 0.821141 | 3 | 3 | 0.917755 |
| 7 | bhv | AVFRconnFromPos | 0.760397 | 3 | 3 | 1.169215 |
| 1 | FWD | AVFRconnFromPos | 0.455954 | 2 | 1 | 0.555808 |
| 2 | FWD | AVFRconnFromPos | 0.374551 | 1 | 1 | 0.788512 |
| 3 | FWD | AVFRconnFromPos | 0.393684 | 1 | 1 | 0.727532 |
| 4 | FWD | AVFRconnFromPos | 0.521763 | 2 | 1 | 0.410406 |
| 5 | FWD | AVFRconnFromPos | 0.104907 | 2 | 1 | 2.629308 |
| 6 | FWD | AVFRconnFromPos | 0.552443 | 2 | 1 | 0.352958 |
| 7 | FWD | AVFRconnFromPos | 0.390819 | 2 | 1 | 0.736392 |
| 1 | Stim | AVFRconnFromPos | 0.3172 | 2 | 1 | 1.000455 |
| 2 | Stim | AVFRconnFromPos | 0.316951 | 2 | 1 | 1.001489 |
| 3 | Stim | AVFRconnFromPos | 0.31698 | 2 | 1 | 1.001367 |
| 4 | Stim | AVFRconnFromPos | 0.317191 | 2 | 1 | 1.000496 |
| 5 | Stim | AVFRconnFromPos | 0.045206 | 2 | 1 | 4.010926 |
| 6 | Stim | AVFRconnFromPos | 0.316951 | 2 | 1 | 1.001489 |
| 7 | Stim | AVFRconnFromPos | 0.31743 | 1 | 1 | 0.999505 |
| 1 | StimDetailed | AVFRconnFromPos | 0.563378 | 1 | 1 | 0.33389 |
| 2 | StimDetailed | AVFRconnFromPos | 0.56322 | 1 | 1 | 0.33416 |
| 3 | StimDetailed | AVFRconnFromPos | 0.56468 | 1 | 1 | 0.331666 |
| 4 | StimDetailed | AVFRconnFromPos | 0.563349 | 1 | 1 | 0.333939 |
| 5 | StimDetailed | AVFRconnFromPos | 0.247926 | 1 | 1 | 1.334952 |
| 6 | StimDetailed | AVFRconnFromPos | 0.56322 | 1 | 1 | 0.33416 |
| 7 | StimDetailed | AVFRconnFromPos | 0.563477 | 1 | 1 | 0.333719 |
| 1 | bhv | AVFRconnFromPos3 | 0.028315 | 3 | 3 | 9.074636 |
| 2 | bhv | AVFRconnFromPos3 | 1.53E-09 | 1 | 3 | 43.96989 |
| 3 | bhv | AVFRconnFromPos3 | 6.80E-05 | 1 | 3 | 21.91259 |
| 4 | bhv | AVFRconnFromPos3 | 0 | 3 | 3 | 233.665 |
| 5 | bhv | AVFRconnFromPos3 | 5.67E-09 | 3 | 3 | 41.29154 |
| 6 | bhv | AVFRconnFromPos3 | 0 | 3 | 3 | 164.9716 |
| 7 | bhv | AVFRconnFromPos3 | 2.91E-09 | 3 | 3 | 42.6603 |
| 1 | FWD | AVFRconnFromPos3 | 0.687188 | 2 | 1 | 0.162147 |
| 2 | FWD | AVFRconnFromPos3 | 1.30E-09 | 1 | 1 | 36.81344 |
| 3 | FWD | AVFRconnFromPos3 | 0.182261 | 1 | 1 | 1.779094 |
| 4 | FWD | AVFRconnFromPos3 | 0 | 2 | 1 | 123.5084 |
| 5 | FWD | AVFRconnFromPos3 | 0.004898 | 2 | 1 | 7.916752 |
| 6 | FWD | AVFRconnFromPos3 | 0 | 2 | 1 | 84.89804 |
| 7 | FWD | AVFRconnFromPos3 | 1.27E-07 | 2 | 1 | 27.90499 |
| 1 | Stim | AVFRconnFromPos3 | 0 | 1 | 1 | 81.07336 |
| 2 | Stim | AVFRconnFromPos3 | 1.57E-13 | 2 | 1 | 54.48424 |
| 3 | Stim | AVFRconnFromPos3 | 0.999273 | 2 | 1 | 8.30E-07 |
| 4 | Stim | AVFRconnFromPos3 | 0 | 1 | 1 | 117.9731 |
| 5 | Stim | AVFRconnFromPos3 | 0.000511 | 2 | 1 | 12.07675 |
| 6 | Stim | AVFRconnFromPos3 | 0 | 1 | 1 | 235.9593 |
| 7 | Stim | AVFRconnFromPos3 | 7.01E-12 | 2 | 1 | 47.02587 |
| 1 | StimDetailed | AVFRconnFromPos3 | 0.154357 | 1 | 1 | 2.028655 |
| 2 | StimDetailed | AVFRconnFromPos3 | 2.01E-05 | 1 | 1 | 18.17941 |
| 3 | StimDetailed | AVFRconnFromPos3 | 0.996217 | 1 | 1 | 2.25E-05 |
| 4 | StimDetailed | AVFRconnFromPos3 | 2.73E-05 | 1 | 1 | 17.59624 |
| 5 | StimDetailed | AVFRconnFromPos3 | 0.044977 | 1 | 1 | 4.019492 |
| 6 | StimDetailed | AVFRconnFromPos3 | 2.22E-16 | 1 | 1 | 67.06295 |
| 7 | StimDetailed | AVFRconnFromPos3 | 7.42E-05 | 1 | 1 | 15.70119 |
| 1 | bhv | AVFRconnTo | 0.153422 | 3 | 3 | 5.264503 |
| 2 | bhv | AVFRconnTo | 0.852213 | 1 | 3 | 0.788512 |
| 3 | bhv | AVFRconnTo | 0.86671 | 1 | 3 | 0.727532 |
| 4 | bhv | AVFRconnTo | 0.007016 | 3 | 3 | 12.1099 |
| 5 | bhv | AVFRconnTo | 0.004201 | 3 | 3 | 13.21129 |
| 6 | bhv | AVFRconnTo | 0.821141 | 3 | 3 | 0.917755 |
| 7 | bhv | AVFRconnTo | 0.760397 | 3 | 3 | 1.169215 |
| 1 | FWD | AVFRconnTo | 0.455954 | 2 | 1 | 0.555808 |
| 2 | FWD | AVFRconnTo | 0.374551 | 1 | 1 | 0.788512 |
| 3 | FWD | AVFRconnTo | 0.393684 | 1 | 1 | 0.727532 |
| 4 | FWD | AVFRconnTo | 0.521763 | 2 | 1 | 0.410406 |
| 5 | FWD | AVFRconnTo | 0.104907 | 2 | 1 | 2.629308 |
| 6 | FWD | AVFRconnTo | 0.552443 | 2 | 1 | 0.352958 |
| 7 | FWD | AVFRconnTo | 0.390819 | 2 | 1 | 0.736392 |
| 1 | Stim | AVFRconnTo | 0.3172 | 2 | 1 | 1.000455 |
| 2 | Stim | AVFRconnTo | 0.316951 | 2 | 1 | 1.001489 |
| 3 | Stim | AVFRconnTo | 0.31698 | 2 | 1 | 1.001367 |
| 4 | Stim | AVFRconnTo | 0.317191 | 2 | 1 | 1.000496 |
| 5 | Stim | AVFRconnTo | 0.045206 | 2 | 1 | 4.010926 |
| 6 | Stim | AVFRconnTo | 0.316951 | 2 | 1 | 1.001489 |
| 7 | Stim | AVFRconnTo | 0.31743 | 1 | 1 | 0.999505 |
| 1 | StimDetailed | AVFRconnTo | 0.563378 | 1 | 1 | 0.33389 |
| 2 | StimDetailed | AVFRconnTo | 0.56322 | 1 | 1 | 0.33416 |
| 3 | StimDetailed | AVFRconnTo | 0.56468 | 1 | 1 | 0.331666 |
| 4 | StimDetailed | AVFRconnTo | 0.563349 | 1 | 1 | 0.333939 |
| 5 | StimDetailed | AVFRconnTo | 0.247926 | 1 | 1 | 1.334952 |
| 6 | StimDetailed | AVFRconnTo | 0.56322 | 1 | 1 | 0.33416 |
| 7 | StimDetailed | AVFRconnTo | 0.563477 | 1 | 1 | 0.333719 |
| 1 | bhv | AVFRconnTo3 | 0.003414 | 3 | 3 | 13.65503 |
| 2 | bhv | AVFRconnTo3 | 0.852213 | 1 | 3 | 0.788512 |
| 3 | bhv | AVFRconnTo3 | 0.86671 | 1 | 3 | 0.727532 |
| 4 | bhv | AVFRconnTo3 | 0.007016 | 3 | 3 | 12.1099 |
| 5 | bhv | AVFRconnTo3 | 0.004201 | 3 | 3 | 13.21129 |
| 6 | bhv | AVFRconnTo3 | 0.821141 | 3 | 3 | 0.917755 |
| 7 | bhv | AVFRconnTo3 | 0.760397 | 3 | 3 | 1.169215 |
| 1 | FWD | AVFRconnTo3 | 0.109545 | 2 | 1 | 2.560777 |
| 2 | FWD | AVFRconnTo3 | 0.374551 | 1 | 1 | 0.788512 |
| 3 | FWD | AVFRconnTo3 | 0.393684 | 1 | 1 | 0.727532 |
| 4 | FWD | AVFRconnTo3 | 0.521763 | 2 | 1 | 0.410406 |
| 5 | FWD | AVFRconnTo3 | 0.104907 | 2 | 1 | 2.629308 |
| 6 | FWD | AVFRconnTo3 | 0.552443 | 2 | 1 | 0.352958 |
| 7 | FWD | AVFRconnTo3 | 0.390819 | 2 | 1 | 0.736392 |
| 1 | Stim | AVFRconnTo3 | 1.88E-08 | 1 | 1 | 31.61775 |
| 2 | Stim | AVFRconnTo3 | 0.316951 | 2 | 1 | 1.001489 |
| 3 | Stim | AVFRconnTo3 | 0.31698 | 2 | 1 | 1.001367 |
| 4 | Stim | AVFRconnTo3 | 0.317191 | 2 | 1 | 1.000496 |
| 5 | Stim | AVFRconnTo3 | 0.045206 | 2 | 1 | 4.010926 |
| 6 | Stim | AVFRconnTo3 | 0.316951 | 2 | 1 | 1.001489 |
| 7 | Stim | AVFRconnTo3 | 0.31743 | 1 | 1 | 0.999505 |
| 1 | StimDetailed | AVFRconnTo3 | 0.000571 | 1 | 1 | 11.86984 |
| 2 | StimDetailed | AVFRconnTo3 | 0.56322 | 1 | 1 | 0.33416 |
| 3 | StimDetailed | AVFRconnTo3 | 0.56468 | 1 | 1 | 0.331666 |
| 4 | StimDetailed | AVFRconnTo3 | 0.563349 | 1 | 1 | 0.333939 |
| 5 | StimDetailed | AVFRconnTo3 | 0.247926 | 1 | 1 | 1.334952 |
| 6 | StimDetailed | AVFRconnTo3 | 0.56322 | 1 | 1 | 0.33416 |
| 7 | StimDetailed | AVFRconnTo3 | 0.563477 | 1 | 1 | 0.333719 |
| 1 | bhv | AVFRconnToNeg | 1.28E-12 | 1 | 3 | 58.41645 |
| 2 | bhv | AVFRconnToNeg | NaN | 1 | 3 | NaN |
| 3 | bhv | AVFRconnToNeg | NaN | 1 | 3 | NaN |
| 4 | bhv | AVFRconnToNeg | NaN | 1 | 3 | NaN |
| 5 | bhv | AVFRconnToNeg | 2.43E-13 | 4 | 3 | 61.79254 |
| 6 | bhv | AVFRconnToNeg | NaN | 1 | 3 | NaN |
| 7 | bhv | AVFRconnToNeg | NaN | 1 | 3 | NaN |
| 1 | FWD | AVFRconnToNeg | 3.85E-10 | 1 | 1 | 39.18842 |
| 2 | FWD | AVFRconnToNeg | NaN | 1 | 1 | NaN |
| 3 | FWD | AVFRconnToNeg | NaN | 1 | 1 | NaN |
| 4 | FWD | AVFRconnToNeg | NaN | 1 | 1 | NaN |
| 5 | FWD | AVFRconnToNeg | 0.006517 | 2 | 1 | 7.401417 |
| 6 | FWD | AVFRconnToNeg | NaN | 1 | 1 | NaN |
| 7 | FWD | AVFRconnToNeg | NaN | 1 | 1 | NaN |
| 1 | Stim | AVFRconnToNeg | 2.45E-05 | 2 | 1 | 17.80395 |
| 2 | Stim | AVFRconnToNeg | NaN | 1 | 1 | NaN |
| 3 | Stim | AVFRconnToNeg | NaN | 1 | 1 | NaN |
| 4 | Stim | AVFRconnToNeg | NaN | 1 | 1 | NaN |
| 5 | Stim | AVFRconnToNeg | 6.97E-06 | 1 | 1 | 20.20175 |
| 6 | Stim | AVFRconnToNeg | NaN | 1 | 1 | NaN |
| 7 | Stim | AVFRconnToNeg | NaN | 1 | 1 | NaN |
| 5 | StimDetailed | AVFRconnToNeg | 0.009514 | 1 | 1 | 6.723726 |
| 5 | StimDetailed | AVFRconnToNeg | 0.009514 | 1 | 1 | 6.723726 |
| 7 | StimDetailed | AVFRconnToNeg | NaN | 1 | 1 | NaN |
| 7 | StimDetailed | AVFRconnToNeg | NaN | 1 | 1 | NaN |
| 1 | bhv | AVFRconnToNeg3 | 1.87E-10 | 1 | 3 | 48.2618 |
| 2 | bhv | AVFRconnToNeg3 | NaN | 1 | 3 | NaN |
| 3 | bhv | AVFRconnToNeg3 | NaN | 1 | 3 | NaN |
| 4 | bhv | AVFRconnToNeg3 | NaN | 1 | 3 | NaN |
| 5 | bhv | AVFRconnToNeg3 | NaN | 1 | 3 | NaN |
| 6 | bhv | AVFRconnToNeg3 | NaN | 1 | 3 | NaN |
| 7 | bhv | AVFRconnToNeg3 | NaN | 1 | 3 | NaN |
| 1 | FWD | AVFRconnToNeg3 | 7.39E-09 | 1 | 1 | 33.42869 |
| 2 | FWD | AVFRconnToNeg3 | NaN | 1 | 1 | NaN |
| 3 | FWD | AVFRconnToNeg3 | NaN | 1 | 1 | NaN |
| 4 | FWD | AVFRconnToNeg3 | NaN | 1 | 1 | NaN |
| 5 | FWD | AVFRconnToNeg3 | NaN | 1 | 1 | NaN |
| 6 | FWD | AVFRconnToNeg3 | NaN | 1 | 1 | NaN |
| 7 | FWD | AVFRconnToNeg3 | NaN | 1 | 1 | NaN |
| 1 | Stim | AVFRconnToNeg3 | 9.48E-06 | 2 | 1 | 19.6139 |
| 2 | Stim | AVFRconnToNeg3 | NaN | 1 | 1 | NaN |
| 3 | Stim | AVFRconnToNeg3 | NaN | 1 | 1 | NaN |
| 4 | Stim | AVFRconnToNeg3 | NaN | 1 | 1 | NaN |
| 5 | Stim | AVFRconnToNeg3 | NaN | 1 | 1 | NaN |
| 6 | Stim | AVFRconnToNeg3 | NaN | 1 | 1 | NaN |
| 7 | Stim | AVFRconnToNeg3 | NaN | 1 | 1 | NaN |
| 5 | StimDetailed | AVFRconnToNeg3 | NaN | 1 | 1 | NaN |
| 5 | StimDetailed | AVFRconnToNeg3 | NaN | 1 | 1 | NaN |
| 7 | StimDetailed | AVFRconnToNeg3 | NaN | 1 | 1 | NaN |
| 7 | StimDetailed | AVFRconnToNeg3 | NaN | 1 | 1 | NaN |
| 1 | bhv | AVFRconnToPos | 0.153422 | 3 | 3 | 5.264503 |
| 2 | bhv | AVFRconnToPos | 0.852213 | 1 | 3 | 0.788512 |
| 3 | bhv | AVFRconnToPos | 0.86671 | 1 | 3 | 0.727532 |
| 4 | bhv | AVFRconnToPos | 0.007016 | 3 | 3 | 12.1099 |
| 5 | bhv | AVFRconnToPos | 0.004201 | 3 | 3 | 13.21129 |
| 6 | bhv | AVFRconnToPos | 0.821141 | 3 | 3 | 0.917755 |
| 7 | bhv | AVFRconnToPos | 0.760397 | 3 | 3 | 1.169215 |
| 1 | FWD | AVFRconnToPos | 0.455954 | 2 | 1 | 0.555808 |
| 2 | FWD | AVFRconnToPos | 0.374551 | 1 | 1 | 0.788512 |
| 3 | FWD | AVFRconnToPos | 0.393684 | 1 | 1 | 0.727532 |
| 4 | FWD | AVFRconnToPos | 0.521763 | 2 | 1 | 0.410406 |
| 5 | FWD | AVFRconnToPos | 0.104907 | 2 | 1 | 2.629308 |
| 6 | FWD | AVFRconnToPos | 0.552443 | 2 | 1 | 0.352958 |
| 7 | FWD | AVFRconnToPos | 0.390819 | 2 | 1 | 0.736392 |
| 1 | Stim | AVFRconnToPos | 0.3172 | 2 | 1 | 1.000455 |
| 2 | Stim | AVFRconnToPos | 0.316951 | 2 | 1 | 1.001489 |
| 3 | Stim | AVFRconnToPos | 0.31698 | 2 | 1 | 1.001367 |
| 4 | Stim | AVFRconnToPos | 0.317191 | 2 | 1 | 1.000496 |
| 5 | Stim | AVFRconnToPos | 0.045206 | 2 | 1 | 4.010926 |
| 6 | Stim | AVFRconnToPos | 0.316951 | 2 | 1 | 1.001489 |
| 7 | Stim | AVFRconnToPos | 0.31743 | 1 | 1 | 0.999505 |
| 1 | StimDetailed | AVFRconnToPos | 0.563378 | 1 | 1 | 0.33389 |
| 2 | StimDetailed | AVFRconnToPos | 0.56322 | 1 | 1 | 0.33416 |
| 3 | StimDetailed | AVFRconnToPos | 0.56468 | 1 | 1 | 0.331666 |
| 4 | StimDetailed | AVFRconnToPos | 0.563349 | 1 | 1 | 0.333939 |
| 5 | StimDetailed | AVFRconnToPos | 0.247926 | 1 | 1 | 1.334952 |
| 6 | StimDetailed | AVFRconnToPos | 0.56322 | 1 | 1 | 0.33416 |
| 7 | StimDetailed | AVFRconnToPos | 0.563477 | 1 | 1 | 0.333719 |
| 1 | bhv | AVFRconnToPos3 | 0.000329 | 3 | 3 | 18.61041 |
| 2 | bhv | AVFRconnToPos3 | 0.852213 | 1 | 3 | 0.788512 |
| 3 | bhv | AVFRconnToPos3 | 0.86671 | 1 | 3 | 0.727532 |
| 4 | bhv | AVFRconnToPos3 | 0.007016 | 3 | 3 | 12.1099 |
| 5 | bhv | AVFRconnToPos3 | 0.004201 | 3 | 3 | 13.21129 |
| 6 | bhv | AVFRconnToPos3 | 0.821141 | 3 | 3 | 0.917755 |
| 7 | bhv | AVFRconnToPos3 | 0.760397 | 3 | 3 | 1.169215 |
| 1 | FWD | AVFRconnToPos3 | 0.071189 | 2 | 1 | 3.2554 |
| 2 | FWD | AVFRconnToPos3 | 0.374551 | 1 | 1 | 0.788512 |
| 3 | FWD | AVFRconnToPos3 | 0.393684 | 1 | 1 | 0.727532 |
| 4 | FWD | AVFRconnToPos3 | 0.521763 | 2 | 1 | 0.410406 |
| 5 | FWD | AVFRconnToPos3 | 0.104907 | 2 | 1 | 2.629308 |
| 6 | FWD | AVFRconnToPos3 | 0.552443 | 2 | 1 | 0.352958 |
| 7 | FWD | AVFRconnToPos3 | 0.390819 | 2 | 1 | 0.736392 |
| 1 | Stim | AVFRconnToPos3 | 2.75E-07 | 1 | 1 | 26.41767 |
| 2 | Stim | AVFRconnToPos3 | 0.316951 | 2 | 1 | 1.001489 |
| 3 | Stim | AVFRconnToPos3 | 0.31698 | 2 | 1 | 1.001367 |
| 4 | Stim | AVFRconnToPos3 | 0.317191 | 2 | 1 | 1.000496 |
| 5 | Stim | AVFRconnToPos3 | 0.045206 | 2 | 1 | 4.010926 |
| 6 | Stim | AVFRconnToPos3 | 0.316951 | 2 | 1 | 1.001489 |
| 7 | Stim | AVFRconnToPos3 | 0.31743 | 1 | 1 | 0.999505 |
| 1 | StimDetailed | AVFRconnToPos3 | 0.000394 | 1 | 1 | 12.55972 |
| 2 | StimDetailed | AVFRconnToPos3 | 0.56322 | 1 | 1 | 0.33416 |
| 3 | StimDetailed | AVFRconnToPos3 | 0.56468 | 1 | 1 | 0.331666 |
| 4 | StimDetailed | AVFRconnToPos3 | 0.563349 | 1 | 1 | 0.333939 |
| 5 | StimDetailed | AVFRconnToPos3 | 0.247926 | 1 | 1 | 1.334952 |
| 6 | StimDetailed | AVFRconnToPos3 | 0.56322 | 1 | 1 | 0.33416 |
| 7 | StimDetailed | AVFRconnToPos3 | 0.563477 | 1 | 1 | 0.333719 |
| 1 | bhv | AVFRyhat | 0 | 1 | 3 | 803.8678 |
| 2 | bhv | AVFRyhat | 0 | 1 | 3 | 828.4552 |
| 3 | bhv | AVFRyhat | 0 | 1 | 3 | 605.7794 |
| 4 | bhv | AVFRyhat | 0 | 1 | 3 | 501.9176 |
| 5 | bhv | AVFRyhat | 0 | 1 | 3 | 177.3765 |
| 6 | bhv | AVFRyhat | 0 | 1 | 3 | 351.3449 |
| 7 | bhv | AVFRyhat | 0 | 3 | 3 | 170.6179 |
| 1 | FWD | AVFRyhat | 0 | 1 | 1 | 664.3134 |
| 2 | FWD | AVFRyhat | 0 | 1 | 1 | 812.1669 |
| 3 | FWD | AVFRyhat | 0 | 1 | 1 | 507.6664 |
| 4 | FWD | AVFRyhat | 0 | 2 | 1 | 220.561 |
| 5 | FWD | AVFRyhat | 0 | 1 | 1 | 146.3624 |
| 6 | FWD | AVFRyhat | 0 | 2 | 1 | 215.9023 |
| 7 | FWD | AVFRyhat | 0 | 2 | 1 | 91.51452 |
| 1 | Stim | AVFRyhat | 0.015033 | 1 | 1 | 5.912648 |
| 2 | Stim | AVFRyhat | 0.640085 | 2 | 1 | 0.218631 |
| 3 | Stim | AVFRyhat | 1.11E-16 | 1 | 1 | 68.16997 |
| 4 | Stim | AVFRyhat | 0 | 1 | 1 | 315.5619 |
| 5 | Stim | AVFRyhat | 0.359371 | 1 | 1 | 0.840091 |
| 6 | Stim | AVFRyhat | 4.04E-06 | 1 | 1 | 21.24404 |
| 7 | Stim | AVFRyhat | 4.44E-16 | 2 | 1 | 65.98585 |
| 1 | StimD | AVFRyhat | 0.076264 | 1 | 1 | 3.142771 |
| 2 | StimD | AVFRyhat | 4.61E-10 | 1 | 1 | 38.83551 |
| 3 | StimD | AVFRyhat | 2.60E-05 | 1 | 1 | 17.69038 |
| 4 | StimD | AVFRyhat | 0 | 1 | 1 | 214.5263 |
| 5 | StimD | AVFRyhat | 2.14E-08 | 1 | 1 | 31.36274 |
| 6 | StimD | AVFRyhat | 0.055553 | 1 | 1 | 3.665423 |
| 7 | StimD | AVFRyhat | 0.33529 | 1 | 1 | 0.928359 |
| 1 | bhv | AVFRyhat3 | 4.20E-06 | 3 | 3 | 27.69767 |
| 2 | bhv | AVFRyhat3 | 0.367791 | 1 | 3 | 3.158747 |
| 3 | bhv | AVFRyhat3 | 0.16415 | 1 | 3 | 5.10668 |
| 4 | bhv | AVFRyhat3 | 8.86E-13 | 4 | 3 | 59.16692 |
| 5 | bhv | AVFRyhat3 | 0 | 4 | 3 | 100.8311 |
| 6 | bhv | AVFRyhat3 | 0 | 4 | 3 | 118.5279 |
| 7 | bhv | AVFRyhat3 | 0 | 4 | 3 | 123.8975 |
| 1 | FWD | AVFRyhat3 | 0.00546 | 2 | 1 | 7.720573 |
| 2 | FWD | AVFRyhat3 | 0.075521 | 1 | 1 | 3.158747 |
| 3 | FWD | AVFRyhat3 | 0.023834 | 1 | 1 | 5.10668 |
| 4 | FWD | AVFRyhat3 | 0.042318 | 2 | 1 | 4.122455 |
| 5 | FWD | AVFRyhat3 | 0.012274 | 2 | 1 | 6.270887 |
| 6 | FWD | AVFRyhat3 | 0.011365 | 2 | 1 | 6.40728 |
| 7 | FWD | AVFRyhat3 | 0.011078 | 2 | 1 | 6.452826 |
| 1 | Stim | AVFRyhat3 | 1.01E-07 | 1 | 1 | 28.36129 |
| 2 | Stim | AVFRyhat3 | 0.045393 | 2 | 1 | 4.003976 |
| 3 | Stim | AVFRyhat3 | 0.008021 | 1 | 1 | 7.02877 |
| 4 | Stim | AVFRyhat3 | 0.001524 | 1 | 1 | 10.0498 |
| 5 | Stim | AVFRyhat3 | 0.000612 | 2 | 1 | 11.73917 |
| 6 | Stim | AVFRyhat3 | 2.01E-05 | 1 | 1 | 18.1801 |
| 7 | Stim | AVFRyhat3 | 0.0003 | 1 | 1 | 13.07129 |
| 1 | StimD | AVFRyhat3 | 0.002094 | 1 | 1 | 9.465236 |
| 2 | StimD | AVFRyhat3 | 0.247275 | 1 | 1 | 1.338634 |
| 3 | StimD | AVFRyhat3 | 0.127063 | 1 | 1 | 2.328024 |
| 4 | StimD | AVFRyhat3 | 0.067027 | 1 | 1 | 3.354362 |
| 5 | StimD | AVFRyhat3 | 2.35E-11 | 2 | 1 | 44.65795 |
| 6 | StimD | AVFRyhat3 | 0.013781 | 1 | 1 | 6.066039 |
| 7 | StimD | AVFRyhat3 | 0.0367 | 1 | 1 | 4.364296 |
| 1 | bhv | BAGLconnFrom | 0.153422 | 3 | 3 | 5.264503 |
| 2 | bhv | BAGLconnFrom | 0.852213 | 1 | 3 | 0.788512 |
| 3 | bhv | BAGLconnFrom | 0.86671 | 1 | 3 | 0.727532 |
| 4 | bhv | BAGLconnFrom | 0.133764 | 3 | 3 | 5.582888 |
| 5 | bhv | BAGLconnFrom | 0.008364 | 3 | 3 | 11.73084 |
| 6 | bhv | BAGLconnFrom | 0 | 3 | 3 | 268.1857 |
| 7 | bhv | BAGLconnFrom | 0.760397 | 3 | 3 | 1.169215 |
| 1 | FWD | BAGLconnFrom | 0.455954 | 2 | 1 | 0.555808 |
| 2 | FWD | BAGLconnFrom | 0.374551 | 1 | 1 | 0.788512 |
| 3 | FWD | BAGLconnFrom | 0.393684 | 1 | 1 | 0.727532 |
| 4 | FWD | BAGLconnFrom | 0.364824 | 2 | 1 | 0.821219 |
| 5 | FWD | BAGLconnFrom | 0.069782 | 2 | 1 | 3.288132 |
| 6 | FWD | BAGLconnFrom | 0 | 2 | 1 | 108.096 |
| 7 | FWD | BAGLconnFrom | 0.390819 | 2 | 1 | 0.736392 |
| 1 | Stim | BAGLconnFrom | 0.3172 | 2 | 1 | 1.000455 |
| 2 | Stim | BAGLconnFrom | 0.316951 | 2 | 1 | 1.001489 |
| 3 | Stim | BAGLconnFrom | 0.31698 | 2 | 1 | 1.001367 |
| 4 | Stim | BAGLconnFrom | 0.157093 | 2 | 1 | 2.001984 |
| 5 | Stim | BAGLconnFrom | 0.025115 | 2 | 1 | 5.01594 |
| 6 | Stim | BAGLconnFrom | 0 | 2 | 1 | 306.7129 |
| 7 | Stim | BAGLconnFrom | 0.31743 | 1 | 1 | 0.999505 |
| 1 | StimDetailed | BAGLconnFrom | 0.563378 | 1 | 1 | 0.33389 |
| 2 | StimDetailed | BAGLconnFrom | 0.56322 | 1 | 1 | 0.33416 |
| 3 | StimDetailed | BAGLconnFrom | 0.56468 | 1 | 1 | 0.331666 |
| 4 | StimDetailed | BAGLconnFrom | 0.413676 | 1 | 1 | 0.66821 |
| 5 | StimDetailed | BAGLconnFrom | 0.196332 | 1 | 1 | 1.66945 |
| 6 | StimDetailed | BAGLconnFrom | 0 | 1 | 1 | 102.339 |
| 7 | StimDetailed | BAGLconnFrom | 0.563477 | 1 | 1 | 0.333719 |
| 1 | bhv | BAGLconnFrom3 | 0.035457 | 3 | 3 | 8.578265 |
| 2 | bhv | BAGLconnFrom3 | 4.44E-16 | 1 | 3 | 74.58049 |
| 3 | bhv | BAGLconnFrom3 | 0 | 3 | 3 | 785.8092 |
| 4 | bhv | BAGLconnFrom3 | 0 | 3 | 3 | 315.0905 |
| 5 | bhv | BAGLconnFrom3 | 0 | 1 | 3 | 306.5793 |
| 6 | bhv | BAGLconnFrom3 | 0 | 1 | 3 | 726.7748 |
| 7 | bhv | BAGLconnFrom3 | 0 | 1 | 3 | 125.6072 |
| 1 | FWD | BAGLconnFrom3 | 0.174965 | 2 | 1 | 1.839876 |
| 2 | FWD | BAGLconnFrom3 | 3.19E-12 | 1 | 1 | 48.57108 |
| 3 | FWD | BAGLconnFrom3 | 0 | 2 | 1 | 784.0925 |
| 4 | FWD | BAGLconnFrom3 | 0 | 2 | 1 | 312.3635 |
| 5 | FWD | BAGLconnFrom3 | 0 | 1 | 1 | 253.808 |
| 6 | FWD | BAGLconnFrom3 | 0 | 1 | 1 | 378.5621 |
| 7 | FWD | BAGLconnFrom3 | 0 | 1 | 1 | 90.66841 |
| 1 | Stim | BAGLconnFrom3 | 0.068785 | 2 | 1 | 3.311777 |
| 2 | Stim | BAGLconnFrom3 | 9.64E-11 | 2 | 1 | 41.89223 |
| 3 | Stim | BAGLconnFrom3 | 0 | 1 | 1 | 77.18278 |
| 4 | Stim | BAGLconnFrom3 | 0 | 2 | 1 | 124.9179 |
| 5 | Stim | BAGLconnFrom3 | 0 | 2 | 1 | 420.0473 |
| 6 | Stim | BAGLconnFrom3 | 0 | 2 | 1 | 136.8379 |
| 7 | Stim | BAGLconnFrom3 | 0 | 2 | 1 | 511.749 |
| 1 | StimDetailed | BAGLconnFrom3 | 0 | 1 | 1 | 76.3616 |
| 2 | StimDetailed | BAGLconnFrom3 | 0.000185 | 1 | 1 | 13.97792 |
| 3 | StimDetailed | BAGLconnFrom3 | 3.35E-13 | 1 | 1 | 52.99006 |
| 4 | StimDetailed | BAGLconnFrom3 | 0 | 1 | 1 | 444.1357 |
| 5 | StimDetailed | BAGLconnFrom3 | 0 | 2 | 1 | 853.7878 |
| 6 | StimDetailed | BAGLconnFrom3 | 0 | 1 | 1 | 111.7575 |
| 7 | StimDetailed | BAGLconnFrom3 | 0 | 1 | 1 | 405.762 |
| 1 | bhv | BAGLconnFromNeg | 0.069563 | 3 | 3 | 7.074426 |
| 2 | bhv | BAGLconnFromNeg | NaN | 1 | 3 | NaN |
| 3 | bhv | BAGLconnFromNeg | NaN | 1 | 3 | NaN |
| 4 | bhv | BAGLconnFromNeg | NaN | 1 | 3 | NaN |
| 5 | bhv | BAGLconnFromNeg | 5.45E-09 | 2 | 3 | 41.37183 |
| 6 | bhv | BAGLconnFromNeg | NaN | 1 | 3 | NaN |
| 7 | bhv | BAGLconnFromNeg | NaN | 1 | 3 | NaN |
| 1 | FWD | BAGLconnFromNeg | 0.441856 | 2 | 1 | 0.591456 |
| 2 | FWD | BAGLconnFromNeg | NaN | 1 | 1 | NaN |
| 3 | FWD | BAGLconnFromNeg | NaN | 1 | 1 | NaN |
| 4 | FWD | BAGLconnFromNeg | NaN | 1 | 1 | NaN |
| 5 | FWD | BAGLconnFromNeg | 0.160333 | 2 | 1 | 1.971084 |
| 6 | FWD | BAGLconnFromNeg | NaN | 1 | 1 | NaN |
| 7 | FWD | BAGLconnFromNeg | NaN | 1 | 1 | NaN |
| 1 | Stim | BAGLconnFromNeg | 5.28E-07 | 1 | 1 | 25.15744 |
| 2 | Stim | BAGLconnFromNeg | NaN | 1 | 1 | NaN |
| 3 | Stim | BAGLconnFromNeg | NaN | 1 | 1 | NaN |
| 4 | Stim | BAGLconnFromNeg | NaN | 1 | 1 | NaN |
| 5 | Stim | BAGLconnFromNeg | 0.082915 | 1 | 1 | 3.006826 |
| 6 | Stim | BAGLconnFromNeg | NaN | 1 | 1 | NaN |
| 7 | Stim | BAGLconnFromNeg | NaN | 1 | 1 | NaN |
| 5 | StimDetailed | BAGLconnFromNeg | 0.317127 | 1 | 1 | 1.000758 |
| 7 | StimDetailed | BAGLconnFromNeg | NaN | 1 | 1 | NaN |
| 1 | bhv | BAGLconnFromNeg3 | 3.59E-13 | 3 | 3 | 61.00187 |
| 2 | bhv | BAGLconnFromNeg3 | NaN | 1 | 3 | NaN |
| 3 | bhv | BAGLconnFromNeg3 | NaN | 1 | 3 | NaN |
| 4 | bhv | BAGLconnFromNeg3 | NaN | 1 | 3 | NaN |
| 5 | bhv | BAGLconnFromNeg3 | NaN | 1 | 3 | NaN |
| 6 | bhv | BAGLconnFromNeg3 | NaN | 1 | 3 | NaN |
| 7 | bhv | BAGLconnFromNeg3 | NaN | 1 | 3 | NaN |
| 1 | FWD | BAGLconnFromNeg3 | 4.11E-13 | 2 | 1 | 52.5879 |
| 2 | FWD | BAGLconnFromNeg3 | NaN | 1 | 1 | NaN |
| 3 | FWD | BAGLconnFromNeg3 | NaN | 1 | 1 | NaN |
| 4 | FWD | BAGLconnFromNeg3 | NaN | 1 | 1 | NaN |
| 5 | FWD | BAGLconnFromNeg3 | NaN | 1 | 1 | NaN |
| 6 | FWD | BAGLconnFromNeg3 | NaN | 1 | 1 | NaN |
| 7 | FWD | BAGLconnFromNeg3 | NaN | 1 | 1 | NaN |
| 1 | Stim | BAGLconnFromNeg3 | 0 | 1 | 1 | 117.7325 |
| 2 | Stim | BAGLconnFromNeg3 | NaN | 1 | 1 | NaN |
| 3 | Stim | BAGLconnFromNeg3 | NaN | 1 | 1 | NaN |
| 4 | Stim | BAGLconnFromNeg3 | NaN | 1 | 1 | NaN |
| 5 | Stim | BAGLconnFromNeg3 | NaN | 1 | 1 | NaN |
| 6 | Stim | BAGLconnFromNeg3 | NaN | 1 | 1 | NaN |
| 7 | Stim | BAGLconnFromNeg3 | NaN | 1 | 1 | NaN |
| 5 | StimDetailed | BAGLconnFromNeg3 | NaN | 1 | 1 | NaN |
| 7 | StimDetailed | BAGLconnFromNeg3 | NaN | 1 | 1 | NaN |
| 1 | bhv | BAGLconnFromPos | 0.153422 | 3 | 3 | 5.264503 |
| 2 | bhv | BAGLconnFromPos | 0.852213 | 1 | 3 | 0.788512 |
| 3 | bhv | BAGLconnFromPos | 0.86671 | 1 | 3 | 0.727532 |
| 4 | bhv | BAGLconnFromPos | 0.133764 | 3 | 3 | 5.582888 |
| 5 | bhv | BAGLconnFromPos | 6.90E-05 | 3 | 3 | 21.88323 |
| 6 | bhv | BAGLconnFromPos | 0 | 3 | 3 | 268.1857 |
| 7 | bhv | BAGLconnFromPos | 0.760397 | 3 | 3 | 1.169215 |
| 1 | FWD | BAGLconnFromPos | 0.455954 | 2 | 1 | 0.555808 |
| 2 | FWD | BAGLconnFromPos | 0.374551 | 1 | 1 | 0.788512 |
| 3 | FWD | BAGLconnFromPos | 0.393684 | 1 | 1 | 0.727532 |
| 4 | FWD | BAGLconnFromPos | 0.364824 | 2 | 1 | 0.821219 |
| 5 | FWD | BAGLconnFromPos | 0.031831 | 2 | 1 | 4.607581 |
| 6 | FWD | BAGLconnFromPos | 0 | 2 | 1 | 108.096 |
| 7 | FWD | BAGLconnFromPos | 0.390819 | 2 | 1 | 0.736392 |
| 1 | Stim | BAGLconnFromPos | 0.3172 | 2 | 1 | 1.000455 |
| 2 | Stim | BAGLconnFromPos | 0.316951 | 2 | 1 | 1.001489 |
| 3 | Stim | BAGLconnFromPos | 0.31698 | 2 | 1 | 1.001367 |
| 4 | Stim | BAGLconnFromPos | 0.157093 | 2 | 1 | 2.001984 |
| 5 | Stim | BAGLconnFromPos | 0.008021 | 2 | 1 | 7.028718 |
| 6 | Stim | BAGLconnFromPos | 0 | 2 | 1 | 306.7129 |
| 7 | Stim | BAGLconnFromPos | 0.31743 | 1 | 1 | 0.999505 |
| 1 | StimDetailed | BAGLconnFromPos | 0.563378 | 1 | 1 | 0.33389 |
| 2 | StimDetailed | BAGLconnFromPos | 0.56322 | 1 | 1 | 0.33416 |
| 3 | StimDetailed | BAGLconnFromPos | 0.56468 | 1 | 1 | 0.331666 |
| 4 | StimDetailed | BAGLconnFromPos | 0.413676 | 1 | 1 | 0.66821 |
| 5 | StimDetailed | BAGLconnFromPos | 0.126141 | 1 | 1 | 2.33936 |
| 6 | StimDetailed | BAGLconnFromPos | 0 | 1 | 1 | 102.339 |
| 7 | StimDetailed | BAGLconnFromPos | 0.563477 | 1 | 1 | 0.333719 |
| 1 | bhv | BAGLconnFromPos3 | 0.530963 | 3 | 3 | 2.205007 |
| 2 | bhv | BAGLconnFromPos3 | 4.44E-16 | 1 | 3 | 74.58049 |
| 3 | bhv | BAGLconnFromPos3 | 0 | 3 | 3 | 785.8092 |
| 4 | bhv | BAGLconnFromPos3 | 0 | 3 | 3 | 315.0905 |
| 5 | bhv | BAGLconnFromPos3 | 0 | 1 | 3 | 306.5793 |
| 6 | bhv | BAGLconnFromPos3 | 0 | 1 | 3 | 726.7748 |
| 7 | bhv | BAGLconnFromPos3 | 0 | 1 | 3 | 125.6072 |
| 1 | FWD | BAGLconnFromPos3 | 0.36935 | 2 | 1 | 0.805851 |
| 2 | FWD | BAGLconnFromPos3 | 3.19E-12 | 1 | 1 | 48.57108 |
| 3 | FWD | BAGLconnFromPos3 | 0 | 2 | 1 | 784.0925 |
| 4 | FWD | BAGLconnFromPos3 | 0 | 2 | 1 | 312.3635 |
| 5 | FWD | BAGLconnFromPos3 | 0 | 1 | 1 | 253.808 |
| 6 | FWD | BAGLconnFromPos3 | 0 | 1 | 1 | 378.5621 |
| 7 | FWD | BAGLconnFromPos3 | 0 | 1 | 1 | 90.66841 |
| 1 | Stim | BAGLconnFromPos3 | 0.006857 | 2 | 1 | 7.310053 |
| 2 | Stim | BAGLconnFromPos3 | 9.64E-11 | 2 | 1 | 41.89223 |
| 3 | Stim | BAGLconnFromPos3 | 0 | 1 | 1 | 77.18278 |
| 4 | Stim | BAGLconnFromPos3 | 0 | 2 | 1 | 124.9179 |
| 5 | Stim | BAGLconnFromPos3 | 0 | 2 | 1 | 420.0473 |
| 6 | Stim | BAGLconnFromPos3 | 0 | 2 | 1 | 136.8379 |
| 7 | Stim | BAGLconnFromPos3 | 0 | 2 | 1 | 511.749 |
| 1 | StimDetailed | BAGLconnFromPos3 | 0 | 1 | 1 | 82.61539 |
| 2 | StimDetailed | BAGLconnFromPos3 | 0.000185 | 1 | 1 | 13.97792 |
| 3 | StimDetailed | BAGLconnFromPos3 | 3.35E-13 | 1 | 1 | 52.99006 |
| 4 | StimDetailed | BAGLconnFromPos3 | 0 | 1 | 1 | 444.1357 |
| 5 | StimDetailed | BAGLconnFromPos3 | 0 | 2 | 1 | 853.7878 |
| 6 | StimDetailed | BAGLconnFromPos3 | 0 | 1 | 1 | 111.7575 |
| 7 | StimDetailed | BAGLconnFromPos3 | 0 | 1 | 1 | 405.762 |
| 1 | bhv | BAGLconnTo | 0.153422 | 3 | 3 | 5.264503 |
| 2 | bhv | BAGLconnTo | 0.852213 | 1 | 3 | 0.788512 |
| 3 | bhv | BAGLconnTo | 0.86671 | 1 | 3 | 0.727532 |
| 4 | bhv | BAGLconnTo | 0.007016 | 3 | 3 | 12.1099 |
| 5 | bhv | BAGLconnTo | 0.004201 | 3 | 3 | 13.21129 |
| 6 | bhv | BAGLconnTo | 0 | 3 | 3 | 91.45084 |
| 7 | bhv | BAGLconnTo | 0.760397 | 3 | 3 | 1.169215 |
| 1 | FWD | BAGLconnTo | 0.455954 | 2 | 1 | 0.555808 |
| 2 | FWD | BAGLconnTo | 0.374551 | 1 | 1 | 0.788512 |
| 3 | FWD | BAGLconnTo | 0.393684 | 1 | 1 | 0.727532 |
| 4 | FWD | BAGLconnTo | 0.521763 | 2 | 1 | 0.410406 |
| 5 | FWD | BAGLconnTo | 0.104907 | 2 | 1 | 2.629308 |
| 6 | FWD | BAGLconnTo | 3.02E-09 | 2 | 1 | 35.17096 |
| 7 | FWD | BAGLconnTo | 0.390819 | 2 | 1 | 0.736392 |
| 1 | Stim | BAGLconnTo | 0.3172 | 2 | 1 | 1.000455 |
| 2 | Stim | BAGLconnTo | 0.316951 | 2 | 1 | 1.001489 |
| 3 | Stim | BAGLconnTo | 0.31698 | 2 | 1 | 1.001367 |
| 4 | Stim | BAGLconnTo | 0.317191 | 2 | 1 | 1.000496 |
| 5 | Stim | BAGLconnTo | 0.045206 | 2 | 1 | 4.010926 |
| 6 | Stim | BAGLconnTo | 0 | 2 | 1 | 99.79453 |
| 7 | Stim | BAGLconnTo | 0.31743 | 1 | 1 | 0.999505 |
| 1 | StimDetailed | BAGLconnTo | 0.563378 | 1 | 1 | 0.33389 |
| 2 | StimDetailed | BAGLconnTo | 0.56322 | 1 | 1 | 0.33416 |
| 3 | StimDetailed | BAGLconnTo | 0.56468 | 1 | 1 | 0.331666 |
| 4 | StimDetailed | BAGLconnTo | 0.563349 | 1 | 1 | 0.333939 |
| 5 | StimDetailed | BAGLconnTo | 0.247926 | 1 | 1 | 1.334952 |
| 6 | StimDetailed | BAGLconnTo | 7.91E-09 | 1 | 1 | 33.29781 |
| 7 | StimDetailed | BAGLconnTo | 0.563477 | 1 | 1 | 0.333719 |
| 1 | bhv | BAGLconnTo3 | 0 | 3 | 3 | 82.77762 |
| 2 | bhv | BAGLconnTo3 | 0.852213 | 1 | 3 | 0.788512 |
| 3 | bhv | BAGLconnTo3 | 1.49E-07 | 1 | 3 | 34.59181 |
| 4 | bhv | BAGLconnTo3 | 0 | 3 | 3 | 344.1666 |
| 5 | bhv | BAGLconnTo3 | 0 | 1 | 3 | 469.5483 |
| 6 | bhv | BAGLconnTo3 | 0 | 1 | 3 | 1256.707 |
| 7 | bhv | BAGLconnTo3 | 0 | 3 | 3 | 174.5673 |
| 1 | FWD | BAGLconnTo3 | 1.68E-14 | 2 | 1 | 58.88097 |
| 2 | FWD | BAGLconnTo3 | 0.374551 | 1 | 1 | 0.788512 |
| 3 | FWD | BAGLconnTo3 | 2.54E-08 | 1 | 1 | 31.03315 |
| 4 | FWD | BAGLconnTo3 | 0 | 2 | 1 | 342.8264 |
| 5 | FWD | BAGLconnTo3 | 0 | 1 | 1 | 449.5595 |
| 6 | FWD | BAGLconnTo3 | 0 | 1 | 1 | 756.9948 |
| 7 | FWD | BAGLconnTo3 | 0 | 2 | 1 | 173.9562 |
| 1 | Stim | BAGLconnTo3 | 0.403364 | 1 | 1 | 0.698274 |
| 2 | Stim | BAGLconnTo3 | 0.316951 | 2 | 1 | 1.001489 |
| 3 | Stim | BAGLconnTo3 | 0.236329 | 1 | 1 | 1.402361 |
| 4 | Stim | BAGLconnTo3 | 0.003525 | 2 | 1 | 8.51377 |
| 5 | Stim | BAGLconnTo3 | 0 | 2 | 1 | 659.1007 |
| 6 | Stim | BAGLconnTo3 | 2.34E-09 | 2 | 1 | 35.66844 |
| 7 | Stim | BAGLconnTo3 | 0 | 2 | 1 | 130.3942 |
| 1 | StimDetailed | BAGLconnTo3 | 4.44E-13 | 1 | 1 | 52.43667 |
| 2 | StimDetailed | BAGLconnTo3 | 0.56322 | 1 | 1 | 0.33416 |
| 3 | StimDetailed | BAGLconnTo3 | 0.00449 | 1 | 1 | 8.074173 |
| 4 | StimDetailed | BAGLconnTo3 | 0 | 1 | 1 | 109.5453 |
| 5 | StimDetailed | BAGLconnTo3 | 0 | 2 | 1 | 560.7455 |
| 6 | StimDetailed | BAGLconnTo3 | 0.010911 | 1 | 1 | 6.479725 |
| 7 | StimDetailed | BAGLconnTo3 | 1.67E-10 | 1 | 1 | 40.82013 |
| 1 | bhv | BAGLconnToNeg | 0.000985 | 3 | 3 | 16.29781 |
| 2 | bhv | BAGLconnToNeg | NaN | 1 | 3 | NaN |
| 3 | bhv | BAGLconnToNeg | NaN | 1 | 3 | NaN |
| 4 | bhv | BAGLconnToNeg | NaN | 1 | 3 | NaN |
| 5 | bhv | BAGLconnToNeg | 4.99E-10 | 3 | 3 | 46.26315 |
| 6 | bhv | BAGLconnToNeg | NaN | 1 | 3 | NaN |
| 7 | bhv | BAGLconnToNeg | 2.04E-10 | 1 | 3 | 48.09132 |
| 1 | FWD | BAGLconnToNeg | 0.002475 | 2 | 1 | 9.158899 |
| 2 | FWD | BAGLconnToNeg | NaN | 1 | 1 | NaN |
| 3 | FWD | BAGLconnToNeg | NaN | 1 | 1 | NaN |
| 4 | FWD | BAGLconnToNeg | NaN | 1 | 1 | NaN |
| 5 | FWD | BAGLconnToNeg | 3.86E-10 | 2 | 1 | 39.18124 |
| 6 | FWD | BAGLconnToNeg | NaN | 1 | 1 | NaN |
| 7 | FWD | BAGLconnToNeg | 0.111068 | 1 | 1 | 2.53896 |
| 1 | Stim | BAGLconnToNeg | 4.60E-12 | 1 | 1 | 47.85242 |
| 2 | Stim | BAGLconnToNeg | NaN | 1 | 1 | NaN |
| 3 | Stim | BAGLconnToNeg | NaN | 1 | 1 | NaN |
| 4 | Stim | BAGLconnToNeg | NaN | 1 | 1 | NaN |
| 5 | Stim | BAGLconnToNeg | 2.79E-10 | 1 | 1 | 39.81288 |
| 6 | Stim | BAGLconnToNeg | NaN | 1 | 1 | NaN |
| 7 | Stim | BAGLconnToNeg | 3.37E-08 | 2 | 1 | 30.48267 |
| 5 | StimDetailed | BAGLconnToNeg | 0.064316 | 1 | 1 | 3.422428 |
| 5 | StimDetailed | BAGLconnToNeg | 0.064316 | 1 | 1 | 3.422428 |
| 7 | StimDetailed | BAGLconnToNeg | 0 | 2 | 1 | 91.29716 |
| 7 | StimDetailed | BAGLconnToNeg | 0 | 2 | 1 | 91.29716 |
| 1 | bhv | BAGLconnToNeg3 | 1.04E-12 | 3 | 3 | 58.83325 |
| 2 | bhv | BAGLconnToNeg3 | NaN | 1 | 3 | NaN |
| 3 | bhv | BAGLconnToNeg3 | NaN | 1 | 3 | NaN |
| 4 | bhv | BAGLconnToNeg3 | NaN | 1 | 3 | NaN |
| 5 | bhv | BAGLconnToNeg3 | 8.62E-14 | 4 | 3 | 63.90237 |
| 6 | bhv | BAGLconnToNeg3 | NaN | 1 | 3 | NaN |
| 7 | bhv | BAGLconnToNeg3 | NaN | 1 | 3 | NaN |
| 1 | FWD | BAGLconnToNeg3 | 1.20E-12 | 2 | 1 | 50.47947 |
| 2 | FWD | BAGLconnToNeg3 | NaN | 1 | 1 | NaN |
| 3 | FWD | BAGLconnToNeg3 | NaN | 1 | 1 | NaN |
| 4 | FWD | BAGLconnToNeg3 | NaN | 1 | 1 | NaN |
| 5 | FWD | BAGLconnToNeg3 | 0.010248 | 2 | 1 | 6.591271 |
| 6 | FWD | BAGLconnToNeg3 | NaN | 1 | 1 | NaN |
| 7 | FWD | BAGLconnToNeg3 | NaN | 1 | 1 | NaN |
| 1 | Stim | BAGLconnToNeg3 | 0 | 1 | 1 | 118.8466 |
| 2 | Stim | BAGLconnToNeg3 | NaN | 1 | 1 | NaN |
| 3 | Stim | BAGLconnToNeg3 | NaN | 1 | 1 | NaN |
| 4 | Stim | BAGLconnToNeg3 | NaN | 1 | 1 | NaN |
| 5 | Stim | BAGLconnToNeg3 | 0.001535 | 2 | 1 | 10.03652 |
| 6 | Stim | BAGLconnToNeg3 | NaN | 1 | 1 | NaN |
| 7 | Stim | BAGLconnToNeg3 | NaN | 1 | 1 | NaN |
| 5 | StimDetailed | BAGLconnToNeg3 | 3.99E-08 | 2 | 1 | 30.15519 |
| 5 | StimDetailed | BAGLconnToNeg3 | 3.99E-08 | 2 | 1 | 30.15519 |
| 7 | StimDetailed | BAGLconnToNeg3 | NaN | 1 | 1 | NaN |
| 7 | StimDetailed | BAGLconnToNeg3 | NaN | 1 | 1 | NaN |
| 1 | bhv | BAGLconnToPos | 0.153422 | 3 | 3 | 5.264503 |
| 2 | bhv | BAGLconnToPos | 0.852213 | 1 | 3 | 0.788512 |
| 3 | bhv | BAGLconnToPos | 0.86671 | 1 | 3 | 0.727532 |
| 4 | bhv | BAGLconnToPos | 0.007016 | 3 | 3 | 12.1099 |
| 5 | bhv | BAGLconnToPos | 1.19E-11 | 3 | 3 | 53.87812 |
| 6 | bhv | BAGLconnToPos | 0 | 3 | 3 | 91.45084 |
| 7 | bhv | BAGLconnToPos | 0.760397 | 3 | 3 | 1.169215 |
| 1 | FWD | BAGLconnToPos | 0.455954 | 2 | 1 | 0.555808 |
| 2 | FWD | BAGLconnToPos | 0.374551 | 1 | 1 | 0.788512 |
| 3 | FWD | BAGLconnToPos | 0.393684 | 1 | 1 | 0.727532 |
| 4 | FWD | BAGLconnToPos | 0.521763 | 2 | 1 | 0.410406 |
| 5 | FWD | BAGLconnToPos | 0.020803 | 2 | 1 | 5.343192 |
| 6 | FWD | BAGLconnToPos | 3.02E-09 | 2 | 1 | 35.17096 |
| 7 | FWD | BAGLconnToPos | 0.390819 | 2 | 1 | 0.736392 |
| 1 | Stim | BAGLconnToPos | 0.3172 | 2 | 1 | 1.000455 |
| 2 | Stim | BAGLconnToPos | 0.316951 | 2 | 1 | 1.001489 |
| 3 | Stim | BAGLconnToPos | 0.31698 | 2 | 1 | 1.001367 |
| 4 | Stim | BAGLconnToPos | 0.317191 | 2 | 1 | 1.000496 |
| 5 | Stim | BAGLconnToPos | 8.30E-07 | 2 | 1 | 24.28664 |
| 6 | Stim | BAGLconnToPos | 0 | 2 | 1 | 99.79453 |
| 7 | Stim | BAGLconnToPos | 0.31743 | 1 | 1 | 0.999505 |
| 1 | StimDetailed | BAGLconnToPos | 0.563378 | 1 | 1 | 0.33389 |
| 2 | StimDetailed | BAGLconnToPos | 0.56322 | 1 | 1 | 0.33416 |
| 3 | StimDetailed | BAGLconnToPos | 0.56468 | 1 | 1 | 0.331666 |
| 4 | StimDetailed | BAGLconnToPos | 0.563349 | 1 | 1 | 0.333939 |
| 5 | StimDetailed | BAGLconnToPos | 0.004468 | 1 | 1 | 8.083296 |
| 6 | StimDetailed | BAGLconnToPos | 7.91E-09 | 1 | 1 | 33.29781 |
| 7 | StimDetailed | BAGLconnToPos | 0.563477 | 1 | 1 | 0.333719 |
| 1 | bhv | BAGLconnToPos3 | 3.33E-16 | 3 | 3 | 75.11481 |
| 2 | bhv | BAGLconnToPos3 | 0.852213 | 1 | 3 | 0.788512 |
| 3 | bhv | BAGLconnToPos3 | 1.49E-07 | 1 | 3 | 34.59181 |
| 4 | bhv | BAGLconnToPos3 | 0 | 3 | 3 | 344.1666 |
| 5 | bhv | BAGLconnToPos3 | 0 | 1 | 3 | 469.5483 |
| 6 | bhv | BAGLconnToPos3 | 0 | 1 | 3 | 1256.707 |
| 7 | bhv | BAGLconnToPos3 | 0 | 3 | 3 | 174.5673 |
| 1 | FWD | BAGLconnToPos3 | 6.11E-15 | 2 | 1 | 60.85144 |
| 2 | FWD | BAGLconnToPos3 | 0.374551 | 1 | 1 | 0.788512 |
| 3 | FWD | BAGLconnToPos3 | 2.54E-08 | 1 | 1 | 31.03315 |
| 4 | FWD | BAGLconnToPos3 | 0 | 2 | 1 | 342.8264 |
| 5 | FWD | BAGLconnToPos3 | 0 | 1 | 1 | 449.5595 |
| 6 | FWD | BAGLconnToPos3 | 0 | 1 | 1 | 756.9948 |
| 7 | FWD | BAGLconnToPos3 | 0 | 2 | 1 | 173.9562 |
| 1 | Stim | BAGLconnToPos3 | 0.482755 | 2 | 1 | 0.492633 |
| 2 | Stim | BAGLconnToPos3 | 0.316951 | 2 | 1 | 1.001489 |
| 3 | Stim | BAGLconnToPos3 | 0.236329 | 1 | 1 | 1.402361 |
| 4 | Stim | BAGLconnToPos3 | 0.003525 | 2 | 1 | 8.51377 |
| 5 | Stim | BAGLconnToPos3 | 0 | 2 | 1 | 659.1007 |
| 6 | Stim | BAGLconnToPos3 | 2.34E-09 | 2 | 1 | 35.66844 |
| 7 | Stim | BAGLconnToPos3 | 0 | 2 | 1 | 130.3942 |
| 1 | StimDetailed | BAGLconnToPos3 | 3.11E-15 | 1 | 1 | 62.16386 |
| 2 | StimDetailed | BAGLconnToPos3 | 0.56322 | 1 | 1 | 0.33416 |
| 3 | StimDetailed | BAGLconnToPos3 | 0.00449 | 1 | 1 | 8.074173 |
| 4 | StimDetailed | BAGLconnToPos3 | 0 | 1 | 1 | 109.5453 |
| 5 | StimDetailed | BAGLconnToPos3 | 0 | 2 | 1 | 560.7455 |
| 6 | StimDetailed | BAGLconnToPos3 | 0.010911 | 1 | 1 | 6.479725 |
| 7 | StimDetailed | BAGLconnToPos3 | 1.67E-10 | 1 | 1 | 40.82013 |
| 1 | bhv | BAGLyhat | 0 | 3 | 3 | 150.0971 |
| 2 | bhv | BAGLyhat | 0 | 1 | 3 | 246.4557 |
| 3 | bhv | BAGLyhat | 0 | 3 | 3 | 1205.009 |
| 4 | bhv | BAGLyhat | 0 | 3 | 3 | 401.6204 |
| 5 | bhv | BAGLyhat | 0 | 1 | 3 | 119.0088 |
| 6 | bhv | BAGLyhat | 5.92E-13 | 3 | 3 | 59.98423 |
| 7 | bhv | BAGLyhat | 0 | 1 | 3 | 85.96362 |
| 1 | FWD | BAGLyhat | 0.082548 | 2 | 1 | 3.014013 |
| 2 | FWD | BAGLyhat | 0 | 2 | 1 | 130.3386 |
| 3 | FWD | BAGLyhat | 0 | 2 | 1 | 1091.556 |
| 4 | FWD | BAGLyhat | 0 | 2 | 1 | 327.1118 |
| 5 | FWD | BAGLyhat | 5.00E-15 | 2 | 1 | 61.26463 |
| 6 | FWD | BAGLyhat | 0.000592 | 2 | 1 | 11.80111 |
| 7 | FWD | BAGLyhat | 1.82E-10 | 2 | 1 | 40.65017 |
| 1 | Stim | BAGLyhat | 1.80E-09 | 2 | 1 | 36.18016 |
| 2 | Stim | BAGLyhat | 4.44E-16 | 2 | 1 | 66.05092 |
| 3 | Stim | BAGLyhat | 0.000431 | 2 | 1 | 12.39183 |
| 4 | Stim | BAGLyhat | 0 | 2 | 1 | 214.551 |
| 5 | Stim | BAGLyhat | 0 | 2 | 1 | 681.7921 |
| 6 | Stim | BAGLyhat | 0 | 2 | 1 | 715.1217 |
| 7 | Stim | BAGLyhat | 0 | 2 | 1 | 203.5076 |
| 1 | StimDetailed | BAGLyhat | 0 | 1 | 1 | 108.9882 |
| 2 | StimDetailed | BAGLyhat | 0.028008 | 1 | 1 | 4.827558 |
| 3 | StimDetailed | BAGLyhat | 0 | 1 | 1 | 170.6217 |
| 4 | StimDetailed | BAGLyhat | 0 | 1 | 1 | 495.1841 |
| 5 | StimDetailed | BAGLyhat | 0 | 1 | 1 | 468.7295 |
| 6 | StimDetailed | BAGLyhat | 0 | 1 | 1 | 522.5093 |
| 7 | StimDetailed | BAGLyhat | 0 | 1 | 1 | 394.984 |
| 1 | bhv | BAGLyhat3 | 1.57E-14 | 1 | 3 | 67.36783 |
| 2 | bhv | BAGLyhat3 | 6.66E-16 | 4 | 3 | 73.78761 |
| 3 | bhv | BAGLyhat3 | 0 | 1 | 3 | 106.9087 |
| 4 | bhv | BAGLyhat3 | 0.000121 | 3 | 3 | 20.71101 |
| 5 | bhv | BAGLyhat3 | 0 | 4 | 3 | 101.5347 |
| 6 | bhv | BAGLyhat3 | 0 | 1 | 3 | 132.938 |
| 7 | bhv | BAGLyhat3 | 0 | 1 | 3 | 97.04037 |
| 1 | FWD | BAGLyhat3 | 1.44E-14 | 1 | 1 | 59.17059 |
| 2 | FWD | BAGLyhat3 | 0.001411 | 2 | 1 | 10.19115 |
| 3 | FWD | BAGLyhat3 | 0.190523 | 1 | 1 | 1.713573 |
| 4 | FWD | BAGLyhat3 | 0.445287 | 2 | 1 | 0.582621 |
| 5 | FWD | BAGLyhat3 | 0.798675 | 2 | 1 | 0.065056 |
| 6 | FWD | BAGLyhat3 | 0 | 1 | 1 | 123.0829 |
| 7 | FWD | BAGLyhat3 | 1.69E-11 | 1 | 1 | 45.29808 |
| 1 | Stim | BAGLyhat3 | 6.88E-15 | 2 | 1 | 60.6274 |
| 2 | Stim | BAGLyhat3 | 0.004616 | 2 | 1 | 8.023896 |
| 3 | Stim | BAGLyhat3 | 1.11E-16 | 2 | 1 | 69.04463 |
| 4 | Stim | BAGLyhat3 | 0 | 2 | 1 | 82.21867 |
| 5 | Stim | BAGLyhat3 | 1.25E-11 | 2 | 1 | 45.89751 |
| 6 | Stim | BAGLyhat3 | 0 | 2 | 1 | 79.97684 |
| 7 | Stim | BAGLyhat3 | 0 | 2 | 1 | 94.29243 |
| 1 | StimDetailed | BAGLyhat3 | 0 | 2 | 1 | 181.6617 |
| 2 | StimDetailed | BAGLyhat3 | 0.998059 | 1 | 1 | 5.92E-06 |
| 3 | StimDetailed | BAGLyhat3 | 0 | 2 | 1 | 208.4596 |
| 4 | StimDetailed | BAGLyhat3 | 0 | 2 | 1 | 246.3304 |
| 5 | StimDetailed | BAGLyhat3 | 0 | 2 | 1 | 137.9012 |
| 6 | StimDetailed | BAGLyhat3 | 0 | 2 | 1 | 239.693 |
| 7 | StimDetailed | BAGLyhat3 | 0 | 2 | 1 | 265.9205 |
| 1 | bhv | BAGRconnFrom | 0.153422 | 3 | 3 | 5.264503 |
| 2 | bhv | BAGRconnFrom | 0.852213 | 1 | 3 | 0.788512 |
| 3 | bhv | BAGRconnFrom | 0.86671 | 1 | 3 | 0.727532 |
| 4 | bhv | BAGRconnFrom | 0.133764 | 3 | 3 | 5.582888 |
| 5 | bhv | BAGRconnFrom | 0.004201 | 3 | 3 | 13.21129 |
| 6 | bhv | BAGRconnFrom | 0 | 3 | 3 | 102.6272 |
| 7 | bhv | BAGRconnFrom | 0.760397 | 3 | 3 | 1.169215 |
| 1 | FWD | BAGRconnFrom | 0.455954 | 2 | 1 | 0.555808 |
| 2 | FWD | BAGRconnFrom | 0.374551 | 1 | 1 | 0.788512 |
| 3 | FWD | BAGRconnFrom | 0.393684 | 1 | 1 | 0.727532 |
| 4 | FWD | BAGRconnFrom | 0.364824 | 2 | 1 | 0.821219 |
| 5 | FWD | BAGRconnFrom | 0.104907 | 2 | 1 | 2.629308 |
| 6 | FWD | BAGRconnFrom | 3.33E-10 | 2 | 1 | 39.46928 |
| 7 | FWD | BAGRconnFrom | 0.390819 | 2 | 1 | 0.736392 |
| 1 | Stim | BAGRconnFrom | 0.3172 | 2 | 1 | 1.000455 |
| 2 | Stim | BAGRconnFrom | 0.316951 | 2 | 1 | 1.001489 |
| 3 | Stim | BAGRconnFrom | 0.31698 | 2 | 1 | 1.001367 |
| 4 | Stim | BAGRconnFrom | 0.157093 | 2 | 1 | 2.001984 |
| 5 | Stim | BAGRconnFrom | 0.045206 | 2 | 1 | 4.010926 |
| 6 | Stim | BAGRconnFrom | 0 | 2 | 1 | 111.9906 |
| 7 | Stim | BAGRconnFrom | 0.31743 | 1 | 1 | 0.999505 |
| 1 | StimDetailed | BAGRconnFrom | 0.563378 | 1 | 1 | 0.33389 |
| 2 | StimDetailed | BAGRconnFrom | 0.56322 | 1 | 1 | 0.33416 |
| 3 | StimDetailed | BAGRconnFrom | 0.56468 | 1 | 1 | 0.331666 |
| 4 | StimDetailed | BAGRconnFrom | 0.413676 | 1 | 1 | 0.66821 |
| 5 | StimDetailed | BAGRconnFrom | 0.247926 | 1 | 1 | 1.334952 |
| 6 | StimDetailed | BAGRconnFrom | 9.79E-10 | 1 | 1 | 37.36721 |
| 7 | StimDetailed | BAGRconnFrom | 0.563477 | 1 | 1 | 0.333719 |
| 1 | bhv | BAGRconnFrom3 | 7.59E-08 | 3 | 3 | 35.97162 |
| 2 | bhv | BAGRconnFrom3 | 0.861703 | 1 | 3 | 0.748663 |
| 3 | bhv | BAGRconnFrom3 | 0 | 3 | 3 | 798.0366 |
| 4 | bhv | BAGRconnFrom3 | 0 | 3 | 3 | 394.8619 |
| 5 | bhv | BAGRconnFrom3 | 0 | 1 | 3 | 292.6434 |
| 6 | bhv | BAGRconnFrom3 | 0 | 1 | 3 | 673.6905 |
| 7 | bhv | BAGRconnFrom3 | 0 | 1 | 3 | 80.29296 |
| 1 | FWD | BAGRconnFrom3 | 1.50E-06 | 2 | 1 | 23.14471 |
| 2 | FWD | BAGRconnFrom3 | 0.865917 | 1 | 1 | 0.028509 |
| 3 | FWD | BAGRconnFrom3 | 0 | 2 | 1 | 796.9195 |
| 4 | FWD | BAGRconnFrom3 | 0 | 2 | 1 | 391.6893 |
| 5 | FWD | BAGRconnFrom3 | 0 | 1 | 1 | 239.5775 |
| 6 | FWD | BAGRconnFrom3 | 0 | 1 | 1 | 330.378 |
| 7 | FWD | BAGRconnFrom3 | 1.07E-13 | 1 | 1 | 55.22841 |
| 1 | Stim | BAGRconnFrom3 | 1.44E-15 | 1 | 1 | 63.77727 |
| 2 | Stim | BAGRconnFrom3 | 0.156888 | 2 | 1 | 2.003971 |
| 3 | Stim | BAGRconnFrom3 | 0 | 1 | 1 | 83.96648 |
| 4 | Stim | BAGRconnFrom3 | 0 | 2 | 1 | 85.08102 |
| 5 | Stim | BAGRconnFrom3 | 0 | 2 | 1 | 404.9418 |
| 6 | Stim | BAGRconnFrom3 | 0 | 2 | 1 | 118.7458 |
| 7 | Stim | BAGRconnFrom3 | 0 | 2 | 1 | 573.3883 |
| 1 | StimDetailed | BAGRconnFrom3 | 1.56E-07 | 1 | 1 | 27.51819 |
| 2 | StimDetailed | BAGRconnFrom3 | 0.413522 | 1 | 1 | 0.668652 |
| 3 | StimDetailed | BAGRconnFrom3 | 1.82E-12 | 1 | 1 | 49.66789 |
| 4 | StimDetailed | BAGRconnFrom3 | 0 | 1 | 1 | 398.2547 |
| 5 | StimDetailed | BAGRconnFrom3 | 0 | 2 | 1 | 869.6359 |
| 6 | StimDetailed | BAGRconnFrom3 | 0 | 1 | 1 | 141.235 |
| 7 | StimDetailed | BAGRconnFrom3 | 0 | 1 | 1 | 425.5057 |
| 1 | bhv | BAGRconnFromNeg | 0.044429 | 3 | 3 | 8.077887 |
| 2 | bhv | BAGRconnFromNeg | NaN | 1 | 3 | NaN |
| 3 | bhv | BAGRconnFromNeg | NaN | 1 | 3 | NaN |
| 4 | bhv | BAGRconnFromNeg | NaN | 1 | 3 | NaN |
| 5 | bhv | BAGRconnFromNeg | 6.26E-12 | 2 | 3 | 55.18754 |
| 6 | bhv | BAGRconnFromNeg | NaN | 1 | 3 | NaN |
| 7 | bhv | BAGRconnFromNeg | NaN | 1 | 3 | NaN |
| 1 | FWD | BAGRconnFromNeg | 0.266248 | 2 | 1 | 1.235974 |
| 2 | FWD | BAGRconnFromNeg | NaN | 1 | 1 | NaN |
| 3 | FWD | BAGRconnFromNeg | NaN | 1 | 1 | NaN |
| 4 | FWD | BAGRconnFromNeg | NaN | 1 | 1 | NaN |
| 5 | FWD | BAGRconnFromNeg | 0.104907 | 2 | 1 | 2.629308 |
| 6 | FWD | BAGRconnFromNeg | NaN | 1 | 1 | NaN |
| 7 | FWD | BAGRconnFromNeg | NaN | 1 | 1 | NaN |
| 1 | Stim | BAGRconnFromNeg | 1.25E-07 | 1 | 1 | 27.93786 |
| 2 | Stim | BAGRconnFromNeg | NaN | 1 | 1 | NaN |
| 3 | Stim | BAGRconnFromNeg | NaN | 1 | 1 | NaN |
| 4 | Stim | BAGRconnFromNeg | NaN | 1 | 1 | NaN |
| 5 | Stim | BAGRconnFromNeg | 0.045206 | 1 | 1 | 4.010926 |
| 6 | Stim | BAGRconnFromNeg | NaN | 1 | 1 | NaN |
| 7 | Stim | BAGRconnFromNeg | NaN | 1 | 1 | NaN |
| 5 | StimDetailed | BAGRconnFromNeg | 0.247926 | 1 | 1 | 1.334952 |
| 7 | StimDetailed | BAGRconnFromNeg | NaN | 1 | 1 | NaN |
| 1 | bhv | BAGRconnFromNeg3 | 3.33E-13 | 3 | 3 | 61.15418 |
| 2 | bhv | BAGRconnFromNeg3 | NaN | 1 | 3 | NaN |
| 3 | bhv | BAGRconnFromNeg3 | NaN | 1 | 3 | NaN |
| 4 | bhv | BAGRconnFromNeg3 | NaN | 1 | 3 | NaN |
| 5 | bhv | BAGRconnFromNeg3 | NaN | 1 | 3 | NaN |
| 6 | bhv | BAGRconnFromNeg3 | NaN | 1 | 3 | NaN |
| 7 | bhv | BAGRconnFromNeg3 | NaN | 1 | 3 | NaN |
| 1 | FWD | BAGRconnFromNeg3 | 5.58E-13 | 2 | 1 | 51.98995 |
| 2 | FWD | BAGRconnFromNeg3 | NaN | 1 | 1 | NaN |
| 3 | FWD | BAGRconnFromNeg3 | NaN | 1 | 1 | NaN |
| 4 | FWD | BAGRconnFromNeg3 | NaN | 1 | 1 | NaN |
| 5 | FWD | BAGRconnFromNeg3 | NaN | 1 | 1 | NaN |
| 6 | FWD | BAGRconnFromNeg3 | NaN | 1 | 1 | NaN |
| 7 | FWD | BAGRconnFromNeg3 | NaN | 1 | 1 | NaN |
| 1 | Stim | BAGRconnFromNeg3 | 0 | 1 | 1 | 116.6196 |
| 2 | Stim | BAGRconnFromNeg3 | NaN | 1 | 1 | NaN |
| 3 | Stim | BAGRconnFromNeg3 | NaN | 1 | 1 | NaN |
| 4 | Stim | BAGRconnFromNeg3 | NaN | 1 | 1 | NaN |
| 5 | Stim | BAGRconnFromNeg3 | NaN | 1 | 1 | NaN |
| 6 | Stim | BAGRconnFromNeg3 | NaN | 1 | 1 | NaN |
| 7 | Stim | BAGRconnFromNeg3 | NaN | 1 | 1 | NaN |
| 5 | StimDetailed | BAGRconnFromNeg3 | NaN | 1 | 1 | NaN |
| 7 | StimDetailed | BAGRconnFromNeg3 | NaN | 1 | 1 | NaN |
| 1 | bhv | BAGRconnFromPos | 0.153422 | 3 | 3 | 5.264503 |
| 2 | bhv | BAGRconnFromPos | 0.852213 | 1 | 3 | 0.788512 |
| 3 | bhv | BAGRconnFromPos | 0.86671 | 1 | 3 | 0.727532 |
| 4 | bhv | BAGRconnFromPos | 0.133764 | 3 | 3 | 5.582888 |
| 5 | bhv | BAGRconnFromPos | 6.90E-05 | 3 | 3 | 21.88323 |
| 6 | bhv | BAGRconnFromPos | 0 | 3 | 3 | 102.6272 |
| 7 | bhv | BAGRconnFromPos | 0.760397 | 3 | 3 | 1.169215 |
| 1 | FWD | BAGRconnFromPos | 0.455954 | 2 | 1 | 0.555808 |
| 2 | FWD | BAGRconnFromPos | 0.374551 | 1 | 1 | 0.788512 |
| 3 | FWD | BAGRconnFromPos | 0.393684 | 1 | 1 | 0.727532 |
| 4 | FWD | BAGRconnFromPos | 0.364824 | 2 | 1 | 0.821219 |
| 5 | FWD | BAGRconnFromPos | 0.031831 | 2 | 1 | 4.607581 |
| 6 | FWD | BAGRconnFromPos | 3.33E-10 | 2 | 1 | 39.46928 |
| 7 | FWD | BAGRconnFromPos | 0.390819 | 2 | 1 | 0.736392 |
| 1 | Stim | BAGRconnFromPos | 0.3172 | 2 | 1 | 1.000455 |
| 2 | Stim | BAGRconnFromPos | 0.316951 | 2 | 1 | 1.001489 |
| 3 | Stim | BAGRconnFromPos | 0.31698 | 2 | 1 | 1.001367 |
| 4 | Stim | BAGRconnFromPos | 0.157093 | 2 | 1 | 2.001984 |
| 5 | Stim | BAGRconnFromPos | 0.008021 | 2 | 1 | 7.028718 |
| 6 | Stim | BAGRconnFromPos | 0 | 2 | 1 | 111.9906 |
| 7 | Stim | BAGRconnFromPos | 0.31743 | 1 | 1 | 0.999505 |
| 1 | StimDetailed | BAGRconnFromPos | 0.563378 | 1 | 1 | 0.33389 |
| 2 | StimDetailed | BAGRconnFromPos | 0.56322 | 1 | 1 | 0.33416 |
| 3 | StimDetailed | BAGRconnFromPos | 0.56468 | 1 | 1 | 0.331666 |
| 4 | StimDetailed | BAGRconnFromPos | 0.413676 | 1 | 1 | 0.66821 |
| 5 | StimDetailed | BAGRconnFromPos | 0.126141 | 1 | 1 | 2.33936 |
| 6 | StimDetailed | BAGRconnFromPos | 9.79E-10 | 1 | 1 | 37.36721 |
| 7 | StimDetailed | BAGRconnFromPos | 0.563477 | 1 | 1 | 0.333719 |
| 1 | bhv | BAGRconnFromPos3 | 3.60E-06 | 3 | 3 | 28.01632 |
| 2 | bhv | BAGRconnFromPos3 | 0.861703 | 1 | 3 | 0.748663 |
| 3 | bhv | BAGRconnFromPos3 | 0 | 3 | 3 | 798.0366 |
| 4 | bhv | BAGRconnFromPos3 | 0 | 3 | 3 | 394.8619 |
| 5 | bhv | BAGRconnFromPos3 | 0 | 1 | 3 | 292.6434 |
| 6 | bhv | BAGRconnFromPos3 | 0 | 1 | 3 | 673.6905 |
| 7 | bhv | BAGRconnFromPos3 | 0 | 1 | 3 | 80.29296 |
| 1 | FWD | BAGRconnFromPos3 | 1.35E-07 | 2 | 1 | 27.79012 |
| 2 | FWD | BAGRconnFromPos3 | 0.865917 | 1 | 1 | 0.028509 |
| 3 | FWD | BAGRconnFromPos3 | 0 | 2 | 1 | 796.9195 |
| 4 | FWD | BAGRconnFromPos3 | 0 | 2 | 1 | 391.6893 |
| 5 | FWD | BAGRconnFromPos3 | 0 | 1 | 1 | 239.5775 |
| 6 | FWD | BAGRconnFromPos3 | 0 | 1 | 1 | 330.378 |
| 7 | FWD | BAGRconnFromPos3 | 1.07E-13 | 1 | 1 | 55.22841 |
| 1 | Stim | BAGRconnFromPos3 | 1.84E-08 | 1 | 1 | 31.65146 |
| 2 | Stim | BAGRconnFromPos3 | 0.156888 | 2 | 1 | 2.003971 |
| 3 | Stim | BAGRconnFromPos3 | 0 | 1 | 1 | 83.96648 |
| 4 | Stim | BAGRconnFromPos3 | 0 | 2 | 1 | 85.08102 |
| 5 | Stim | BAGRconnFromPos3 | 0 | 2 | 1 | 404.9418 |
| 6 | Stim | BAGRconnFromPos3 | 0 | 2 | 1 | 118.7458 |
| 7 | Stim | BAGRconnFromPos3 | 0 | 2 | 1 | 573.3883 |
| 1 | StimDetailed | BAGRconnFromPos3 | 6.48E-09 | 1 | 1 | 33.68641 |
| 2 | StimDetailed | BAGRconnFromPos3 | 0.413522 | 1 | 1 | 0.668652 |
| 3 | StimDetailed | BAGRconnFromPos3 | 1.82E-12 | 1 | 1 | 49.66789 |
| 4 | StimDetailed | BAGRconnFromPos3 | 0 | 1 | 1 | 398.2547 |
| 5 | StimDetailed | BAGRconnFromPos3 | 0 | 2 | 1 | 869.6359 |
| 6 | StimDetailed | BAGRconnFromPos3 | 0 | 1 | 1 | 141.235 |
| 7 | StimDetailed | BAGRconnFromPos3 | 0 | 1 | 1 | 425.5057 |
| 1 | bhv | BAGRconnTo | 0.153422 | 3 | 3 | 5.264503 |
| 2 | bhv | BAGRconnTo | 0.852213 | 1 | 3 | 0.788512 |
| 3 | bhv | BAGRconnTo | 0.86671 | 1 | 3 | 0.727532 |
| 4 | bhv | BAGRconnTo | 0.007016 | 3 | 3 | 12.1099 |
| 5 | bhv | BAGRconnTo | 0.004201 | 3 | 3 | 13.21129 |
| 6 | bhv | BAGRconnTo | 0.821141 | 3 | 3 | 0.917755 |
| 7 | bhv | BAGRconnTo | 0.760397 | 3 | 3 | 1.169215 |
| 1 | FWD | BAGRconnTo | 0.455954 | 2 | 1 | 0.555808 |
| 2 | FWD | BAGRconnTo | 0.374551 | 1 | 1 | 0.788512 |
| 3 | FWD | BAGRconnTo | 0.393684 | 1 | 1 | 0.727532 |
| 4 | FWD | BAGRconnTo | 0.521763 | 2 | 1 | 0.410406 |
| 5 | FWD | BAGRconnTo | 0.104907 | 2 | 1 | 2.629308 |
| 6 | FWD | BAGRconnTo | 0.552443 | 2 | 1 | 0.352958 |
| 7 | FWD | BAGRconnTo | 0.390819 | 2 | 1 | 0.736392 |
| 1 | Stim | BAGRconnTo | 0.3172 | 2 | 1 | 1.000455 |
| 2 | Stim | BAGRconnTo | 0.316951 | 2 | 1 | 1.001489 |
| 3 | Stim | BAGRconnTo | 0.31698 | 2 | 1 | 1.001367 |
| 4 | Stim | BAGRconnTo | 0.317191 | 2 | 1 | 1.000496 |
| 5 | Stim | BAGRconnTo | 0.045206 | 2 | 1 | 4.010926 |
| 6 | Stim | BAGRconnTo | 0.316951 | 2 | 1 | 1.001489 |
| 7 | Stim | BAGRconnTo | 0.31743 | 1 | 1 | 0.999505 |
| 1 | StimDetailed | BAGRconnTo | 0.563378 | 1 | 1 | 0.33389 |
| 2 | StimDetailed | BAGRconnTo | 0.56322 | 1 | 1 | 0.33416 |
| 3 | StimDetailed | BAGRconnTo | 0.56468 | 1 | 1 | 0.331666 |
| 4 | StimDetailed | BAGRconnTo | 0.563349 | 1 | 1 | 0.333939 |
| 5 | StimDetailed | BAGRconnTo | 0.247926 | 1 | 1 | 1.334952 |
| 6 | StimDetailed | BAGRconnTo | 0.56322 | 1 | 1 | 0.33416 |
| 7 | StimDetailed | BAGRconnTo | 0.563477 | 1 | 1 | 0.333719 |
| 1 | bhv | BAGRconnTo3 | 2.37E-06 | 3 | 3 | 28.88709 |
| 2 | bhv | BAGRconnTo3 | 0.852213 | 1 | 3 | 0.788512 |
| 3 | bhv | BAGRconnTo3 | 0.034043 | 1 | 3 | 8.668266 |
| 4 | bhv | BAGRconnTo3 | 0 | 3 | 3 | 815.3059 |
| 5 | bhv | BAGRconnTo3 | 0 | 1 | 3 | 266.4603 |
| 6 | bhv | BAGRconnTo3 | 0 | 1 | 3 | 1313.302 |
| 7 | bhv | BAGRconnTo3 | 0 | 3 | 3 | 232.6008 |
| 1 | FWD | BAGRconnTo3 | 2.15E-07 | 2 | 1 | 26.89487 |
| 2 | FWD | BAGRconnTo3 | 0.374551 | 1 | 1 | 0.788512 |
| 3 | FWD | BAGRconnTo3 | 0.007863 | 1 | 1 | 7.064456 |
| 4 | FWD | BAGRconnTo3 | 0 | 2 | 1 | 805.3392 |
| 5 | FWD | BAGRconnTo3 | 0 | 1 | 1 | 240.268 |
| 6 | FWD | BAGRconnTo3 | 0 | 1 | 1 | 771.4798 |
| 7 | FWD | BAGRconnTo3 | 0 | 2 | 1 | 232.5979 |
| 1 | Stim | BAGRconnTo3 | 4.21E-08 | 1 | 1 | 30.05113 |
| 2 | Stim | BAGRconnTo3 | 0.316951 | 2 | 1 | 1.001489 |
| 3 | Stim | BAGRconnTo3 | 0.007374 | 1 | 1 | 7.179508 |
| 4 | Stim | BAGRconnTo3 | 0.308309 | 2 | 1 | 1.037908 |
| 5 | Stim | BAGRconnTo3 | 0 | 2 | 1 | 640.8911 |
| 6 | Stim | BAGRconnTo3 | 1.96E-08 | 2 | 1 | 31.53523 |
| 7 | Stim | BAGRconnTo3 | 0 | 2 | 1 | 174.7778 |
| 1 | StimDetailed | BAGRconnTo3 | 1.04E-08 | 1 | 1 | 32.77293 |
| 2 | StimDetailed | BAGRconnTo3 | 0.56322 | 1 | 1 | 0.33416 |
| 3 | StimDetailed | BAGRconnTo3 | 0.000628 | 1 | 1 | 11.69025 |
| 4 | StimDetailed | BAGRconnTo3 | 0 | 1 | 1 | 140.133 |
| 5 | StimDetailed | BAGRconnTo3 | 0 | 2 | 1 | 599.8086 |
| 6 | StimDetailed | BAGRconnTo3 | 0.006914 | 1 | 1 | 7.295128 |
| 7 | StimDetailed | BAGRconnTo3 | 2.19E-14 | 1 | 1 | 58.35553 |
| 1 | bhv | BAGRconnToNeg | 0.000985 | 3 | 3 | 16.29781 |
| 2 | bhv | BAGRconnToNeg | NaN | 1 | 3 | NaN |
| 3 | bhv | BAGRconnToNeg | NaN | 1 | 3 | NaN |
| 4 | bhv | BAGRconnToNeg | NaN | 1 | 3 | NaN |
| 5 | bhv | BAGRconnToNeg | 1.10E-13 | 4 | 3 | 63.41412 |
| 6 | bhv | BAGRconnToNeg | NaN | 1 | 3 | NaN |
| 7 | bhv | BAGRconnToNeg | 2.41E-14 | 1 | 3 | 66.48456 |
| 1 | FWD | BAGRconnToNeg | 0.002475 | 2 | 1 | 9.158899 |
| 2 | FWD | BAGRconnToNeg | NaN | 1 | 1 | NaN |
| 3 | FWD | BAGRconnToNeg | NaN | 1 | 1 | NaN |
| 4 | FWD | BAGRconnToNeg | NaN | 1 | 1 | NaN |
| 5 | FWD | BAGRconnToNeg | 6.25E-06 | 2 | 1 | 20.40971 |
| 6 | FWD | BAGRconnToNeg | NaN | 1 | 1 | NaN |
| 7 | FWD | BAGRconnToNeg | 0.023225 | 1 | 1 | 5.151569 |
| 1 | Stim | BAGRconnToNeg | 4.60E-12 | 1 | 1 | 47.85242 |
| 2 | Stim | BAGRconnToNeg | NaN | 1 | 1 | NaN |
| 3 | Stim | BAGRconnToNeg | NaN | 1 | 1 | NaN |
| 4 | Stim | BAGRconnToNeg | NaN | 1 | 1 | NaN |
| 5 | Stim | BAGRconnToNeg | 0.004038 | 1 | 1 | 8.2666 |
| 6 | Stim | BAGRconnToNeg | NaN | 1 | 1 | NaN |
| 7 | Stim | BAGRconnToNeg | 5.66E-11 | 2 | 1 | 42.93477 |
| 5 | StimDetailed | BAGRconnToNeg | 0.711202 | 1 | 1 | 0.13708 |
| 5 | StimDetailed | BAGRconnToNeg | 0.711202 | 1 | 1 | 0.13708 |
| 7 | StimDetailed | BAGRconnToNeg | 0 | 2 | 1 | 128.5918 |
| 7 | StimDetailed | BAGRconnToNeg | 0 | 2 | 1 | 128.5918 |
| 1 | bhv | BAGRconnToNeg3 | 3.59E-13 | 3 | 3 | 61.00187 |
| 2 | bhv | BAGRconnToNeg3 | NaN | 1 | 3 | NaN |
| 3 | bhv | BAGRconnToNeg3 | NaN | 1 | 3 | NaN |
| 4 | bhv | BAGRconnToNeg3 | NaN | 1 | 3 | NaN |
| 5 | bhv | BAGRconnToNeg3 | 2.48E-08 | 4 | 3 | 38.27155 |
| 6 | bhv | BAGRconnToNeg3 | NaN | 1 | 3 | NaN |
| 7 | bhv | BAGRconnToNeg3 | NaN | 1 | 3 | NaN |
| 1 | FWD | BAGRconnToNeg3 | 4.11E-13 | 2 | 1 | 52.5879 |
| 2 | FWD | BAGRconnToNeg3 | NaN | 1 | 1 | NaN |
| 3 | FWD | BAGRconnToNeg3 | NaN | 1 | 1 | NaN |
| 4 | FWD | BAGRconnToNeg3 | NaN | 1 | 1 | NaN |
| 5 | FWD | BAGRconnToNeg3 | 0.04694 | 2 | 1 | 3.947556 |
| 6 | FWD | BAGRconnToNeg3 | NaN | 1 | 1 | NaN |
| 7 | FWD | BAGRconnToNeg3 | NaN | 1 | 1 | NaN |
| 1 | Stim | BAGRconnToNeg3 | 0 | 1 | 1 | 117.7325 |
| 2 | Stim | BAGRconnToNeg3 | NaN | 1 | 1 | NaN |
| 3 | Stim | BAGRconnToNeg3 | NaN | 1 | 1 | NaN |
| 4 | Stim | BAGRconnToNeg3 | NaN | 1 | 1 | NaN |
| 5 | Stim | BAGRconnToNeg3 | 0.014217 | 2 | 1 | 6.010936 |
| 6 | Stim | BAGRconnToNeg3 | NaN | 1 | 1 | NaN |
| 7 | Stim | BAGRconnToNeg3 | NaN | 1 | 1 | NaN |
| 5 | StimDetailed | BAGRconnToNeg3 | 2.14E-05 | 2 | 1 | 18.06014 |
| 5 | StimDetailed | BAGRconnToNeg3 | 2.14E-05 | 2 | 1 | 18.06014 |
| 7 | StimDetailed | BAGRconnToNeg3 | NaN | 1 | 1 | NaN |
| 7 | StimDetailed | BAGRconnToNeg3 | NaN | 1 | 1 | NaN |
| 1 | bhv | BAGRconnToPos | 0.153422 | 3 | 3 | 5.264503 |
| 2 | bhv | BAGRconnToPos | 0.852213 | 1 | 3 | 0.788512 |
| 3 | bhv | BAGRconnToPos | 0.86671 | 1 | 3 | 0.727532 |
| 4 | bhv | BAGRconnToPos | 0.007016 | 3 | 3 | 12.1099 |
| 5 | bhv | BAGRconnToPos | 2.59E-12 | 3 | 3 | 56.98301 |
| 6 | bhv | BAGRconnToPos | 0.821141 | 3 | 3 | 0.917755 |
| 7 | bhv | BAGRconnToPos | 0.760397 | 3 | 3 | 1.169215 |
| 1 | FWD | BAGRconnToPos | 0.455954 | 2 | 1 | 0.555808 |
| 2 | FWD | BAGRconnToPos | 0.374551 | 1 | 1 | 0.788512 |
| 3 | FWD | BAGRconnToPos | 0.393684 | 1 | 1 | 0.727532 |
| 4 | FWD | BAGRconnToPos | 0.521763 | 2 | 1 | 0.410406 |
| 5 | FWD | BAGRconnToPos | 0.002105 | 2 | 1 | 9.45589 |
| 6 | FWD | BAGRconnToPos | 0.552443 | 2 | 1 | 0.352958 |
| 7 | FWD | BAGRconnToPos | 0.390819 | 2 | 1 | 0.736392 |
| 1 | Stim | BAGRconnToPos | 0.3172 | 2 | 1 | 1.000455 |
| 2 | Stim | BAGRconnToPos | 0.316951 | 2 | 1 | 1.001489 |
| 3 | Stim | BAGRconnToPos | 0.31698 | 2 | 1 | 1.001367 |
| 4 | Stim | BAGRconnToPos | 0.317191 | 2 | 1 | 1.000496 |
| 5 | Stim | BAGRconnToPos | 1.19E-05 | 2 | 1 | 19.18287 |
| 6 | Stim | BAGRconnToPos | 0.316951 | 2 | 1 | 1.001489 |
| 7 | Stim | BAGRconnToPos | 0.31743 | 1 | 1 | 0.999505 |
| 1 | StimDetailed | BAGRconnToPos | 0.563378 | 1 | 1 | 0.33389 |
| 2 | StimDetailed | BAGRconnToPos | 0.56322 | 1 | 1 | 0.33416 |
| 3 | StimDetailed | BAGRconnToPos | 0.56468 | 1 | 1 | 0.331666 |
| 4 | StimDetailed | BAGRconnToPos | 0.563349 | 1 | 1 | 0.333939 |
| 5 | StimDetailed | BAGRconnToPos | 0.011511 | 1 | 1 | 6.384612 |
| 6 | StimDetailed | BAGRconnToPos | 0.56322 | 1 | 1 | 0.33416 |
| 7 | StimDetailed | BAGRconnToPos | 0.563477 | 1 | 1 | 0.333719 |
| 1 | bhv | BAGRconnToPos3 | 1.84E-08 | 3 | 3 | 38.87753 |
| 2 | bhv | BAGRconnToPos3 | 0.852213 | 1 | 3 | 0.788512 |
| 3 | bhv | BAGRconnToPos3 | 0.034043 | 1 | 3 | 8.668266 |
| 4 | bhv | BAGRconnToPos3 | 0 | 3 | 3 | 815.3059 |
| 5 | bhv | BAGRconnToPos3 | 0 | 1 | 3 | 266.4603 |
| 6 | bhv | BAGRconnToPos3 | 0 | 1 | 3 | 1313.302 |
| 7 | bhv | BAGRconnToPos3 | 0 | 3 | 3 | 232.6008 |
| 1 | FWD | BAGRconnToPos3 | 6.32E-08 | 2 | 1 | 29.2641 |
| 2 | FWD | BAGRconnToPos3 | 0.374551 | 1 | 1 | 0.788512 |
| 3 | FWD | BAGRconnToPos3 | 0.007863 | 1 | 1 | 7.064456 |
| 4 | FWD | BAGRconnToPos3 | 0 | 2 | 1 | 805.3392 |
| 5 | FWD | BAGRconnToPos3 | 0 | 1 | 1 | 240.268 |
| 6 | FWD | BAGRconnToPos3 | 0 | 1 | 1 | 771.4798 |
| 7 | FWD | BAGRconnToPos3 | 0 | 2 | 1 | 232.5979 |
| 1 | Stim | BAGRconnToPos3 | 0.001868 | 1 | 1 | 9.67507 |
| 2 | Stim | BAGRconnToPos3 | 0.316951 | 2 | 1 | 1.001489 |
| 3 | Stim | BAGRconnToPos3 | 0.007374 | 1 | 1 | 7.179508 |
| 4 | Stim | BAGRconnToPos3 | 0.308309 | 2 | 1 | 1.037908 |
| 5 | Stim | BAGRconnToPos3 | 0 | 2 | 1 | 640.8911 |
| 6 | Stim | BAGRconnToPos3 | 1.96E-08 | 2 | 1 | 31.53523 |
| 7 | Stim | BAGRconnToPos3 | 0 | 2 | 1 | 174.7778 |
| 1 | StimDetailed | BAGRconnToPos3 | 9.18E-11 | 1 | 1 | 41.98804 |
| 2 | StimDetailed | BAGRconnToPos3 | 0.56322 | 1 | 1 | 0.33416 |
| 3 | StimDetailed | BAGRconnToPos3 | 0.000628 | 1 | 1 | 11.69025 |
| 4 | StimDetailed | BAGRconnToPos3 | 0 | 1 | 1 | 140.133 |
| 5 | StimDetailed | BAGRconnToPos3 | 0 | 2 | 1 | 599.8086 |
| 6 | StimDetailed | BAGRconnToPos3 | 0.006914 | 1 | 1 | 7.295128 |
| 7 | StimDetailed | BAGRconnToPos3 | 2.19E-14 | 1 | 1 | 58.35553 |
| 1 | bhv | BAGRyhat | 0 | 1 | 3 | 133.4093 |
| 2 | bhv | BAGRyhat | 0 | 3 | 3 | 571.557 |
| 3 | bhv | BAGRyhat | 0 | 3 | 3 | 1214.179 |
| 4 | bhv | BAGRyhat | 0 | 3 | 3 | 524.8399 |
| 5 | bhv | BAGRyhat | 3.49E-13 | 3 | 3 | 61.06041 |
| 6 | bhv | BAGRyhat | 0 | 3 | 3 | 120.3225 |
| 7 | bhv | BAGRyhat | 1.47E-09 | 3 | 3 | 44.04825 |
| 1 | FWD | BAGRyhat | 2.35E-05 | 2 | 1 | 17.88393 |
| 2 | FWD | BAGRyhat | 0 | 2 | 1 | 397.5158 |
| 3 | FWD | BAGRyhat | 0 | 2 | 1 | 1091.556 |
| 4 | FWD | BAGRyhat | 0 | 2 | 1 | 378.6382 |
| 5 | FWD | BAGRyhat | 0.094266 | 2 | 1 | 2.799979 |
| 6 | FWD | BAGRyhat | 0.00011 | 2 | 1 | 14.95702 |
| 7 | FWD | BAGRyhat | 0.271371 | 2 | 1 | 1.209805 |
| 1 | Stim | BAGRyhat | 6.00E-07 | 2 | 1 | 24.91174 |
| 2 | Stim | BAGRyhat | 0 | 1 | 1 | 131.4951 |
| 3 | Stim | BAGRyhat | 0.020047 | 2 | 1 | 5.407786 |
| 4 | Stim | BAGRyhat | 0 | 2 | 1 | 137.1041 |
| 5 | Stim | BAGRyhat | 0 | 2 | 1 | 419.5913 |
| 6 | Stim | BAGRyhat | 0 | 2 | 1 | 439.0087 |
| 7 | Stim | BAGRyhat | 0 | 2 | 1 | 229.6988 |
| 1 | StimDetailed | BAGRyhat | 0 | 1 | 1 | 169.0113 |
| 2 | StimDetailed | BAGRyhat | 0.43315 | 1 | 1 | 0.614361 |
| 3 | StimDetailed | BAGRyhat | 0 | 1 | 1 | 162.9758 |
| 4 | StimDetailed | BAGRyhat | 0 | 1 | 1 | 472.5759 |
| 5 | StimDetailed | BAGRyhat | 0 | 1 | 1 | 539.4291 |
| 6 | StimDetailed | BAGRyhat | 0 | 1 | 1 | 463.2034 |
| 7 | StimDetailed | BAGRyhat | 0 | 1 | 1 | 476.4153 |
| 1 | bhv | BAGRyhat3 | 6.99E-15 | 1 | 3 | 68.98668 |
| 2 | bhv | BAGRyhat3 | NaN | 1 | 3 | NaN |
| 3 | bhv | BAGRyhat3 | 0 | 1 | 3 | 108.2079 |
| 4 | bhv | BAGRyhat3 | 9.75E-05 | 3 | 3 | 21.16052 |
| 5 | bhv | BAGRyhat3 | 0 | 4 | 3 | 106.5017 |
| 6 | bhv | BAGRyhat3 | 0 | 1 | 3 | 115.6404 |
| 7 | bhv | BAGRyhat3 | 0 | 1 | 3 | 104.023 |
| 1 | FWD | BAGRyhat3 | 5.88E-15 | 1 | 1 | 60.92229 |
| 2 | FWD | BAGRyhat3 | NaN | 1 | 1 | NaN |
| 3 | FWD | BAGRyhat3 | 0.225503 | 1 | 1 | 1.469003 |
| 4 | FWD | BAGRyhat3 | 0.962007 | 2 | 1 | 0.002269 |
| 5 | FWD | BAGRyhat3 | 0.521478 | 2 | 1 | 0.410969 |
| 6 | FWD | BAGRyhat3 | 0 | 1 | 1 | 103.5235 |
| 7 | FWD | BAGRyhat3 | 9.35E-11 | 1 | 1 | 41.95374 |
| 1 | Stim | BAGRyhat3 | 4.00E-15 | 2 | 1 | 61.68382 |
| 2 | Stim | BAGRyhat3 | NaN | 1 | 1 | NaN |
| 3 | Stim | BAGRyhat3 | 1.11E-16 | 2 | 1 | 67.9822 |
| 4 | Stim | BAGRyhat3 | 0 | 2 | 1 | 77.89501 |
| 5 | Stim | BAGRyhat3 | 3.61E-11 | 2 | 1 | 43.81698 |
| 6 | Stim | BAGRyhat3 | 0 | 2 | 1 | 74.5913 |
| 7 | Stim | BAGRyhat3 | 0 | 2 | 1 | 96.48785 |
| 1 | StimDetailed | BAGRyhat3 | 0 | 2 | 1 | 184.8271 |
| 2 | StimDetailed | BAGRyhat3 | NaN | 1 | 1 | NaN |
| 3 | StimDetailed | BAGRyhat3 | 0 | 2 | 1 | 205.2519 |
| 4 | StimDetailed | BAGRyhat3 | 0 | 2 | 1 | 233.3765 |
| 5 | StimDetailed | BAGRyhat3 | 0 | 2 | 1 | 131.6502 |
| 6 | StimDetailed | BAGRyhat3 | 0 | 2 | 1 | 223.5523 |
| 7 | StimDetailed | BAGRyhat3 | 0 | 2 | 1 | 272.4734 |
| 2 | bhv | RIGLconnFrom3 | 0 | 1 | 3 | 112.807 |
| 3 | bhv | RIGLconnFrom3 | 0 | 1 | 3 | 285.2349 |
| 4 | bhv | RIGLconnFrom3 | 0 | 3 | 3 | 160.4864 |
| 6 | bhv | RIGLconnFrom3 | 0 | 3 | 3 | 199.3679 |
| 2 | FWD | RIGLconnFrom3 | 0 | 1 | 1 | 112.7293 |
| 3 | FWD | RIGLconnFrom3 | 0 | 1 | 1 | 240.2124 |
| 4 | FWD | RIGLconnFrom3 | 0 | 2 | 1 | 76.63883 |
| 6 | FWD | RIGLconnFrom3 | 0 | 2 | 1 | 102.9789 |
| 2 | Stim | RIGLconnFrom3 | 0 | 1 | 1 | 141.0149 |
| 3 | Stim | RIGLconnFrom3 | 1.54E-14 | 1 | 1 | 59.04495 |
| 4 | Stim | RIGLconnFrom3 | 4.76E-07 | 1 | 1 | 25.35942 |
| 6 | Stim | RIGLconnFrom3 | 0 | 1 | 1 | 278.0465 |
| 2 | StimDetailed | RIGLconnFrom3 | 0.014779 | 1 | 1 | 5.942574 |
| 3 | StimDetailed | RIGLconnFrom3 | 1.78E-05 | 1 | 1 | 18.4078 |
| 4 | StimDetailed | RIGLconnFrom3 | 0.683721 | 1 | 1 | 0.165968 |
| 6 | StimDetailed | RIGLconnFrom3 | 0 | 1 | 1 | 80.89194 |
| 2 | bhv | RIGLconnFromNeg | NaN | 1 | 3 | NaN |
| 3 | bhv | RIGLconnFromNeg | NaN | 1 | 3 | NaN |
| 4 | bhv | RIGLconnFromNeg | NaN | 1 | 3 | NaN |
| 6 | bhv | RIGLconnFromNeg | NaN | 1 | 3 | NaN |
| 2 | FWD | RIGLconnFromNeg | NaN | 1 | 1 | NaN |
| 3 | FWD | RIGLconnFromNeg | NaN | 1 | 1 | NaN |
| 4 | FWD | RIGLconnFromNeg | NaN | 1 | 1 | NaN |
| 6 | FWD | RIGLconnFromNeg | NaN | 1 | 1 | NaN |
| 2 | Stim | RIGLconnFromNeg | NaN | 1 | 1 | NaN |
| 3 | Stim | RIGLconnFromNeg | NaN | 1 | 1 | NaN |
| 4 | Stim | RIGLconnFromNeg | NaN | 1 | 1 | NaN |
| 6 | Stim | RIGLconnFromNeg | NaN | 1 | 1 | NaN |
| 2 | bhv | RIGLconnFromNeg3 | NaN | 1 | 3 | NaN |
| 3 | bhv | RIGLconnFromNeg3 | NaN | 1 | 3 | NaN |
| 4 | bhv | RIGLconnFromNeg3 | NaN | 1 | 3 | NaN |
| 6 | bhv | RIGLconnFromNeg3 | NaN | 1 | 3 | NaN |
| 2 | FWD | RIGLconnFromNeg3 | NaN | 1 | 1 | NaN |
| 3 | FWD | RIGLconnFromNeg3 | NaN | 1 | 1 | NaN |
| 4 | FWD | RIGLconnFromNeg3 | NaN | 1 | 1 | NaN |
| 6 | FWD | RIGLconnFromNeg3 | NaN | 1 | 1 | NaN |
| 2 | Stim | RIGLconnFromNeg3 | NaN | 1 | 1 | NaN |
| 3 | Stim | RIGLconnFromNeg3 | NaN | 1 | 1 | NaN |
| 4 | Stim | RIGLconnFromNeg3 | NaN | 1 | 1 | NaN |
| 6 | Stim | RIGLconnFromNeg3 | NaN | 1 | 1 | NaN |
| 2 | bhv | RIGLconnFromPos | 0.852213 | 1 | 3 | 0.788512 |
| 3 | bhv | RIGLconnFromPos | 0.010611 | 1 | 3 | 11.21651 |
| 4 | bhv | RIGLconnFromPos | 0.007016 | 3 | 3 | 12.1099 |
| 6 | bhv | RIGLconnFromPos | 0.821141 | 3 | 3 | 0.917755 |
| 2 | FWD | RIGLconnFromPos | 0.374551 | 1 | 1 | 0.788512 |
| 3 | FWD | RIGLconnFromPos | 0.820963 | 1 | 1 | 0.051215 |
| 4 | FWD | RIGLconnFromPos | 0.521763 | 2 | 1 | 0.410406 |
| 6 | FWD | RIGLconnFromPos | 0.552443 | 2 | 1 | 0.352958 |
| 2 | Stim | RIGLconnFromPos | 0.316951 | 2 | 1 | 1.001489 |
| 3 | Stim | RIGLconnFromPos | 0.156921 | 2 | 1 | 2.003645 |
| 4 | Stim | RIGLconnFromPos | 0.317191 | 2 | 1 | 1.000496 |
| 6 | Stim | RIGLconnFromPos | 0.316951 | 2 | 1 | 1.001489 |
| 2 | StimDetailed | RIGLconnFromPos | 0.56322 | 1 | 1 | 0.33416 |
| 3 | StimDetailed | RIGLconnFromPos | 0.41528 | 1 | 1 | 0.663634 |
| 4 | StimDetailed | RIGLconnFromPos | 0.563349 | 1 | 1 | 0.333939 |
| 6 | StimDetailed | RIGLconnFromPos | 0.56322 | 1 | 1 | 0.33416 |
| 2 | bhv | RIGLconnFromPos3 | 0 | 1 | 3 | 112.807 |
| 3 | bhv | RIGLconnFromPos3 | 0 | 1 | 3 | 285.2349 |
| 4 | bhv | RIGLconnFromPos3 | 0 | 3 | 3 | 160.4864 |
| 6 | bhv | RIGLconnFromPos3 | 0 | 3 | 3 | 199.3679 |
| 2 | FWD | RIGLconnFromPos3 | 0 | 1 | 1 | 112.7293 |
| 3 | FWD | RIGLconnFromPos3 | 0 | 1 | 1 | 240.2124 |
| 4 | FWD | RIGLconnFromPos3 | 0 | 2 | 1 | 76.63883 |
| 6 | FWD | RIGLconnFromPos3 | 0 | 2 | 1 | 102.9789 |
| 2 | Stim | RIGLconnFromPos3 | 0 | 1 | 1 | 141.0149 |
| 3 | Stim | RIGLconnFromPos3 | 1.54E-14 | 1 | 1 | 59.04495 |
| 4 | Stim | RIGLconnFromPos3 | 4.76E-07 | 1 | 1 | 25.35942 |
| 6 | Stim | RIGLconnFromPos3 | 0 | 1 | 1 | 278.0465 |
| 2 | StimDetailed | RIGLconnFromPos3 | 0.014779 | 1 | 1 | 5.942574 |
| 3 | StimDetailed | RIGLconnFromPos3 | 1.78E-05 | 1 | 1 | 18.4078 |
| 4 | StimDetailed | RIGLconnFromPos3 | 0.683721 | 1 | 1 | 0.165968 |
| 6 | StimDetailed | RIGLconnFromPos3 | 0 | 1 | 1 | 80.89194 |
| 2 | bhv | RIGLconnTo3 | 0.852213 | 1 | 3 | 0.788512 |
| 3 | bhv | RIGLconnTo3 | 0.86671 | 1 | 3 | 0.727532 |
| 4 | bhv | RIGLconnTo3 | 0.007016 | 3 | 3 | 12.1099 |
| 6 | bhv | RIGLconnTo3 | 1.47E-08 | 3 | 3 | 39.34592 |
| 2 | FWD | RIGLconnTo3 | 0.374551 | 1 | 1 | 0.788512 |
| 3 | FWD | RIGLconnTo3 | 0.393684 | 1 | 1 | 0.727532 |
| 4 | FWD | RIGLconnTo3 | 0.521763 | 2 | 1 | 0.410406 |
| 6 | FWD | RIGLconnTo3 | 0.0001 | 2 | 1 | 15.132 |
| 2 | Stim | RIGLconnTo3 | 0.316951 | 2 | 1 | 1.001489 |
| 3 | Stim | RIGLconnTo3 | 0.31698 | 2 | 1 | 1.001367 |
| 4 | Stim | RIGLconnTo3 | 0.317191 | 2 | 1 | 1.000496 |
| 6 | Stim | RIGLconnTo3 | 4.54E-10 | 1 | 1 | 38.86488 |
| 2 | StimDetailed | RIGLconnTo3 | 0.56322 | 1 | 1 | 0.33416 |
| 3 | StimDetailed | RIGLconnTo3 | 0.56468 | 1 | 1 | 0.331666 |
| 4 | StimDetailed | RIGLconnTo3 | 0.563349 | 1 | 1 | 0.333939 |
| 6 | StimDetailed | RIGLconnTo3 | 0.048377 | 1 | 1 | 3.89685 |
| 2 | bhv | RIGLconnToNeg | NaN | 1 | 3 | NaN |
| 3 | bhv | RIGLconnToNeg | NaN | 1 | 3 | NaN |
| 4 | bhv | RIGLconnToNeg | NaN | 1 | 3 | NaN |
| 6 | bhv | RIGLconnToNeg | NaN | 1 | 3 | NaN |
| 2 | FWD | RIGLconnToNeg | NaN | 1 | 1 | NaN |
| 3 | FWD | RIGLconnToNeg | NaN | 1 | 1 | NaN |
| 4 | FWD | RIGLconnToNeg | NaN | 1 | 1 | NaN |
| 6 | FWD | RIGLconnToNeg | NaN | 1 | 1 | NaN |
| 2 | Stim | RIGLconnToNeg | NaN | 1 | 1 | NaN |
| 3 | Stim | RIGLconnToNeg | NaN | 1 | 1 | NaN |
| 4 | Stim | RIGLconnToNeg | NaN | 1 | 1 | NaN |
| 6 | Stim | RIGLconnToNeg | NaN | 1 | 1 | NaN |
| 2 | bhv | RIGLconnToNeg3 | NaN | 1 | 3 | NaN |
| 3 | bhv | RIGLconnToNeg3 | NaN | 1 | 3 | NaN |
| 4 | bhv | RIGLconnToNeg3 | NaN | 1 | 3 | NaN |
| 6 | bhv | RIGLconnToNeg3 | NaN | 1 | 3 | NaN |
| 2 | FWD | RIGLconnToNeg3 | NaN | 1 | 1 | NaN |
| 3 | FWD | RIGLconnToNeg3 | NaN | 1 | 1 | NaN |
| 4 | FWD | RIGLconnToNeg3 | NaN | 1 | 1 | NaN |
| 6 | FWD | RIGLconnToNeg3 | NaN | 1 | 1 | NaN |
| 2 | Stim | RIGLconnToNeg3 | NaN | 1 | 1 | NaN |
| 3 | Stim | RIGLconnToNeg3 | NaN | 1 | 1 | NaN |
| 4 | Stim | RIGLconnToNeg3 | NaN | 1 | 1 | NaN |
| 6 | Stim | RIGLconnToNeg3 | NaN | 1 | 1 | NaN |
| 2 | bhv | RIGLconnToPos | 0.852213 | 1 | 3 | 0.788512 |
| 3 | bhv | RIGLconnToPos | 0.86671 | 1 | 3 | 0.727532 |
| 4 | bhv | RIGLconnToPos | 0.007016 | 3 | 3 | 12.1099 |
| 6 | bhv | RIGLconnToPos | 0.821141 | 3 | 3 | 0.917755 |
| 2 | FWD | RIGLconnToPos | 0.374551 | 1 | 1 | 0.788512 |
| 3 | FWD | RIGLconnToPos | 0.393684 | 1 | 1 | 0.727532 |
| 4 | FWD | RIGLconnToPos | 0.521763 | 2 | 1 | 0.410406 |
| 6 | FWD | RIGLconnToPos | 0.552443 | 2 | 1 | 0.352958 |
| 2 | Stim | RIGLconnToPos | 0.316951 | 2 | 1 | 1.001489 |
| 3 | Stim | RIGLconnToPos | 0.31698 | 2 | 1 | 1.001367 |
| 4 | Stim | RIGLconnToPos | 0.317191 | 2 | 1 | 1.000496 |
| 6 | Stim | RIGLconnToPos | 0.316951 | 2 | 1 | 1.001489 |
| 2 | StimDetailed | RIGLconnToPos | 0.56322 | 1 | 1 | 0.33416 |
| 3 | StimDetailed | RIGLconnToPos | 0.56468 | 1 | 1 | 0.331666 |
| 4 | StimDetailed | RIGLconnToPos | 0.563349 | 1 | 1 | 0.333939 |
| 6 | StimDetailed | RIGLconnToPos | 0.56322 | 1 | 1 | 0.33416 |
| 2 | bhv | RIGLconnToPos3 | 0.852213 | 1 | 3 | 0.788512 |
| 3 | bhv | RIGLconnToPos3 | 0.86671 | 1 | 3 | 0.727532 |
| 4 | bhv | RIGLconnToPos3 | 0.007016 | 3 | 3 | 12.1099 |
| 6 | bhv | RIGLconnToPos3 | 1.47E-08 | 3 | 3 | 39.34592 |
| 2 | FWD | RIGLconnToPos3 | 0.374551 | 1 | 1 | 0.788512 |
| 3 | FWD | RIGLconnToPos3 | 0.393684 | 1 | 1 | 0.727532 |
| 4 | FWD | RIGLconnToPos3 | 0.521763 | 2 | 1 | 0.410406 |
| 6 | FWD | RIGLconnToPos3 | 0.0001 | 2 | 1 | 15.132 |
| 2 | Stim | RIGLconnToPos3 | 0.316951 | 2 | 1 | 1.001489 |
| 3 | Stim | RIGLconnToPos3 | 0.31698 | 2 | 1 | 1.001367 |
| 4 | Stim | RIGLconnToPos3 | 0.317191 | 2 | 1 | 1.000496 |
| 6 | Stim | RIGLconnToPos3 | 4.54E-10 | 1 | 1 | 38.86488 |
| 2 | StimDetailed | RIGLconnToPos3 | 0.56322 | 1 | 1 | 0.33416 |
| 3 | StimDetailed | RIGLconnToPos3 | 0.56468 | 1 | 1 | 0.331666 |
| 4 | StimDetailed | RIGLconnToPos3 | 0.563349 | 1 | 1 | 0.333939 |
| 6 | StimDetailed | RIGLconnToPos3 | 0.048377 | 1 | 1 | 3.89685 |
| 2 | bhv | RIGLyhat | 0 | 1 | 3 | 264.9874 |
| 3 | bhv | RIGLyhat | 0 | 1 | 3 | 472.0281 |
| 4 | bhv | RIGLyhat | 0 | 3 | 3 | 352.085 |
| 6 | bhv | RIGLyhat | 0 | 1 | 3 | 404.9562 |
| 2 | FWD | RIGLyhat | 0 | 2 | 1 | 87.48912 |
| 3 | FWD | RIGLyhat | 0 | 1 | 1 | 363.8374 |
| 4 | FWD | RIGLyhat | 0.259605 | 2 | 1 | 1.270864 |
| 6 | FWD | RIGLyhat | 0 | 2 | 1 | 286.4555 |
| 2 | Stim | RIGLyhat | 0 | 1 | 1 | 175.521 |
| 3 | Stim | RIGLyhat | 0 | 1 | 1 | 86.12903 |
| 4 | Stim | RIGLyhat | 1.57E-05 | 1 | 1 | 18.65015 |
| 6 | Stim | RIGLyhat | 4.61E-05 | 1 | 1 | 16.60347 |
| 2 | StimD | RIGLyhat | 0.375292 | 1 | 1 | 0.786067 |
| 3 | StimD | RIGLyhat | 0.041838 | 1 | 1 | 4.141763 |
| 4 | StimD | RIGLyhat | 1.33E-10 | 1 | 1 | 41.26776 |
| 6 | StimD | RIGLyhat | 0.055553 | 1 | 1 | 3.665423 |
| 2 | bhv | RIGLyhat3 | NaN | 1 | 3 | NaN |
| 3 | bhv | RIGLyhat3 | NaN | 1 | 3 | NaN |
| 4 | bhv | RIGLyhat3 | 0.00021 | 3 | 3 | 19.5572 |
| 6 | bhv | RIGLyhat3 | 0 | 1 | 3 | 103.9001 |
| 2 | FWD | RIGLyhat3 | NaN | 1 | 1 | NaN |
| 3 | FWD | RIGLyhat3 | NaN | 1 | 1 | NaN |
| 4 | FWD | RIGLyhat3 | 0.023014 | 2 | 1 | 5.167429 |
| 6 | FWD | RIGLyhat3 | 0 | 1 | 1 | 103.9001 |
| 2 | Stim | RIGLyhat3 | NaN | 1 | 1 | NaN |
| 3 | Stim | RIGLyhat3 | NaN | 1 | 1 | NaN |
| 4 | Stim | RIGLyhat3 | 1.61E-13 | 2 | 1 | 54.42952 |
| 6 | Stim | RIGLyhat3 | 1.44E-09 | 2 | 1 | 36.61789 |
| 2 | StimD | RIGLyhat3 | NaN | 1 | 1 | NaN |
| 3 | StimD | RIGLyhat3 | NaN | 1 | 1 | NaN |
| 4 | StimD | RIGLyhat3 | 0 | 2 | 1 | 163.073 |
| 6 | StimD | RIGLyhat3 | 0 | 2 | 1 | 86.70126 |
| 1 | bhv | RISconnFrom3 | 0.000962 | 3 | 3 | 16.34804 |
| 2 | bhv | RISconnFrom3 | 0 | 1 | 3 | 1058.83 |
| 3 | bhv | RISconnFrom3 | 0 | 1 | 3 | 1124.39 |
| 4 | bhv | RISconnFrom3 | 0 | 1 | 3 | 369.8879 |
| 5 | bhv | RISconnFrom3 | 0 | 1 | 3 | 1074.343 |
| 6 | bhv | RISconnFrom3 | 0 | 1 | 3 | 1101.669 |
| 1 | FWD | RISconnFrom3 | 0.000982 | 2 | 1 | 10.86146 |
| 2 | FWD | RISconnFrom3 | 0 | 1 | 1 | 1040.901 |
| 3 | FWD | RISconnFrom3 | 0 | 1 | 1 | 1004.043 |
| 4 | FWD | RISconnFrom3 | 0 | 2 | 1 | 175.868 |
| 5 | FWD | RISconnFrom3 | 0 | 1 | 1 | 1044.536 |
| 6 | FWD | RISconnFrom3 | 0 | 1 | 1 | 785.4569 |
| 1 | Stim | RISconnFrom3 | 1.25E-09 | 1 | 1 | 36.8891 |
| 2 | Stim | RISconnFrom3 | 1.07E-14 | 2 | 1 | 59.77444 |
| 3 | Stim | RISconnFrom3 | 0 | 2 | 1 | 125.8139 |
| 4 | Stim | RISconnFrom3 | 0 | 2 | 1 | 297.7872 |
| 5 | Stim | RISconnFrom3 | 0 | 2 | 1 | 744.8344 |
| 6 | Stim | RISconnFrom3 | 1.28E-09 | 2 | 1 | 36.85035 |
| 1 | StimDetailed | RISconnFrom3 | 9.00E-07 | 1 | 1 | 24.13013 |
| 2 | StimDetailed | RISconnFrom3 | 0.0109 | 1 | 1 | 6.481579 |
| 3 | StimDetailed | RISconnFrom3 | 2.22E-16 | 1 | 1 | 67.36094 |
| 4 | StimDetailed | RISconnFrom3 | 0.014009 | 1 | 1 | 6.036948 |
| 5 | StimDetailed | RISconnFrom3 | 0 | 1 | 1 | 294.2571 |
| 6 | StimDetailed | RISconnFrom3 | 0.009288 | 1 | 1 | 6.766642 |
| 1 | bhv | RISconnFromNeg | 0 | 1 | 3 | 79.92578 |
| 2 | bhv | RISconnFromNeg | NaN | 1 | 3 | NaN |
| 3 | bhv | RISconnFromNeg | NaN | 1 | 3 | NaN |
| 4 | bhv | RISconnFromNeg | NaN | 1 | 3 | NaN |
| 5 | bhv | RISconnFromNeg | 0 | 4 | 3 | 113.7796 |
| 6 | bhv | RISconnFromNeg | NaN | 1 | 3 | NaN |
| 1 | FWD | RISconnFromNeg | 4.11E-15 | 1 | 1 | 61.62271 |
| 2 | FWD | RISconnFromNeg | NaN | 1 | 1 | NaN |
| 3 | FWD | RISconnFromNeg | NaN | 1 | 1 | NaN |
| 4 | FWD | RISconnFromNeg | NaN | 1 | 1 | NaN |
| 5 | FWD | RISconnFromNeg | 6.10E-05 | 2 | 1 | 16.07148 |
| 6 | FWD | RISconnFromNeg | NaN | 1 | 1 | NaN |
| 1 | Stim | RISconnFromNeg | 2.78E-10 | 2 | 1 | 39.82159 |
| 2 | Stim | RISconnFromNeg | NaN | 1 | 1 | NaN |
| 3 | Stim | RISconnFromNeg | NaN | 1 | 1 | NaN |
| 4 | Stim | RISconnFromNeg | NaN | 1 | 1 | NaN |
| 5 | Stim | RISconnFromNeg | 0.849784 | 1 | 1 | 0.03587 |
| 6 | Stim | RISconnFromNeg | NaN | 1 | 1 | NaN |
| 5 | StimDetailed | RISconnFromNeg | 0.003547 | 1 | 1 | 8.502054 |
| 1 | bhv | RISconnFromNeg3 | 7.55E-11 | 1 | 3 | 50.11433 |
| 2 | bhv | RISconnFromNeg3 | NaN | 1 | 3 | NaN |
| 3 | bhv | RISconnFromNeg3 | NaN | 1 | 3 | NaN |
| 4 | bhv | RISconnFromNeg3 | NaN | 1 | 3 | NaN |
| 5 | bhv | RISconnFromNeg3 | 1.09E-09 | 4 | 3 | 44.6705 |
| 6 | bhv | RISconnFromNeg3 | NaN | 1 | 3 | NaN |
| 1 | FWD | RISconnFromNeg3 | 4.74E-11 | 1 | 1 | 43.28066 |
| 2 | FWD | RISconnFromNeg3 | NaN | 1 | 1 | NaN |
| 3 | FWD | RISconnFromNeg3 | NaN | 1 | 1 | NaN |
| 4 | FWD | RISconnFromNeg3 | NaN | 1 | 1 | NaN |
| 5 | FWD | RISconnFromNeg3 | 0.031831 | 2 | 1 | 4.607581 |
| 6 | FWD | RISconnFromNeg3 | NaN | 1 | 1 | NaN |
| 1 | Stim | RISconnFromNeg3 | 6.60E-10 | 2 | 1 | 38.13633 |
| 2 | Stim | RISconnFromNeg3 | NaN | 1 | 1 | NaN |
| 3 | Stim | RISconnFromNeg3 | NaN | 1 | 1 | NaN |
| 4 | Stim | RISconnFromNeg3 | NaN | 1 | 1 | NaN |
| 5 | Stim | RISconnFromNeg3 | 0.008079 | 2 | 1 | 7.015955 |
| 6 | Stim | RISconnFromNeg3 | NaN | 1 | 1 | NaN |
| 5 | StimDetailed | RISconnFromNeg3 | 4.41E-06 | 2 | 1 | 21.07977 |
| 1 | bhv | RISconnFromPos | 0.153422 | 3 | 3 | 5.264503 |
| 2 | bhv | RISconnFromPos | 0 | 1 | 3 | 104.5495 |
| 3 | bhv | RISconnFromPos | 0.077833 | 1 | 3 | 6.820848 |
| 4 | bhv | RISconnFromPos | 0.007016 | 3 | 3 | 12.1099 |
| 5 | bhv | RISconnFromPos | 0 | 3 | 3 | 178.7878 |
| 6 | bhv | RISconnFromPos | 0.821141 | 3 | 3 | 0.917755 |
| 1 | FWD | RISconnFromPos | 0.455954 | 2 | 1 | 0.555808 |
| 2 | FWD | RISconnFromPos | 1.45E-14 | 1 | 1 | 59.15543 |
| 3 | FWD | RISconnFromPos | 0.388511 | 1 | 1 | 0.743596 |
| 4 | FWD | RISconnFromPos | 0.521763 | 2 | 1 | 0.410406 |
| 5 | FWD | RISconnFromPos | 8.18E-12 | 2 | 1 | 46.72267 |
| 6 | FWD | RISconnFromPos | 0.552443 | 2 | 1 | 0.352958 |
| 1 | Stim | RISconnFromPos | 0.3172 | 2 | 1 | 1.000455 |
| 2 | Stim | RISconnFromPos | 1.66E-08 | 2 | 1 | 31.85334 |
| 3 | Stim | RISconnFromPos | 0.082914 | 2 | 1 | 3.006838 |
| 4 | Stim | RISconnFromPos | 0.317191 | 2 | 1 | 1.000496 |
| 5 | Stim | RISconnFromPos | 0 | 2 | 1 | 267.0984 |
| 6 | Stim | RISconnFromPos | 0.316951 | 2 | 1 | 1.001489 |
| 1 | StimDetailed | RISconnFromPos | 0.563378 | 1 | 1 | 0.33389 |
| 2 | StimDetailed | RISconnFromPos | 0.015251 | 1 | 1 | 5.88721 |
| 3 | StimDetailed | RISconnFromPos | 0.318303 | 1 | 1 | 0.995905 |
| 4 | StimDetailed | RISconnFromPos | 0.563349 | 1 | 1 | 0.333939 |
| 5 | StimDetailed | RISconnFromPos | 0 | 1 | 1 | 88.89807 |
| 6 | StimDetailed | RISconnFromPos | 0.56322 | 1 | 1 | 0.33416 |
| 1 | bhv | RISconnFromPos3 | 0.010412 | 3 | 3 | 11.25754 |
| 2 | bhv | RISconnFromPos3 | 0 | 1 | 3 | 1058.83 |
| 3 | bhv | RISconnFromPos3 | 0 | 1 | 3 | 1124.39 |
| 4 | bhv | RISconnFromPos3 | 0 | 1 | 3 | 369.8879 |
| 5 | bhv | RISconnFromPos3 | 0 | 1 | 3 | 1074.343 |
| 6 | bhv | RISconnFromPos3 | 0 | 1 | 3 | 1101.669 |
| 1 | FWD | RISconnFromPos3 | 0.513743 | 2 | 1 | 0.426435 |
| 2 | FWD | RISconnFromPos3 | 0 | 1 | 1 | 1040.901 |
| 3 | FWD | RISconnFromPos3 | 0 | 1 | 1 | 1004.043 |
| 4 | FWD | RISconnFromPos3 | 0 | 2 | 1 | 175.868 |
| 5 | FWD | RISconnFromPos3 | 0 | 1 | 1 | 1044.536 |
| 6 | FWD | RISconnFromPos3 | 0 | 1 | 1 | 785.4569 |
| 1 | Stim | RISconnFromPos3 | 1.11E-16 | 1 | 1 | 69.93889 |
| 2 | Stim | RISconnFromPos3 | 1.07E-14 | 2 | 1 | 59.77444 |
| 3 | Stim | RISconnFromPos3 | 0 | 2 | 1 | 125.8139 |
| 4 | Stim | RISconnFromPos3 | 0 | 2 | 1 | 297.7872 |
| 5 | Stim | RISconnFromPos3 | 0 | 2 | 1 | 744.8344 |
| 6 | Stim | RISconnFromPos3 | 1.28E-09 | 2 | 1 | 36.85035 |
| 1 | StimDetailed | RISconnFromPos3 | 0.051391 | 1 | 1 | 3.795473 |
| 2 | StimDetailed | RISconnFromPos3 | 0.0109 | 1 | 1 | 6.481579 |
| 3 | StimDetailed | RISconnFromPos3 | 2.22E-16 | 1 | 1 | 67.36094 |
| 4 | StimDetailed | RISconnFromPos3 | 0.014009 | 1 | 1 | 6.036948 |
| 5 | StimDetailed | RISconnFromPos3 | 0 | 1 | 1 | 294.2571 |
| 6 | StimDetailed | RISconnFromPos3 | 0.009288 | 1 | 1 | 6.766642 |
| 1 | bhv | RISconnTo3 | 0.000469 | 3 | 3 | 17.86335 |
| 2 | bhv | RISconnTo3 | 0 | 1 | 3 | 883.0234 |
| 3 | bhv | RISconnTo3 | 0 | 1 | 3 | 152.6555 |
| 4 | bhv | RISconnTo3 | 7.78E-05 | 3 | 3 | 21.63331 |
| 5 | bhv | RISconnTo3 | 0 | 1 | 3 | 1081.612 |
| 6 | bhv | RISconnTo3 | 0 | 1 | 3 | 1257.173 |
| 1 | FWD | RISconnTo3 | 0.011174 | 2 | 1 | 6.437474 |
| 2 | FWD | RISconnTo3 | 0 | 1 | 1 | 785.6922 |
| 3 | FWD | RISconnTo3 | 0 | 1 | 1 | 133.6314 |
| 4 | FWD | RISconnTo3 | 0.000132 | 2 | 1 | 14.6105 |
| 5 | FWD | RISconnTo3 | 0 | 1 | 1 | 1049.527 |
| 6 | FWD | RISconnTo3 | 0 | 2 | 1 | 610.0056 |
| 1 | Stim | RISconnTo3 | 9.37E-11 | 1 | 1 | 41.94769 |
| 2 | Stim | RISconnTo3 | 0.477601 | 1 | 1 | 0.504336 |
| 3 | Stim | RISconnTo3 | 0 | 1 | 1 | 79.27358 |
| 4 | Stim | RISconnTo3 | 1.83E-08 | 2 | 1 | 31.66345 |
| 5 | Stim | RISconnTo3 | 0 | 2 | 1 | 731.444 |
| 6 | Stim | RISconnTo3 | 0.007911 | 2 | 1 | 7.053413 |
| 1 | StimDetailed | RISconnTo3 | 0.00036 | 1 | 1 | 12.72979 |
| 2 | StimDetailed | RISconnTo3 | 2.66E-15 | 1 | 1 | 62.47534 |
| 3 | StimDetailed | RISconnTo3 | 5.19E-09 | 1 | 1 | 34.11829 |
| 4 | StimDetailed | RISconnTo3 | 0.002256 | 1 | 1 | 9.328616 |
| 5 | StimDetailed | RISconnTo3 | 0 | 1 | 1 | 287.4945 |
| 6 | StimDetailed | RISconnTo3 | 0.038278 | 1 | 1 | 4.292617 |
| 1 | bhv | RISconnToNeg | 9.56E-12 | 1 | 3 | 54.32583 |
| 2 | bhv | RISconnToNeg | NaN | 1 | 3 | NaN |
| 3 | bhv | RISconnToNeg | NaN | 1 | 3 | NaN |
| 4 | bhv | RISconnToNeg | NaN | 1 | 3 | NaN |
| 5 | bhv | RISconnToNeg | 3.47E-09 | 4 | 3 | 42.29563 |
| 6 | bhv | RISconnToNeg | NaN | 1 | 3 | NaN |
| 1 | FWD | RISconnToNeg | 1.18E-08 | 1 | 1 | 32.51581 |
| 2 | FWD | RISconnToNeg | NaN | 1 | 1 | NaN |
| 3 | FWD | RISconnToNeg | NaN | 1 | 1 | NaN |
| 4 | FWD | RISconnToNeg | NaN | 1 | 1 | NaN |
| 5 | FWD | RISconnToNeg | 0.066323 | 2 | 1 | 3.371761 |
| 6 | FWD | RISconnToNeg | NaN | 1 | 1 | NaN |
| 1 | Stim | RISconnToNeg | 0.000208 | 2 | 1 | 13.75399 |
| 2 | Stim | RISconnToNeg | NaN | 1 | 1 | NaN |
| 3 | Stim | RISconnToNeg | NaN | 1 | 1 | NaN |
| 4 | Stim | RISconnToNeg | NaN | 1 | 1 | NaN |
| 5 | Stim | RISconnToNeg | 3.44E-08 | 1 | 1 | 30.44221 |
| 6 | Stim | RISconnToNeg | NaN | 1 | 1 | NaN |
| 5 | StimDetailed | RISconnToNeg | 0.001457 | 1 | 1 | 10.13204 |
| 5 | StimDetailed | RISconnToNeg | 0.001457 | 1 | 1 | 10.13204 |
| 1 | bhv | RISconnToNeg3 | 1.96E-07 | 4 | 3 | 34.02338 |
| 2 | bhv | RISconnToNeg3 | NaN | 1 | 3 | NaN |
| 3 | bhv | RISconnToNeg3 | NaN | 1 | 3 | NaN |
| 4 | bhv | RISconnToNeg3 | NaN | 1 | 3 | NaN |
| 5 | bhv | RISconnToNeg3 | NaN | 1 | 3 | NaN |
| 6 | bhv | RISconnToNeg3 | NaN | 1 | 3 | NaN |
| 1 | FWD | RISconnToNeg3 | 0.124954 | 2 | 1 | 2.354094 |
| 2 | FWD | RISconnToNeg3 | NaN | 1 | 1 | NaN |
| 3 | FWD | RISconnToNeg3 | NaN | 1 | 1 | NaN |
| 4 | FWD | RISconnToNeg3 | NaN | 1 | 1 | NaN |
| 5 | FWD | RISconnToNeg3 | NaN | 1 | 1 | NaN |
| 6 | FWD | RISconnToNeg3 | NaN | 1 | 1 | NaN |
| 1 | Stim | RISconnToNeg3 | 0.000299 | 1 | 1 | 13.07735 |
| 2 | Stim | RISconnToNeg3 | NaN | 1 | 1 | NaN |
| 3 | Stim | RISconnToNeg3 | NaN | 1 | 1 | NaN |
| 4 | Stim | RISconnToNeg3 | NaN | 1 | 1 | NaN |
| 5 | Stim | RISconnToNeg3 | NaN | 1 | 1 | NaN |
| 6 | Stim | RISconnToNeg3 | NaN | 1 | 1 | NaN |
| 5 | StimDetailed | RISconnToNeg3 | NaN | 1 | 1 | NaN |
| 5 | StimDetailed | RISconnToNeg3 | NaN | 1 | 1 | NaN |
| 1 | bhv | RISconnToPos | 0.153422 | 3 | 3 | 5.264503 |
| 2 | bhv | RISconnToPos | 0.852213 | 1 | 3 | 0.788512 |
| 3 | bhv | RISconnToPos | 0.86671 | 1 | 3 | 0.727532 |
| 4 | bhv | RISconnToPos | 0.007016 | 3 | 3 | 12.1099 |
| 5 | bhv | RISconnToPos | 0.004201 | 3 | 3 | 13.21129 |
| 6 | bhv | RISconnToPos | 0.821141 | 3 | 3 | 0.917755 |
| 1 | FWD | RISconnToPos | 0.455954 | 2 | 1 | 0.555808 |
| 2 | FWD | RISconnToPos | 0.374551 | 1 | 1 | 0.788512 |
| 3 | FWD | RISconnToPos | 0.393684 | 1 | 1 | 0.727532 |
| 4 | FWD | RISconnToPos | 0.521763 | 2 | 1 | 0.410406 |
| 5 | FWD | RISconnToPos | 0.104907 | 2 | 1 | 2.629308 |
| 6 | FWD | RISconnToPos | 0.552443 | 2 | 1 | 0.352958 |
| 1 | Stim | RISconnToPos | 0.3172 | 2 | 1 | 1.000455 |
| 2 | Stim | RISconnToPos | 0.316951 | 2 | 1 | 1.001489 |
| 3 | Stim | RISconnToPos | 0.31698 | 2 | 1 | 1.001367 |
| 4 | Stim | RISconnToPos | 0.317191 | 2 | 1 | 1.000496 |
| 5 | Stim | RISconnToPos | 0.045206 | 2 | 1 | 4.010926 |
| 6 | Stim | RISconnToPos | 0.316951 | 2 | 1 | 1.001489 |
| 1 | StimDetailed | RISconnToPos | 0.563378 | 1 | 1 | 0.33389 |
| 2 | StimDetailed | RISconnToPos | 0.56322 | 1 | 1 | 0.33416 |
| 3 | StimDetailed | RISconnToPos | 0.56468 | 1 | 1 | 0.331666 |
| 4 | StimDetailed | RISconnToPos | 0.563349 | 1 | 1 | 0.333939 |
| 5 | StimDetailed | RISconnToPos | 0.247926 | 1 | 1 | 1.334952 |
| 6 | StimDetailed | RISconnToPos | 0.56322 | 1 | 1 | 0.33416 |
| 1 | bhv | RISconnToPos3 | 3.46E-05 | 3 | 3 | 23.32004 |
| 2 | bhv | RISconnToPos3 | 0 | 1 | 3 | 883.0234 |
| 3 | bhv | RISconnToPos3 | 0 | 1 | 3 | 152.6555 |
| 4 | bhv | RISconnToPos3 | 7.78E-05 | 3 | 3 | 21.63331 |
| 5 | bhv | RISconnToPos3 | 0 | 1 | 3 | 1081.612 |
| 6 | bhv | RISconnToPos3 | 0 | 1 | 3 | 1257.173 |
| 1 | FWD | RISconnToPos3 | 0.007581 | 2 | 1 | 7.129898 |
| 2 | FWD | RISconnToPos3 | 0 | 1 | 1 | 785.6922 |
| 3 | FWD | RISconnToPos3 | 0 | 1 | 1 | 133.6314 |
| 4 | FWD | RISconnToPos3 | 0.000132 | 2 | 1 | 14.6105 |
| 5 | FWD | RISconnToPos3 | 0 | 1 | 1 | 1049.527 |
| 6 | FWD | RISconnToPos3 | 0 | 2 | 1 | 610.0056 |
| 1 | Stim | RISconnToPos3 | 2.80E-06 | 1 | 1 | 21.94659 |
| 2 | Stim | RISconnToPos3 | 0.477601 | 1 | 1 | 0.504336 |
| 3 | Stim | RISconnToPos3 | 0 | 1 | 1 | 79.27358 |
| 4 | Stim | RISconnToPos3 | 1.83E-08 | 2 | 1 | 31.66345 |
| 5 | Stim | RISconnToPos3 | 0 | 2 | 1 | 731.444 |
| 6 | Stim | RISconnToPos3 | 0.007911 | 2 | 1 | 7.053413 |
| 1 | StimDetailed | RISconnToPos3 | 6.88E-05 | 1 | 1 | 15.84228 |
| 2 | StimDetailed | RISconnToPos3 | 2.66E-15 | 1 | 1 | 62.47534 |
| 3 | StimDetailed | RISconnToPos3 | 5.19E-09 | 1 | 1 | 34.11829 |
| 4 | StimDetailed | RISconnToPos3 | 0.002256 | 1 | 1 | 9.328616 |
| 5 | StimDetailed | RISconnToPos3 | 0 | 1 | 1 | 287.4945 |
| 6 | StimDetailed | RISconnToPos3 | 0.038278 | 1 | 1 | 4.292617 |
| 1 | bhv | RISyhat | 0 | 1 | 3 | 418.5486 |
| 2 | bhv | RISyhat | 0 | 1 | 3 | 94.02 |
| 3 | bhv | RISyhat | 0 | 1 | 3 | 1374.75 |
| 4 | bhv | RISyhat | 0 | 1 | 3 | 440.4671 |
| 5 | bhv | RISyhat | 0 | 1 | 3 | 371.1937 |
| 6 | bhv | RISyhat | 0 | 1 | 3 | 377.1536 |
| 1 | FWD | RISyhat | 0 | 1 | 1 | 395.9414 |
| 2 | FWD | RISyhat | 7.33E-15 | 1 | 1 | 60.50586 |
| 3 | FWD | RISyhat | 0 | 1 | 1 | 1340.91 |
| 4 | FWD | RISyhat | 0 | 2 | 1 | 337.836 |
| 5 | FWD | RISyhat | 0 | 1 | 1 | 370.3772 |
| 6 | FWD | RISyhat | 0 | 2 | 1 | 374.7798 |
| 1 | Stim | RISyhat | 0.029582 | 2 | 1 | 4.733394 |
| 2 | Stim | RISyhat | 0 | 2 | 1 | 439.0087 |
| 3 | Stim | RISyhat | 0 | 2 | 1 | 280.4845 |
| 4 | Stim | RISyhat | 0 | 2 | 1 | 199.1853 |
| 5 | Stim | RISyhat | 0 | 2 | 1 | 1519.873 |
| 6 | Stim | RISyhat | 0 | 2 | 1 | 541.4103 |
| 1 | StimD | RISyhat | 4.15E-11 | 1 | 1 | 43.5429 |
| 2 | StimD | RISyhat | 5.43E-10 | 1 | 1 | 38.5154 |
| 3 | StimD | RISyhat | 0 | 1 | 1 | 106.0757 |
| 4 | StimD | RISyhat | 0.136128 | 1 | 1 | 2.221196 |
| 5 | StimD | RISyhat | 0 | 1 | 1 | 431.1425 |
| 6 | StimD | RISyhat | 0 | 1 | 1 | 355.3057 |
| 1 | bhv | RISyhat3 | 3.62E-07 | 1 | 3 | 32.7617 |
| 2 | bhv | RISyhat3 | 3.33E-16 | 1 | 3 | 75.10756 |
| 3 | bhv | RISyhat3 | NaN | 1 | 3 | NaN |
| 4 | bhv | RISyhat3 | 0 | 1 | 3 | 221.6459 |
| 5 | bhv | RISyhat3 | 0.054122 | 1 | 3 | 7.637764 |
| 6 | bhv | RISyhat3 | 0 | 1 | 3 | 127.5039 |
| 1 | FWD | RISyhat3 | 4.94E-06 | 1 | 1 | 20.85961 |
| 2 | FWD | RISyhat3 | 0 | 1 | 1 | 75.10756 |
| 3 | FWD | RISyhat3 | NaN | 1 | 1 | NaN |
| 4 | FWD | RISyhat3 | 0 | 1 | 1 | 221.6459 |
| 5 | FWD | RISyhat3 | 0.005716 | 1 | 1 | 7.637764 |
| 6 | FWD | RISyhat3 | 0 | 1 | 1 | 127.5039 |
| 1 | Stim | RISyhat3 | 0.523285 | 2 | 1 | 0.407414 |
| 2 | Stim | RISyhat3 | 0 | 2 | 1 | 95.20514 |
| 3 | Stim | RISyhat3 | NaN | 1 | 1 | NaN |
| 4 | Stim | RISyhat3 | 0 | 2 | 1 | 90.91973 |
| 5 | Stim | RISyhat3 | 0.025247 | 2 | 1 | 5.006833 |
| 6 | Stim | RISyhat3 | 2.04E-11 | 2 | 1 | 44.93667 |
| 1 | StimD | RISyhat3 | 9.98E-06 | 2 | 1 | 19.51441 |
| 2 | StimD | RISyhat3 | 6.88E-08 | 1 | 1 | 29.09683 |
| 3 | StimD | RISyhat3 | NaN | 1 | 1 | NaN |
| 4 | StimD | RISyhat3 | 0.001231 | 1 | 1 | 10.44286 |
| 5 | StimD | RISyhat3 | 0.000105 | 2 | 1 | 15.04326 |
| 6 | StimD | RISyhat3 | 0 | 2 | 1 | 134.6765 |
| 1 | bhv | RMEDconnFrom | 0.153422 | 3 | 3 | 5.264503 |
| 2 | bhv | RMEDconnFrom | 0 | 1 | 3 | 458.822 |
| 3 | bhv | RMEDconnFrom | 0.86671 | 1 | 3 | 0.727532 |
| 4 | bhv | RMEDconnFrom | 0.007016 | 3 | 3 | 12.1099 |
| 5 | bhv | RMEDconnFrom | 0.004201 | 3 | 3 | 13.21129 |
| 6 | bhv | RMEDconnFrom | 0 | 1 | 3 | 857.5053 |
| 7 | bhv | RMEDconnFrom | 0 | 1 | 3 | 227.3615 |
| 1 | FWD | RMEDconnFrom | 0.455954 | 2 | 1 | 0.555808 |
| 2 | FWD | RMEDconnFrom | 0 | 1 | 1 | 453.185 |
| 3 | FWD | RMEDconnFrom | 0.393684 | 1 | 1 | 0.727532 |
| 4 | FWD | RMEDconnFrom | 0.521763 | 2 | 1 | 0.410406 |
| 5 | FWD | RMEDconnFrom | 0.104907 | 2 | 1 | 2.629308 |
| 6 | FWD | RMEDconnFrom | 0 | 2 | 1 | 382.3572 |
| 7 | FWD | RMEDconnFrom | 0 | 2 | 1 | 180.8015 |
| 1 | Stim | RMEDconnFrom | 0.3172 | 2 | 1 | 1.000455 |
| 2 | Stim | RMEDconnFrom | 0 | 2 | 1 | 380.3011 |
| 3 | Stim | RMEDconnFrom | 0.31698 | 2 | 1 | 1.001367 |
| 4 | Stim | RMEDconnFrom | 0.317191 | 2 | 1 | 1.000496 |
| 5 | Stim | RMEDconnFrom | 0.045206 | 2 | 1 | 4.010926 |
| 6 | Stim | RMEDconnFrom | 0.021195 | 2 | 1 | 5.310667 |
| 7 | Stim | RMEDconnFrom | 3.74E-08 | 2 | 1 | 30.28146 |
| 1 | StimDetailed | RMEDconnFrom | 0.563378 | 1 | 1 | 0.33389 |
| 2 | StimDetailed | RMEDconnFrom | 0 | 1 | 1 | 120.7377 |
| 3 | StimDetailed | RMEDconnFrom | 0.56468 | 1 | 1 | 0.331666 |
| 4 | StimDetailed | RMEDconnFrom | 0.563349 | 1 | 1 | 0.333939 |
| 5 | StimDetailed | RMEDconnFrom | 0.247926 | 1 | 1 | 1.334952 |
| 6 | StimDetailed | RMEDconnFrom | 0.027036 | 1 | 1 | 4.88848 |
| 7 | StimDetailed | RMEDconnFrom | 0.003307 | 1 | 1 | 8.629741 |
| 1 | bhv | RMEDconnFrom3 | 0.153422 | 3 | 3 | 5.264503 |
| 2 | bhv | RMEDconnFrom3 | 0 | 1 | 3 | 1685.895 |
| 3 | bhv | RMEDconnFrom3 | 0.010611 | 1 | 3 | 11.21651 |
| 4 | bhv | RMEDconnFrom3 | 0 | 1 | 3 | 992.8515 |
| 5 | bhv | RMEDconnFrom3 | 0.004201 | 3 | 3 | 13.21129 |
| 6 | bhv | RMEDconnFrom3 | 0 | 1 | 3 | 993.031 |
| 7 | bhv | RMEDconnFrom3 | 0 | 1 | 3 | 1207.939 |
| 1 | FWD | RMEDconnFrom3 | 0.455954 | 2 | 1 | 0.555808 |
| 2 | FWD | RMEDconnFrom3 | 0 | 1 | 1 | 1647.339 |
| 3 | FWD | RMEDconnFrom3 | 0.820963 | 1 | 1 | 0.051215 |
| 4 | FWD | RMEDconnFrom3 | 0 | 2 | 1 | 496.936 |
| 5 | FWD | RMEDconnFrom3 | 0.104907 | 2 | 1 | 2.629308 |
| 6 | FWD | RMEDconnFrom3 | 0 | 1 | 1 | 617.4302 |
| 7 | FWD | RMEDconnFrom3 | 0 | 1 | 1 | 1066.678 |
| 1 | Stim | RMEDconnFrom3 | 0.3172 | 2 | 1 | 1.000455 |
| 2 | Stim | RMEDconnFrom3 | 0.877386 | 2 | 1 | 0.023804 |
| 3 | Stim | RMEDconnFrom3 | 0.999486 | 2 | 1 | 4.15E-07 |
| 4 | Stim | RMEDconnFrom3 | 0.024262 | 1 | 1 | 5.075775 |
| 5 | Stim | RMEDconnFrom3 | 0.045206 | 2 | 1 | 4.010926 |
| 6 | Stim | RMEDconnFrom3 | 6.85E-07 | 2 | 1 | 24.65737 |
| 7 | Stim | RMEDconnFrom3 | 2.02E-06 | 2 | 1 | 22.57714 |
| 1 | StimDetailed | RMEDconnFrom3 | 0.563378 | 1 | 1 | 0.33389 |
| 2 | StimDetailed | RMEDconnFrom3 | 4.80E-08 | 1 | 1 | 29.7958 |
| 3 | StimDetailed | RMEDconnFrom3 | 0.41528 | 1 | 1 | 0.663634 |
| 4 | StimDetailed | RMEDconnFrom3 | 0.001178 | 1 | 1 | 10.52442 |
| 5 | StimDetailed | RMEDconnFrom3 | 0.247926 | 1 | 1 | 1.334952 |
| 6 | StimDetailed | RMEDconnFrom3 | 0.003259 | 1 | 1 | 8.656482 |
| 7 | StimDetailed | RMEDconnFrom3 | 3.34E-07 | 1 | 1 | 26.04279 |
| 1 | bhv | RMEDconnFromNeg | 4.25E-13 | 1 | 3 | 60.65948 |
| 2 | bhv | RMEDconnFromNeg | NaN | 1 | 3 | NaN |
| 3 | bhv | RMEDconnFromNeg | NaN | 1 | 3 | NaN |
| 4 | bhv | RMEDconnFromNeg | NaN | 1 | 3 | NaN |
| 5 | bhv | RMEDconnFromNeg | 2.54E-05 | 3 | 3 | 23.96288 |
| 6 | bhv | RMEDconnFromNeg | NaN | 1 | 3 | NaN |
| 7 | bhv | RMEDconnFromNeg | NaN | 1 | 3 | NaN |
| 1 | FWD | RMEDconnFromNeg | 1.96E-10 | 1 | 1 | 40.50183 |
| 2 | FWD | RMEDconnFromNeg | NaN | 1 | 1 | NaN |
| 3 | FWD | RMEDconnFromNeg | NaN | 1 | 1 | NaN |
| 4 | FWD | RMEDconnFromNeg | NaN | 1 | 1 | NaN |
| 5 | FWD | RMEDconnFromNeg | 0.000356 | 2 | 1 | 12.7481 |
| 6 | FWD | RMEDconnFromNeg | NaN | 1 | 1 | NaN |
| 7 | FWD | RMEDconnFromNeg | NaN | 1 | 1 | NaN |
| 1 | Stim | RMEDconnFromNeg | 1.43E-05 | 2 | 1 | 18.83403 |
| 2 | Stim | RMEDconnFromNeg | NaN | 1 | 1 | NaN |
| 3 | Stim | RMEDconnFromNeg | NaN | 1 | 1 | NaN |
| 4 | Stim | RMEDconnFromNeg | NaN | 1 | 1 | NaN |
| 5 | Stim | RMEDconnFromNeg | 8.30E-07 | 1 | 1 | 24.28664 |
| 6 | Stim | RMEDconnFromNeg | NaN | 1 | 1 | NaN |
| 7 | Stim | RMEDconnFromNeg | NaN | 1 | 1 | NaN |
| 5 | StimDetailed | RMEDconnFromNeg | 0.004468 | 1 | 1 | 8.083296 |
| 7 | StimDetailed | RMEDconnFromNeg | NaN | 1 | 1 | NaN |
| 1 | bhv | RMEDconnFromNeg3 | 1.38E-10 | 1 | 3 | 48.89102 |
| 2 | bhv | RMEDconnFromNeg3 | NaN | 1 | 3 | NaN |
| 3 | bhv | RMEDconnFromNeg3 | NaN | 1 | 3 | NaN |
| 4 | bhv | RMEDconnFromNeg3 | NaN | 1 | 3 | NaN |
| 5 | bhv | RMEDconnFromNeg3 | NaN | 1 | 3 | NaN |
| 6 | bhv | RMEDconnFromNeg3 | NaN | 1 | 3 | NaN |
| 7 | bhv | RMEDconnFromNeg3 | NaN | 1 | 3 | NaN |
| 1 | FWD | RMEDconnFromNeg3 | 8.20E-09 | 1 | 1 | 33.22673 |
| 2 | FWD | RMEDconnFromNeg3 | NaN | 1 | 1 | NaN |
| 3 | FWD | RMEDconnFromNeg3 | NaN | 1 | 1 | NaN |
| 4 | FWD | RMEDconnFromNeg3 | NaN | 1 | 1 | NaN |
| 5 | FWD | RMEDconnFromNeg3 | NaN | 1 | 1 | NaN |
| 6 | FWD | RMEDconnFromNeg3 | NaN | 1 | 1 | NaN |
| 7 | FWD | RMEDconnFromNeg3 | NaN | 1 | 1 | NaN |
| 1 | Stim | RMEDconnFromNeg3 | 1.33E-05 | 2 | 1 | 18.96674 |
| 2 | Stim | RMEDconnFromNeg3 | NaN | 1 | 1 | NaN |
| 3 | Stim | RMEDconnFromNeg3 | NaN | 1 | 1 | NaN |
| 4 | Stim | RMEDconnFromNeg3 | NaN | 1 | 1 | NaN |
| 5 | Stim | RMEDconnFromNeg3 | NaN | 1 | 1 | NaN |
| 6 | Stim | RMEDconnFromNeg3 | NaN | 1 | 1 | NaN |
| 7 | Stim | RMEDconnFromNeg3 | NaN | 1 | 1 | NaN |
| 5 | StimDetailed | RMEDconnFromNeg3 | NaN | 1 | 1 | NaN |
| 7 | StimDetailed | RMEDconnFromNeg3 | NaN | 1 | 1 | NaN |
| 1 | bhv | RMEDconnFromPos | 0.153422 | 3 | 3 | 5.264503 |
| 2 | bhv | RMEDconnFromPos | 0 | 1 | 3 | 458.822 |
| 3 | bhv | RMEDconnFromPos | 0.86671 | 1 | 3 | 0.727532 |
| 4 | bhv | RMEDconnFromPos | 0.007016 | 3 | 3 | 12.1099 |
| 5 | bhv | RMEDconnFromPos | 0.004201 | 3 | 3 | 13.21129 |
| 6 | bhv | RMEDconnFromPos | 0 | 1 | 3 | 857.5053 |
| 7 | bhv | RMEDconnFromPos | 0 | 1 | 3 | 227.3615 |
| 1 | FWD | RMEDconnFromPos | 0.455954 | 2 | 1 | 0.555808 |
| 2 | FWD | RMEDconnFromPos | 0 | 1 | 1 | 453.185 |
| 3 | FWD | RMEDconnFromPos | 0.393684 | 1 | 1 | 0.727532 |
| 4 | FWD | RMEDconnFromPos | 0.521763 | 2 | 1 | 0.410406 |
| 5 | FWD | RMEDconnFromPos | 0.104907 | 2 | 1 | 2.629308 |
| 6 | FWD | RMEDconnFromPos | 0 | 2 | 1 | 382.3572 |
| 7 | FWD | RMEDconnFromPos | 0 | 2 | 1 | 180.8015 |
| 1 | Stim | RMEDconnFromPos | 0.3172 | 2 | 1 | 1.000455 |
| 2 | Stim | RMEDconnFromPos | 0 | 2 | 1 | 380.3011 |
| 3 | Stim | RMEDconnFromPos | 0.31698 | 2 | 1 | 1.001367 |
| 4 | Stim | RMEDconnFromPos | 0.317191 | 2 | 1 | 1.000496 |
| 5 | Stim | RMEDconnFromPos | 0.045206 | 2 | 1 | 4.010926 |
| 6 | Stim | RMEDconnFromPos | 0.021195 | 2 | 1 | 5.310667 |
| 7 | Stim | RMEDconnFromPos | 3.74E-08 | 2 | 1 | 30.28146 |
| 1 | StimDetailed | RMEDconnFromPos | 0.563378 | 1 | 1 | 0.33389 |
| 2 | StimDetailed | RMEDconnFromPos | 0 | 1 | 1 | 120.7377 |
| 3 | StimDetailed | RMEDconnFromPos | 0.56468 | 1 | 1 | 0.331666 |
| 4 | StimDetailed | RMEDconnFromPos | 0.563349 | 1 | 1 | 0.333939 |
| 5 | StimDetailed | RMEDconnFromPos | 0.247926 | 1 | 1 | 1.334952 |
| 6 | StimDetailed | RMEDconnFromPos | 0.027036 | 1 | 1 | 4.88848 |
| 7 | StimDetailed | RMEDconnFromPos | 0.003307 | 1 | 1 | 8.629741 |
| 1 | bhv | RMEDconnFromPos3 | 0.000191 | 3 | 3 | 19.75565 |
| 2 | bhv | RMEDconnFromPos3 | 0 | 1 | 3 | 1685.895 |
| 3 | bhv | RMEDconnFromPos3 | 0.010611 | 1 | 3 | 11.21651 |
| 4 | bhv | RMEDconnFromPos3 | 0 | 1 | 3 | 992.8515 |
| 5 | bhv | RMEDconnFromPos3 | 0.004201 | 3 | 3 | 13.21129 |
| 6 | bhv | RMEDconnFromPos3 | 0 | 1 | 3 | 993.031 |
| 7 | bhv | RMEDconnFromPos3 | 0 | 1 | 3 | 1207.939 |
| 1 | FWD | RMEDconnFromPos3 | 0.708993 | 2 | 1 | 0.139285 |
| 2 | FWD | RMEDconnFromPos3 | 0 | 1 | 1 | 1647.339 |
| 3 | FWD | RMEDconnFromPos3 | 0.820963 | 1 | 1 | 0.051215 |
| 4 | FWD | RMEDconnFromPos3 | 0 | 2 | 1 | 496.936 |
| 5 | FWD | RMEDconnFromPos3 | 0.104907 | 2 | 1 | 2.629308 |
| 6 | FWD | RMEDconnFromPos3 | 0 | 1 | 1 | 617.4302 |
| 7 | FWD | RMEDconnFromPos3 | 0 | 1 | 1 | 1066.678 |
| 1 | Stim | RMEDconnFromPos3 | 0.051509 | 2 | 1 | 3.791656 |
| 2 | Stim | RMEDconnFromPos3 | 0.877386 | 2 | 1 | 0.023804 |
| 3 | Stim | RMEDconnFromPos3 | 0.999486 | 2 | 1 | 4.15E-07 |
| 4 | Stim | RMEDconnFromPos3 | 0.024262 | 1 | 1 | 5.075775 |
| 5 | Stim | RMEDconnFromPos3 | 0.045206 | 2 | 1 | 4.010926 |
| 6 | Stim | RMEDconnFromPos3 | 6.85E-07 | 2 | 1 | 24.65737 |
| 7 | Stim | RMEDconnFromPos3 | 2.02E-06 | 2 | 1 | 22.57714 |
| 1 | StimDetailed | RMEDconnFromPos3 | 0.870948 | 1 | 1 | 0.026391 |
| 2 | StimDetailed | RMEDconnFromPos3 | 4.80E-08 | 1 | 1 | 29.7958 |
| 3 | StimDetailed | RMEDconnFromPos3 | 0.41528 | 1 | 1 | 0.663634 |
| 4 | StimDetailed | RMEDconnFromPos3 | 0.001178 | 1 | 1 | 10.52442 |
| 5 | StimDetailed | RMEDconnFromPos3 | 0.247926 | 1 | 1 | 1.334952 |
| 6 | StimDetailed | RMEDconnFromPos3 | 0.003259 | 1 | 1 | 8.656482 |
| 7 | StimDetailed | RMEDconnFromPos3 | 3.34E-07 | 1 | 1 | 26.04279 |
| 1 | bhv | RMEDconnTo | 0.153422 | 3 | 3 | 5.264503 |
| 2 | bhv | RMEDconnTo | 0.852213 | 1 | 3 | 0.788512 |
| 3 | bhv | RMEDconnTo | 0.86671 | 1 | 3 | 0.727532 |
| 4 | bhv | RMEDconnTo | 0.007016 | 3 | 3 | 12.1099 |
| 5 | bhv | RMEDconnTo | 0.004201 | 3 | 3 | 13.21129 |
| 6 | bhv | RMEDconnTo | 0 | 3 | 3 | 477.9279 |
| 7 | bhv | RMEDconnTo | 0 | 1 | 3 | 121.2423 |
| 1 | FWD | RMEDconnTo | 0.455954 | 2 | 1 | 0.555808 |
| 2 | FWD | RMEDconnTo | 0.374551 | 1 | 1 | 0.788512 |
| 3 | FWD | RMEDconnTo | 0.393684 | 1 | 1 | 0.727532 |
| 4 | FWD | RMEDconnTo | 0.521763 | 2 | 1 | 0.410406 |
| 5 | FWD | RMEDconnTo | 0.104907 | 2 | 1 | 2.629308 |
| 6 | FWD | RMEDconnTo | 0 | 2 | 1 | 222.2191 |
| 7 | FWD | RMEDconnTo | 0 | 2 | 1 | 88.92835 |
| 1 | Stim | RMEDconnTo | 0.3172 | 2 | 1 | 1.000455 |
| 2 | Stim | RMEDconnTo | 0.316951 | 2 | 1 | 1.001489 |
| 3 | Stim | RMEDconnTo | 0.31698 | 2 | 1 | 1.001367 |
| 4 | Stim | RMEDconnTo | 0.317191 | 2 | 1 | 1.000496 |
| 5 | Stim | RMEDconnTo | 0.045206 | 2 | 1 | 4.010926 |
| 6 | Stim | RMEDconnTo | 0.000856 | 1 | 1 | 11.11514 |
| 7 | Stim | RMEDconnTo | 0.034362 | 2 | 1 | 4.476616 |
| 1 | StimDetailed | RMEDconnTo | 0.563378 | 1 | 1 | 0.33389 |
| 2 | StimDetailed | RMEDconnTo | 0.56322 | 1 | 1 | 0.33416 |
| 3 | StimDetailed | RMEDconnTo | 0.56468 | 1 | 1 | 0.331666 |
| 4 | StimDetailed | RMEDconnTo | 0.563349 | 1 | 1 | 0.333939 |
| 5 | StimDetailed | RMEDconnTo | 0.247926 | 1 | 1 | 1.334952 |
| 6 | StimDetailed | RMEDconnTo | 0.287016 | 1 | 1 | 1.133564 |
| 7 | StimDetailed | RMEDconnTo | 0.433997 | 1 | 1 | 0.612102 |
| 1 | bhv | RMEDconnTo3 | 0.20907 | 3 | 3 | 4.536361 |
| 2 | bhv | RMEDconnTo3 | 0 | 1 | 3 | 1675.188 |
| 3 | bhv | RMEDconnTo3 | 0.86671 | 1 | 3 | 0.727532 |
| 4 | bhv | RMEDconnTo3 | 0 | 3 | 3 | 209.5953 |
| 5 | bhv | RMEDconnTo3 | 0.004201 | 3 | 3 | 13.21129 |
| 6 | bhv | RMEDconnTo3 | 0 | 1 | 3 | 1005.758 |
| 7 | bhv | RMEDconnTo3 | 0 | 1 | 3 | 701.7116 |
| 1 | FWD | RMEDconnTo3 | 0.583886 | 2 | 1 | 0.299994 |
| 2 | FWD | RMEDconnTo3 | 0 | 1 | 1 | 1626.351 |
| 3 | FWD | RMEDconnTo3 | 0.393684 | 1 | 1 | 0.727532 |
| 4 | FWD | RMEDconnTo3 | 0 | 2 | 1 | 83.54662 |
| 5 | FWD | RMEDconnTo3 | 0.104907 | 2 | 1 | 2.629308 |
| 6 | FWD | RMEDconnTo3 | 0 | 2 | 1 | 607.0503 |
| 7 | FWD | RMEDconnTo3 | 0 | 1 | 1 | 642.106 |
| 1 | Stim | RMEDconnTo3 | 0.094902 | 1 | 1 | 2.789199 |
| 2 | Stim | RMEDconnTo3 | 0.835944 | 1 | 1 | 0.042884 |
| 3 | Stim | RMEDconnTo3 | 0.31698 | 2 | 1 | 1.001367 |
| 4 | Stim | RMEDconnTo3 | 0.076017 | 1 | 1 | 3.148057 |
| 5 | Stim | RMEDconnTo3 | 0.045206 | 2 | 1 | 4.010926 |
| 6 | Stim | RMEDconnTo3 | 1.68E-06 | 2 | 1 | 22.92894 |
| 7 | Stim | RMEDconnTo3 | 1.89E-15 | 2 | 1 | 63.19318 |
| 1 | StimDetailed | RMEDconnTo3 | 0.082447 | 1 | 1 | 3.01599 |
| 2 | StimDetailed | RMEDconnTo3 | 1.01E-08 | 1 | 1 | 32.8181 |
| 3 | StimDetailed | RMEDconnTo3 | 0.56468 | 1 | 1 | 0.331666 |
| 4 | StimDetailed | RMEDconnTo3 | 4.44E-16 | 1 | 1 | 66.25375 |
| 5 | StimDetailed | RMEDconnTo3 | 0.247926 | 1 | 1 | 1.334952 |
| 6 | StimDetailed | RMEDconnTo3 | 0.002324 | 1 | 1 | 9.273905 |
| 7 | StimDetailed | RMEDconnTo3 | 4.05E-14 | 1 | 1 | 57.1444 |
| 1 | bhv | RMEDconnToNeg | 0.000777 | 1 | 3 | 16.80092 |
| 2 | bhv | RMEDconnToNeg | NaN | 1 | 3 | NaN |
| 3 | bhv | RMEDconnToNeg | NaN | 1 | 3 | NaN |
| 4 | bhv | RMEDconnToNeg | NaN | 1 | 3 | NaN |
| 5 | bhv | RMEDconnToNeg | 2.34E-12 | 4 | 3 | 57.19401 |
| 6 | bhv | RMEDconnToNeg | NaN | 1 | 3 | NaN |
| 7 | bhv | RMEDconnToNeg | 1.81E-11 | 1 | 3 | 53.02995 |
| 1 | FWD | RMEDconnToNeg | 0.002185 | 2 | 1 | 9.387521 |
| 2 | FWD | RMEDconnToNeg | NaN | 1 | 1 | NaN |
| 3 | FWD | RMEDconnToNeg | NaN | 1 | 1 | NaN |
| 4 | FWD | RMEDconnToNeg | NaN | 1 | 1 | NaN |
| 5 | FWD | RMEDconnToNeg | 2.52E-05 | 2 | 1 | 17.75105 |
| 6 | FWD | RMEDconnToNeg | NaN | 1 | 1 | NaN |
| 7 | FWD | RMEDconnToNeg | 2.70E-12 | 1 | 1 | 48.89589 |
| 1 | Stim | RMEDconnToNeg | 0.581952 | 2 | 1 | 0.303089 |
| 2 | Stim | RMEDconnToNeg | NaN | 1 | 1 | NaN |
| 3 | Stim | RMEDconnToNeg | NaN | 1 | 1 | NaN |
| 4 | Stim | RMEDconnToNeg | NaN | 1 | 1 | NaN |
| 5 | Stim | RMEDconnToNeg | 0.000216 | 1 | 1 | 13.68833 |
| 6 | Stim | RMEDconnToNeg | NaN | 1 | 1 | NaN |
| 7 | Stim | RMEDconnToNeg | 1.14E-11 | 2 | 1 | 46.07145 |
| 5 | StimDetailed | RMEDconnToNeg | 0.438514 | 1 | 1 | 0.600167 |
| 5 | StimDetailed | RMEDconnToNeg | 0.438514 | 1 | 1 | 0.600167 |
| 7 | StimDetailed | RMEDconnToNeg | 8.92E-05 | 1 | 1 | 15.35208 |
| 7 | StimDetailed | RMEDconnToNeg | 8.92E-05 | 1 | 1 | 15.35208 |
| 1 | bhv | RMEDconnToNeg3 | 1.96E-07 | 4 | 3 | 34.02338 |
| 2 | bhv | RMEDconnToNeg3 | NaN | 1 | 3 | NaN |
| 3 | bhv | RMEDconnToNeg3 | NaN | 1 | 3 | NaN |
| 4 | bhv | RMEDconnToNeg3 | NaN | 1 | 3 | NaN |
| 5 | bhv | RMEDconnToNeg3 | NaN | 1 | 3 | NaN |
| 6 | bhv | RMEDconnToNeg3 | NaN | 1 | 3 | NaN |
| 7 | bhv | RMEDconnToNeg3 | NaN | 1 | 3 | NaN |
| 1 | FWD | RMEDconnToNeg3 | 0.124954 | 2 | 1 | 2.354094 |
| 2 | FWD | RMEDconnToNeg3 | NaN | 1 | 1 | NaN |
| 3 | FWD | RMEDconnToNeg3 | NaN | 1 | 1 | NaN |
| 4 | FWD | RMEDconnToNeg3 | NaN | 1 | 1 | NaN |
| 5 | FWD | RMEDconnToNeg3 | NaN | 1 | 1 | NaN |
| 6 | FWD | RMEDconnToNeg3 | NaN | 1 | 1 | NaN |
| 7 | FWD | RMEDconnToNeg3 | NaN | 1 | 1 | NaN |
| 1 | Stim | RMEDconnToNeg3 | 0.000299 | 1 | 1 | 13.07735 |
| 2 | Stim | RMEDconnToNeg3 | NaN | 1 | 1 | NaN |
| 3 | Stim | RMEDconnToNeg3 | NaN | 1 | 1 | NaN |
| 4 | Stim | RMEDconnToNeg3 | NaN | 1 | 1 | NaN |
| 5 | Stim | RMEDconnToNeg3 | NaN | 1 | 1 | NaN |
| 6 | Stim | RMEDconnToNeg3 | NaN | 1 | 1 | NaN |
| 7 | Stim | RMEDconnToNeg3 | NaN | 1 | 1 | NaN |
| 5 | StimDetailed | RMEDconnToNeg3 | NaN | 1 | 1 | NaN |
| 5 | StimDetailed | RMEDconnToNeg3 | NaN | 1 | 1 | NaN |
| 7 | StimDetailed | RMEDconnToNeg3 | NaN | 1 | 1 | NaN |
| 7 | StimDetailed | RMEDconnToNeg3 | NaN | 1 | 1 | NaN |
| 1 | bhv | RMEDconnToPos | 0.153422 | 3 | 3 | 5.264503 |
| 2 | bhv | RMEDconnToPos | 0.852213 | 1 | 3 | 0.788512 |
| 3 | bhv | RMEDconnToPos | 0.86671 | 1 | 3 | 0.727532 |
| 4 | bhv | RMEDconnToPos | 0.007016 | 3 | 3 | 12.1099 |
| 5 | bhv | RMEDconnToPos | 0.004201 | 3 | 3 | 13.21129 |
| 6 | bhv | RMEDconnToPos | 0 | 3 | 3 | 477.9279 |
| 7 | bhv | RMEDconnToPos | 0 | 1 | 3 | 121.2423 |
| 1 | FWD | RMEDconnToPos | 0.455954 | 2 | 1 | 0.555808 |
| 2 | FWD | RMEDconnToPos | 0.374551 | 1 | 1 | 0.788512 |
| 3 | FWD | RMEDconnToPos | 0.393684 | 1 | 1 | 0.727532 |
| 4 | FWD | RMEDconnToPos | 0.521763 | 2 | 1 | 0.410406 |
| 5 | FWD | RMEDconnToPos | 0.104907 | 2 | 1 | 2.629308 |
| 6 | FWD | RMEDconnToPos | 0 | 2 | 1 | 222.2191 |
| 7 | FWD | RMEDconnToPos | 0 | 2 | 1 | 88.92835 |
| 1 | Stim | RMEDconnToPos | 0.3172 | 2 | 1 | 1.000455 |
| 2 | Stim | RMEDconnToPos | 0.316951 | 2 | 1 | 1.001489 |
| 3 | Stim | RMEDconnToPos | 0.31698 | 2 | 1 | 1.001367 |
| 4 | Stim | RMEDconnToPos | 0.317191 | 2 | 1 | 1.000496 |
| 5 | Stim | RMEDconnToPos | 0.045206 | 2 | 1 | 4.010926 |
| 6 | Stim | RMEDconnToPos | 0.000856 | 1 | 1 | 11.11514 |
| 7 | Stim | RMEDconnToPos | 0.034362 | 2 | 1 | 4.476616 |
| 1 | StimDetailed | RMEDconnToPos | 0.563378 | 1 | 1 | 0.33389 |
| 2 | StimDetailed | RMEDconnToPos | 0.56322 | 1 | 1 | 0.33416 |
| 3 | StimDetailed | RMEDconnToPos | 0.56468 | 1 | 1 | 0.331666 |
| 4 | StimDetailed | RMEDconnToPos | 0.563349 | 1 | 1 | 0.333939 |
| 5 | StimDetailed | RMEDconnToPos | 0.247926 | 1 | 1 | 1.334952 |
| 6 | StimDetailed | RMEDconnToPos | 0.287016 | 1 | 1 | 1.133564 |
| 7 | StimDetailed | RMEDconnToPos | 0.433997 | 1 | 1 | 0.612102 |
| 1 | bhv | RMEDconnToPos3 | 0.000244 | 3 | 3 | 19.23521 |
| 2 | bhv | RMEDconnToPos3 | 0 | 1 | 3 | 1675.188 |
| 3 | bhv | RMEDconnToPos3 | 0.86671 | 1 | 3 | 0.727532 |
| 4 | bhv | RMEDconnToPos3 | 0 | 3 | 3 | 209.5953 |
| 5 | bhv | RMEDconnToPos3 | 0.004201 | 3 | 3 | 13.21129 |
| 6 | bhv | RMEDconnToPos3 | 0 | 1 | 3 | 1005.758 |
| 7 | bhv | RMEDconnToPos3 | 0 | 1 | 3 | 701.7116 |
| 1 | FWD | RMEDconnToPos3 | 0.832351 | 2 | 1 | 0.044811 |
| 2 | FWD | RMEDconnToPos3 | 0 | 1 | 1 | 1626.351 |
| 3 | FWD | RMEDconnToPos3 | 0.393684 | 1 | 1 | 0.727532 |
| 4 | FWD | RMEDconnToPos3 | 0 | 2 | 1 | 83.54662 |
| 5 | FWD | RMEDconnToPos3 | 0.104907 | 2 | 1 | 2.629308 |
| 6 | FWD | RMEDconnToPos3 | 0 | 2 | 1 | 607.0503 |
| 7 | FWD | RMEDconnToPos3 | 0 | 1 | 1 | 642.106 |
| 1 | Stim | RMEDconnToPos3 | 0.343795 | 2 | 1 | 0.896228 |
| 2 | Stim | RMEDconnToPos3 | 0.835944 | 1 | 1 | 0.042884 |
| 3 | Stim | RMEDconnToPos3 | 0.31698 | 2 | 1 | 1.001367 |
| 4 | Stim | RMEDconnToPos3 | 0.076017 | 1 | 1 | 3.148057 |
| 5 | Stim | RMEDconnToPos3 | 0.045206 | 2 | 1 | 4.010926 |
| 6 | Stim | RMEDconnToPos3 | 1.68E-06 | 2 | 1 | 22.92894 |
| 7 | Stim | RMEDconnToPos3 | 1.89E-15 | 2 | 1 | 63.19318 |
| 1 | StimDetailed | RMEDconnToPos3 | 0.013852 | 1 | 1 | 6.056883 |
| 2 | StimDetailed | RMEDconnToPos3 | 1.01E-08 | 1 | 1 | 32.8181 |
| 3 | StimDetailed | RMEDconnToPos3 | 0.56468 | 1 | 1 | 0.331666 |
| 4 | StimDetailed | RMEDconnToPos3 | 4.44E-16 | 1 | 1 | 66.25375 |
| 5 | StimDetailed | RMEDconnToPos3 | 0.247926 | 1 | 1 | 1.334952 |
| 6 | StimDetailed | RMEDconnToPos3 | 0.002324 | 1 | 1 | 9.273905 |
| 7 | StimDetailed | RMEDconnToPos3 | 4.05E-14 | 1 | 1 | 57.1444 |
| 1 | bhv | RMEDyhat | 0 | 1 | 3 | 1039.321 |
| 2 | bhv | RMEDyhat | 0 | 1 | 3 | 1448.827 |
| 3 | bhv | RMEDyhat | 0 | 1 | 3 | 718.4859 |
| 4 | bhv | RMEDyhat | 0 | 1 | 3 | 1189.659 |
| 5 | bhv | RMEDyhat | 0 | 1 | 3 | 1542.741 |
| 6 | bhv | RMEDyhat | 0 | 1 | 3 | 1195.449 |
| 7 | bhv | RMEDyhat | 0 | 1 | 3 | 1746.502 |
| 1 | FWD | RMEDyhat | 0 | 1 | 1 | 962.6788 |
| 2 | FWD | RMEDyhat | 0 | 1 | 1 | 1444.488 |
| 3 | FWD | RMEDyhat | 0 | 1 | 1 | 563.653 |
| 4 | FWD | RMEDyhat | 0 | 1 | 1 | 805.3795 |
| 5 | FWD | RMEDyhat | 0 | 1 | 1 | 1358.04 |
| 6 | FWD | RMEDyhat | 0 | 1 | 1 | 678.7785 |
| 7 | FWD | RMEDyhat | 0 | 1 | 1 | 1484.568 |
| 1 | Stim | RMEDyhat | 0 | 2 | 1 | 135.6306 |
| 2 | Stim | RMEDyhat | 0.462476 | 2 | 1 | 0.539895 |
| 3 | Stim | RMEDyhat | 3.24E-11 | 1 | 1 | 44.02427 |
| 4 | Stim | RMEDyhat | 5.96E-11 | 1 | 1 | 42.83251 |
| 5 | Stim | RMEDyhat | 0 | 2 | 1 | 132.9762 |
| 6 | Stim | RMEDyhat | 0 | 2 | 1 | 82.93539 |
| 7 | Stim | RMEDyhat | 0.000562 | 1 | 1 | 11.89938 |
| 1 | StimD | RMEDyhat | 1.57E-14 | 1 | 1 | 59.01426 |
| 2 | StimD | RMEDyhat | 8.84E-10 | 1 | 1 | 37.56491 |
| 3 | StimD | RMEDyhat | 0.720607 | 1 | 1 | 0.127912 |
| 4 | StimD | RMEDyhat | 0 | 1 | 1 | 117.586 |
| 5 | StimD | RMEDyhat | 1.67E-13 | 1 | 1 | 54.35565 |
| 6 | StimD | RMEDyhat | 1.97E-09 | 1 | 1 | 36.00591 |
| 7 | StimD | RMEDyhat | 0.002329 | 1 | 1 | 9.270383 |
| 1 | bhv | RMEDyhat3 | 6.11E-08 | 4 | 3 | 36.41908 |
| 2 | bhv | RMEDyhat3 | NaN | 1 | 3 | NaN |
| 3 | bhv | RMEDyhat3 | 0.692525 | 1 | 3 | 1.455728 |
| 4 | bhv | RMEDyhat3 | 0 | 1 | 3 | 96.94859 |
| 5 | bhv | RMEDyhat3 | 0 | 4 | 3 | 120.4831 |
| 6 | bhv | RMEDyhat3 | NaN | 1 | 3 | NaN |
| 7 | bhv | RMEDyhat3 | NaN | 1 | 3 | NaN |
| 1 | FWD | RMEDyhat3 | 0.093215 | 2 | 1 | 2.817949 |
| 2 | FWD | RMEDyhat3 | NaN | 1 | 1 | NaN |
| 3 | FWD | RMEDyhat3 | 0.227611 | 1 | 1 | 1.455728 |
| 4 | FWD | RMEDyhat3 | 0 | 1 | 1 | 96.94859 |
| 5 | FWD | RMEDyhat3 | 0.001033 | 2 | 1 | 10.76832 |
| 6 | FWD | RMEDyhat3 | NaN | 1 | 1 | NaN |
| 7 | FWD | RMEDyhat3 | NaN | 1 | 1 | NaN |
| 1 | Stim | RMEDyhat3 | 0.000174 | 1 | 1 | 14.08974 |
| 2 | Stim | RMEDyhat3 | NaN | 1 | 1 | NaN |
| 3 | Stim | RMEDyhat3 | 0.999486 | 1 | 1 | 4.15E-07 |
| 4 | Stim | RMEDyhat3 | 2.86E-10 | 1 | 1 | 39.76857 |
| 5 | Stim | RMEDyhat3 | 4.17E-06 | 2 | 1 | 21.18304 |
| 6 | Stim | RMEDyhat3 | NaN | 1 | 1 | NaN |
| 7 | Stim | RMEDyhat3 | NaN | 1 | 1 | NaN |
| 1 | StimD | RMEDyhat3 | 0.030123 | 1 | 1 | 4.702281 |
| 2 | StimD | RMEDyhat3 | NaN | 1 | 1 | NaN |
| 3 | StimD | RMEDyhat3 | 0.411453 | 1 | 1 | 0.674598 |
| 4 | StimD | RMEDyhat3 | 0.000269 | 1 | 1 | 13.27371 |
| 5 | StimD | RMEDyhat3 | 1.44E-15 | 2 | 1 | 63.64543 |
| 6 | StimD | RMEDyhat3 | NaN | 1 | 1 | NaN |
| 7 | StimD | RMEDyhat3 | NaN | 1 | 1 | NaN |
| 1 | bhv | SABDconnFrom3 | 0 | 3 | 3 | 1273.557 |
| 1 | FWD | SABDconnFrom3 | 0 | 2 | 1 | 1245.587 |
| 1 | Stim | SABDconnFrom3 | 7.93E-14 | 1 | 1 | 55.82267 |
| 1 | StimDetailed | SABDconnFrom3 | 1.10E-10 | 1 | 1 | 41.63502 |
| 1 | bhv | SABDconnFromNeg | 0 | 1 | 3 | 128.9231 |
| 1 | FWD | SABDconnFromNeg | 0 | 1 | 1 | 125.4333 |
| 1 | Stim | SABDconnFromNeg | 0 | 2 | 1 | 93.8425 |
| 1 | bhv | SABDconnFromNeg3 | 0 | 1 | 3 | 85.67865 |
| 1 | FWD | SABDconnFromNeg3 | 0 | 1 | 1 | 82.86649 |
| 1 | Stim | SABDconnFromNeg3 | 1.33E-15 | 2 | 1 | 63.79963 |
| 1 | bhv | SABDconnFromPos | 4.11E-12 | 3 | 3 | 56.04617 |
| 1 | FWD | SABDconnFromPos | 9.09E-13 | 2 | 1 | 51.03212 |
| 1 | Stim | SABDconnFromPos | 3.24E-11 | 1 | 1 | 44.02653 |
| 1 | StimDetailed | SABDconnFromPos | 0 | 1 | 1 | 135.8548 |
| 1 | bhv | SABDconnFromPos3 | 0 | 3 | 3 | 1449.883 |
| 1 | FWD | SABDconnFromPos3 | 0 | 2 | 1 | 1416.868 |
| 1 | Stim | SABDconnFromPos3 | 0 | 1 | 1 | 102.2355 |
| 1 | StimDetailed | SABDconnFromPos3 | 0 | 1 | 1 | 125.0272 |
| 1 | bhv | SABDconnTo3 | 0 | 3 | 3 | 1296.674 |
| 1 | FWD | SABDconnTo3 | 0 | 2 | 1 | 1278.101 |
| 1 | Stim | SABDconnTo3 | 0 | 1 | 1 | 83.59023 |
| 1 | StimDetailed | SABDconnTo3 | 2.55E-15 | 1 | 1 | 62.61824 |
| 1 | bhv | SABDconnToNeg | 0 | 1 | 3 | 110.8257 |
| 1 | FWD | SABDconnToNeg | 0 | 1 | 1 | 107.7602 |
| 1 | Stim | SABDconnToNeg | 0 | 2 | 1 | 80.86981 |
| 1 | bhv | SABDconnToNeg3 | 8.68E-13 | 1 | 3 | 59.20722 |
| 1 | FWD | SABDconnToNeg3 | 9.24E-14 | 1 | 1 | 55.52343 |
| 1 | Stim | SABDconnToNeg3 | 4.20E-12 | 2 | 1 | 48.02696 |
| 1 | bhv | SABDconnToPos | 6.53E-14 | 3 | 3 | 64.46438 |
| 1 | FWD | SABDconnToPos | 9.88E-15 | 2 | 1 | 59.9219 |
| 1 | Stim | SABDconnToPos | 2.26E-12 | 1 | 1 | 49.24188 |
| 1 | StimDetailed | SABDconnToPos | 0 | 1 | 1 | 151.505 |
| 1 | bhv | SABDconnToPos3 | 0 | 3 | 3 | 1419.49 |
| 1 | FWD | SABDconnToPos3 | 0 | 2 | 1 | 1396.352 |
| 1 | Stim | SABDconnToPos3 | 0 | 1 | 1 | 123.7109 |
| 1 | StimDetailed | SABDconnToPos3 | 0 | 1 | 1 | 132.1645 |
| 1 | bhv | SABDyhat | 0 | 3 | 3 | 1568.195 |
| 1 | FWD | SABDyhat | 0 | 2 | 1 | 1088.059 |
| 1 | Stim | SABDyhat | 0 | 1 | 1 | 81.02093 |
| 1 | StimD | SABDyhat | 0 | 1 | 1 | 108.9882 |
| 1 | bhv | SABDyhat3 | 2.79E-06 | 4 | 3 | 28.54217 |
| 1 | FWD | SABDyhat3 | 0.16733 | 2 | 1 | 1.906706 |
| 1 | Stim | SABDyhat3 | 0.000514 | 1 | 1 | 12.06587 |
| 1 | StimD | SABDyhat3 | 0.044782 | 1 | 1 | 4.026839 |
| 1 | bhv | stimTimes | 0 | 1 | 3 | 124.5294 |
| 2 | bhv | stimTimes | 0.005427 | 1 | 3 | 12.66214 |
| 3 | bhv | stimTimes | 0 | 1 | 3 | 93.35077 |
| 4 | bhv | stimTimes | 0.002228 | 3 | 3 | 14.56619 |
| 5 | bhv | stimTimes | 0 | 1 | 3 | 247.7468 |
| 6 | bhv | stimTimes | 1.55E-13 | 3 | 3 | 62.71312 |
| 7 | bhv | stimTimes | 6.01E-10 | 3 | 3 | 45.88311 |
| 1 | FWD | stimTimes | 0 | 2 | 1 | 117.0115 |
| 2 | FWD | stimTimes | 0.144226 | 1 | 1 | 2.132278 |
| 3 | FWD | stimTimes | 0 | 1 | 1 | 91.54593 |
| 4 | FWD | stimTimes | 0.00072 | 2 | 1 | 11.43778 |
| 5 | FWD | stimTimes | 0 | 1 | 1 | 229.4815 |
| 6 | FWD | stimTimes | 6.57E-13 | 2 | 1 | 51.66812 |
| 7 | FWD | stimTimes | 1.63E-11 | 2 | 1 | 45.37399 |
| 1 | bhv | stimTimesDetailed | 0 | 1 | 3 | 117.2001 |
| 2 | bhv | stimTimesDetailed | 5.05E-08 | 1 | 3 | 36.80785 |
| 3 | bhv | stimTimesDetailed | 7.11E-14 | 1 | 3 | 64.29177 |
| 4 | bhv | stimTimesDetailed | 2.38E-14 | 3 | 3 | 66.51937 |
| 5 | bhv | stimTimesDetailed | 3.97E-07 | 1 | 3 | 32.56672 |
| 6 | bhv | stimTimesDetailed | 3.49E-05 | 3 | 3 | 23.30617 |
| 7 | bhv | stimTimesDetailed | 3.38E-09 | 3 | 3 | 42.34827 |
| 1 | FWD | stimTimesDetailed | 1.11E-16 | 1 | 1 | 68.56674 |
| 2 | FWD | stimTimesDetailed | 4.49E-05 | 2 | 1 | 16.65269 |
| 3 | FWD | stimTimesDetailed | 9.01E-10 | 1 | 1 | 37.52745 |
| 4 | FWD | stimTimesDetailed | 1.55E-14 | 2 | 1 | 59.02509 |
| 5 | FWD | stimTimesDetailed | 5.63E-06 | 2 | 1 | 20.60837 |
| 6 | FWD | stimTimesDetailed | 2.14E-05 | 2 | 1 | 18.06051 |
| 7 | FWD | stimTimesDetailed | 1.89E-07 | 2 | 1 | 27.14673 |

**Table 6:**
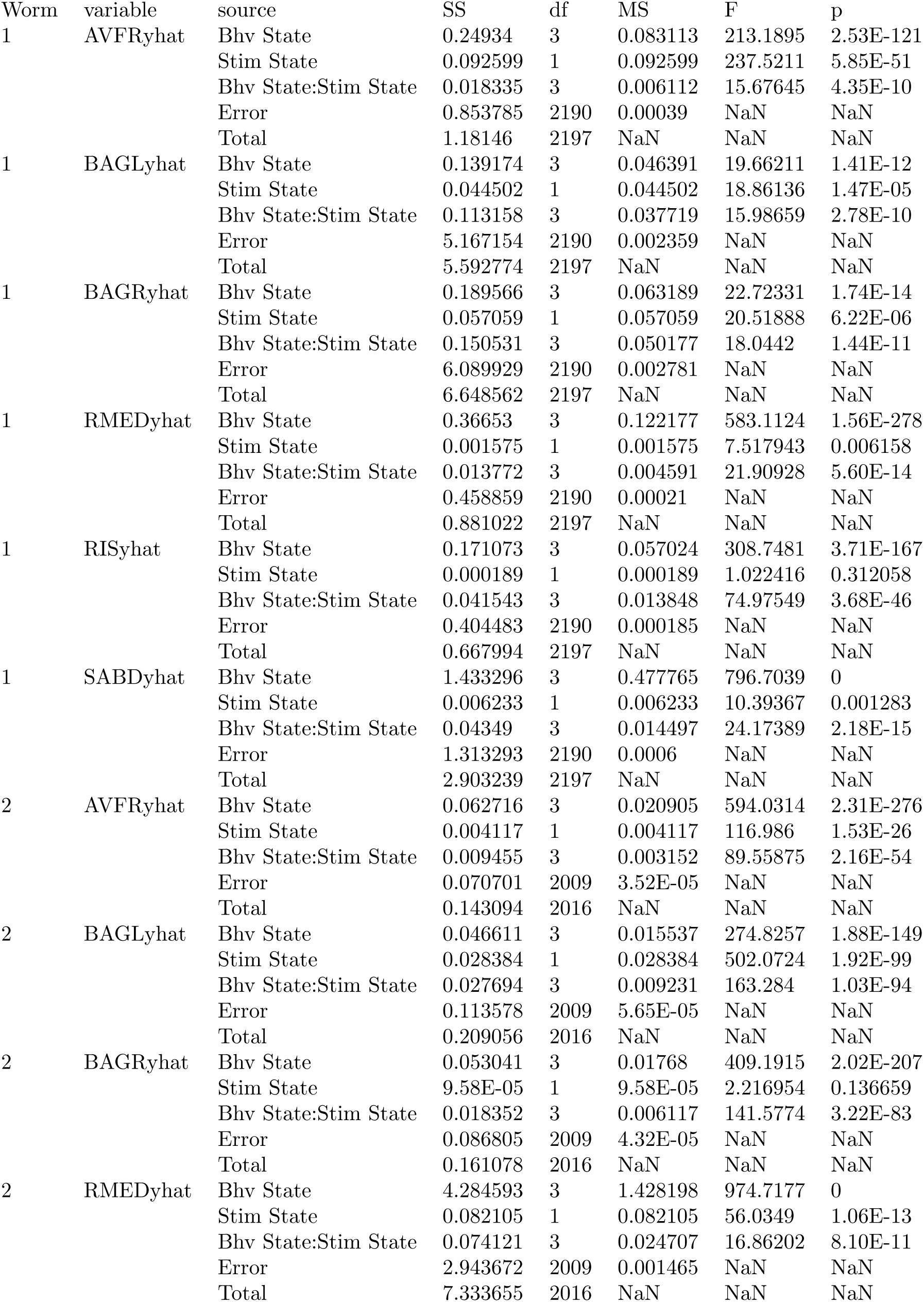
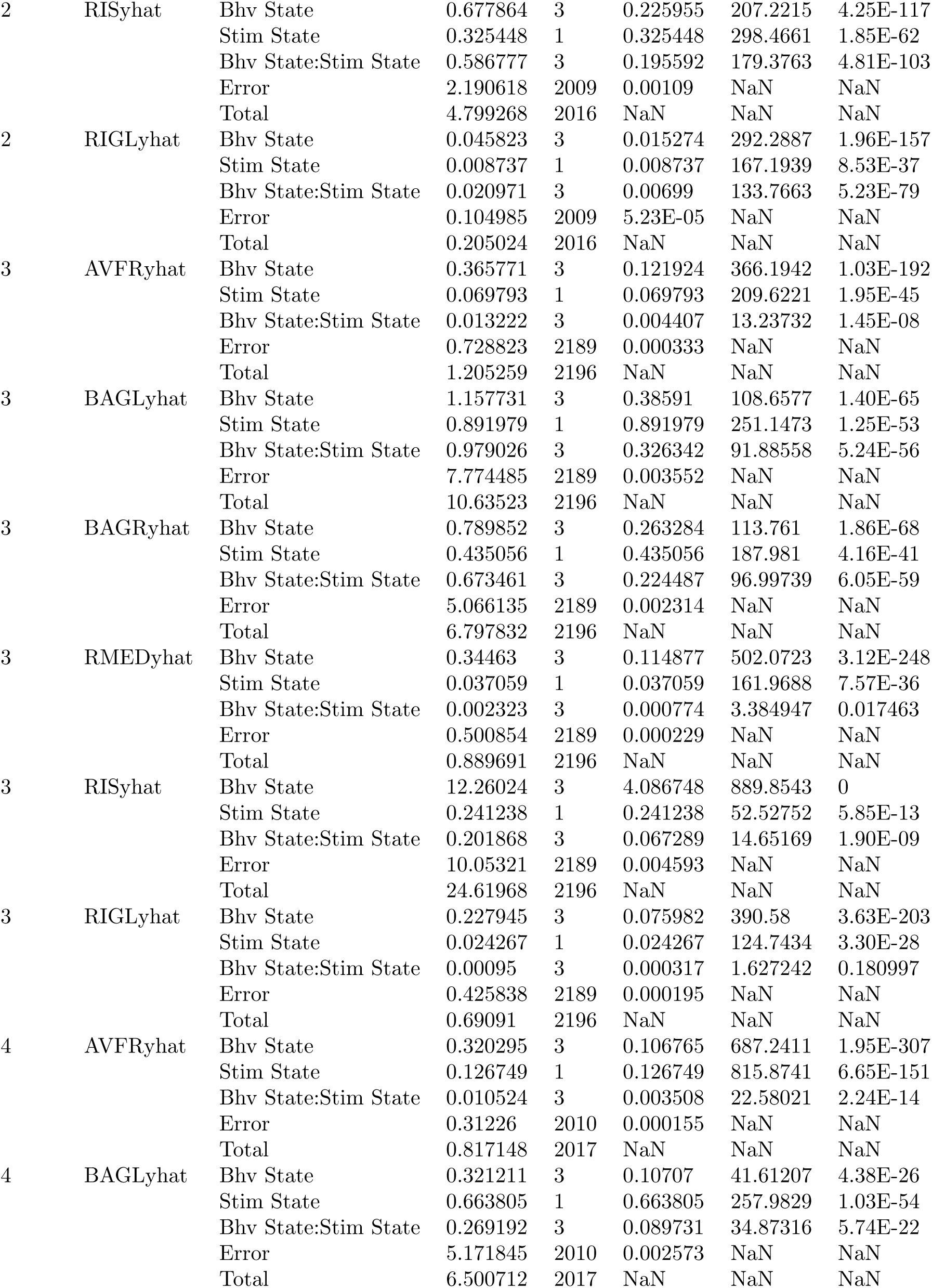
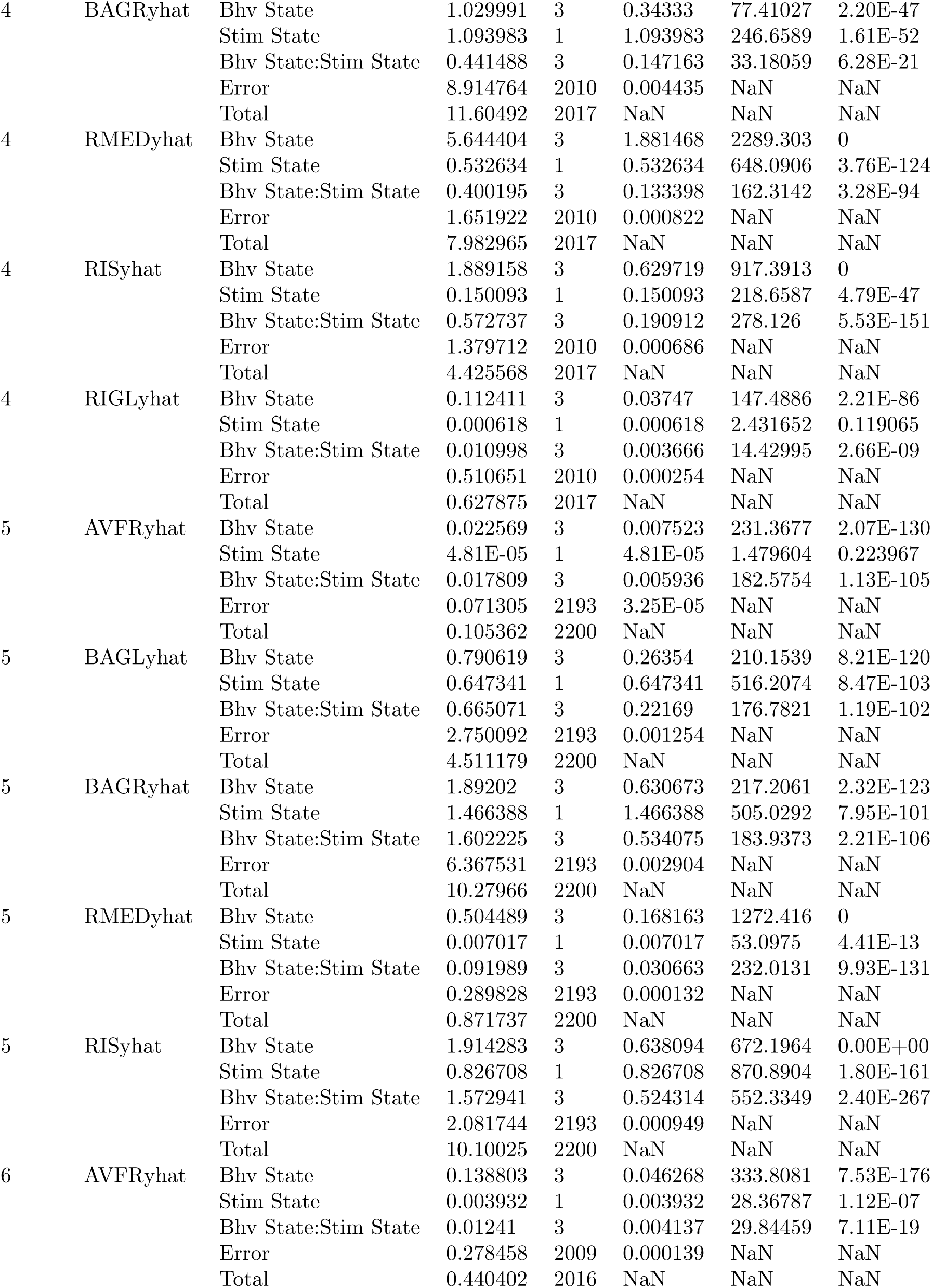
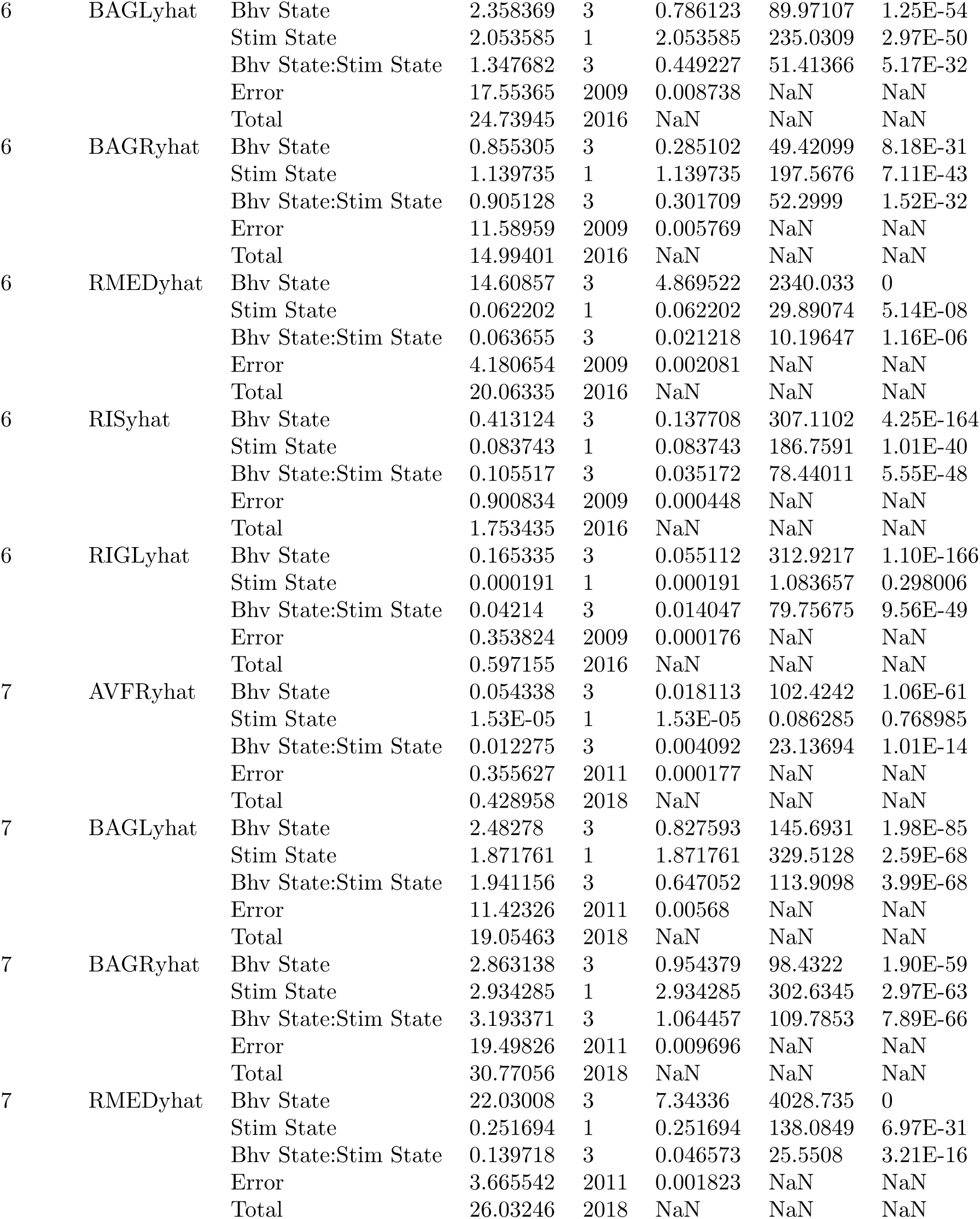
ANOVA tests for AVFR, BAGL, BAGR, RIGL, RIS, RMED, and SABD activity and connectivity vs. behavior and oxygen stimulation factor levels. Glossary: [neuron]yhat: reconstructed activity, Bhv: behavior states (1-4), Stim: stim states (21 percent oxygen only first half trial, or alternating oxygen levels second half), df: degrees of freedom, SS: sum of squares, MS: mean squared, F: test statistic, p: p-value.

**Table 7:**
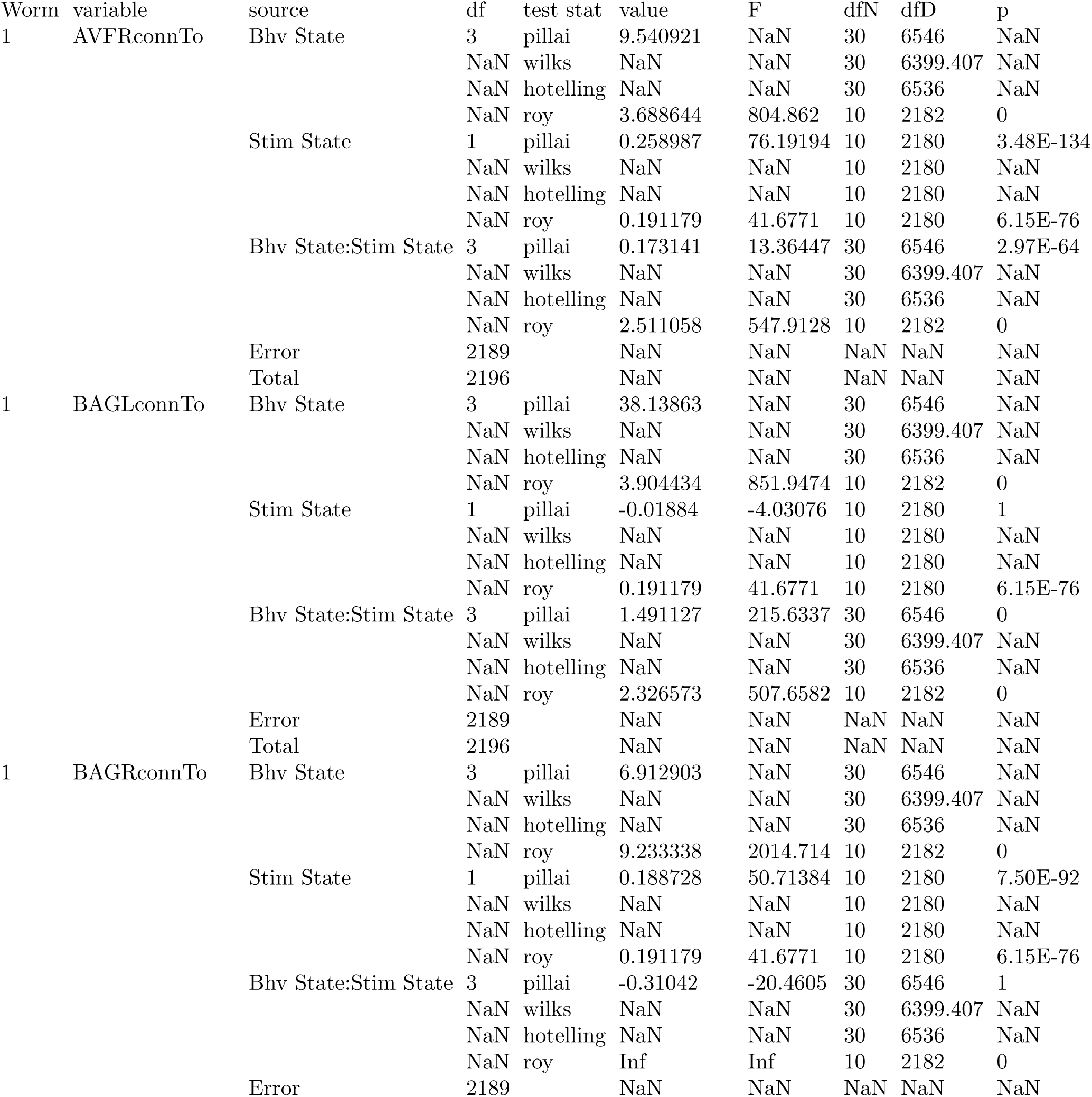
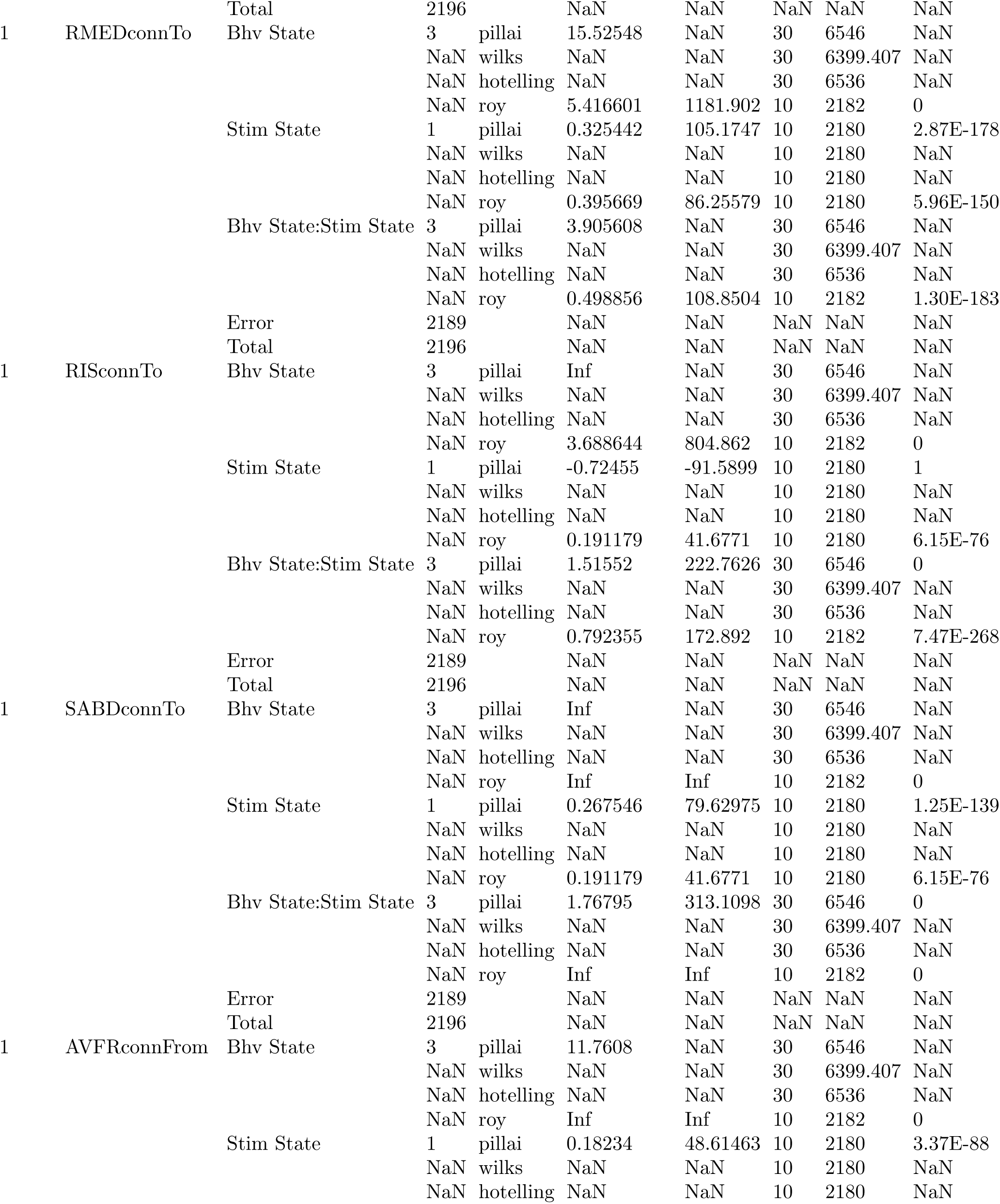
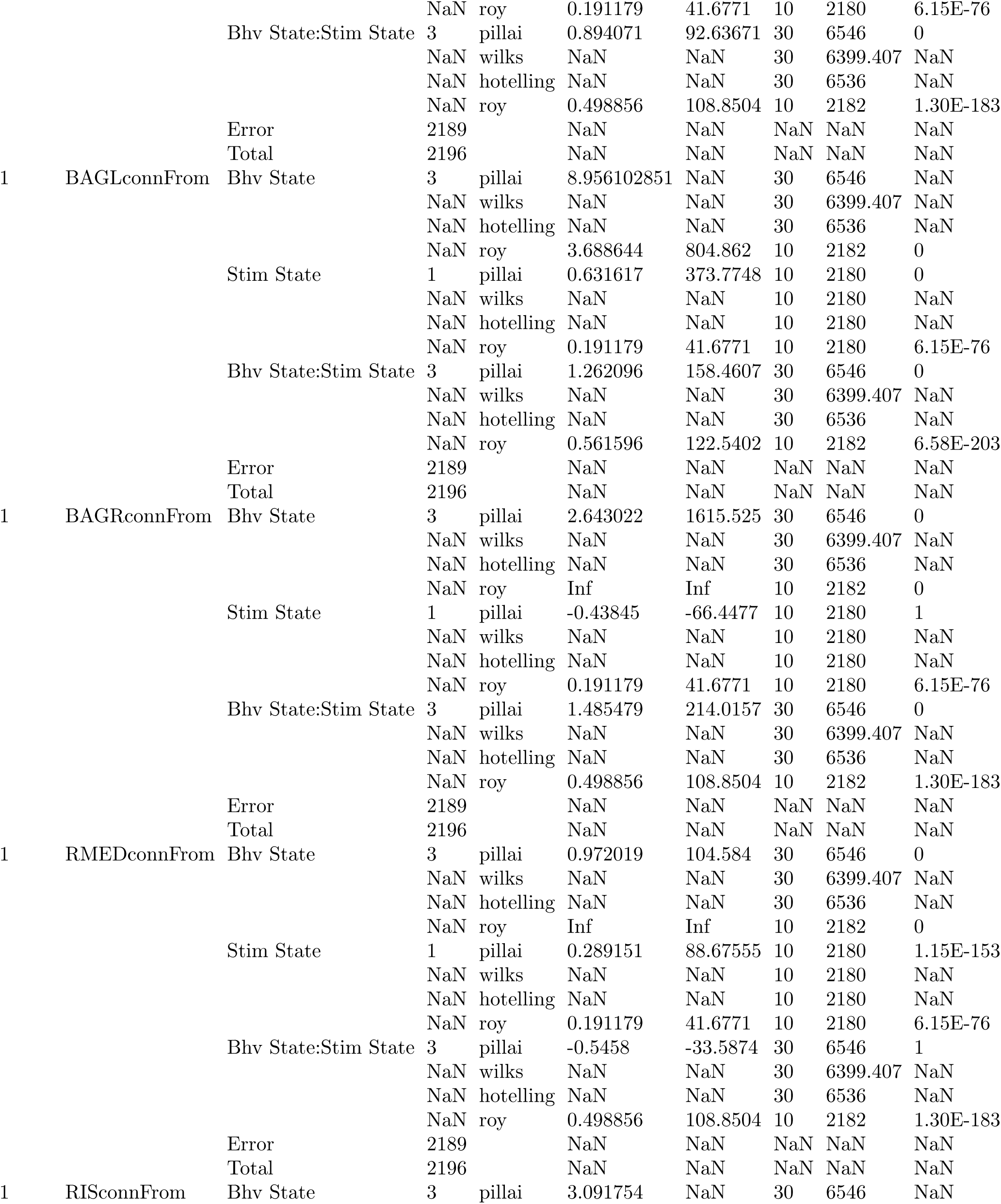
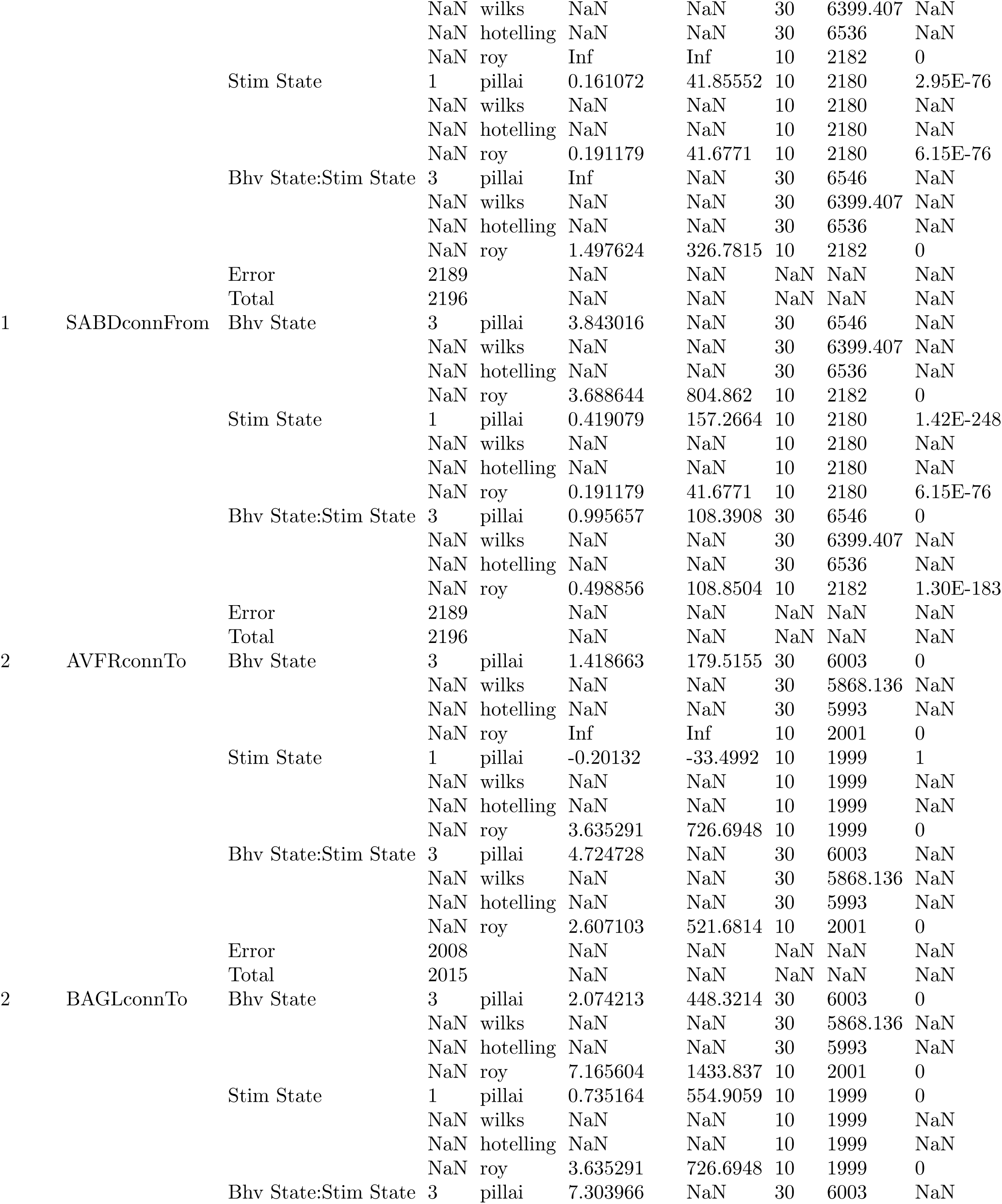
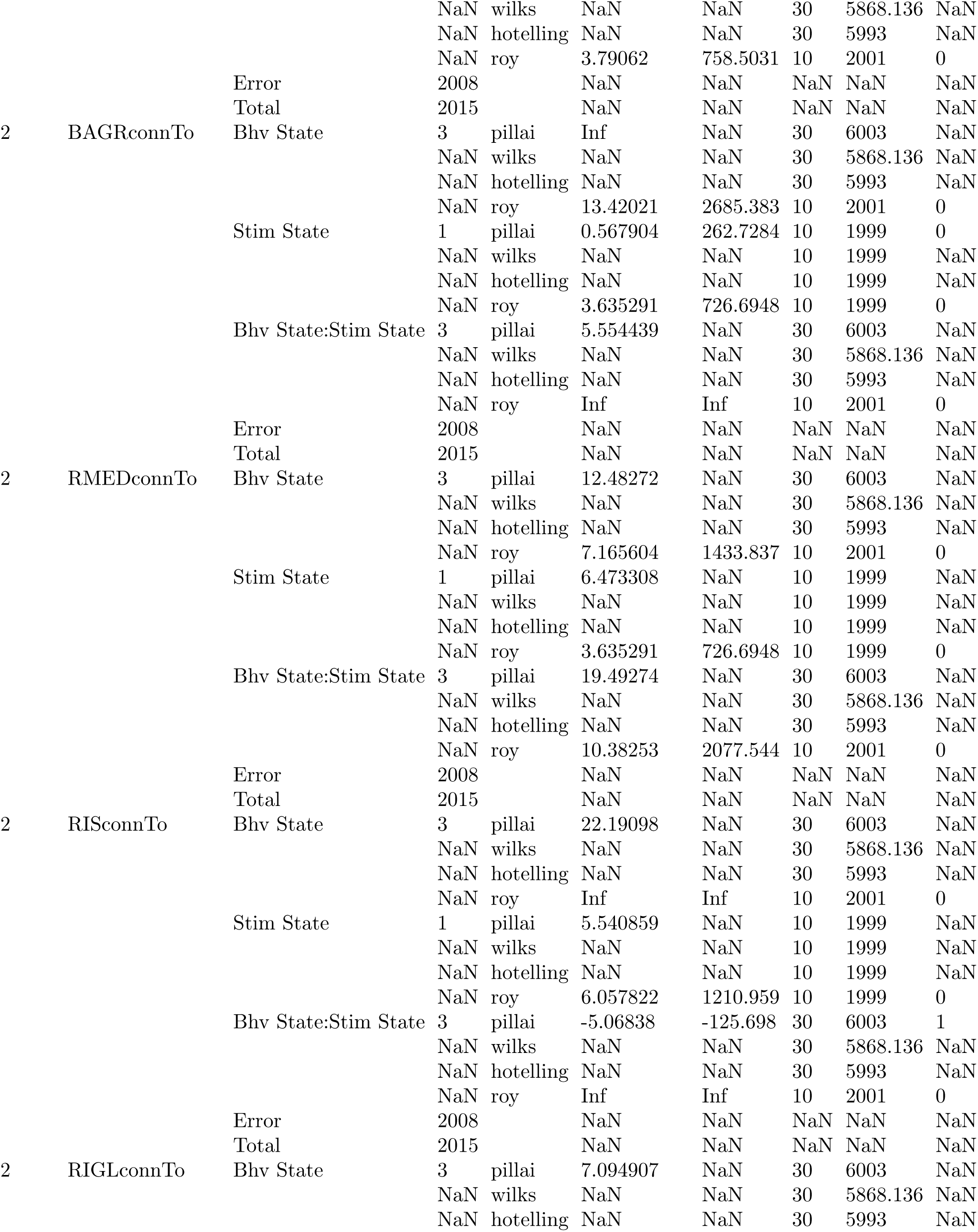
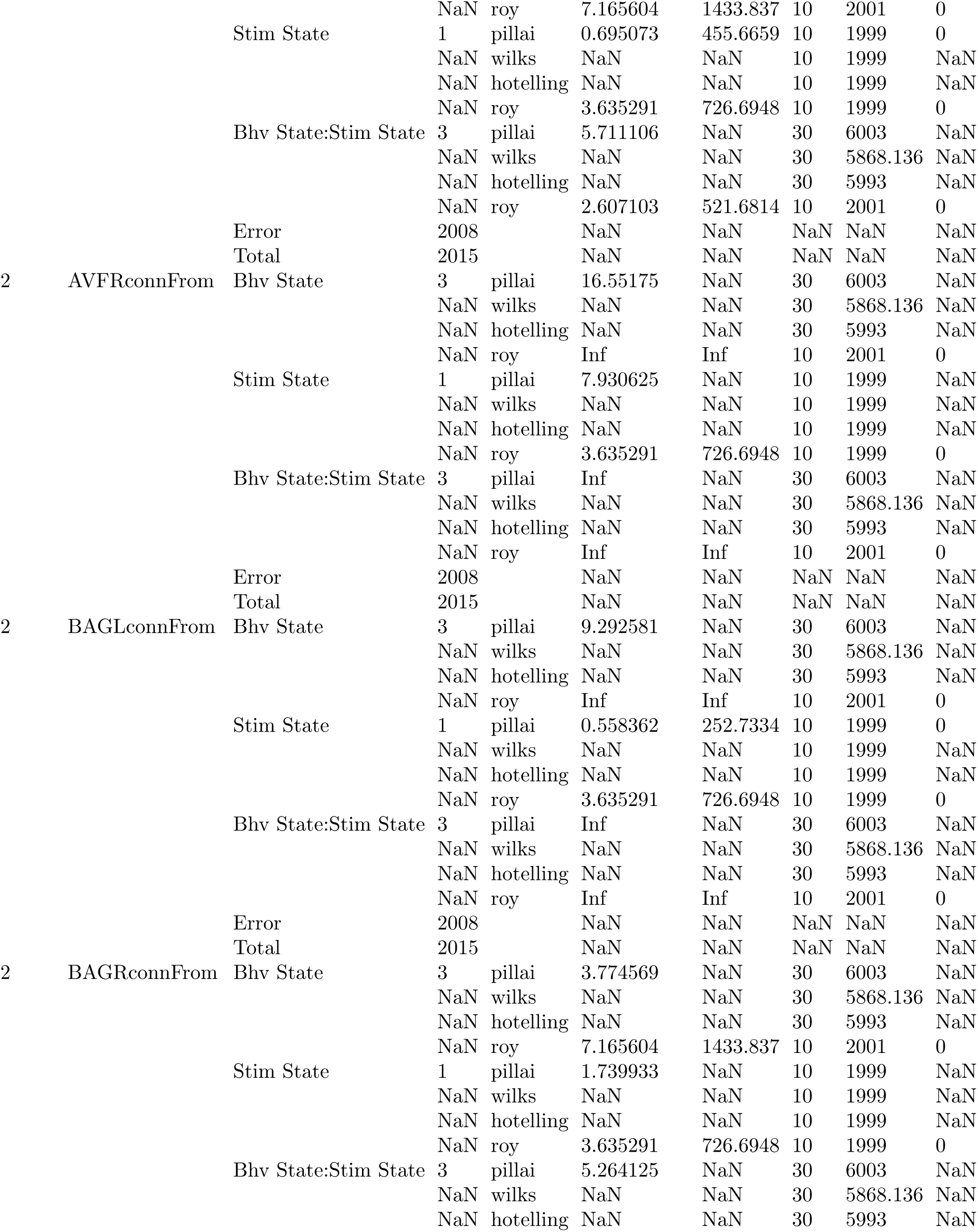
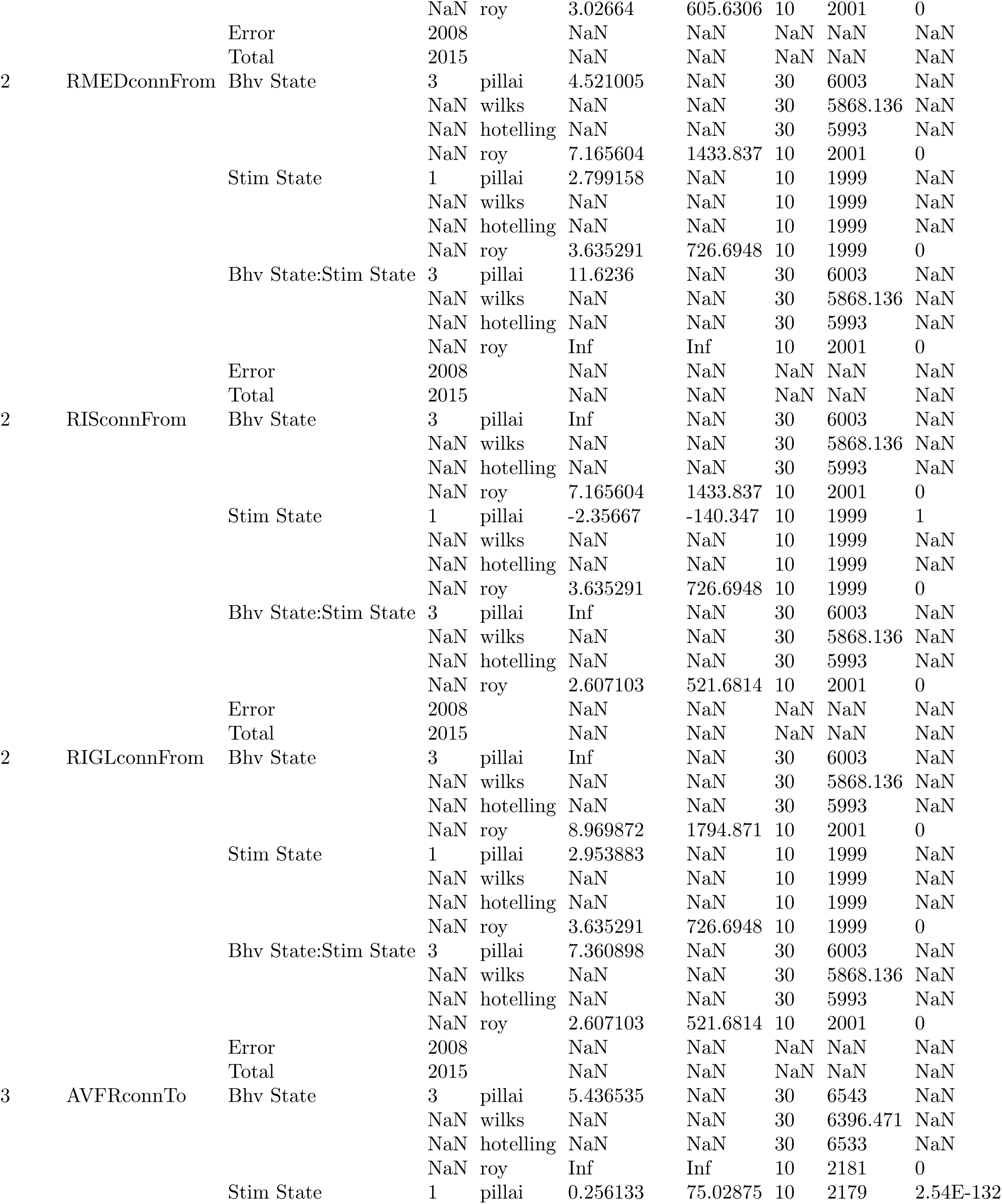
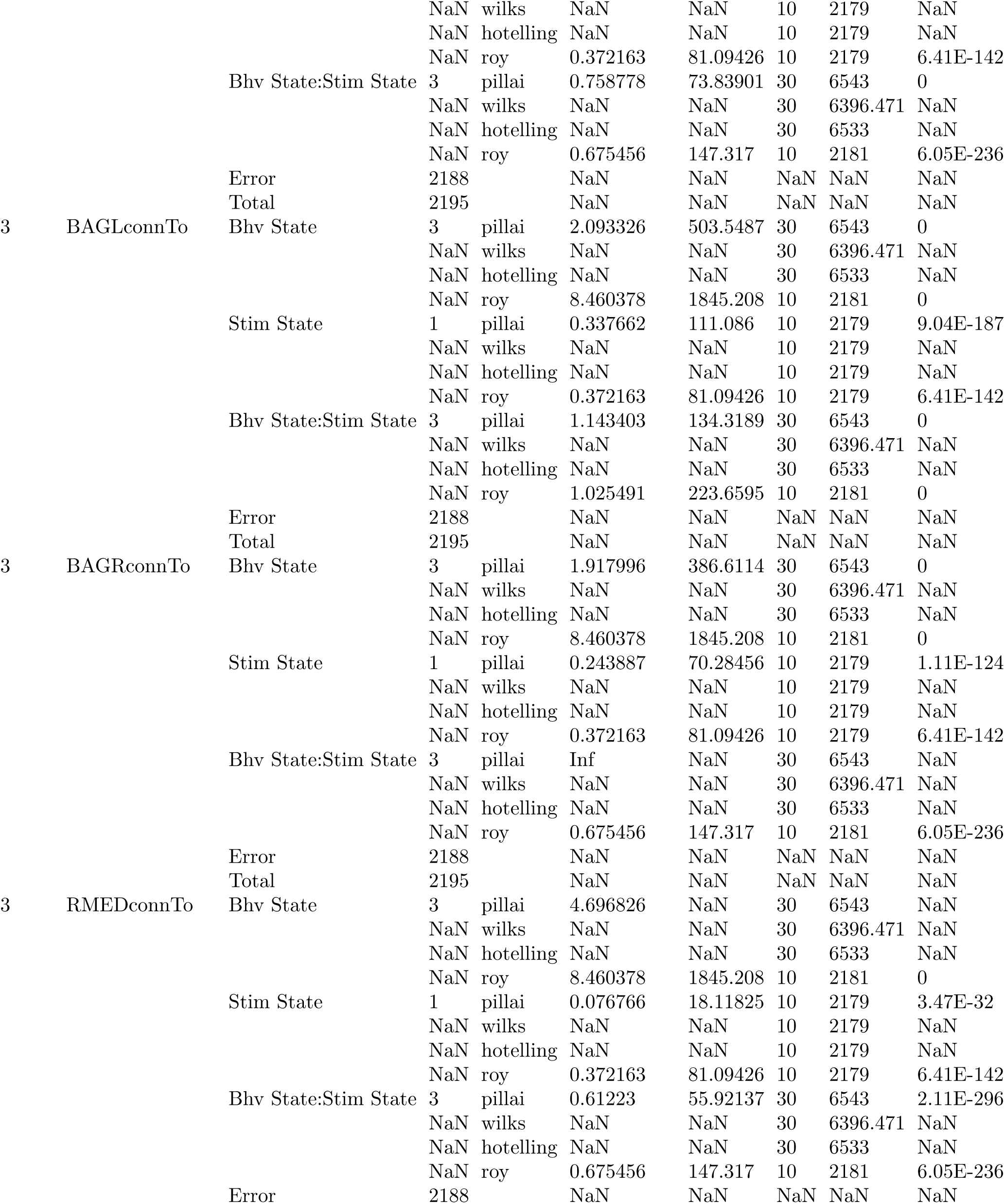
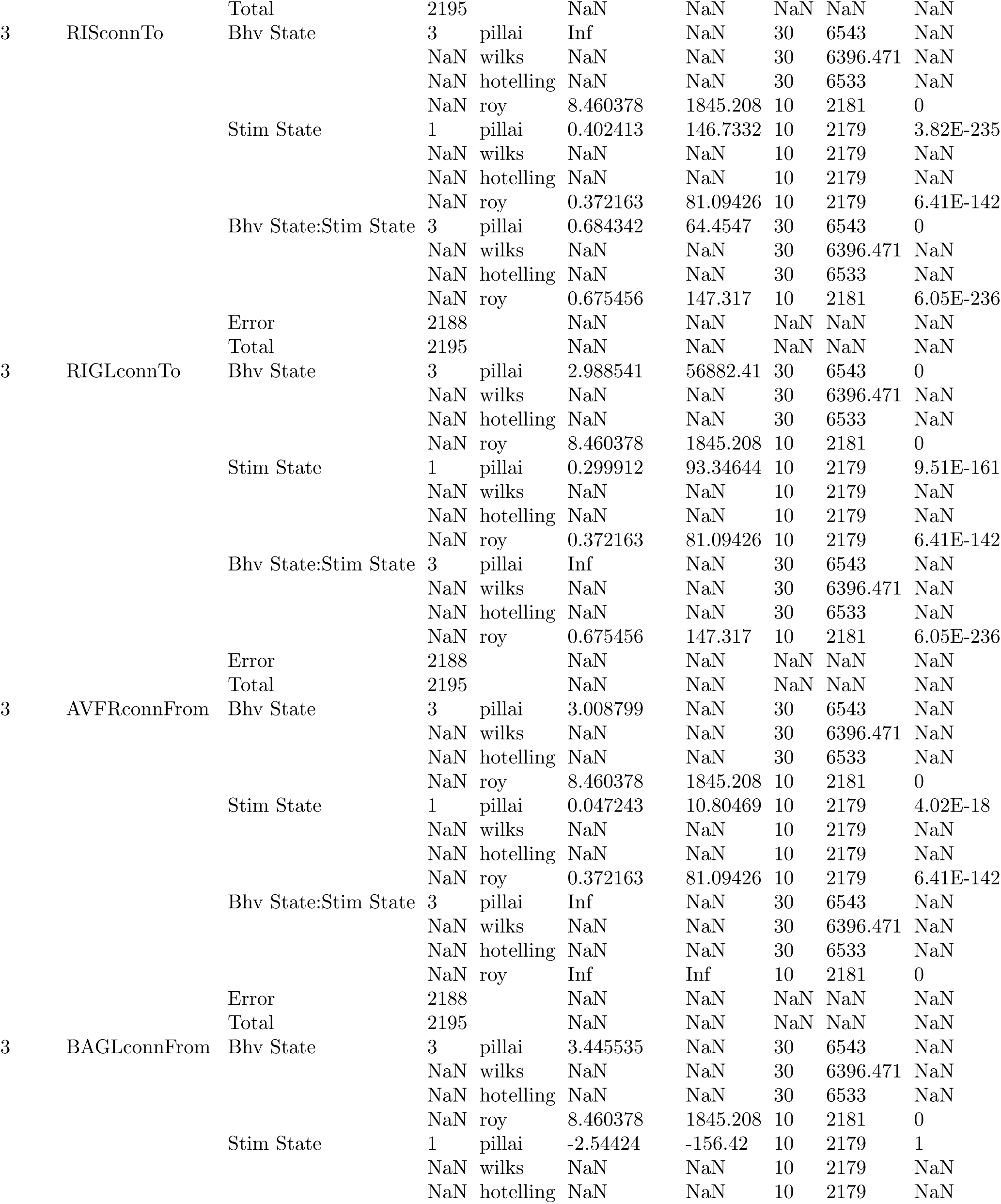
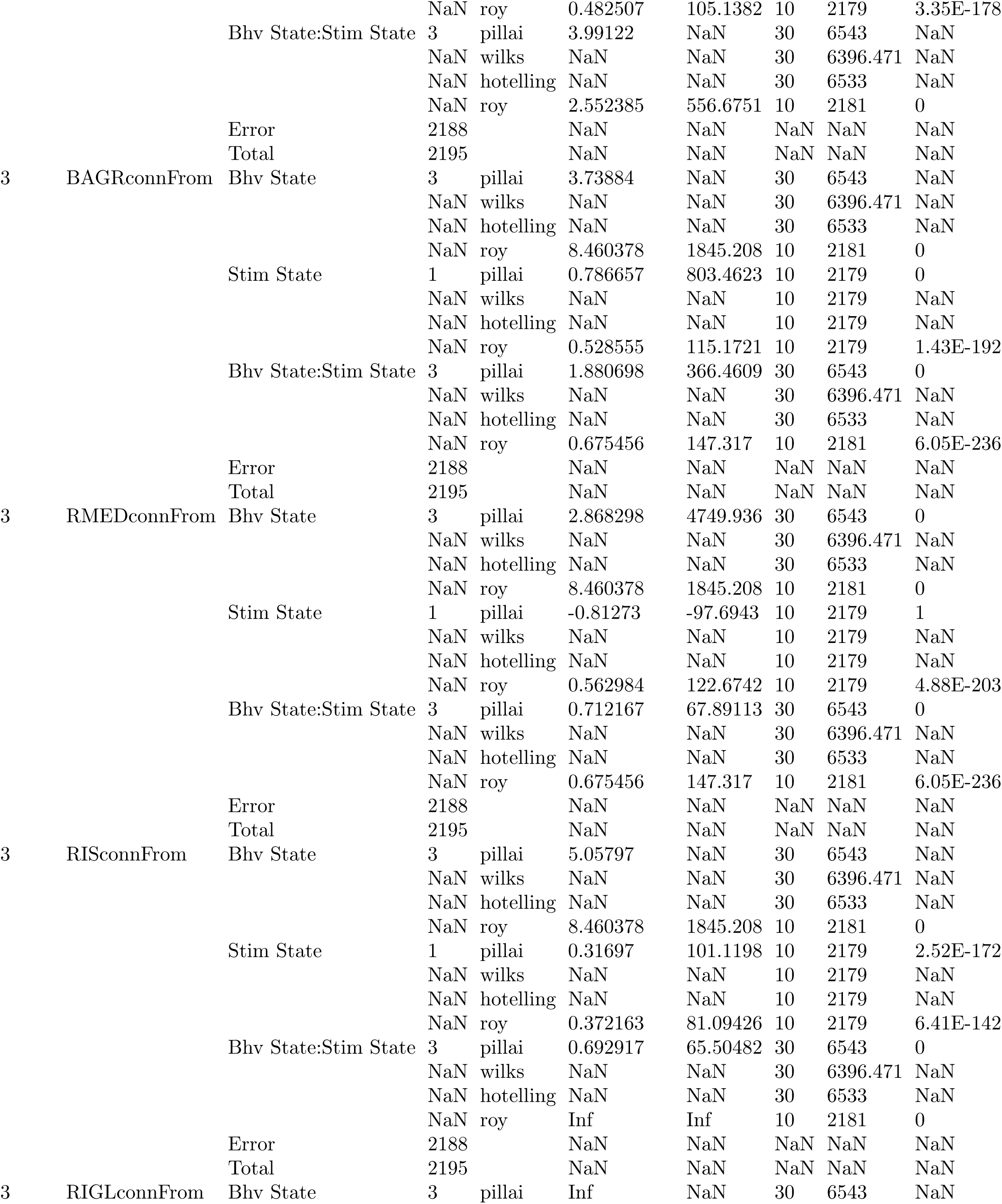
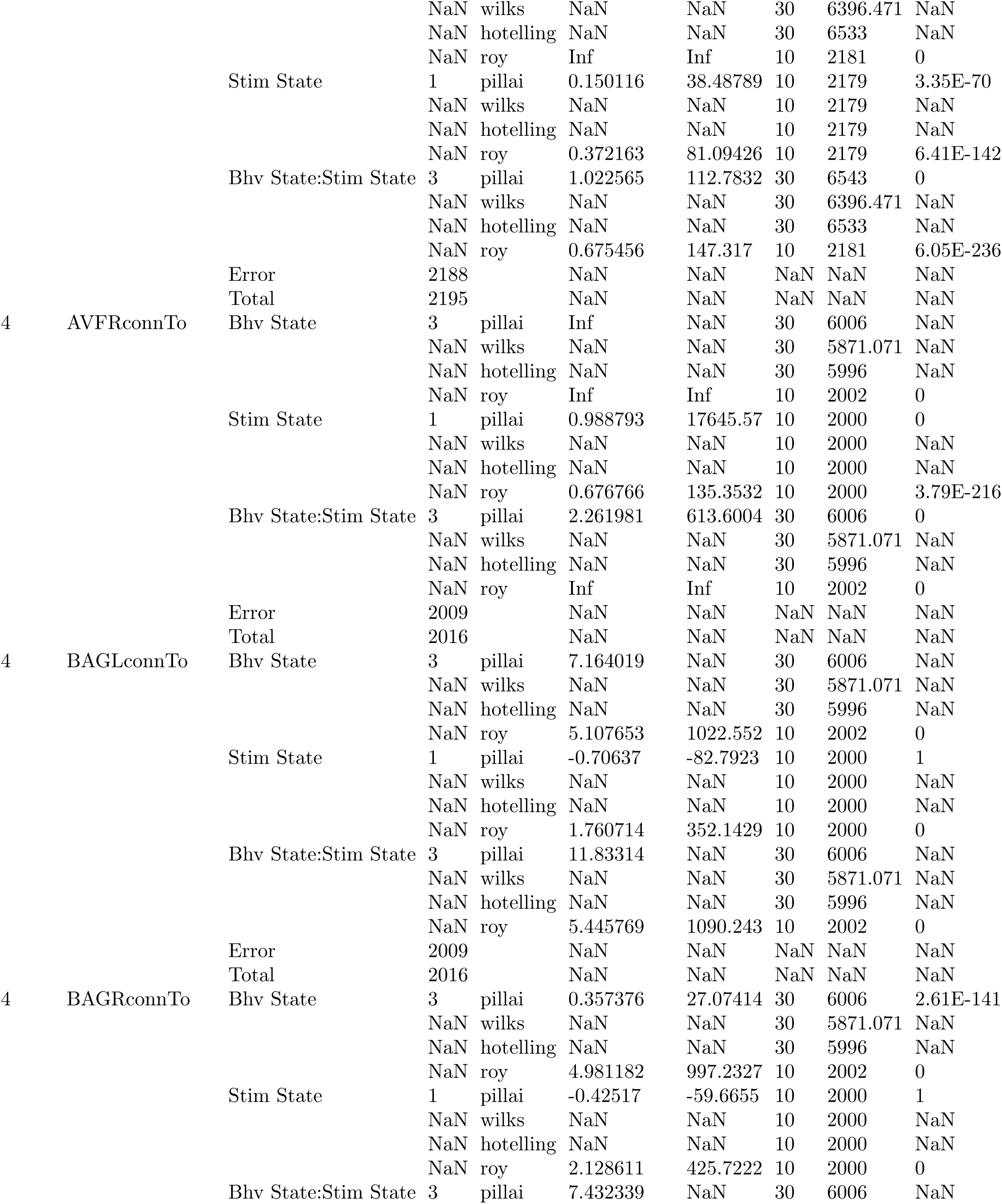
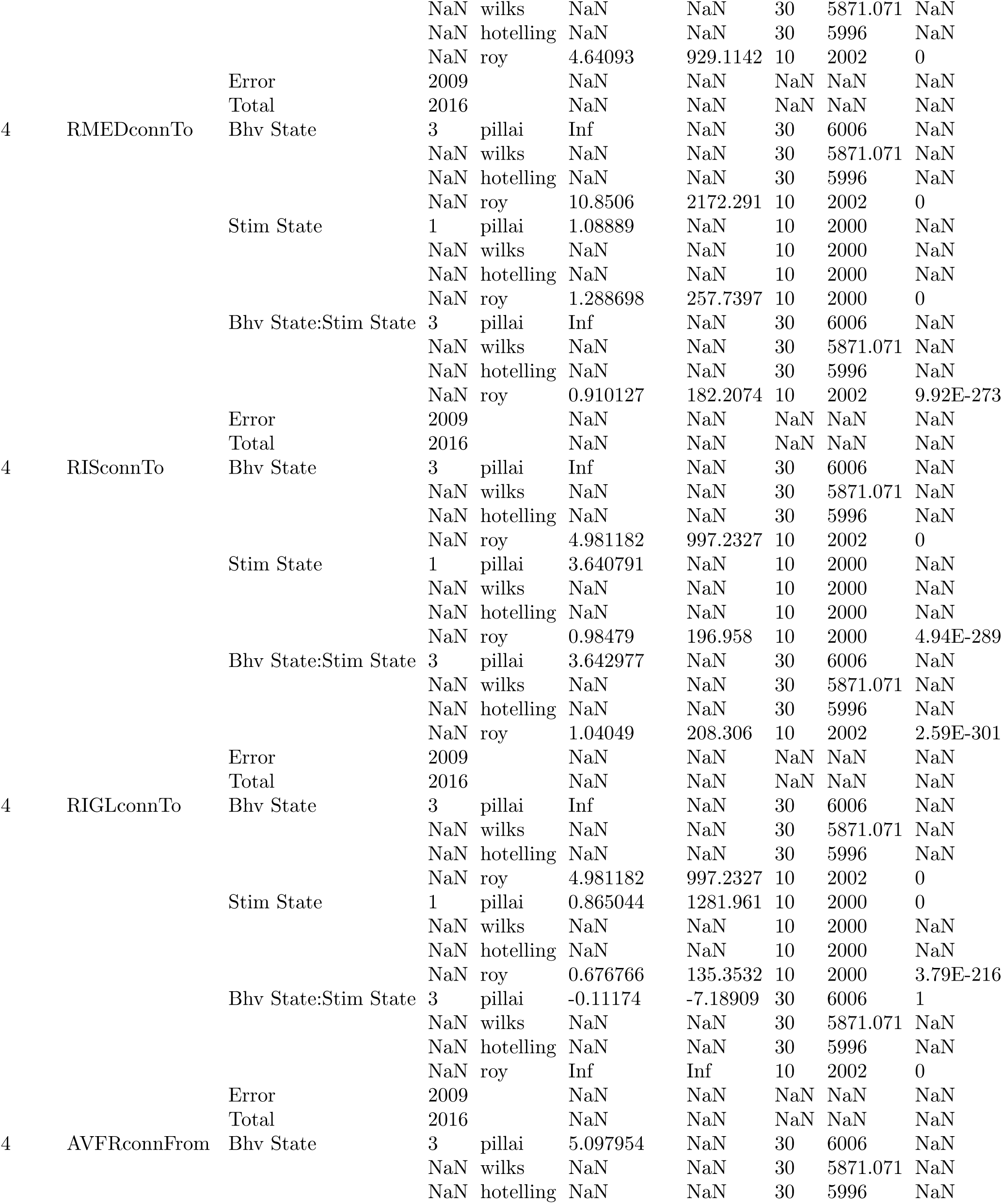
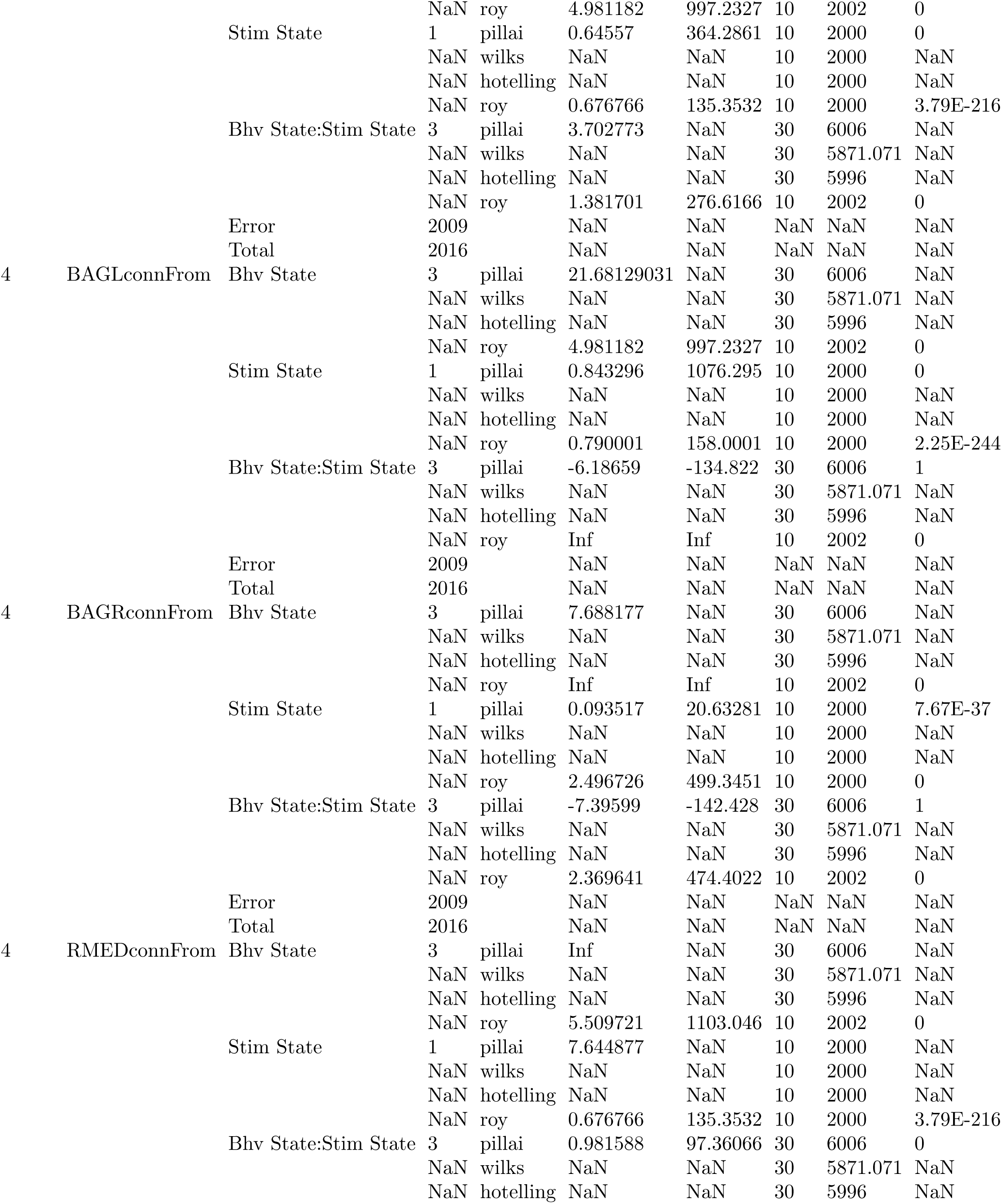
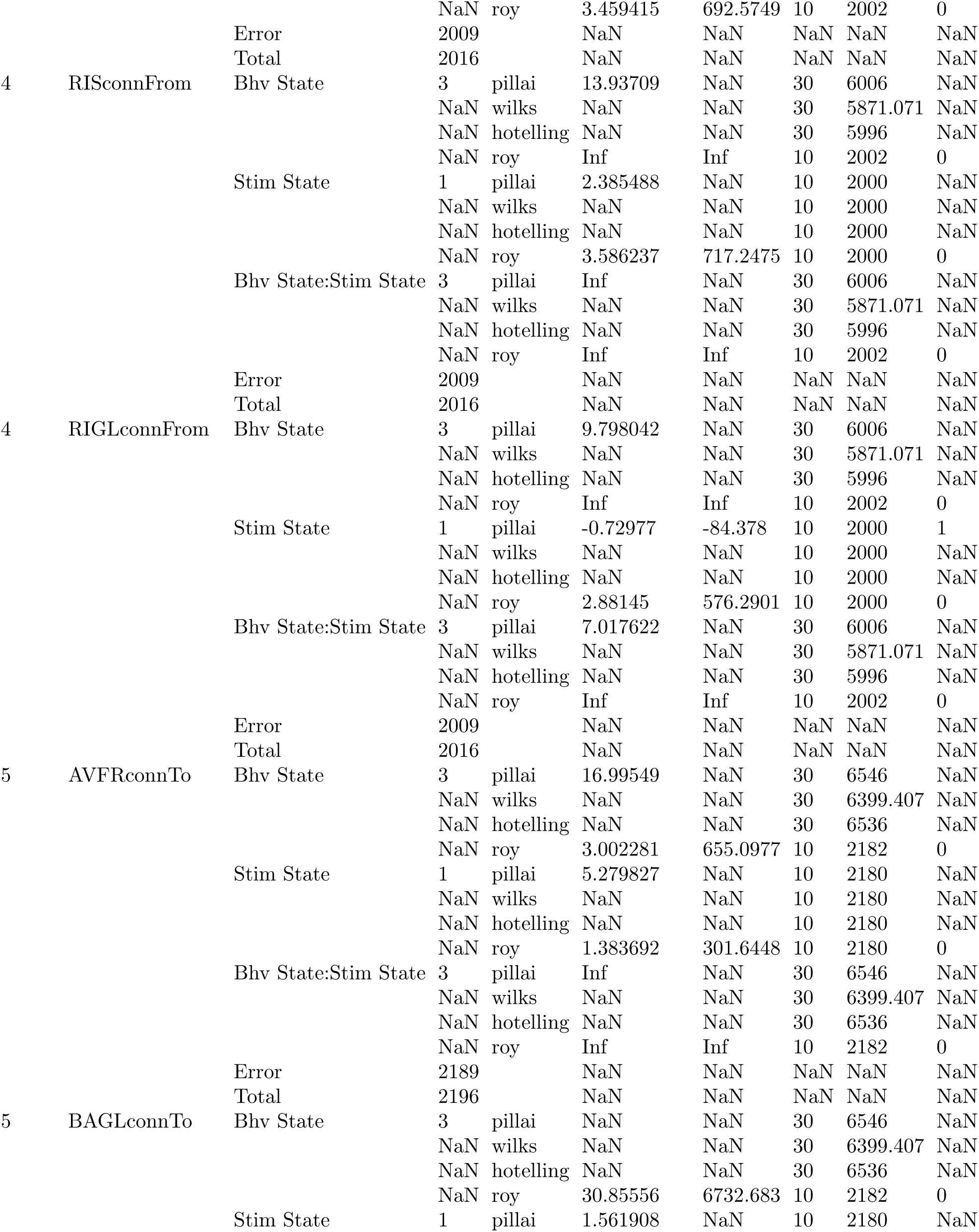
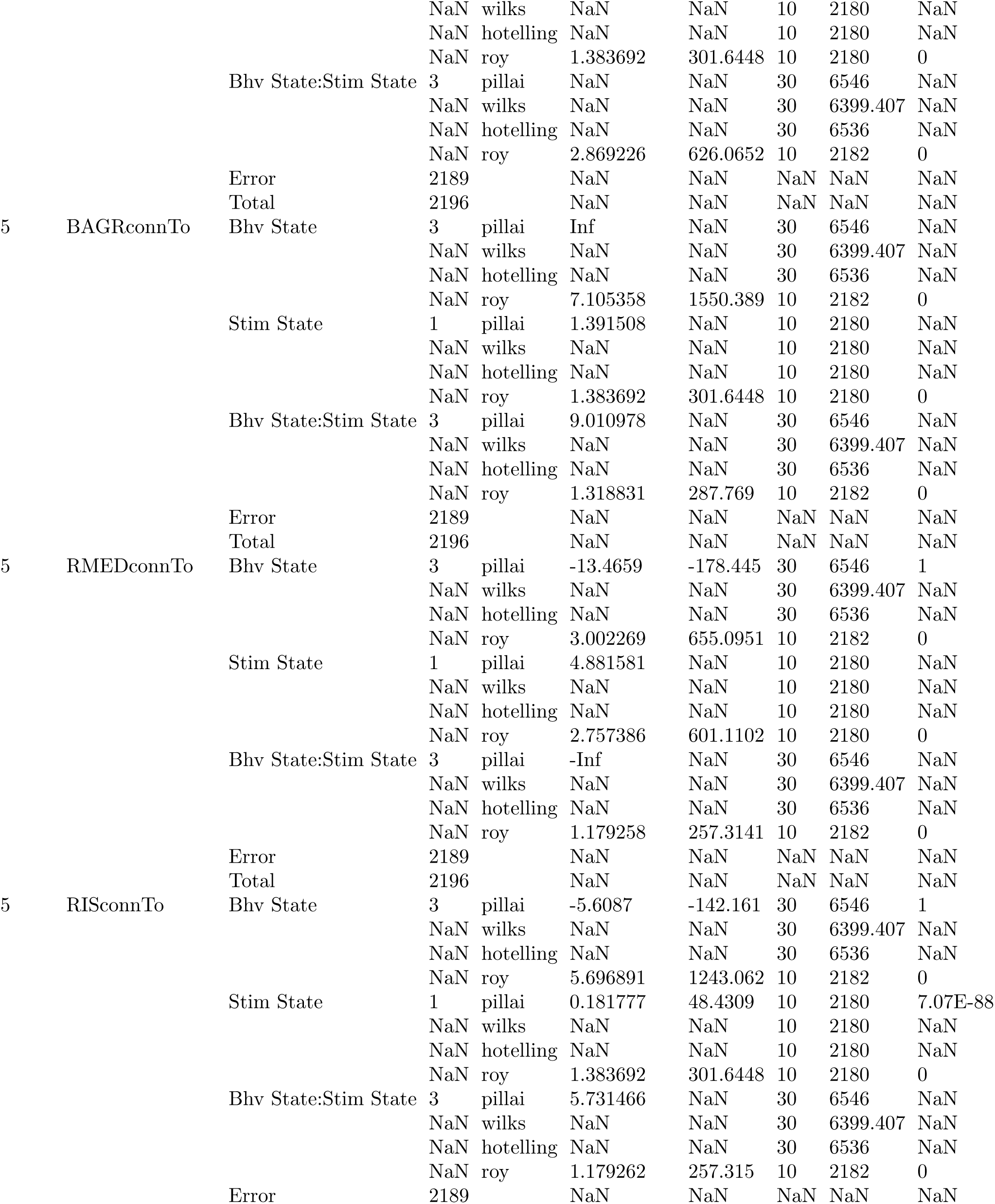
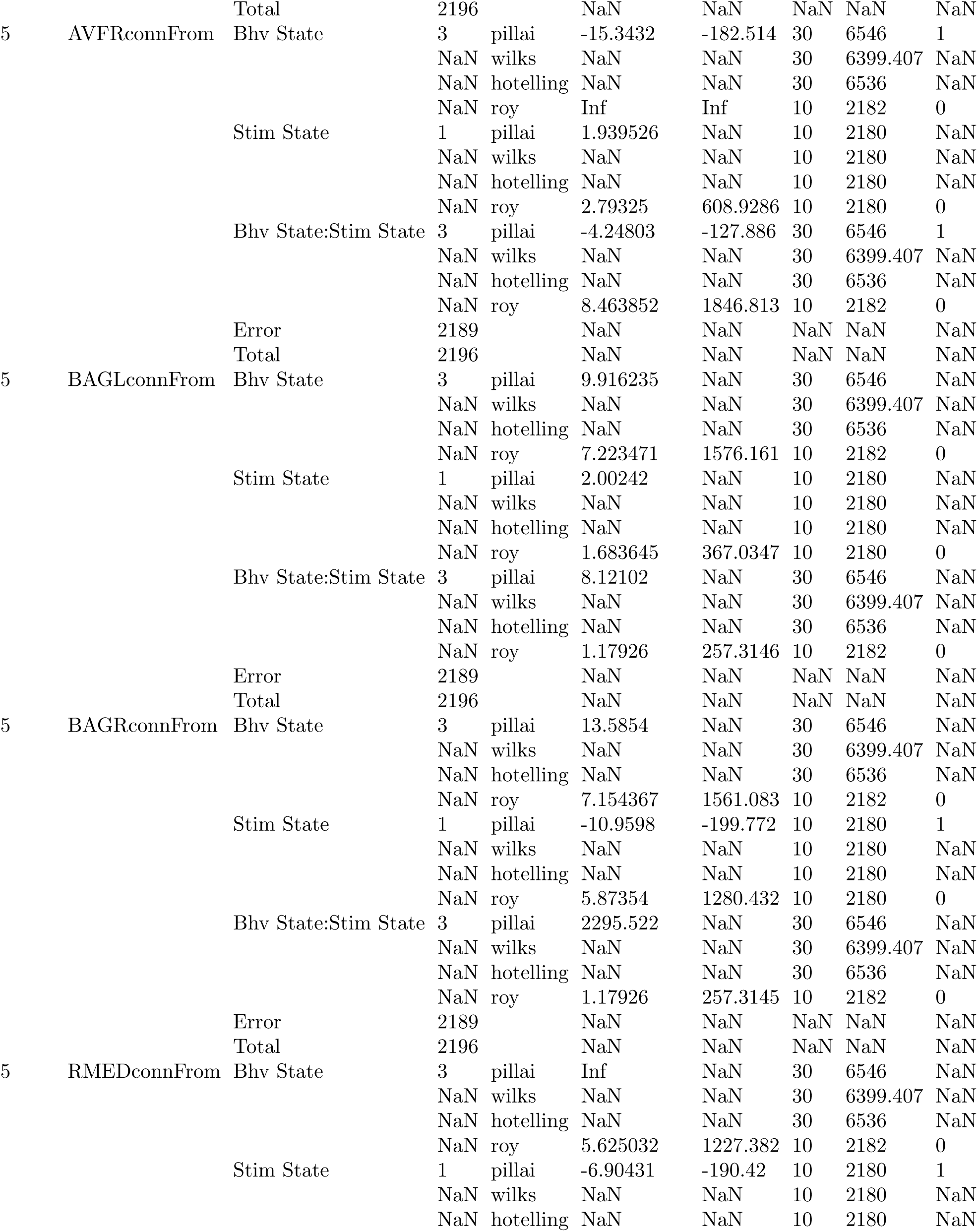
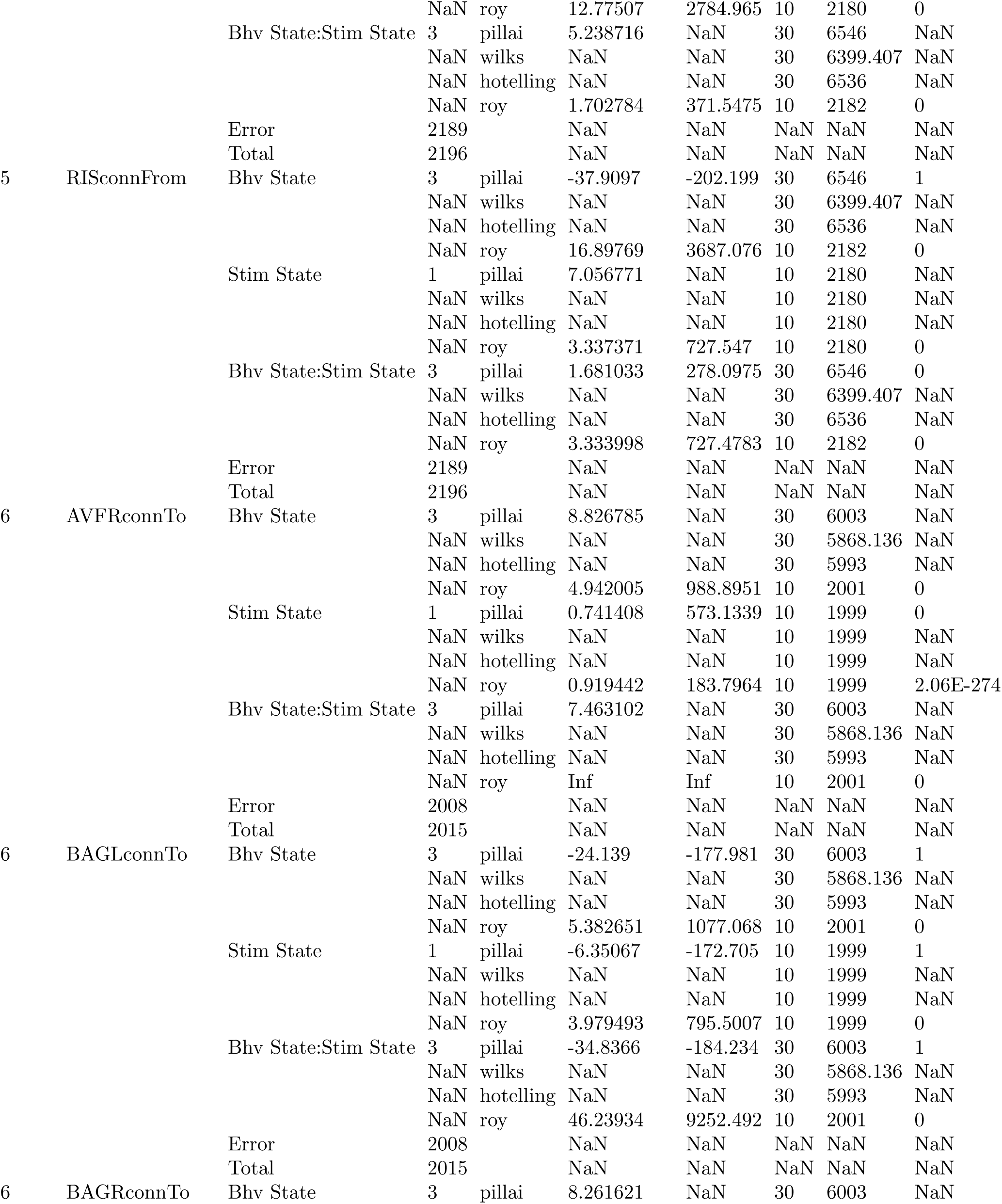
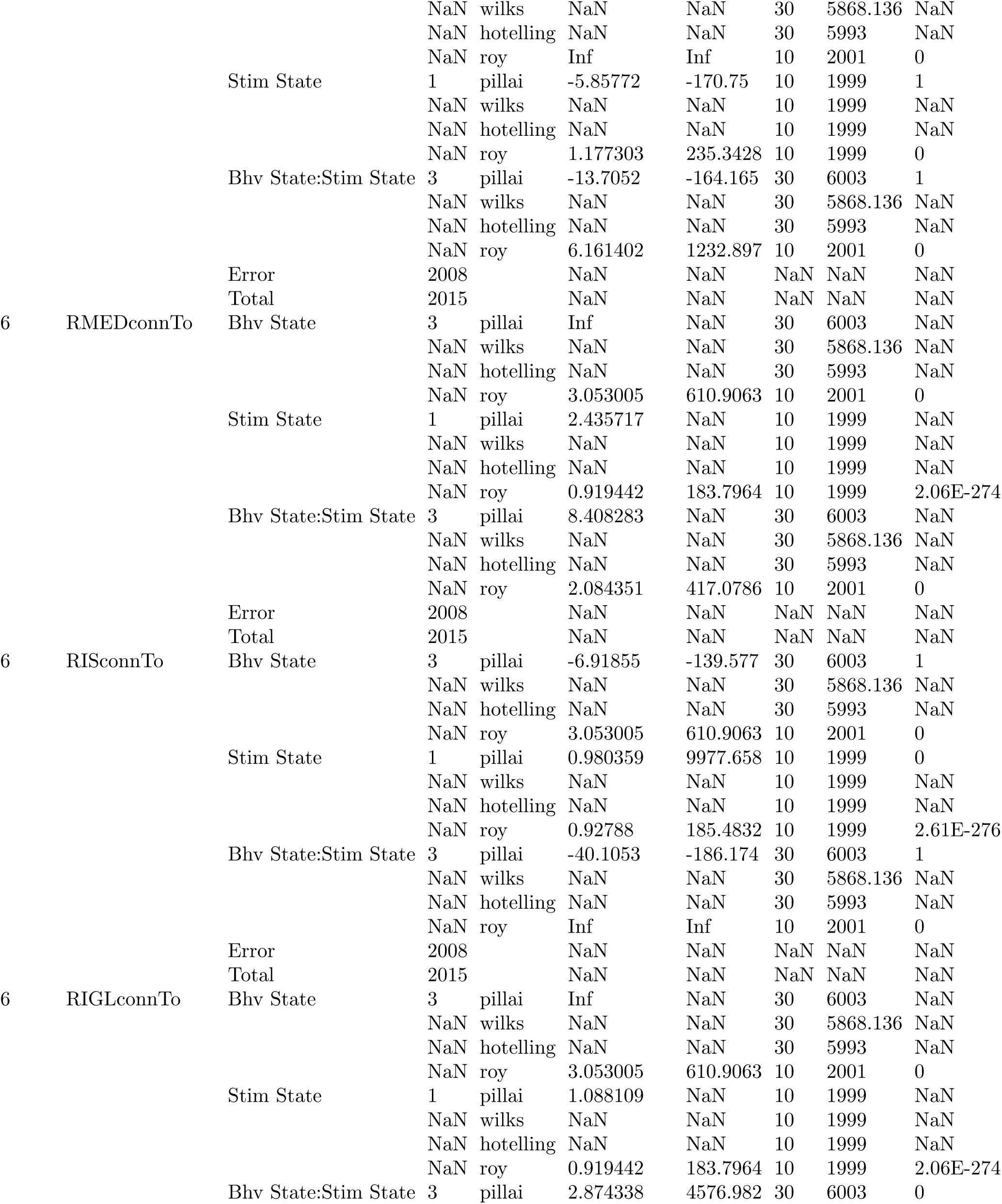
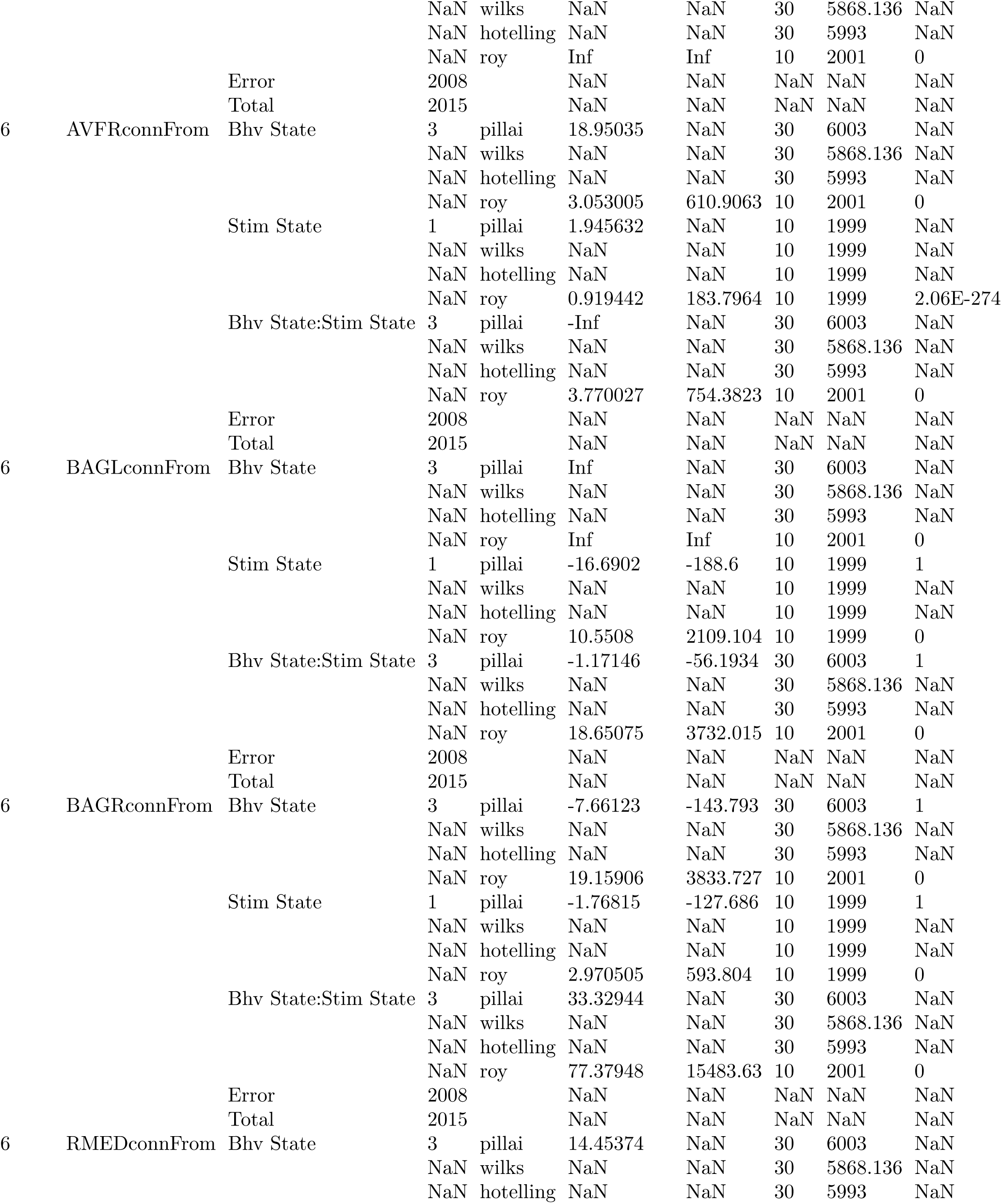
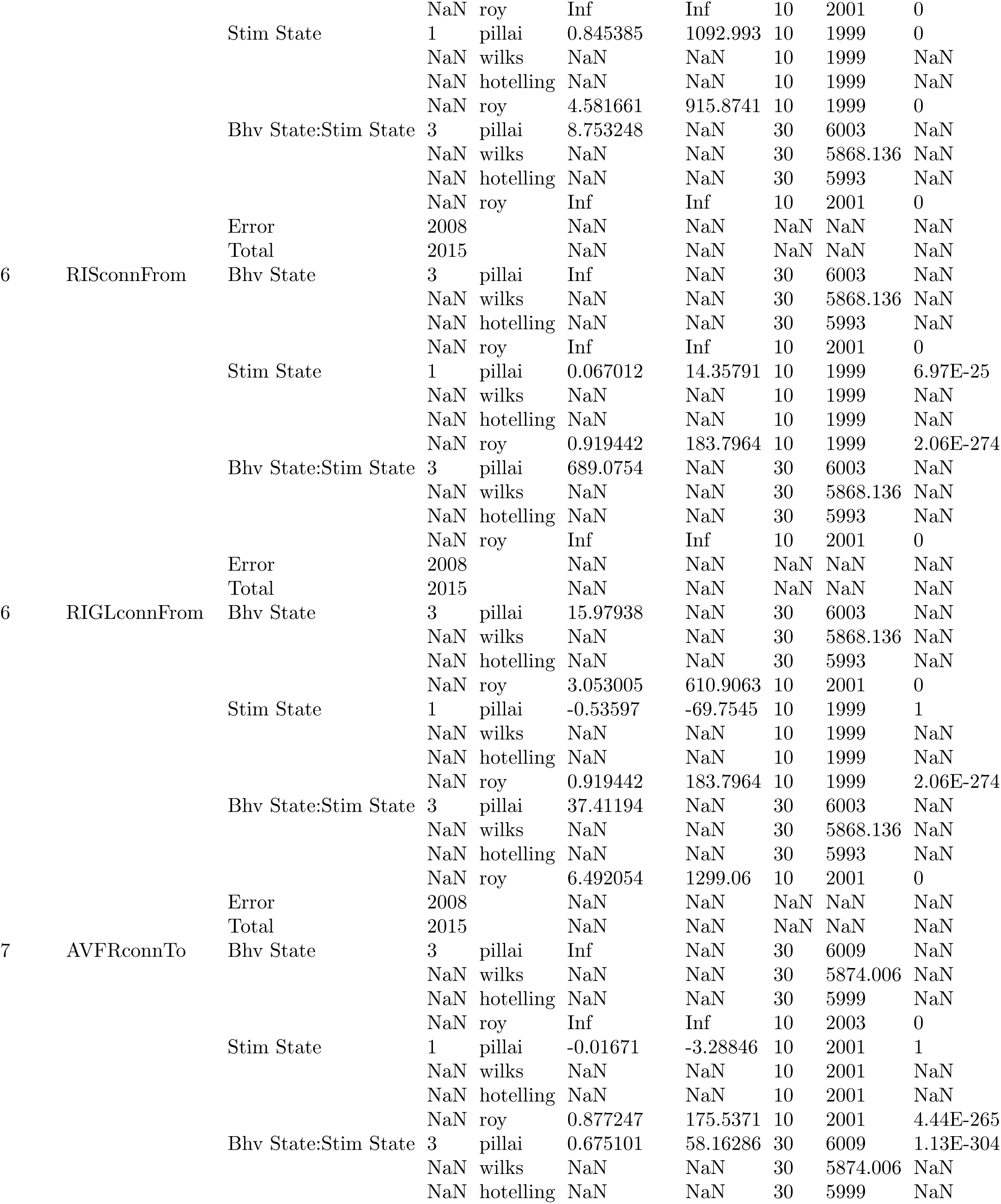
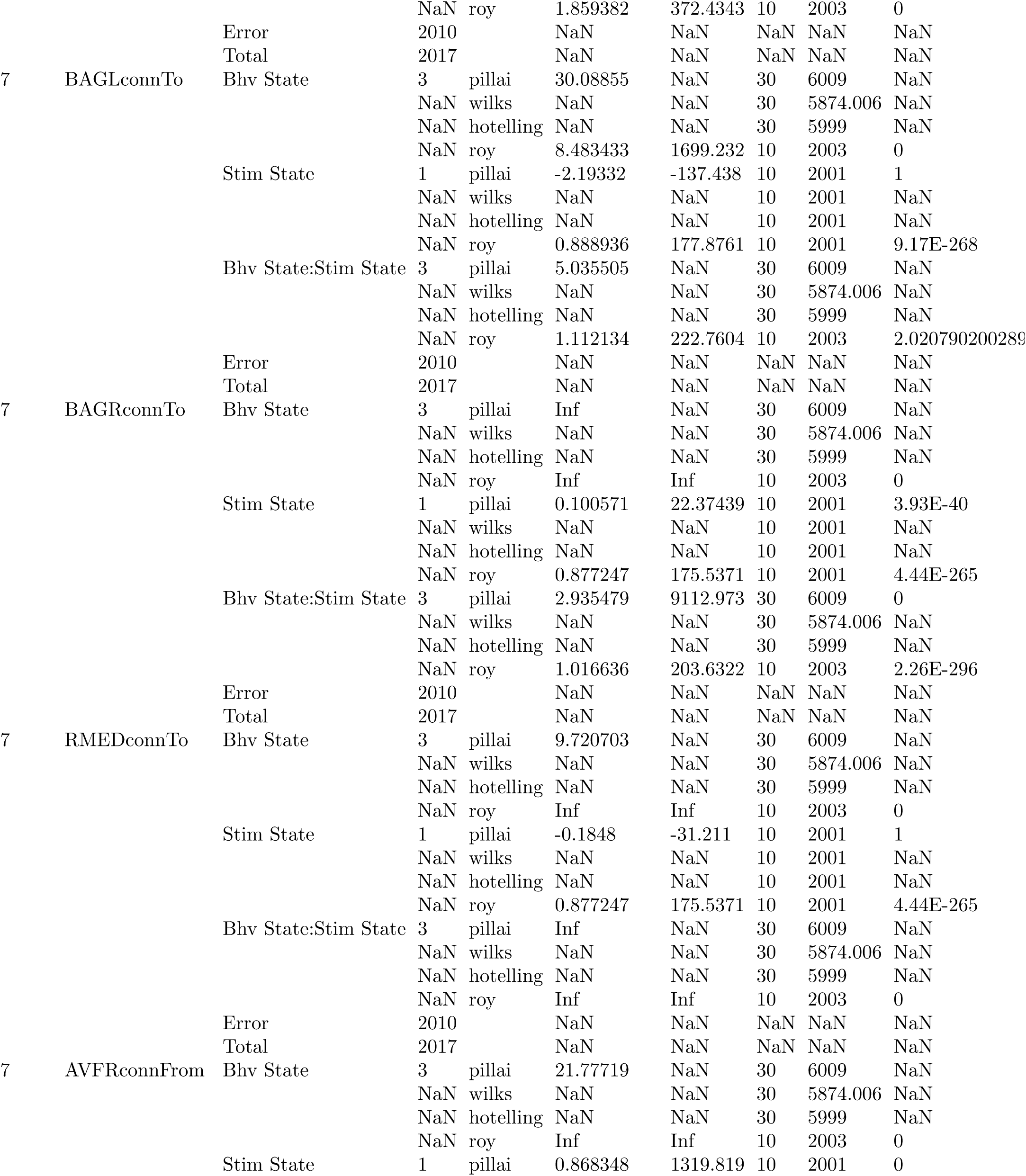
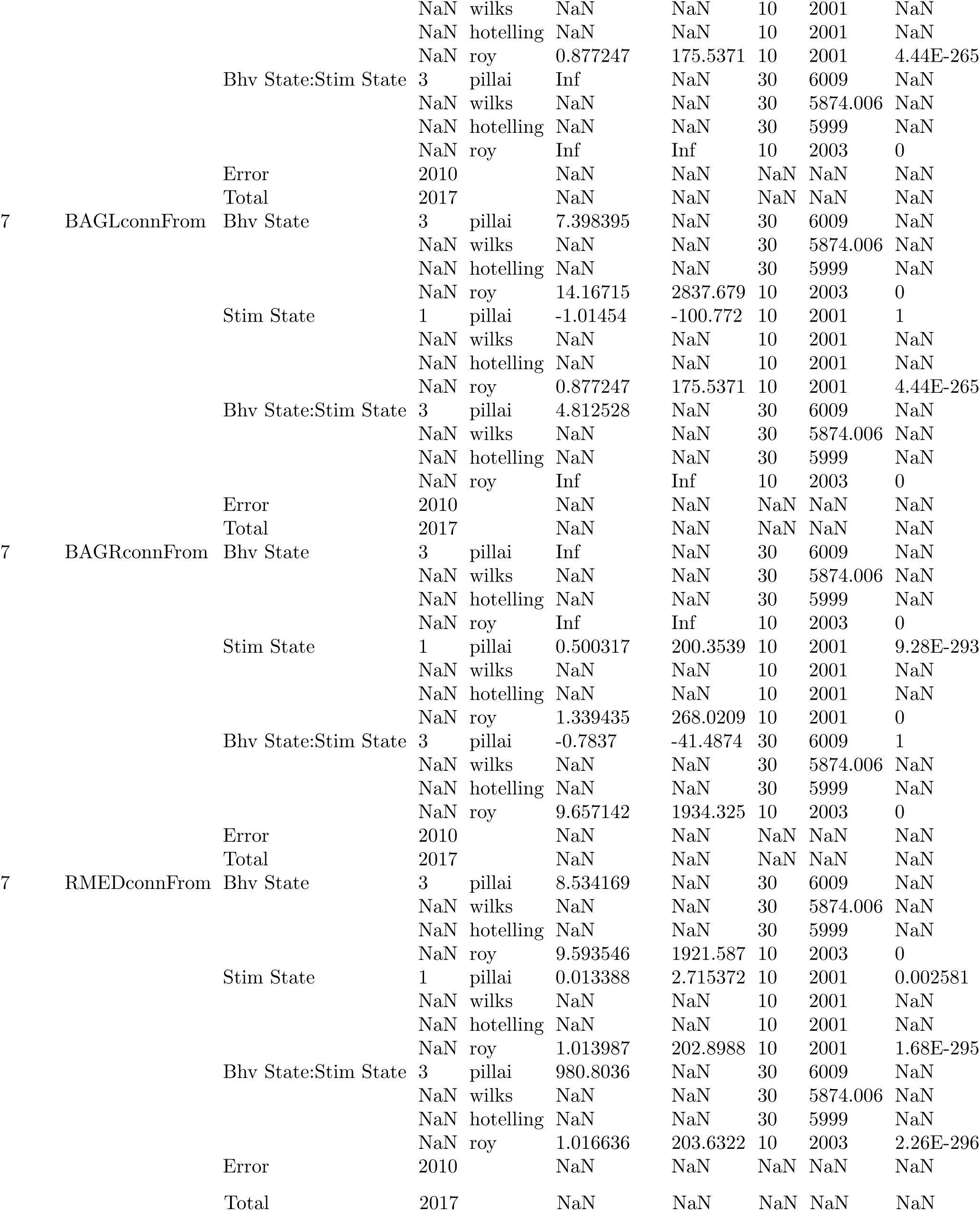
MANOVA tests for AVFR, BAGL, BAGR, RIGL, RIS, RMED, and SABD activity and connectivity vs. behavior and oxygen stimulation factor levels. Glossary: [neuron]connFrom: strongest connectivity value (absolute value) from that neuron, [neuron]connTo: strongest connectivity value to that neuron, Bhv: behavior states (1-4), Stim: stim states (21 percent oxygen only first half trial, or alternating oxygen levels second half), Bhv:Stim: Behavior: Stimulus interaction term, df: degrees of freedom, test stat: test statistic, value: value of test statistic, F: corresponding F value, dfN: degrees of freedom of numerator, dfD: degrees of freedom of denominator, p: p-value.

Considering the example of the interneuron AVFR (Fig. 6), we saw evidence of how these behavior and stimulation effects play out differently compared to sensory and motor neurons, and that dynamic encoding of behavior in this interneuron really depended on the overall, dynamic state of the animal.

Whereas the connectivity to and from sensory neuron BAGR was somewhat correlated with behavior and was strongly correlated with sensory information (Fig. 4), AVFR’s connectivity appeared to correlate strongly with behavior cycles, but transitioned into a different regime (negative values) during the acute hypoxia periods, where connectivity to and from AVFR was also strongly influenced by the stimulation. Likewise, whereas the connectivity to motor neuron RMED was hierarchically correlated to behavior state and stimulation state but strongly sent out behavior-related connectivity, the connectivity to and from interneuron AVFR was strongly correlated to both behavior and stimulation. Thus, we can hypothesize that AVFR participates strongly in relaying both sensory and motor information, and can dynamically change functional regimes.

### 2.5 *Validation:* dLDS dynamic connectivity maps identify novel connections compared to stationary anatomical and functional connectivity atlases

While the *C. elegans* anatomical connectome has been available for over a decade [7], questions remain as to context-dependent functions, nonlinear relationships, and differences between individuals in terms of how those neurons work together. More recently, the pairwise stationary functional connectome has been mapped by stimulating each neuron and observing the responses of every other neuron individually [11]. Along with greater understanding of the role of long-range neuropeptide signaling [15; 17], this functional connectome work has revealed novel relationships that are not mediated by direct synapses.

Using dLDS, we investigated dynamic connectivity, which we define as the graph where each vertex is a single neuron and each edge from nodes *i* to *j* is the directional local influence of neuron *i* on neuron *j*. As dLDS is a nonstationary model, this definition of connectivity allows each edge to be time-varying and to capture the changes in network structure across behaviors. Moreover, the ability of dLDS to represent nonlinear dynamics can further capture effects such as two or more neurons needing to be active to activate another downstream neuron (AND gate)—an effect that cannot be determined from one-to-one stationary functional connectomes.

We compared the dynamic connections found by the subset of *C. elegans* head neurons recorded in this dataset to the aforementioned anatomical and functional connectomes. The magnitudes of dynamic connections that were also found in the respective reference were kept positive (red), while connections that were unique to the dLDS model were given a negative coefficient (blue). We observed that, across the Stim and NoStim worms, a majority of the connections identified by dLDS did not match those in the anatomical or functional connectivity reference atlases, although some connections did (Supplementary Figs. 34, 35, 36, 37). These dLDS-only connections could be avenues for future exploration.

### 2.6 *Novel capabilities:* dLDS enables the alignment of latent manifolds across individuals

Per-worm models are useful for characterizing the dynamics of individual worms and even show some commonalities in terms of the dynamic connectivity maps across worms corresponding to different behavioral states or environmental variables. However, we wanted to investigate how a *shared* latent neural manifold might reflect more general principles of dynamic connectivity across worms. We modified dLDS to learn one set of DOs across all 7 Stim worms, with an individual observation matrix for each worm, resulting in one aligned model per worm (Fig. 7). We kept all the same hyperparameters as for the individual models, with the exception of allowing 70 DOs (thus permitting the possibility of 10 completely unique operators per worm).

**Figure 7:**
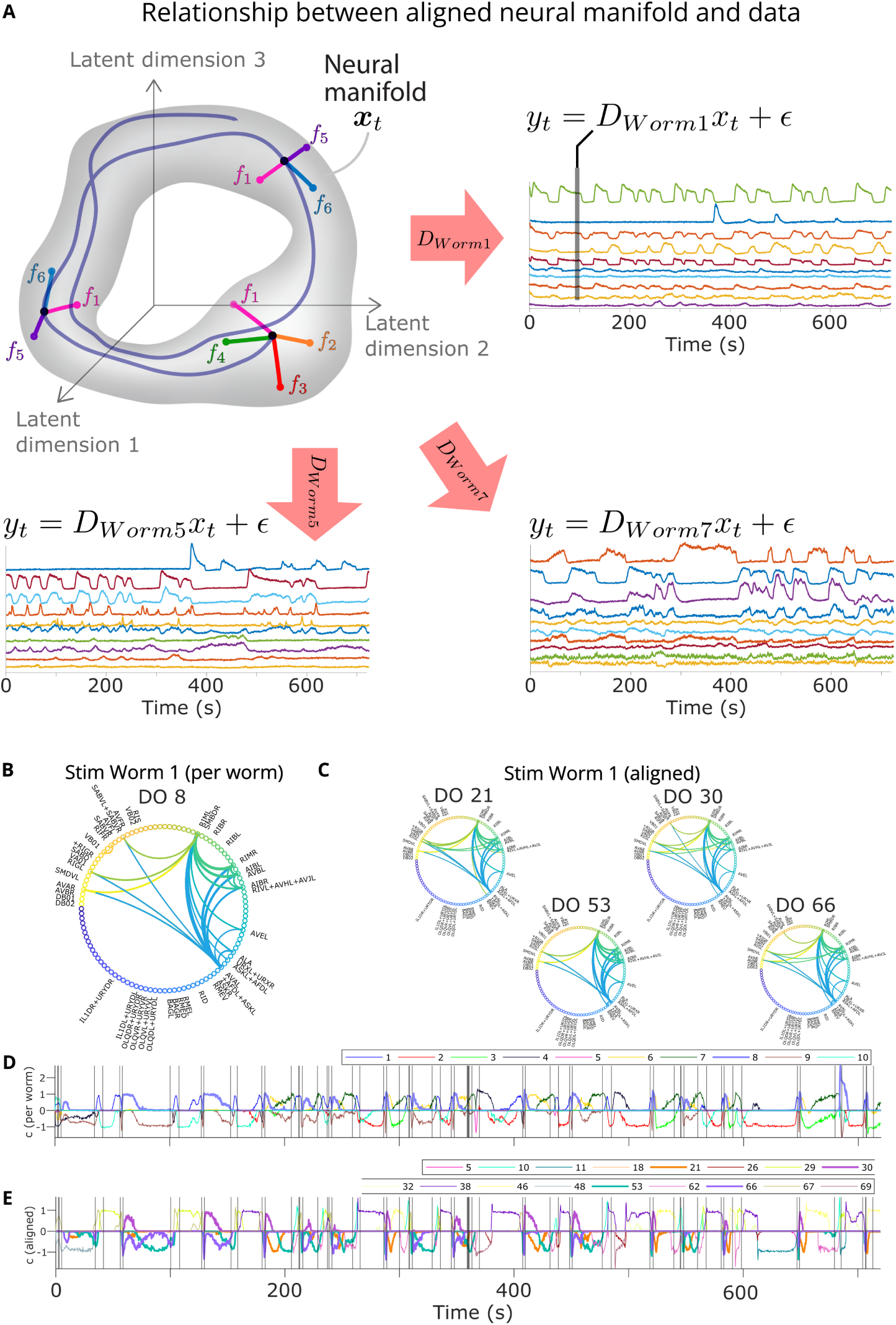
Aligned latent manifold and Stim Worm 1 example. **A:** We learn a shared set of DOs and latent dimensions, but a unique observation matrix and set of latent states and DO coefficients for each worm. **B,C:** We compare the connectivity maps for DO 8 in the per-worm model and the aligned DOs that co-occur with it. **D,E:** The DO coefficients themselves.

We found that, while the aligned models’ inferred DO coefficients were sparser than the total possible number of DOs, they were usually less sparse than the per-worm models (Supplementary Figs. 8, 9, 13); this was as expected, given that we did not scale up the sparsity hyperparameters for this larger model.

We also investigated whether each DO had a different “meaning” in terms of neuron-space dynamic connectivity in each worm, since each worm had a different projection out from the latent space (i.e., the per-worm representations ***D***). (e.g., DO 29 in Supplementary Figs. 38,39,40, 41, 42, 43, 44, and correlations between DOs vs. coefficients in Supplementary Figs. 45, 46). We thus tested whether the aligned DOs were washing out any structure, splitting up dynamic connectivity maps in space, or subdividing the activity of the per-worm operators in time, reflecting nonstationarity in a finer-grained way. By tallying features of the aligned DO dynamic connectivity maps (e.g., BAG and RMED participation), we observed that the aligned model operators shared key dynamic connectivity map features with the per-worm models. Therefore, even though the aligned model DOs used the same directions on the shared manifold to generate different connectivity maps, the aligned DOs overall for each worm still captured important and similar trends in which neurons were coactive and the relationships between them through time (e.g., Supplementary Figs. 38,39,40, 41, 42, 43, 44). We also observed smooth transitions in the middle of behavioral states in the aligned models, reminiscent of those in the per-worm models. This led us to reject the hypothesis that the aligned model with 70 DOs was washing out structure entirely.

Then, for each pair of per-worm and aligned DO coefficient traces, we calculated the probability of the intersection of their activity divided by the activity of one of the two DOs (e.g., Stim Worm 1, Supplementary Fig. 47). We saw a marked asymmetry: each aligned DO represented a small percentage of a per-worm DO utilization probability, but whenever an aligned DO was active, it was strongly coactive with at least one per-worm DO, and sometimes multiple. This indicated that the aligned DOs split up the per-worm DOs utilization patterns in time, as well as subdividing and reusing in multiple ways the dynamic connectivity maps associated with the per-worm DOs. In conclusion, when aligning dynamics across individuals using dLDS, model size can have an impact on how key dynamics are split up and represented, but the key dynamics will still be decipherable and will still represent key latent neural processes such that shared activity patterns across individuals are preserved.

## 3. Discussion, Limitations, and Future Work

In this work, we study the time-varying and contextual nature of *C. elegans* neural function. We focused on the novel characterization afforded by recent advances in time-varying dyanmical systems estimation—decomposed linear dynamical systems (dLDS)—to identify instantaneous “dynamic connectivity” that was able to track changing functional connections across behaviors. The decomposed nature of (dLDS) was able to uncover simultaneous neural processes, the dynamic operators (DOs) where each DO represents one fundament set of interconnections that ebb, flow, and combine to represent the complex nature of moment-by-moment neural dynamics across the full imaged population. We found that the DOs strongly corresponded to either specific motor or sensory tasks, and recombined in different ways across trials and across individuals to capture a diverse set of nonstationary interactions.

A key contribution is the combination of the instantaneous nature of our modeling framework with an interpretable locally linear dynamics model. These properties enabled us to define the dynamic connectivity, i.e., the moment-by-moment directional interactions between the full set of neurons. By interpreting directional dynamics in the latent state via their effects in the natural neural space, we reveal evidence that changes in instantaneous, dynamic connectivity can be the mechanism for sensory neuron adaptation and interneuron function across multiple circuits and task variables.

In general, understanding context-dependent functions or nonlinear relationships between neurons has been a challenging endeavor in systems neuroscience. Specifically, context-dependent relationships can overlay and interact with context-independent operators, interacting in ways that would be impossible to unravel in current RNNs or switched dynamical systems. We demonstrate that exploring the neural interactions through the dLDS dynamic connectivity maps across behavioral states or conditions can reveal these context-dependent functions or nonlinear relationships. Using the dLDS-derived dynamic connectivity, we identified many novel connections not mapped by direct anatomical synapses or one-to-one functional connections, emphasizing the possibility that neurons also communicate indirectly and nonlinearly with other neurons (e.g., as AND, OR, XOR gates, etc.). Moreover, we developed an extension to dLDS that allows latent neural processes to be aligned across individuals. This model extension enables greater generalizability of the learned models.

Our analysis’ unique perspective results in the following findings:

- **Neurons can have dynamic functional roles in circuits, rather than a single or stable function or set of functions:** All of the neurons highlighted in this paper (sensory neurons BAGL and BAGR, interneurons AVFR, RIGL, RIS, interneuron/motor neuron SABD, and motor neuron RMED) are active and connected in both a behavioral and environmental context-dependent way.
- **Changes in interneuron connectivity mediate efficient task-switching:** Some interneurons (e.g., AVFR) may rapidly change their connectivity in different regimes based on the task-related sensory vs. motor information they receive, without obvious changes to their activity levels. While this looks like mixed selectivity when time-averaged, instantaneous dynamic connectivity shows that they act as relay stations that may rapidly switch connectivity regimes, instead. Thus, these highly connected neurons can efficiently reuse their connections to relay information from different circuits. In general, principles of efficient interneuron connectivity and task-selective function are an important question because interneurons are highly connected and they only constitute 27% of the *C. elegans* nervous system [47] and only 20-30% of primate cortical neurons [48].
- **Changes in sensory neuron connectivity show a mechanism of adaptation:** Sensory neurons (e.g., BAG) may change their dynamic connectivity to adapt to stimuli over time, first signaling broadly to many neurons, and then signaling more to a small number of connected interneurons, motor neurons and proprioceptive neurons. Moreover, using the connectivity strengths at each time point, we can quantitatively measure the connectivity shift from external sensing to proprioception as the worm adapts to the experimental protocol.
- **Worm individuality must be modeled to understand adaptation:** Despite the many features shared by these worms, their similar responses to stimuli, and the similar reconstruction performance achieved by the per-worm and aligned dLDS models, the data were nevertheless not identical, and no aligned model used the dynamic connectivity in the same way. We posit that these individual differences are *not* noise; they are essential to understand worm adaptation to their environments. Worm individuality and adaptation (or even learning) is supported by prior work [4; 6; 12; 13; 14; 30; 31; 32; 33; 34].

These findings based on the Kato 2015 data [3] can also lead to new avenues for experimentation. Contextual time-varying neural connectivity patterns across individuals during adaptation can be further investigated across species with perturbation experiments, e.g., using a mixture of new protocols and optical stimulation.

Taken together, our results show that dLDS can be used to reveal a detailed story of lower-dimensional latent neural processes such as sensation that are hierarchically overlaid on those latent neural processes regulating motor behavior states, and the complex roles of interneurons in between. We can use dLDS to gain insight into internal cognitive processes that are otherwise difficult to disentangle. Our analysis complements prior and concurrent work to show, in a data-driven way without time-averaging across behavioral or experimental conditions, that distributed coding in *C. elegans* instantaneously changes to create behavior-related neural activity, and that the connections effecting these changes adapt throughout a trial, even across outwardly similar behaviors.

While not explored in this study due to the confounding variable of behavior base rates potentially impacting nonstationarity in addition to environmental variables, it is possible that the amplitude of DO coefficients might reflect habituation or sensitization to a sensory stimulus, evidence accumulation, a change in wakefulness or attention, etc. Future attempts at applying dLDS to neural data to interpret quantitative differences in the DO coefficients would also benefit from detailed, quantitative measures of behavior and environment variables.

More generally, dLDS complements the space of current models by enabling both interpretability and expressivity. Switched LDS models can model nonstationary processes in terms of temporally local linear systems, but only one system can be active at a time. dLDS enables the flexibility to put together a combination of linear operators, which allows us to track continuously varying changes in the network even with single behavioral bouts. Moreover, it allows the modeling of simultaneous parallel processes, such as if an animal is monitoring for potential threats while also navigating its environment for food—or in our case turning while being puffed with oxygen.

## 4. Data and Code Availability

The code for both the discrete and the continuous models can be found at https://github.com/ dLDS-Decomposed-Linear-Dynamics. The code used to run the *C. elegans* data can be found at https://github.com/dLDS-Decomposed-Linear-Dynamics/dLDS-Discrete-Matlab-Model. The discrete code can also be pip-installed using the **dLDS-discrete** Python package, as described in https://pypi.org/project/dLDS-discrete-2022/.

The *C. elegans* calcium imaging data that support the findings of this study are available in the “Kato2015 whole brain imaging data” repository with the identifier http://dx.doi.org/10.17605/osf.io/2395t [3; 49].

The SSM Python package from the Linderman lab was used to run rSLDS [50].

The code for generating the circular dynamic connectivity graphs is available from the MATLAB File Exchange [51].

## Acknowledgments

EY and ASC were supported in part by NSF CAREER award 2340338 and NM was supported by a Kavli Fellowship. We would like to acknowledge Sue Ann Koay for the original version of the manifold illustration in Figure 1 and Francesco Randi for the advice on how to use the Leifer Lab repositories. We would further like to thank Andrew Gordus for his helpful review of our paper and literature recommendations, our reviewers for their guidance, and Olya Hakobyan, Esther Whang, and Matthew Creamer for their comments on our manuscript.

## 5 Methods

### 5.1 Decomposed Linear Dynamical Systems (dLDS)

In our framework [42], we rely on the neural manifold hypothesis, that is that neural activity comes from an underlying low-dimensional space that evolves characteristically in time via trajectories, which we call dynamics. We observe neural activity during behavior as neural time series traces ***y* ∈** R, which are modeled as a linear function of the latent neural states ***x*** via the observation matrix ***D***. (Note that here we define a neural latent “state” as a larger process in the brain that can occur simultaneously with other processes, as opposed to one whole-brain or whole-animal state at a time.) If trajectories in latent space are a useful model of neural processes, it is necessary to understand them as traversing a manifold. If we look at the tangent space at each latent state time point, we can break down the dynamics trajectories into a linear combination of operators G in continuous time. However, the matrix exponential becomes cumbersome to compute with even a small number of dimensions, so we use the discrete time approximation of this model to work with real neural data. dLDS uses an expectation maximization algorithm to alternate between jointly inferring the latent states ***x*** and the dynamics coefficients ***c***, and updating the randomly initialized dynamics dictionary ***F*** and observation matrix ***D*** via projected gradient descent. Inputs from the user are just the neural activity ***y*** and some parameters, including the hypothesized latent dimensionality and number of operators, as well as some regularization terms for sparsity and smoothness of the latent states and dynamics (Table 2). Sparsity is important because it helps with interpretability – it constrains the model to use a small number of linear operators to represent each time point.

1. Randomly initialize ***F*** and ***D***.
2. Infer ***x*** and ***c*** via LASSO (Equations 6 and 7):

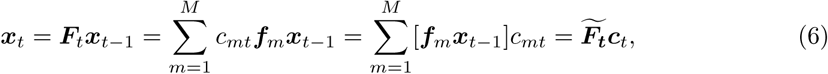

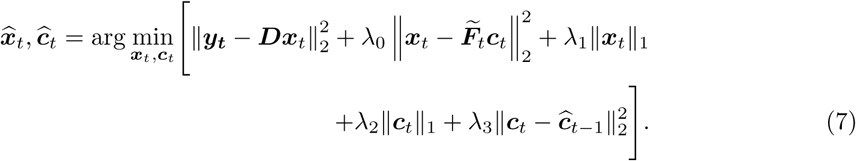

3. Update ***D*** and ***F*** via projected gradient descent (Equations 8, 9):

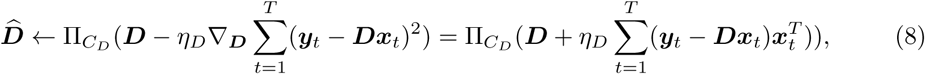

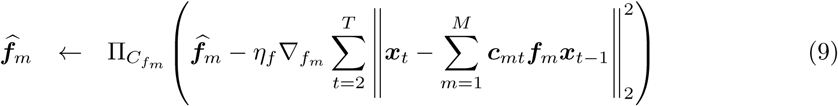

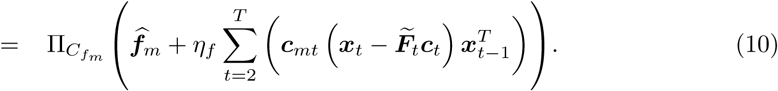

4. Repeat steps 2 and 3 until ***D*** and ***F*** each converge.

In the aligned model, we modify this procedure by inferring and storing ***x***,***c***, and ***D*** separately for each worm on each update iteration, whereas ***F*** is updated using a cell array containing the data from all of the worms, essentially concatenated as one trial after another.

We applied dLDS to each worm delta fluorescence trace individually on the CIS cluster. Preprocessing was minimal - negative values (up to 3% of each trace) were set to 0 because calcium-based imaging is a non-negative measure of activity. All values were scaled between 0 and 1 (maximum fluorescence value per worm) to promote even reconstruction performance.

We first ran 3000 iterations of dLDS and observed that after 500 iterations, all of the models achieved change in ***D*** and change in ***F*** on the order of 10*^−^*^3^. ***D*** and ***F*** were stored and used to perform inference one last time after dynamics learning to obtain the latent states (***x***) and dynamics coefficients (*c*). The number of latent dimensions and dynamics were initialized heuristically as 10 because there were 4-8 behavioral states labelled and because the median number of dynamics active at a time converged to 3-8 with a range of dimensionalities (6, 8, 10, 20), so an intermediate value (10) was chosen in favor of the smallest stable number of coefficients without artificially constraining the median number of active coefficients at any given time (Table 2). Similarly, a range of regularization parameters was explored (from near 0 to greater than 1) before settling on the fairly strong regularization for the smoothness of the states and coefficients. Since ***D*** and ***F*** are randomly initialized, multiple examples were generated for three of the worms and compared visually.

rSLDS was also run for 500 iterations on each worm with 10 latent dimensions and 4 output states for the Stim worms (or 8 for the NoStim worms).

For the reconstructions (Table 1), the product of D with the latent states was rescaled back to the original dF/F range (*R*^2^ between 0.7 and 0.9 for each of the worms). The dynamics coefficients were also compared in terms of their utilization during the experimental settings of the experiment (histograms) and during various behavioral states.

### 5.2 C. elegans data

*C. elegans* calcium imaging data was obtained from [49] (Table 3).

### 5.3 Validation: Decoding other variables from dLDS outputs

We validated the decoding of other additional variables, such as speed or oxygen concentration, from the DO coefficients or latent states. First, we trained an SVM to predict nonstationarity, i.e., whether individual time points were in the first or second half of the trial based on either 1) the DO coefficients (“c vals”) or 2) the latent states (“x vals”) at only that time point. These SVMs correctly classified at 85% mean accuracy for the DO coefficients and 87% mean accuracy for the latent states (Supplementary Figs. 4, 5). However, we saw slightly decreased mean accuracy (84% for the DO coefficients, 85% for the latent states) in the prediction of precise oxygen levels (detailed stimulation times in the Stim experiments only, rather than first half vs. second half), where 4% oxygen times were primarily misclassified as 21% oxygen (Supplementary Fig. 6). This is not entirely surprising, as it is not possible to change the oxygen concentration in a chamber uniformly and instantaneously, and the base rate of 21% oxygen exposure was higher in these experiments. Despite these confounds, the latent states and DO coefficients are clearly nonstationary between the two halves of all the Stim experiments and 4 out of the 5 NoStim experiments in a way that an SVM can decode (Supplementary Figs. 4, 5). Since nonstationarity was also observed in the NoStim worms, dLDS may be picking up on nonstationarity in the data not only due to the oxygen protocol, but also likely due to other time-dependent factors.

As far as classifying speed (forward crawling vs. forward slowing), SVMs trained on either the DO coefficients or the latent state values were also successful (mean accuracy: 83% for the DO coefficients, 82% for the latent states), but were mostly biased toward the forward slowing states (Supplementary Fig. 7). Overall, we found, qualitatively and quantitatively, that the DO coefficients do encode features of the latent neural states that reflect both the discrete and continuous nature of the behaviors they generate.

### 5.4 Computing dynamic connectivity

In this paper we feature dynamic connectivity maps based on neuron location in the *C. elegans* head, which also show time-averaged simulated activity of the neurons of interest. We also use circular dynamic connectivity graphs [51] that only show connections, without activity.

#### 5.4.1 Connectivity mapped to *C. elegans* head

Each labelled neural trace was matched to a three-dimensional location in the worm and a class, as labelled by [11]. The connections between neurons were calculated as a covariance matrix between traces (from rows to columns). Recall that in a latent dynamics model where ***y****_t_* = ***Dx****_t_*, ***D*** is the mapping from the latent space to the ambient (e.g., neural recording) space. In typical connectivity maps, the system (i.e., 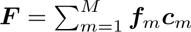) is considered stationary, but because of the time-varying DO coefficients ***c****_t_* in dLDS, the system can be modeled in a time-dependent way. Our dynamic connectivity maps can show not only the dynamic (i.e., time-varying) connectivity (via lines between neurons), but also the reconstructed time-averaged or instantaneous activity of those neurons (via neuron opacity) resulting from that particular linear combination of DOs. For any linear combination of DOs ***F***, the correlations in the ambient space of an equivalent linear time-invariant dynamical system are calculated as ***D*** ∗ ***F*** ∗ ***D****^T^*. To produce the time-averaged simulated fluorescence values ***x***^ generated only by a particular combination of DOs of interest, simulated latent states were calculated by applying an impulse function (vector of ones) to the combined DO ***F***, taking the dot product iteratively for 500 time points, and then taking the mean reconstructed neural activity for each neuron. To produce stable combined DOs, ***F*** was rescaled to unit norm (after linearly combining the component DOs, but before applying the impulse function). Then the simulated latent states were multiplied by ***D*** to produce the simulated fluorescence.

The dynamic connectivity maps were produced for only the labelled traces (less than 50% of the recorded traces). If multiple neurons were present in that trace, the activity of that trace was plotted redundantly in each of the locations associated with the corresponding component neurons. The simulated fluorescences plotted were the mean for each trace. Strong thresholding on both the fluorescence values and the connectivity was performed to declutter the plotting - i.e., only the strongest 0.1-0.5% of connections are shown. Thresholding was also performed on the automatic neuron trace labels so that the strongest activity and strongest connections were labelled.

#### 5.4.2 Circular connectivity graphs

The circular connectivity graphs [51] are another way of visualizing the dynamic connectivity, without spatial information or information about neuron activity. The lines go from source to target neuron (or neuron combination, or unlabelled trace). The color of each line denotes its source.

### 5.5 BAGL/R dynamic connectivity tally

For each behavior or experimental condition context analyzed, the corresponding active DO traces were visually selected for each worm model. Then a dynamic connectivity map was made for each trace, and the presence of key neurons was tallied across those maps, both in terms of their appearance in DOs across worms and whether they appeared in each worm model (Tables 4, 8).

**Table 8:** Tally of RMED presence in dynamic connectivity maps across conditions.

| subset | per-worm DOs | worms | aligned DOs | worms |
| --- | --- | --- | --- | --- |
| Stim FWD | 22/31 | 6/7 | 35/46 | 7/7 |
| Stim REVSUS | 4/36 | 3/7 | 8/54 | 3/7 |
| Stim only 1st half | 0/3 | 0/7 | 1/6 | 1/7 |
| Stim mostly 1st half | 3/11 | 2/7 | 4/8 | 2/7 |
| Stim mostly 2nd half | 4/8 | 2/7 | 6/9 | 3/7 |
| Stim only 2nd half | 8/9 | 4/7 | 12/20 | 7/7 |
| Stim sign flip between halves | 2/3 | 2/7 | 0/1 | 0/7 |
| NoStim FWD | 17/21 | 5/5 |  |  |
| NoStim SLOW | 20/27 | 5/5 |  |  |
| NoStim REVSUS | 3/23 | 3/5 |  |  |

Table 4 is a tally of strong connectivity with or activity in BAGL and/or BAGR across the worms’ dynamic connectivity maps for each DO.

In the per-worm models, BAGL and BAGR were strongly present in DOs that were only active in the second half of the trials (alternative oxygen levels) in more than half of the worms and in DOs that were mostly active in the second half in more than a third of the worms (Supplementary Figs. 16, 19, Table 4). Furthermore, they were present in *none* of the operators that were exclusively or mostly in the first half (Supplementary Figs. 21, 24).

We also observed that BAGL and BAGR were not evenly used between behavioral states; DOs featuring BAGL/R utilized during forward crawling were present in all 7 Stim worms (Fig. 27), but only during reverse sustained crawling in 2 out of 7 worms (Fig. 30). This brings up the question of whether BAGL/R activity and connectivity is dependent on behavioral context. This bias of BAG-featuring DOs toward forward crawling could be a result of the base rates of those two behavioral states, which were biased toward forward crawling in the second half of the experiments, indicating that BAGL/R utilization is independent of behavior and is simply more present during forward crawling because forward crawling occurred more often during hypoxia. However, the bias of BAG-featuring DOs toward forward crawling states could indicate context-dependent BAG utilization. This also called into question whether nonstationarity generally was a side effect of when forward crawling was occurring.

### 5.6 *χ*^2^ tests

In order to test the independence of individual neuron activity and connectivity levels (on/off) from categorical variables such as oxygen stimulation settings and behavior states, we conducted *χ*^2^ tests [52] for each Stim worm per-worm model (Table 5). The null hypothesis of the *χ*^2^ test is that there is no difference in expected value of the observed variable (in our case, reconstructed activity or connectivity) between groups (behavior states, stimulation states - in the stim half of the trial or not, or a detailed readout of either of those categories). Rejecting the null hypothesis reflects a significant difference in observed vs. expected values, indicating that that category significantly impacts the observed values. Most tests were found to be significant, even when the alpha value for rejection was set at an extremely conservative 9e-5, based on a Bonferroni correction from 0.05 by the number of tests per worm.

First, we compared behavior states to stimulation levels and found that the stimulation times significantly impacted the behavior states in most of the worms, which validates the idea that the stimulation protocol could be used to entrain worm behavior (see “bhv” and “FWD” vs. “stimTimes” and “stimTimesDetailed” in Table 5). For most of the 7 Stim worms, in terms of reconstructed activity (*y*^), at a weak threshold for “active” (above median value), both BAGL and BAGR were influenced by state 3 (REVSUS), but at a strong threshold (median plus 3 standard deviations), BAGL was most influenced by state 1 (FWD). At both levels, they were most active during the second half of the experiments (alternating oxygen stimulation protocol (“Stim”= 1)). The stimulation protocol results and the strong threshold results align with the visual observations from the connectivity maps above, which only showed the strongest activity levels and connections. As far as the connections to and from BAGR and BAGL, the *χ*^2^ test results were inconclusive as far as whether they were dependent on stimulation protocol state at the lower threshold for “connected”, but at the higher threshold, connectivity to and from BAGR and BAGR in most worms significantly interacted with both the stimulation protocol states and behavior states.

### 5.7 ANOVA and MANOVA tests

In order to test effects of categorical factors such as oxygen stimulation settings and behavior states on individual neuron activity and connectivity, we conducted ANOVA and MANOVA tests (with Roy’s Largest Root, which most consistently returned non-NaN or Inf values, and which only uses the variance from the dimension of greatest separation between the groups to create the test statistic) [53; 54] for each Stim worm per-worm model (p-values in Tables 6, 7). All of the tests with connectivity to and from each neuron were found to be significant for Behavior (1-4), Stimulation (0 or 1, i.e., 21% oxygen first half vs. alternating oxygen levels second half), and interaction between the levels of Behavior and Stimulation. Most of the activity tests were also significant to varying degrees (*α* = 1 ∗ 10*^−^*^4^ to 0.05, depending on the correction for multiple tests).

All of the neurons, including BAGL and BAGR, showed significant effects of Behavior and Stimulation levels on the connectivity to and from across all 7 Stim worms. As far as reconstructed neural activity, BAGL showed consistently significant effects of both the behavior levels, stimulation protocol, and the interaction between the two, with the exception of Worm 2’s BAGR activity, despite affecting the connectivity.

### 5.8 RMED dynamic connectivity tally

Table 8 is a tally of strong connectivity with or activity in RMED across the worms’ dynamic connectivity maps for each DO.

RMED was utilized in 22/31 forward crawling DOs and during forward crawling in 6/7 Stim worms, but only in 4/36 reverse sustained crawling DOs and during reverse sustained crawling in 3/7 worms (Supplementary Figs. 27, 30, Table 8). RMED-featuring DOs were not at all present in the small number of DOs exclusively active in the first half of the experiments and were present in DOs exclusively active in the second half in 4/7 worms, but otherwise they were utilized in a more balanced way between halves of the experiment (Supplementary Figs. 16, 19, 21, 24). Notably, this relationship between RMED and forward crawling was even stronger in the NoStim worms, which were not exposed to hypoxia. It would seem that RMED should have shown the opposite bias (toward the first half of the Stim experiment) if its nonstationarity were being affected by the oxygenation levels; this observed bias toward the second half of the experiment could be a result of the corresponding bias in the base rates of the forward crawling state(s).

## Footnotes

1 We use the term “aligned” in this paper to mean that the latent states are inferred across individuals such that they have shared relationships with each other, i.e., a shared set of dynamics operators. We do not refer to “alignment” in the machine learning sense of alignment to human values [43].

2 There is some bias due to the base rates of forward and reverse crawling vs. relatively short bouts of turning.

