## Supplementary Figures for "Decomposed Linear Dynamical Systems (dLDS) models reveal instantaneous, context-dependent dynamic connectivity in *C. elegans*"

### 1. Supplementary Information

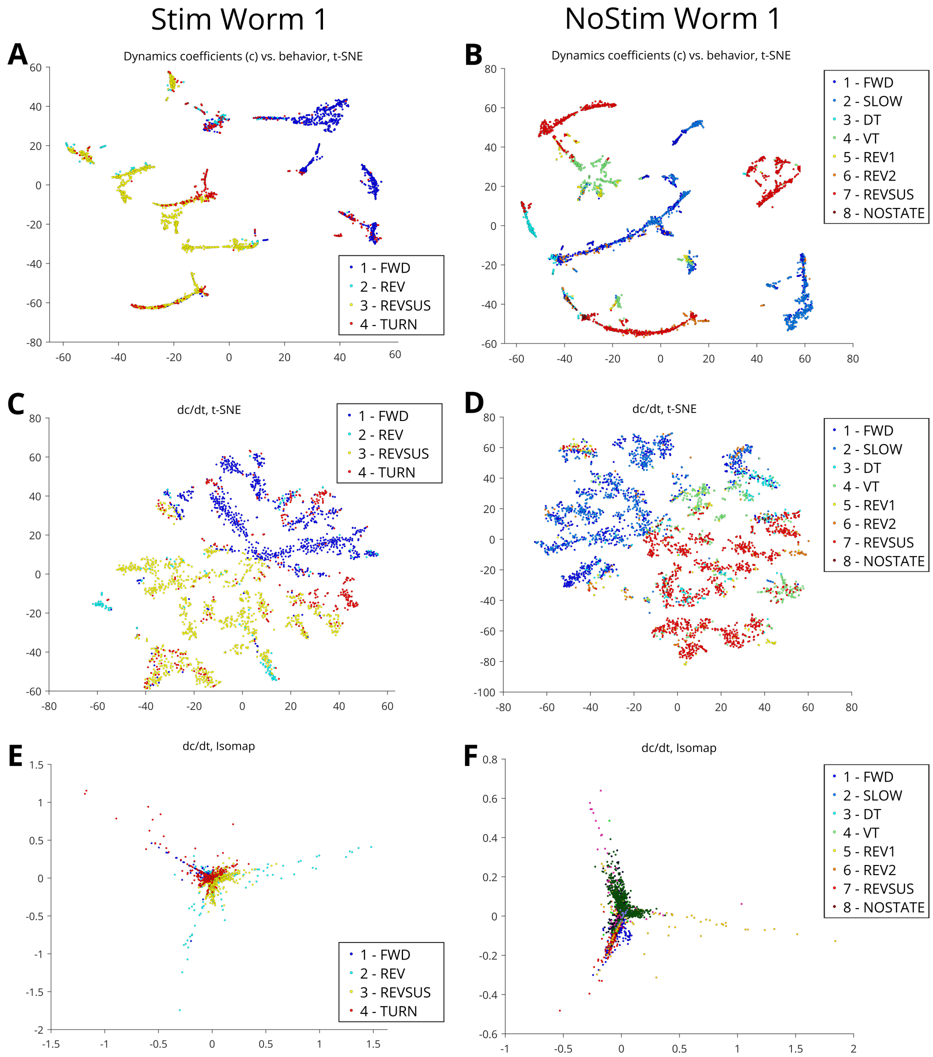


Supplementary Figure 1: **Clustering of DO coefficients ($c$) and first derivative ($\frac{dc}{dt}$) by behavior labels. Left:** Stim Worm 1. **Right:** NoStim Worm 1. **A,B:** DO coefficients vs. behavior, t-SNE. **C,D:** First derivative vs. behavior, t-SNE. **E,F:** First derivative vs. behavior, Isomap. Each dot represents a time point in DO coefficient trace; see Section 6.1 and Table 3 for model parameter settings and counts. By clustering and coloring by behavioral states, t-SNE appears to show a continuum from forward to backward crawling (DO coefficients first derivative $\frac{dc}{dt}$). Isomap embedding reveals a spoke-like structure in the first derivative.

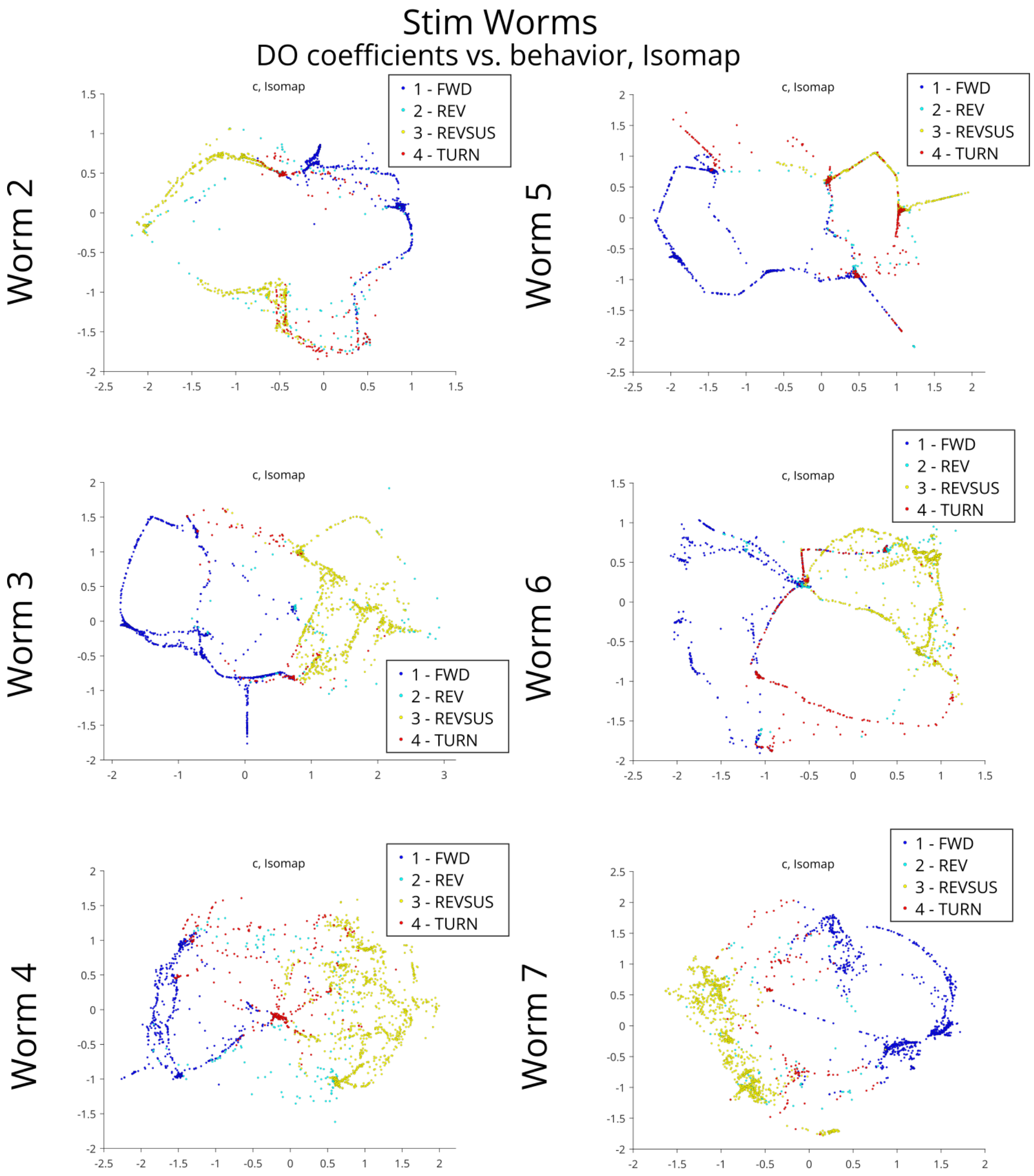


Supplementary Figure 2: **Clustering of DO coefficients ($c$) by behavior labels, Isomap only, remaining Stim Worms.** Each dot represents a time point in DO coefficient trace; see Section 6.1 and Table 3 for model parameter settings and counts. Isomap embedding reveals a ring-like structure from state to state (with additional loops) in the coefficients and a spoke-like structure in the first derivative.

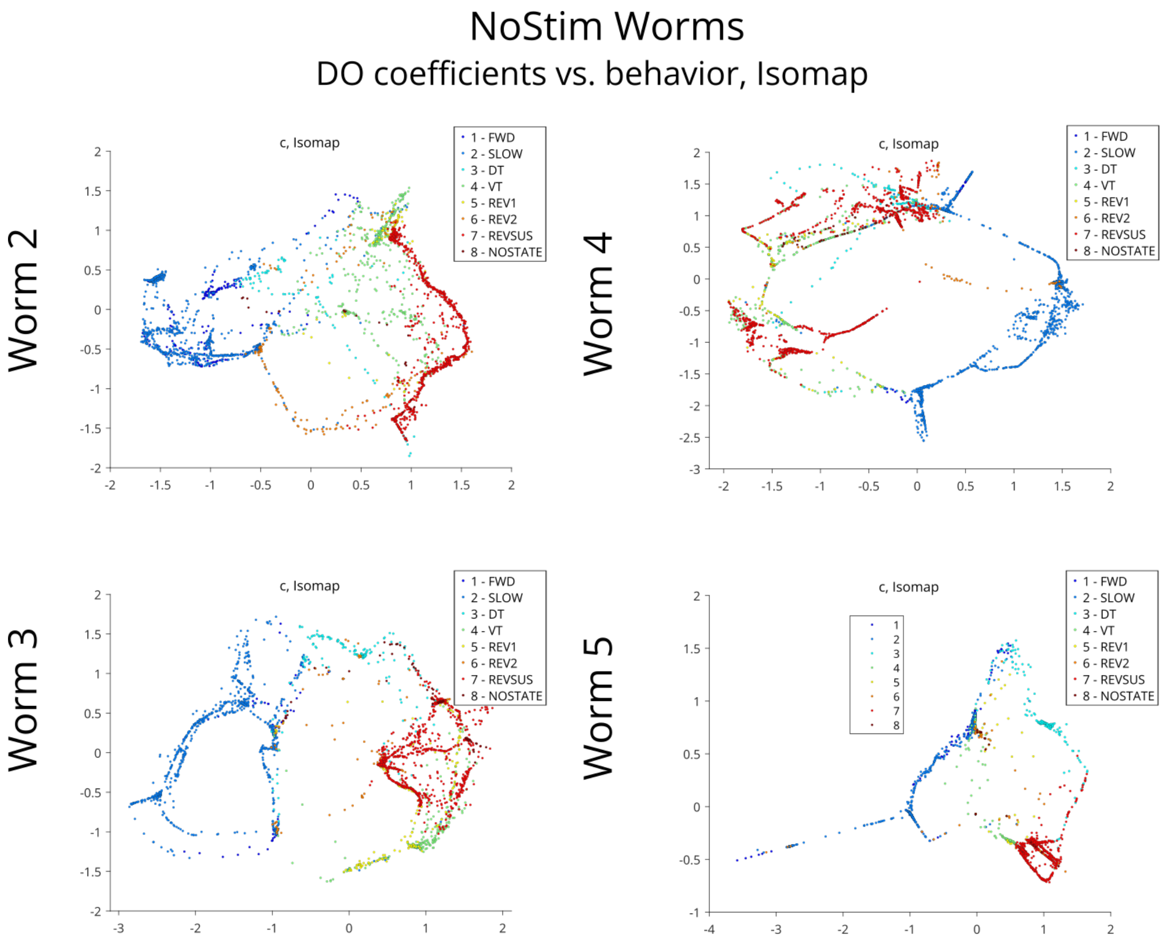


Supplementary Figure 3: **Clustering of DO coefficients ($c$) by behavior labels, Isomap only, remaining NoStim Worms.** Each dot represents a time point in DO coefficient trace; see Section 6.1 and Table 3 for model parameter settings and counts.

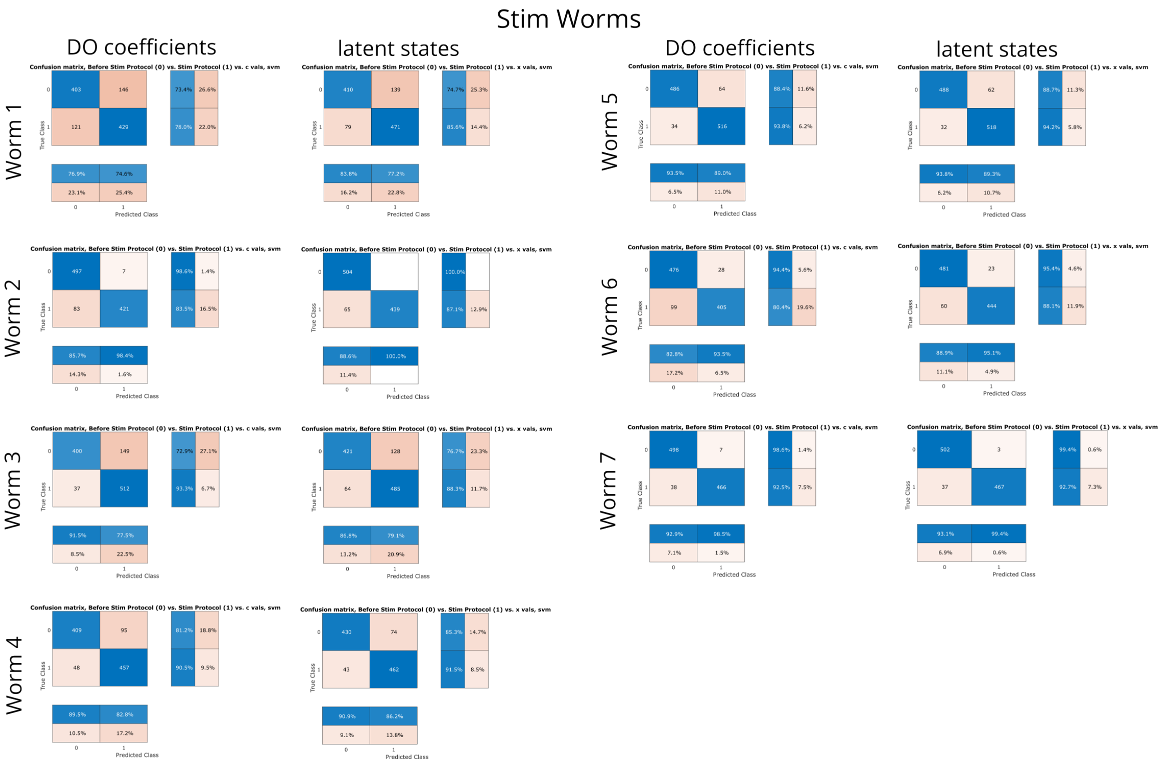


Supplementary Figure 4: **Confusion matrices from SVMs, decoding nonstationarity (first half vs. second half of trials) from dynamics coefficients $c$ and latent states $x$, Stim worms.**

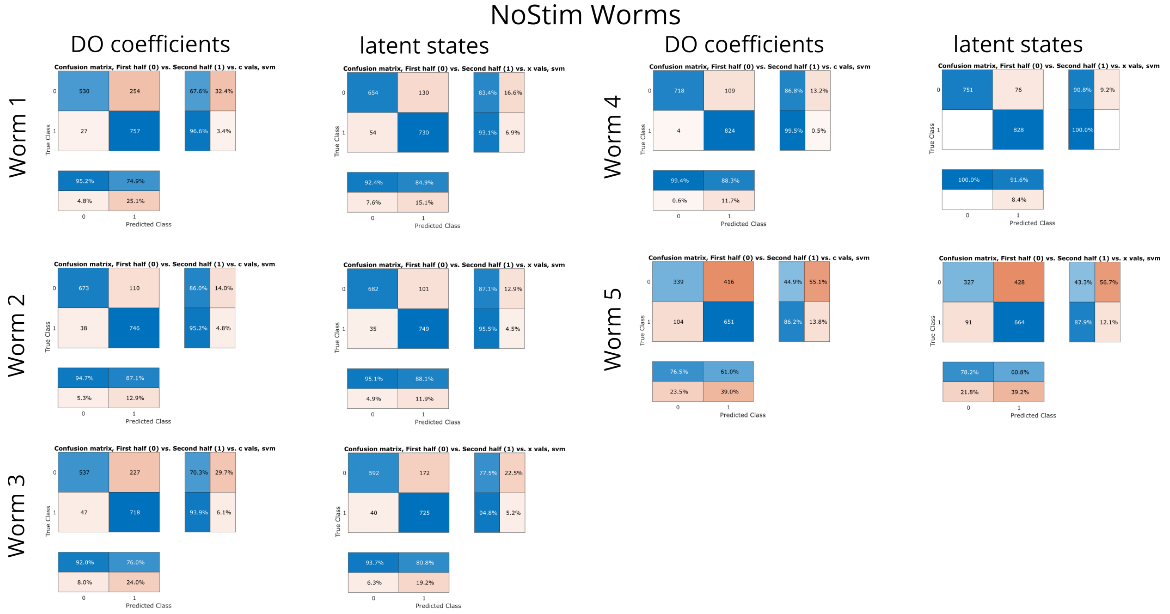


Supplementary Figure 5: **Confusion matrices from SVMs, decoding nonstationarity (first half vs. second half of trials) from dynamics coefficients $c$ and latent states $x$, NoStim worms.**

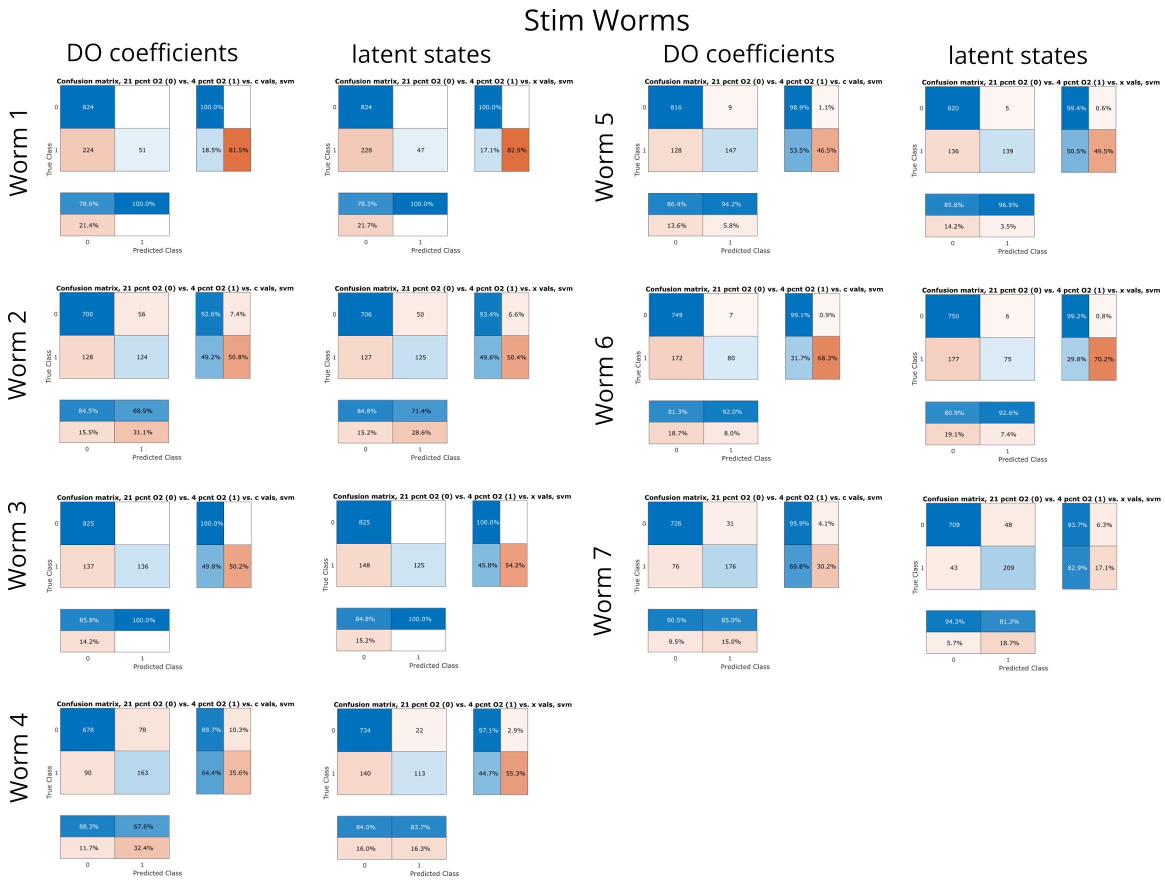


Supplementary Figure 6: **Confusion matrices from SVMs, decoding oxygen concentration (21% vs. 4%) from dynamics coefficients, Stim worms only.**

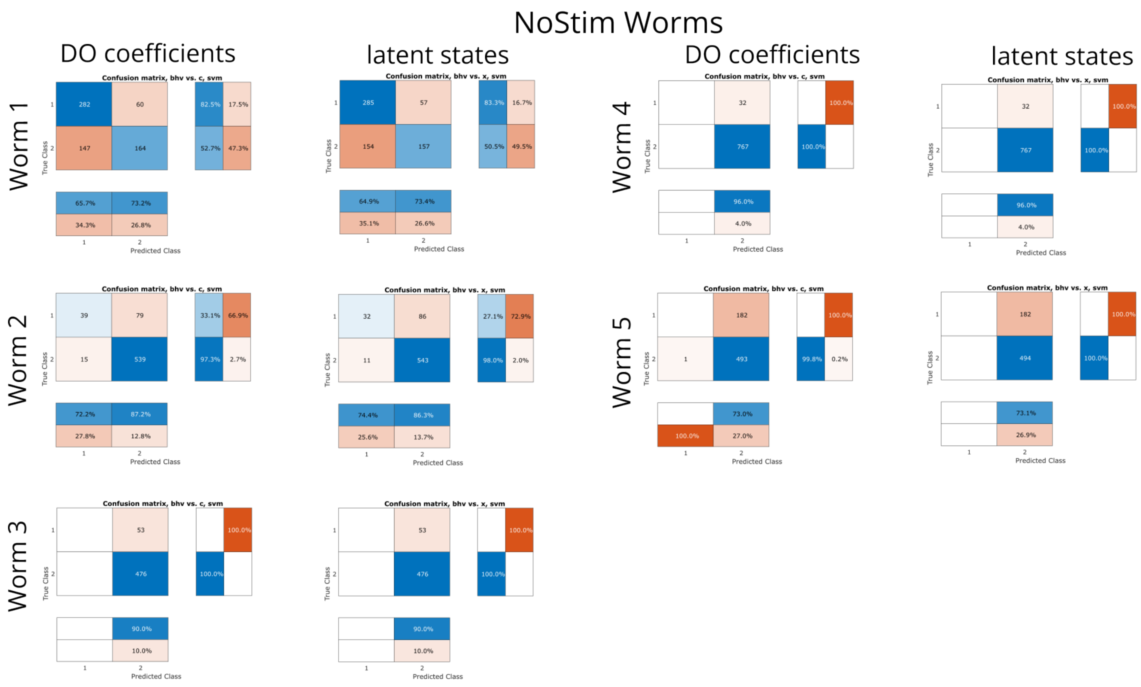


Supplementary Figure 7: **Confusion matrices from SVMs, decoding speed (Forward Crawling vs. Forward Slowing) from dynamics coefficients, NoStim worms only.**

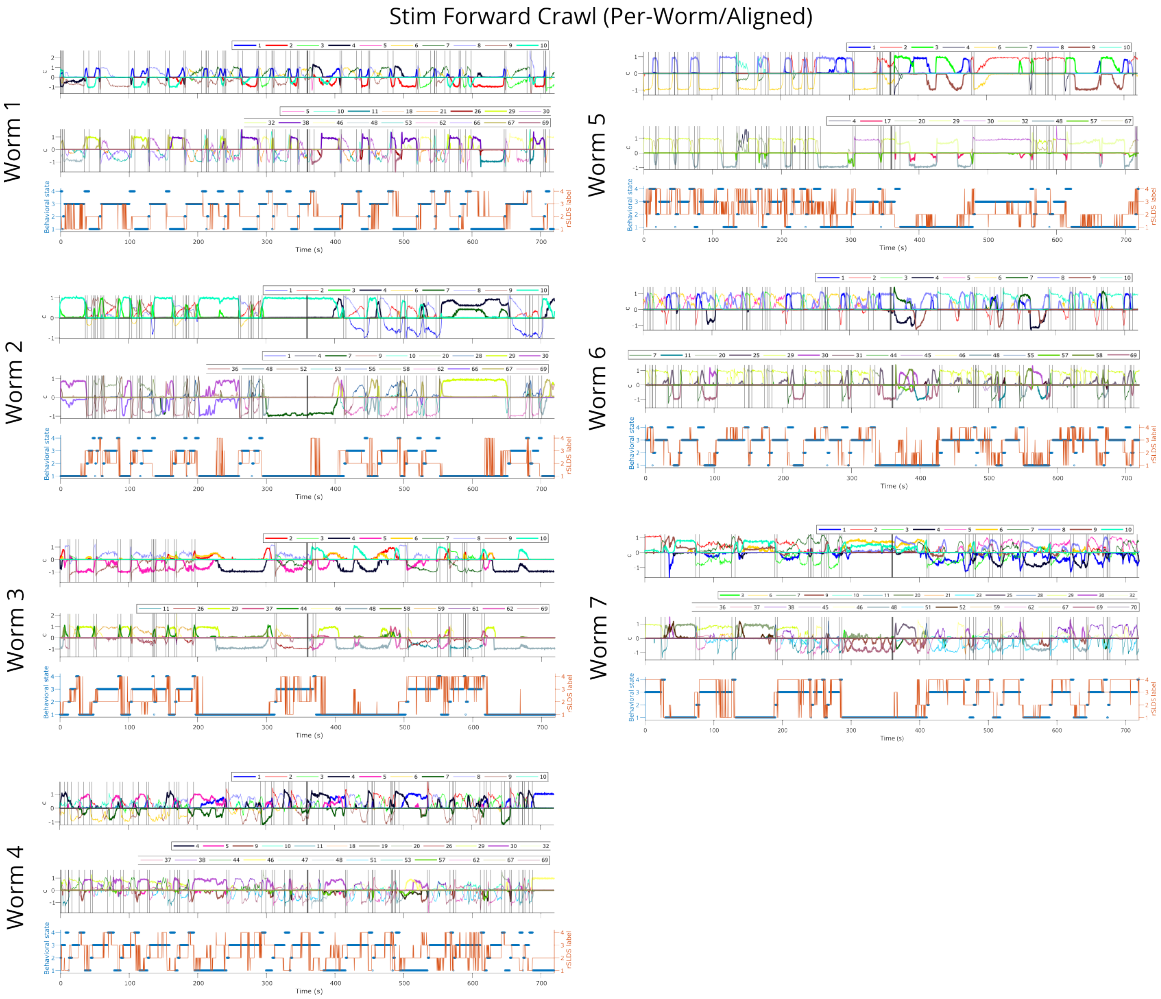


Supplementary Figure 8: **Stim worm dynamics comparison summary during behavior 1 - Forward Crawling.** Dynamics coefficients (**top:** per-worm, **second:** aligned), rSLDS labels (orange) and behavior labels (blue) (**bottom**).

Stim Reverse Sustained Crawl (Per-Worm/Aligned)

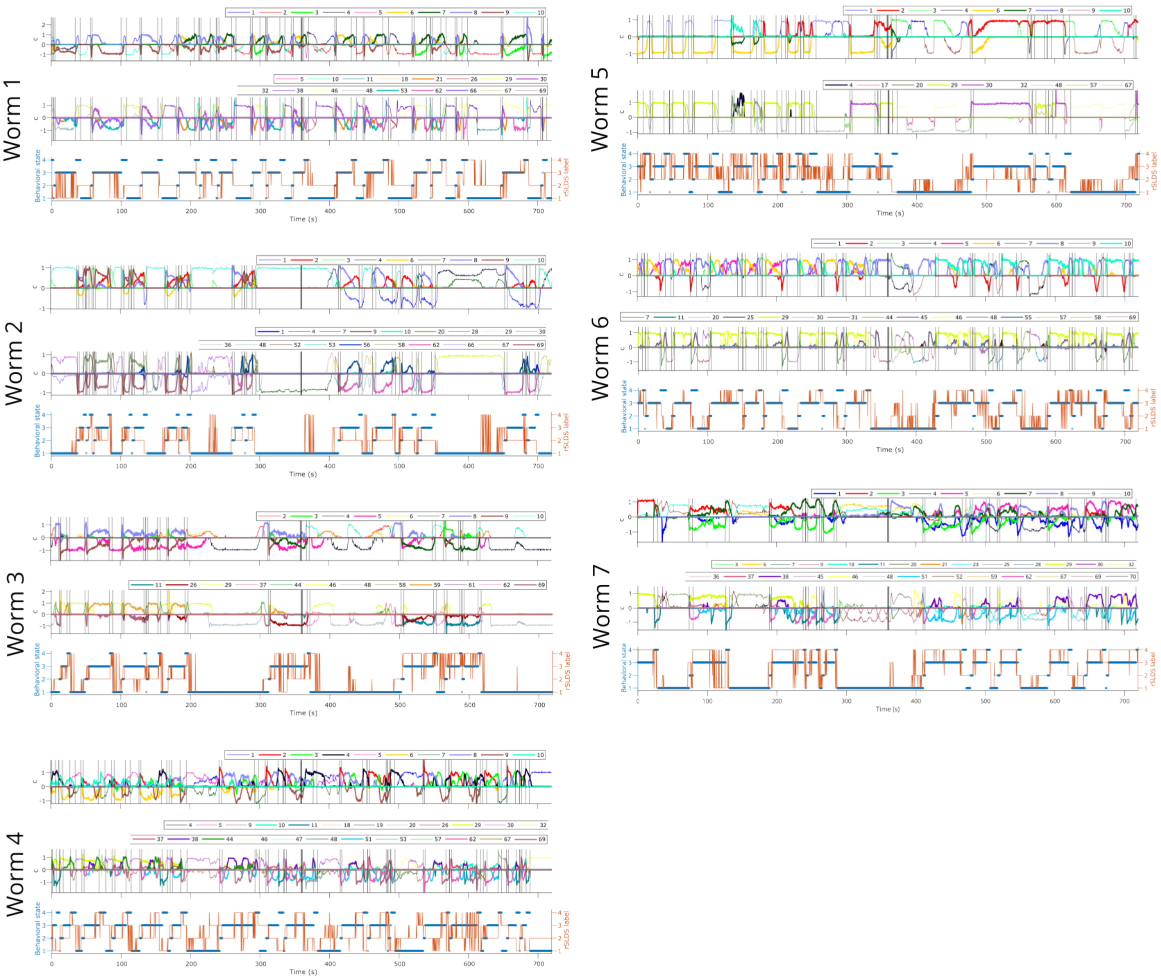


Supplementary Figure 9: **Stim worm dynamics comparison summary during behavior 3 - Reverse Sustained Crawling.** Dynamics coefficients (**top:** per-worm, **second:** aligned), rSLDS labels (orange) and behavior labels (blue) (**bottom**).

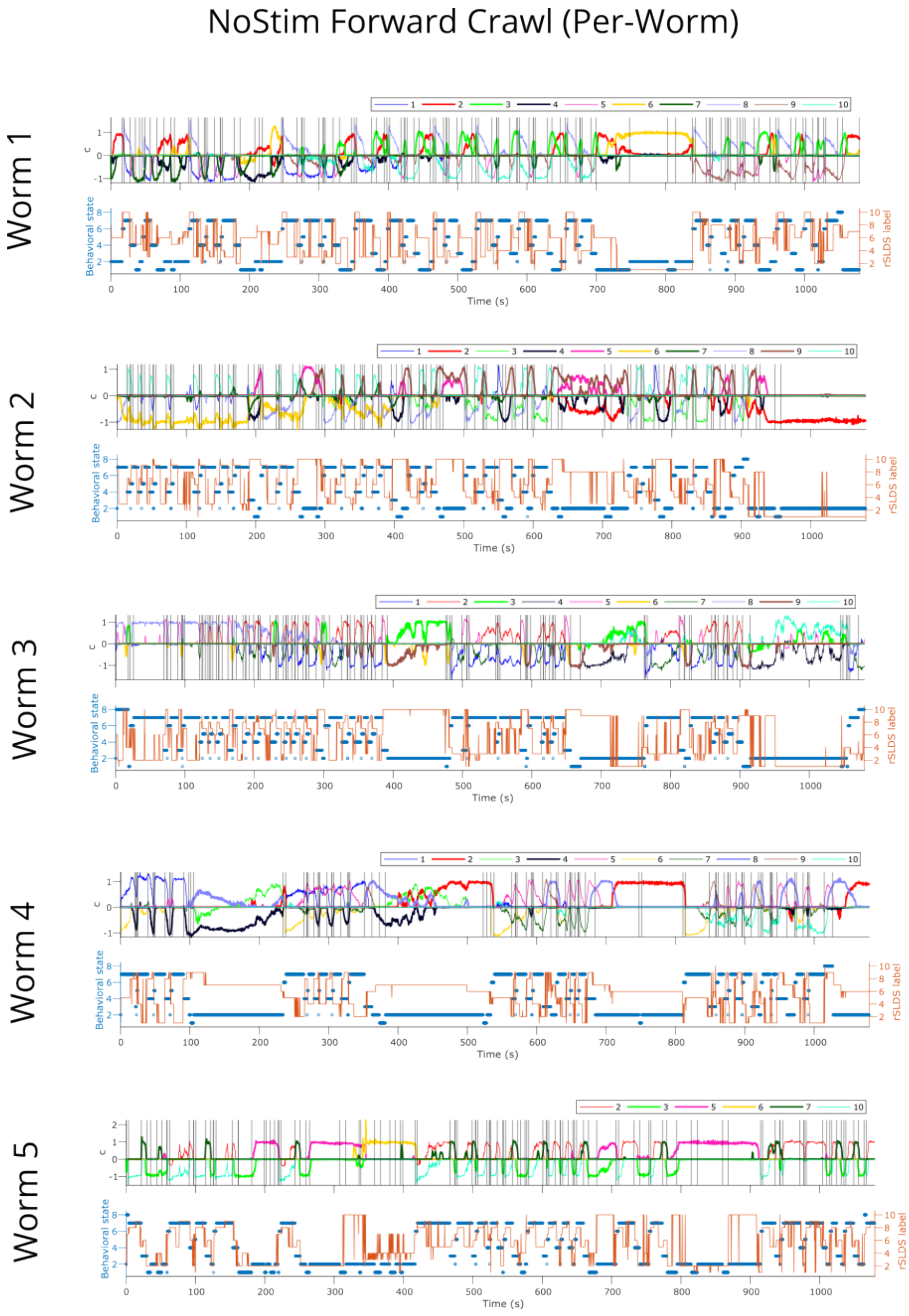


Supplementary Figure 10: **NoStim worm dynamics comparison summary during behavior 1 - Forward Crawling.** Dynamics coefficients (**top**), rSLDS labels (orange) and behavior labels (blue) (**bottom**).

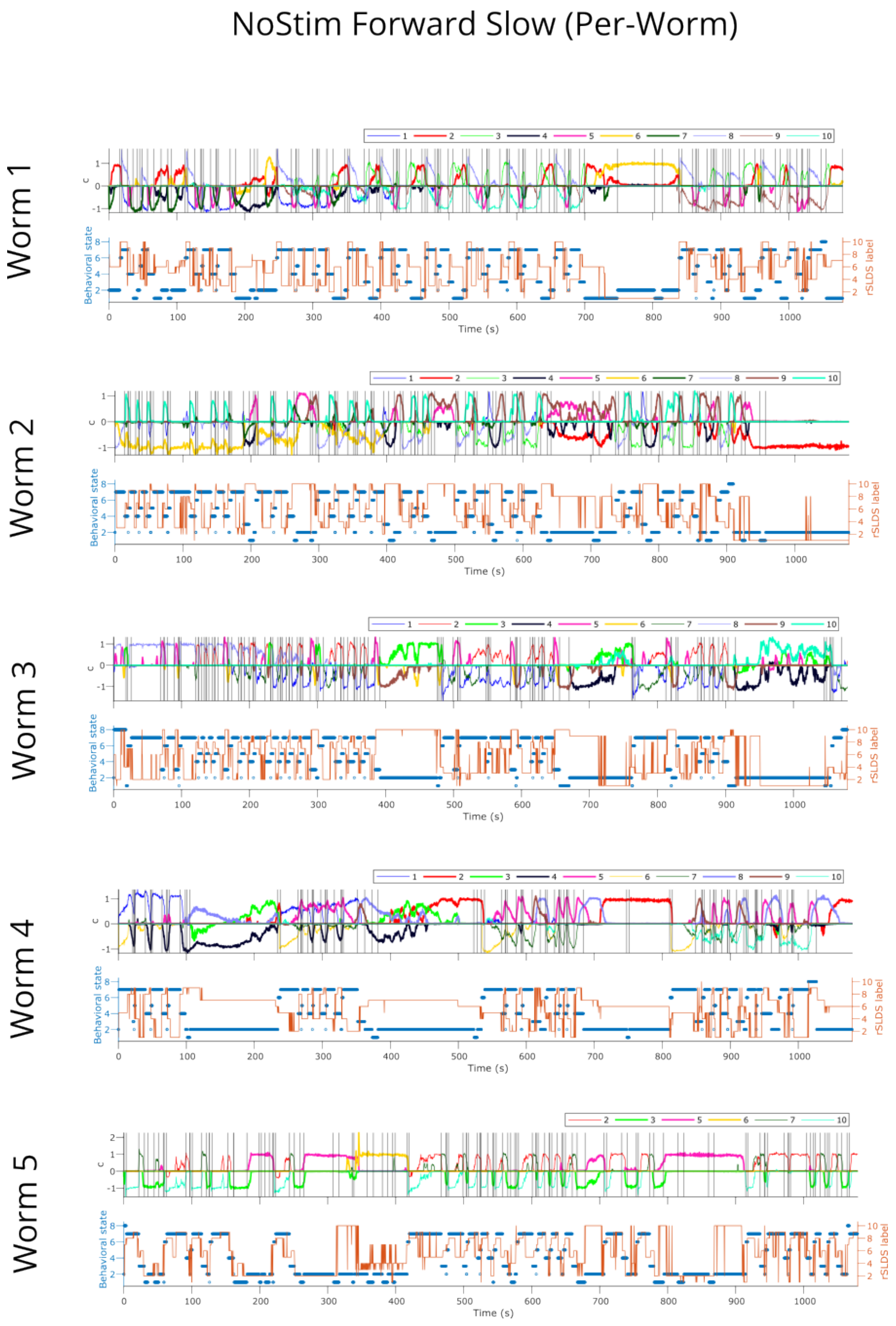


Supplementary Figure 11: **NoStim worm dynamics comparison summary during behavior 2 - Forward Slowing.** Dynamics coefficients (**top**), rSLDS labels (orange) and behavior labels (blue) (**bottom**).

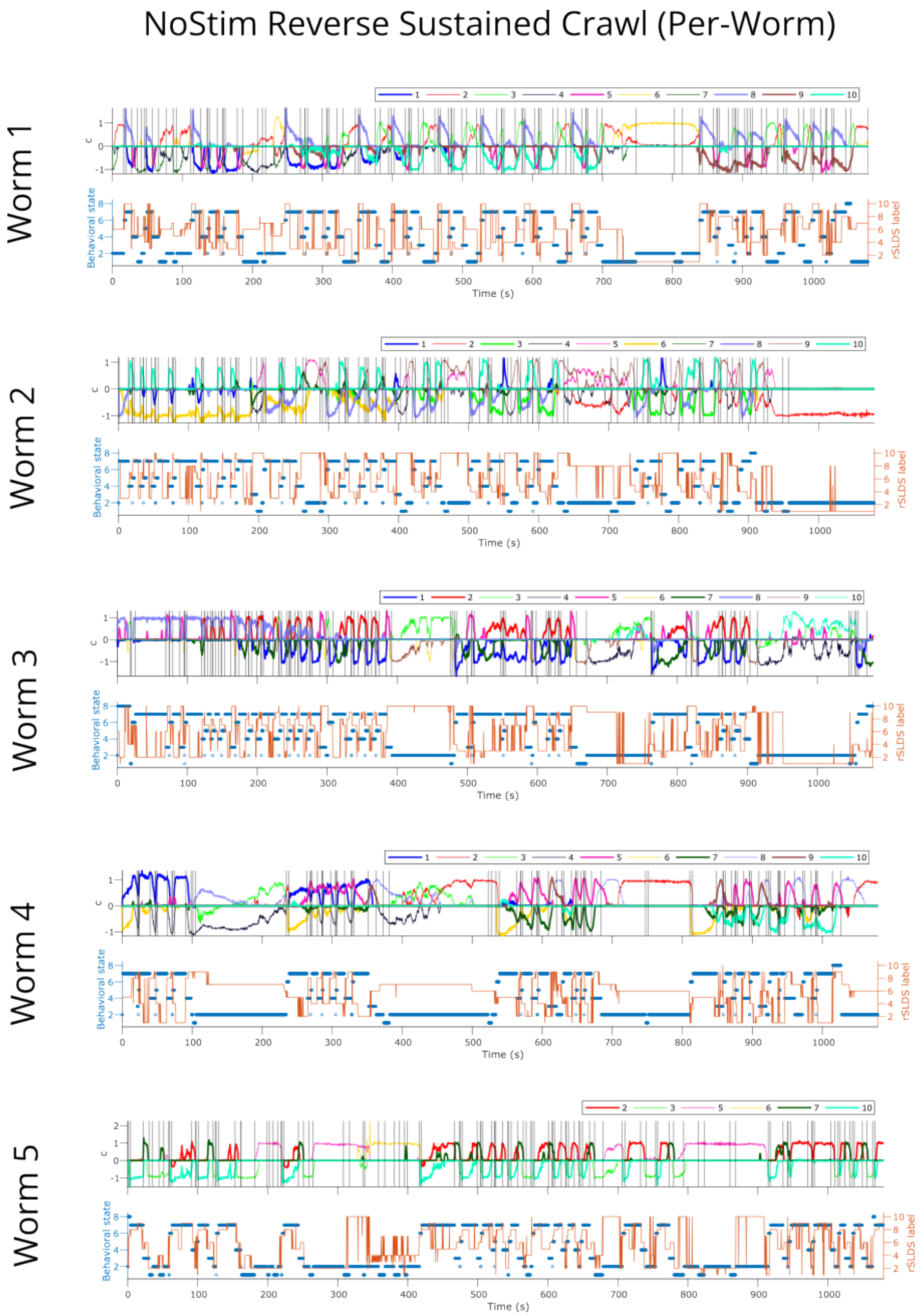


Supplementary Figure 12: **NoStim worm dynamics comparison summary during behavior 3 - Reverse Sustained Crawling.** Dynamics coefficients (**top**), rSLDS labels (orange) and behavior labels (blue) (**bottom**).

Oxygen Sensitivity or Nonstationarity (Per-Worm/Aligned)

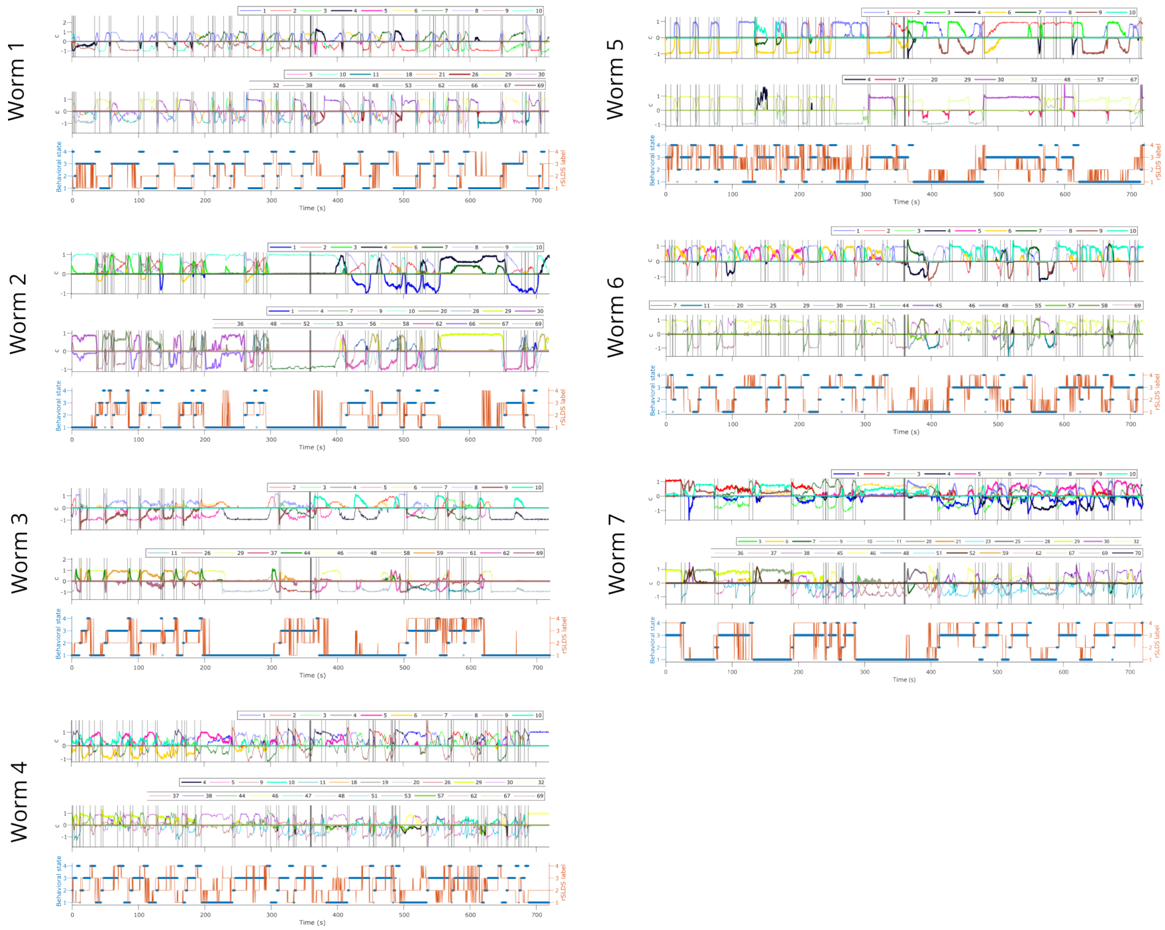


Supplementary Figure 13: **Stim worm dynamics comparison summary highlighting non-stationarity.** Dynamics coefficients (**top:** per-worm, **second:** aligned), rSLDS labels (orange) and behavior labels (blue) (**bottom**).

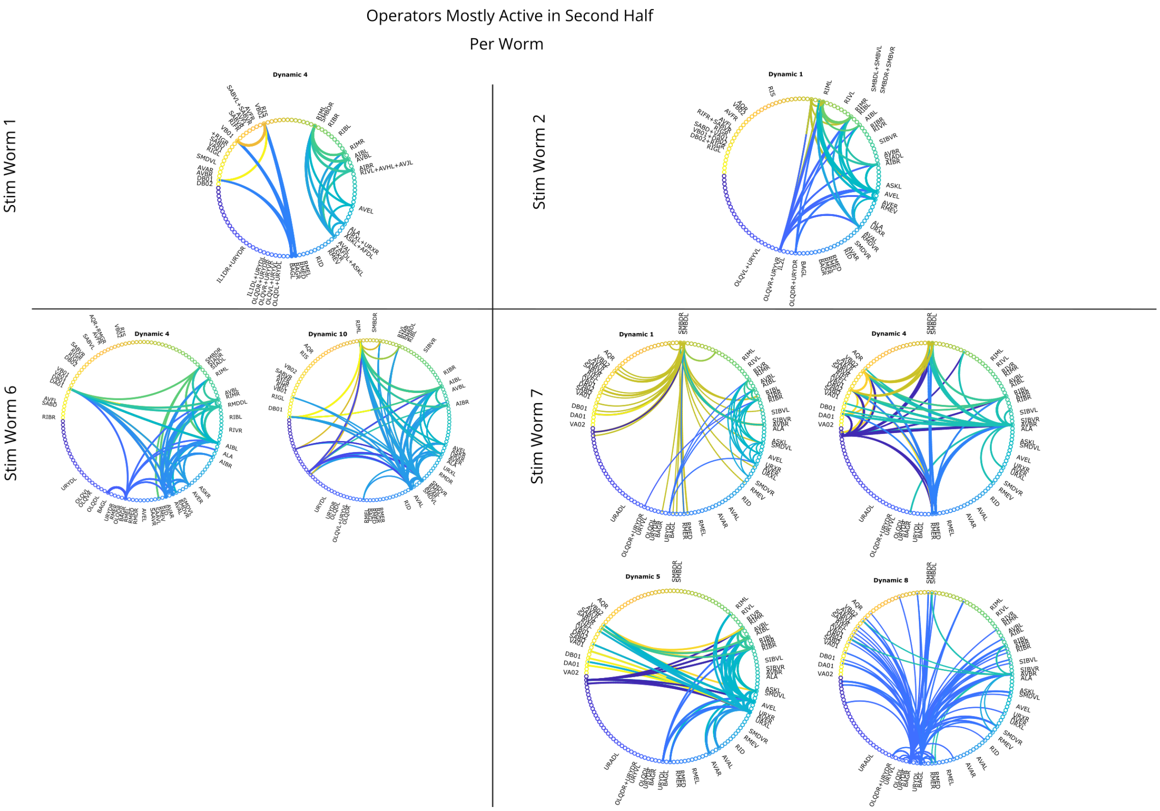


Supplementary Figure 14: **Connectivity maps, DOs active mostly during the alternating oxygen stimulation protocol (second half).** Stim Worms, per-worm DOs. Strength of connection (absolute value) shown by line width; source of line indicated by color.

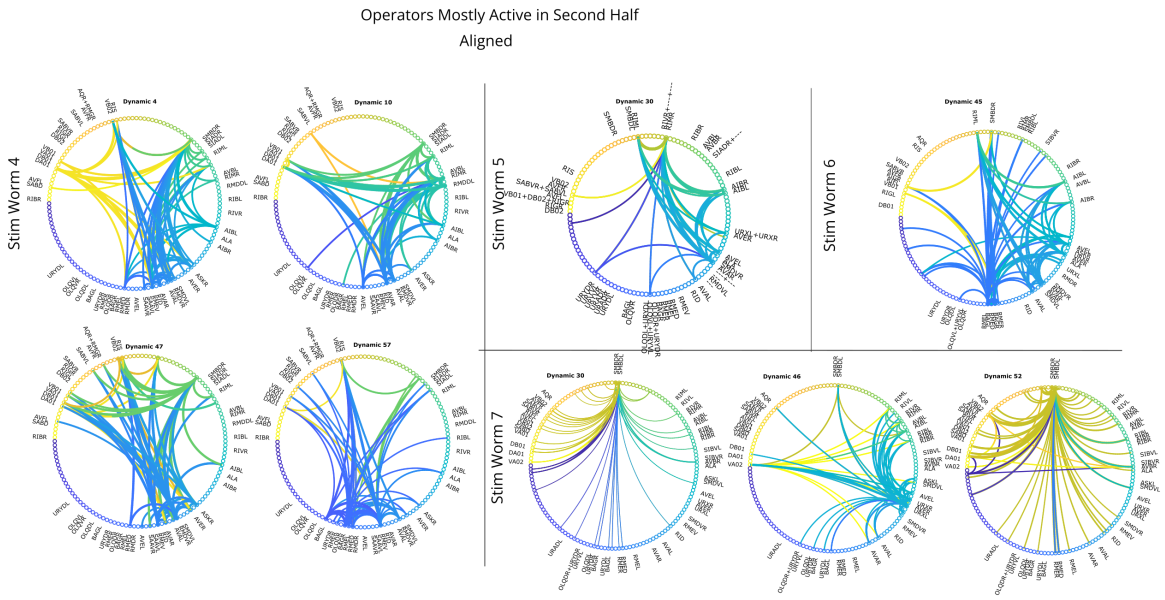


Supplementary Figure 15: **Connectivity maps, DOs active mostly during the alternating oxygen stimulation protocol (second half).** Stim Worms, aligned DOs. Strength of connection (absolute value) shown by line width; source of line indicated by color.

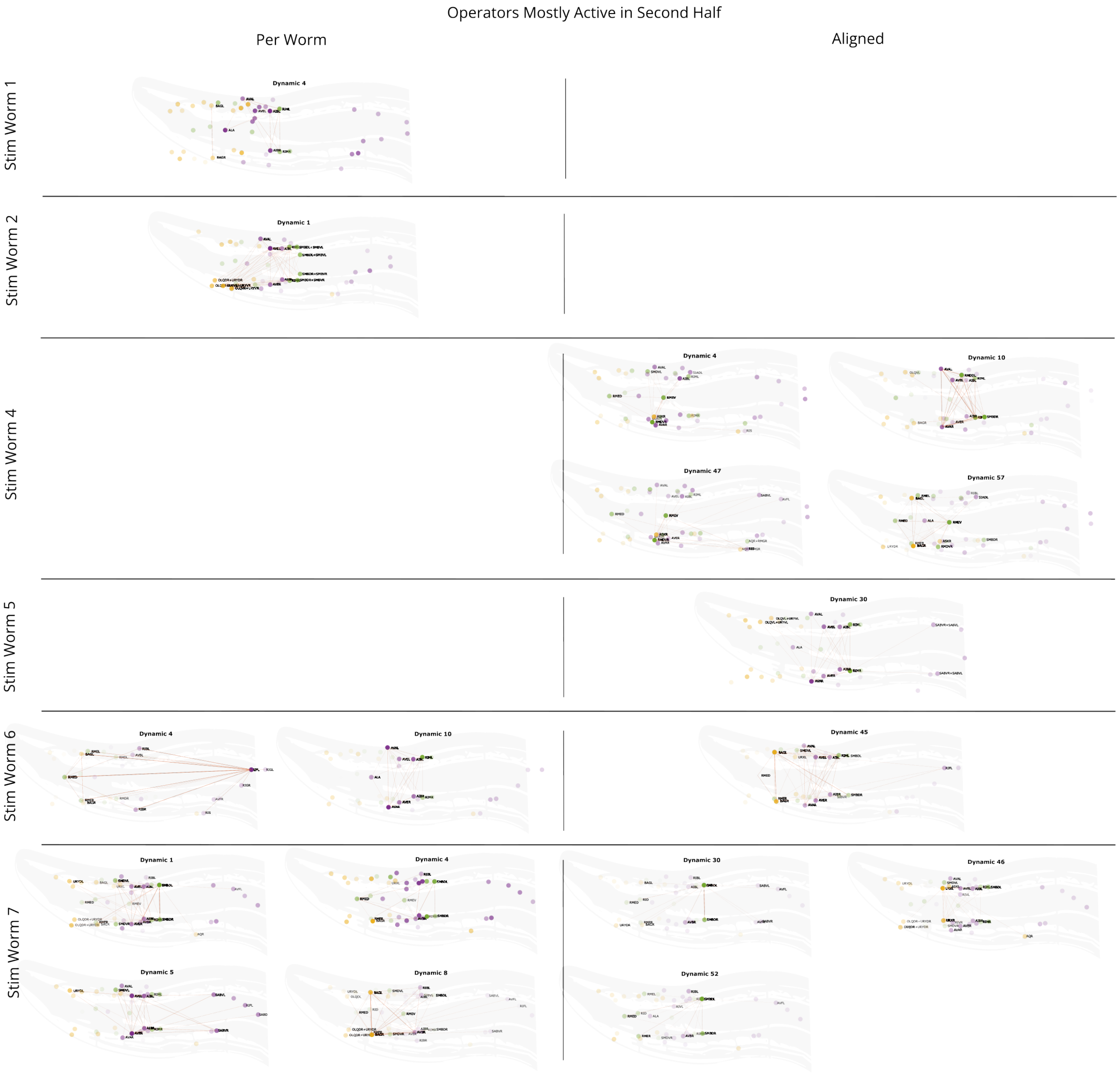


Supplementary Figure 16: **Connectivity maps, DOs active mostly during the alternating oxygen stimulation protocol (second half).** Stim Worms, per-worm and aligned DOs, time-averaged. Strength of connection (absolute value) shown by orange line width. Neuron opacity corresponds to reconstructed activity. Neuron color corresponds to class: yellow sensory, green interneuron, purple motor.

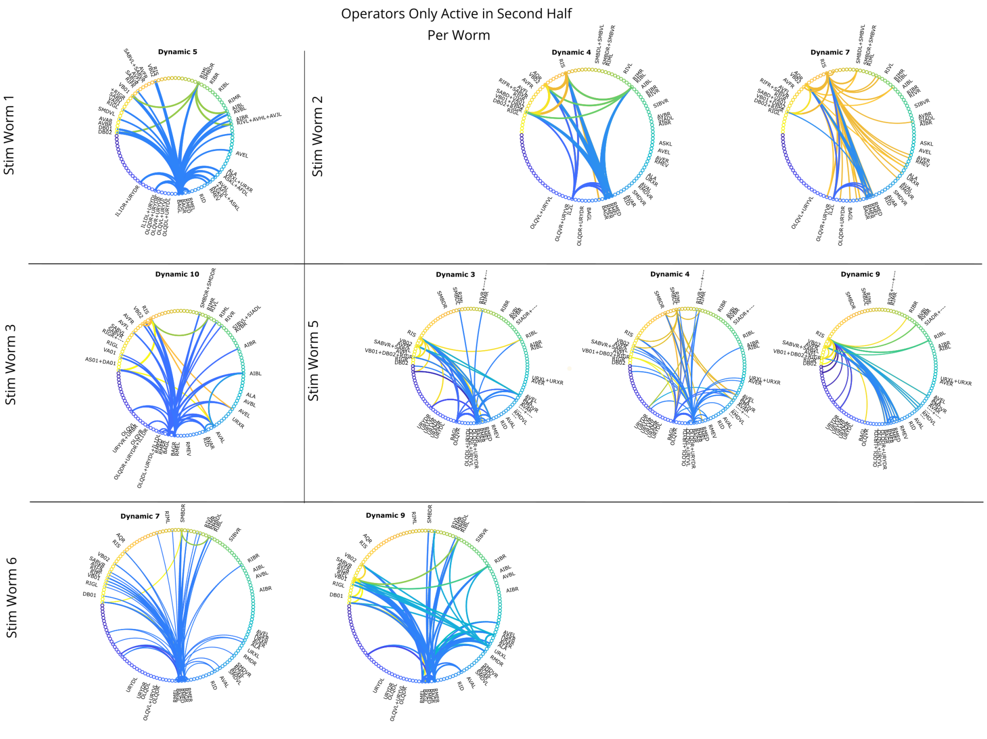


Supplementary Figure 17: **Connectivity maps, DOs active only during the alternating oxygen stimulation protocol (second half).** Stim Worms, per-worm DOs. Strength of connection (absolute value) shown by line width; source of line indicated by color.

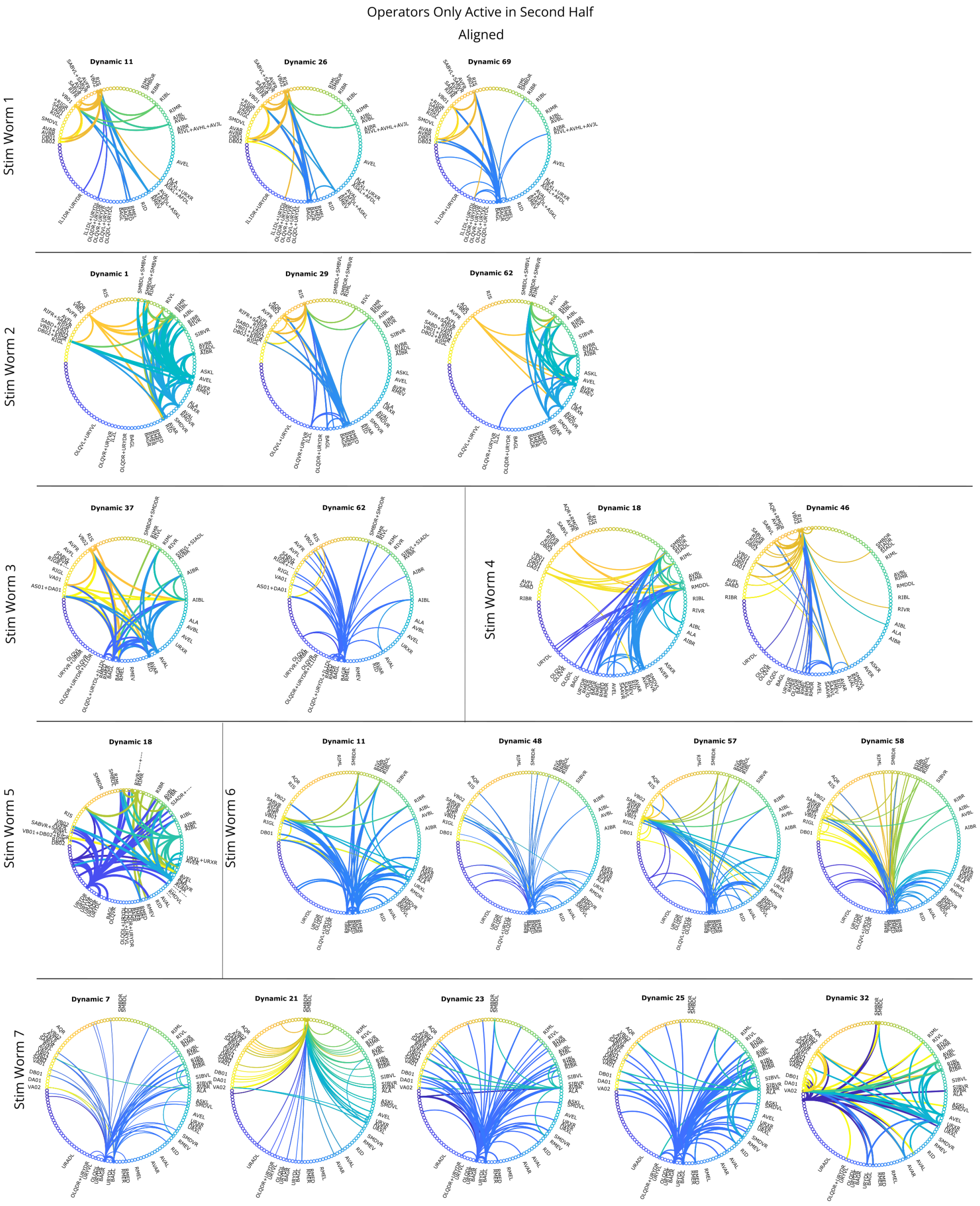


Supplementary Figure 18: **Connectivity maps, DOs active only during the alternating oxygen stimulation protocol (second half).** Stim Worms, aligned DOs. Strength of connection (absolute value) shown by line width; source of line indicated by color.

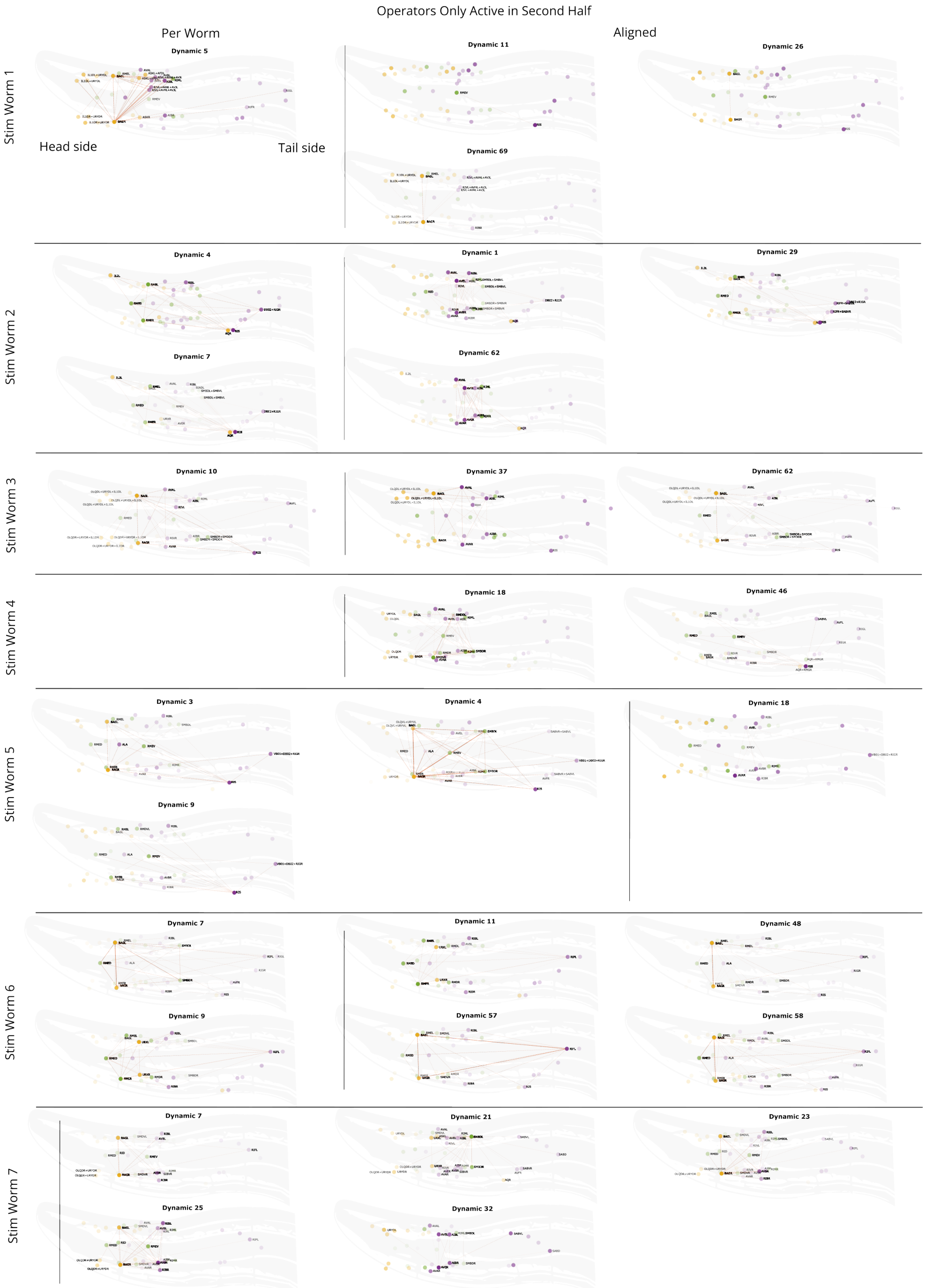


Supplementary Figure 19: **Connectivity maps, DOs active only during the alternating oxygen stimulation protocol (second half).** Stim worms, per-worm and aligned DOs, time-averaged. Strength of connection (absolute value) shown by orange line width. Neuron opacity corresponds to reconstructed activity. Neuron color corresponds to class: yellow sensory, green interneuron, purple motor.

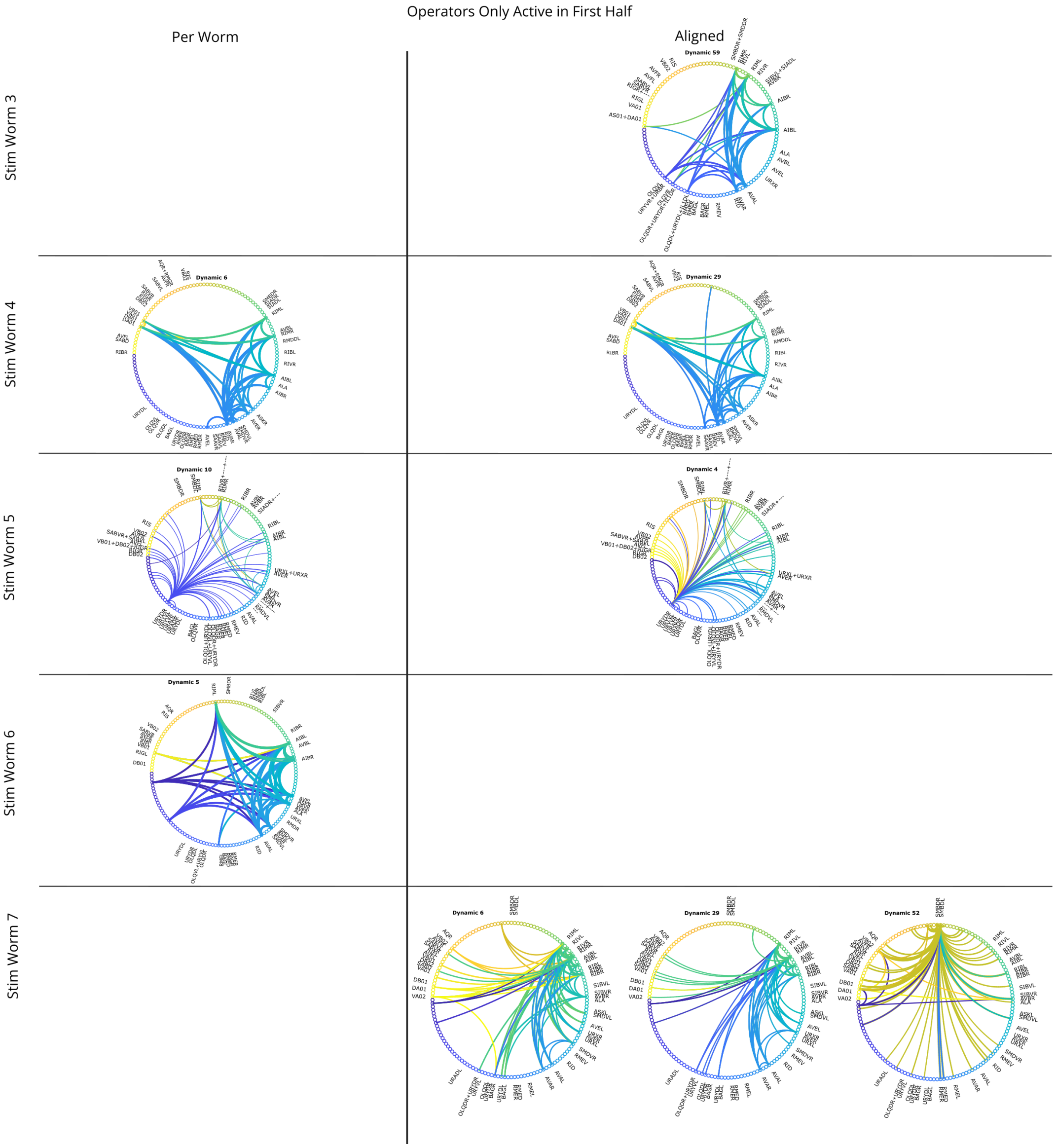


Supplementary Figure 20: **Connectivity maps, DOs active only before the alternating oxygen stimulation protocol (first half).** Stim Worms, per-worm and aligned DOs. Strength of connection (absolute value) shown by line width; source of line indicated by color.

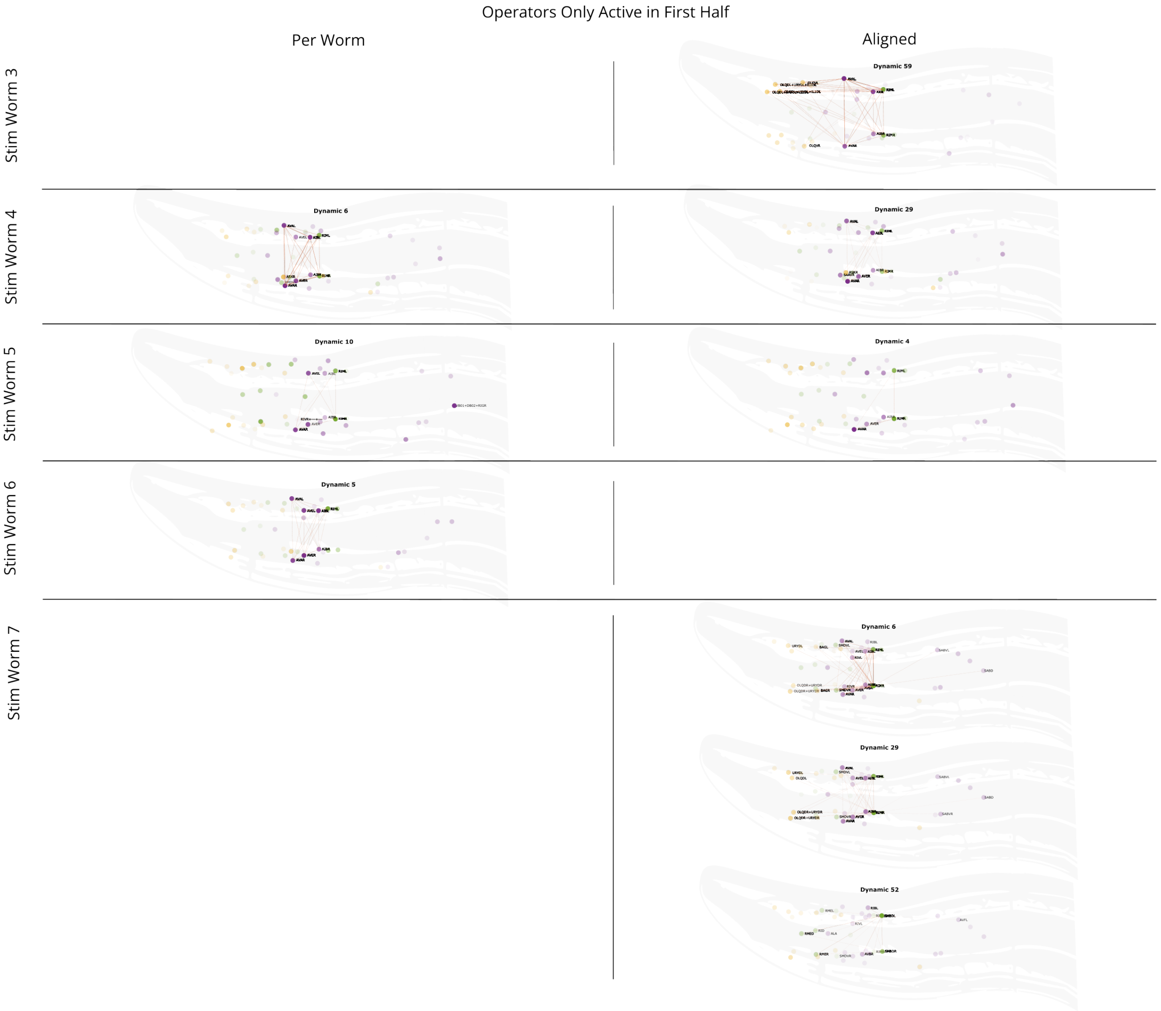


Supplementary Figure 21: **Connectivity maps, DOs active only before the alternating oxygen stimulation protocol (first half).** Stim worms, per-worm and aligned DOs, time-averaged. Strength of connection (absolute value) shown by orange line width. Neuron opacity corresponds to reconstructed activity. Neuron color corresponds to class: yellow sensory, green interneuron, purple motor.

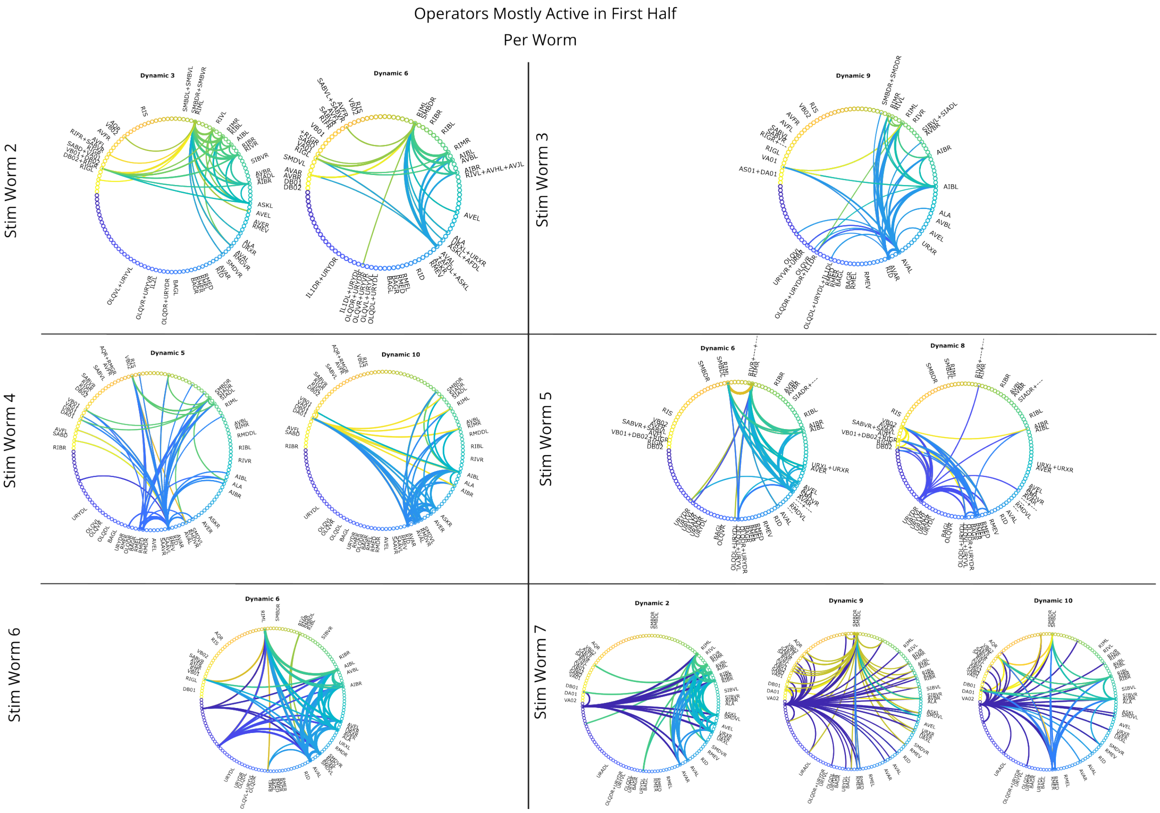


Supplementary Figure 22: **Connectivity maps, DOs active mostly before the alternating oxygen stimulation protocol (first half).** Stim Worms, per-worm DOs. Strength of connection (absolute value) shown by line width; source of line indicated by color.

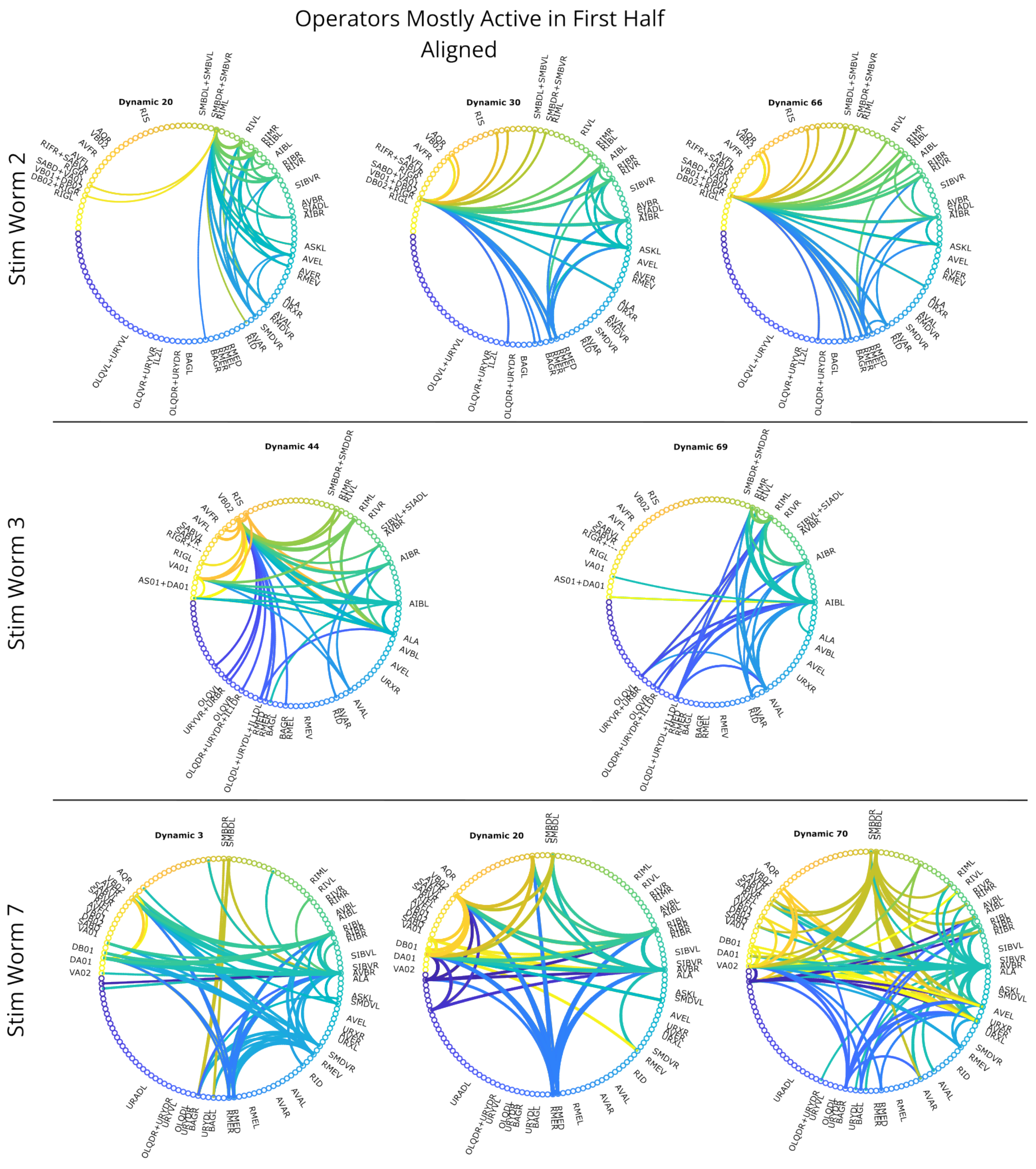


Supplementary Figure 23: **Connectivity maps, DOs active mostly before the alternating oxygen stimulation protocol (first half).** Stim Worms, aligned DOs. Strength of connection (absolute value) shown by line width; source of line indicated by color.

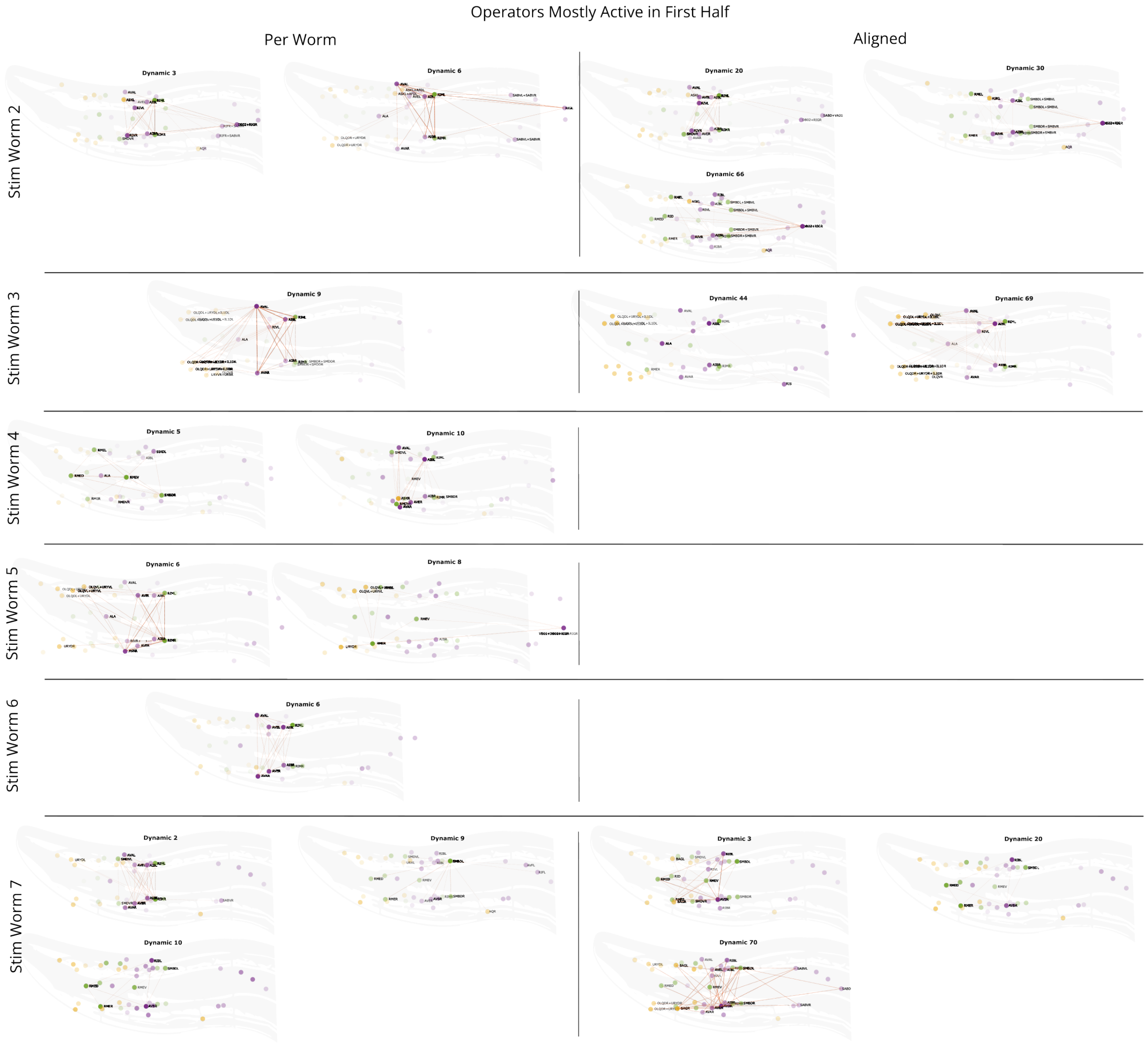


Supplementary Figure 24: **Connectivity maps, DOs active mostly before the alternating oxygen stimulation protocol (first half).** Stim Worms, per-worm and aligned DOs, time-averaged. Strength of connection (absolute value) shown by orange line width. Neuron opacity corresponds to reconstructed activity. Neuron color corresponds to class: yellow sensory, green interneuron, purple motor.

Forward Crawl (Per-Worm)

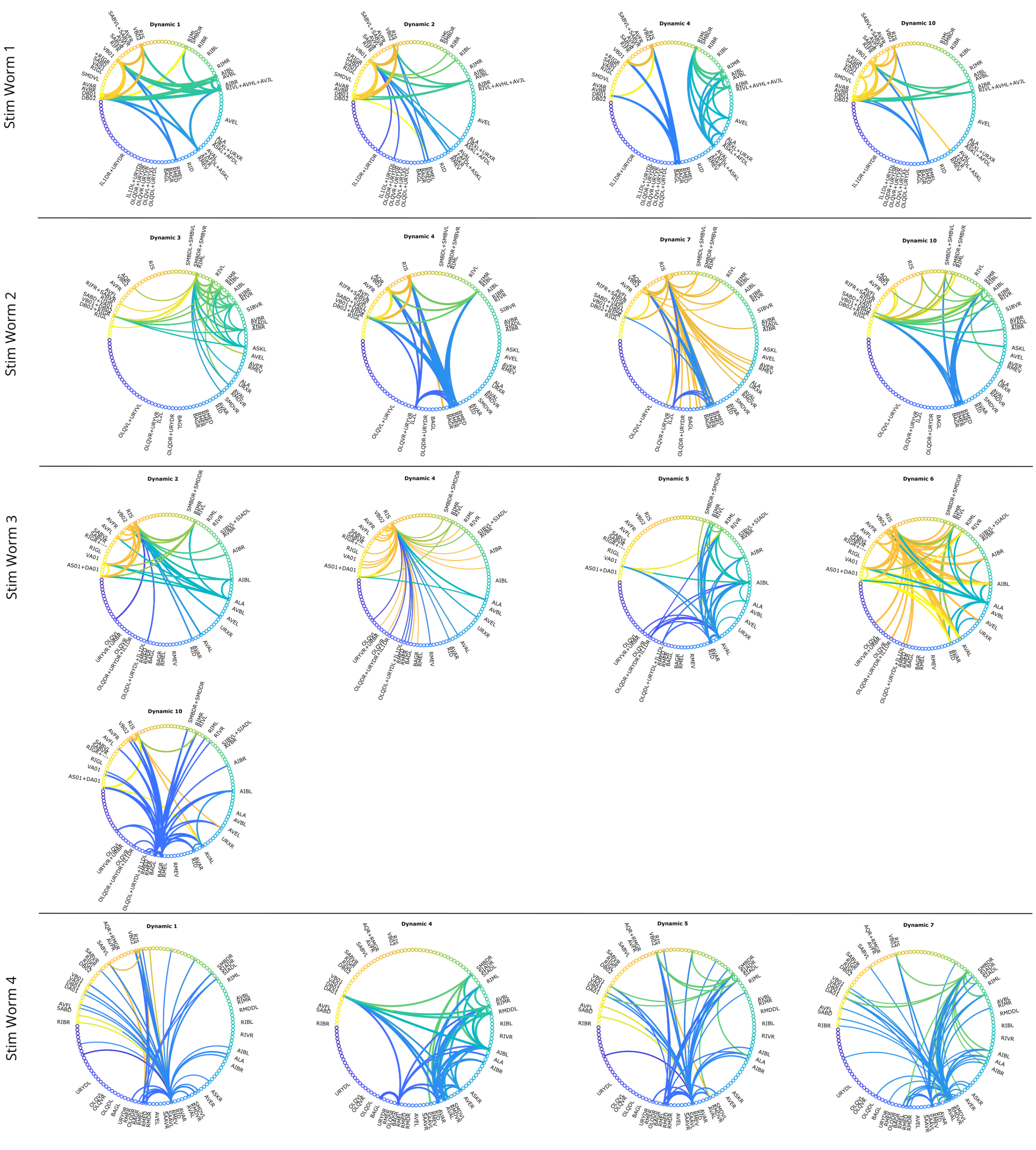


Supplementary Figure 25: **Connectivity maps by per-worm dynamics operator for each Stim worm during Forward Crawl (State 1), part 1.** Strength of connection (absolute value) shown by line width; source of line indicated by color.

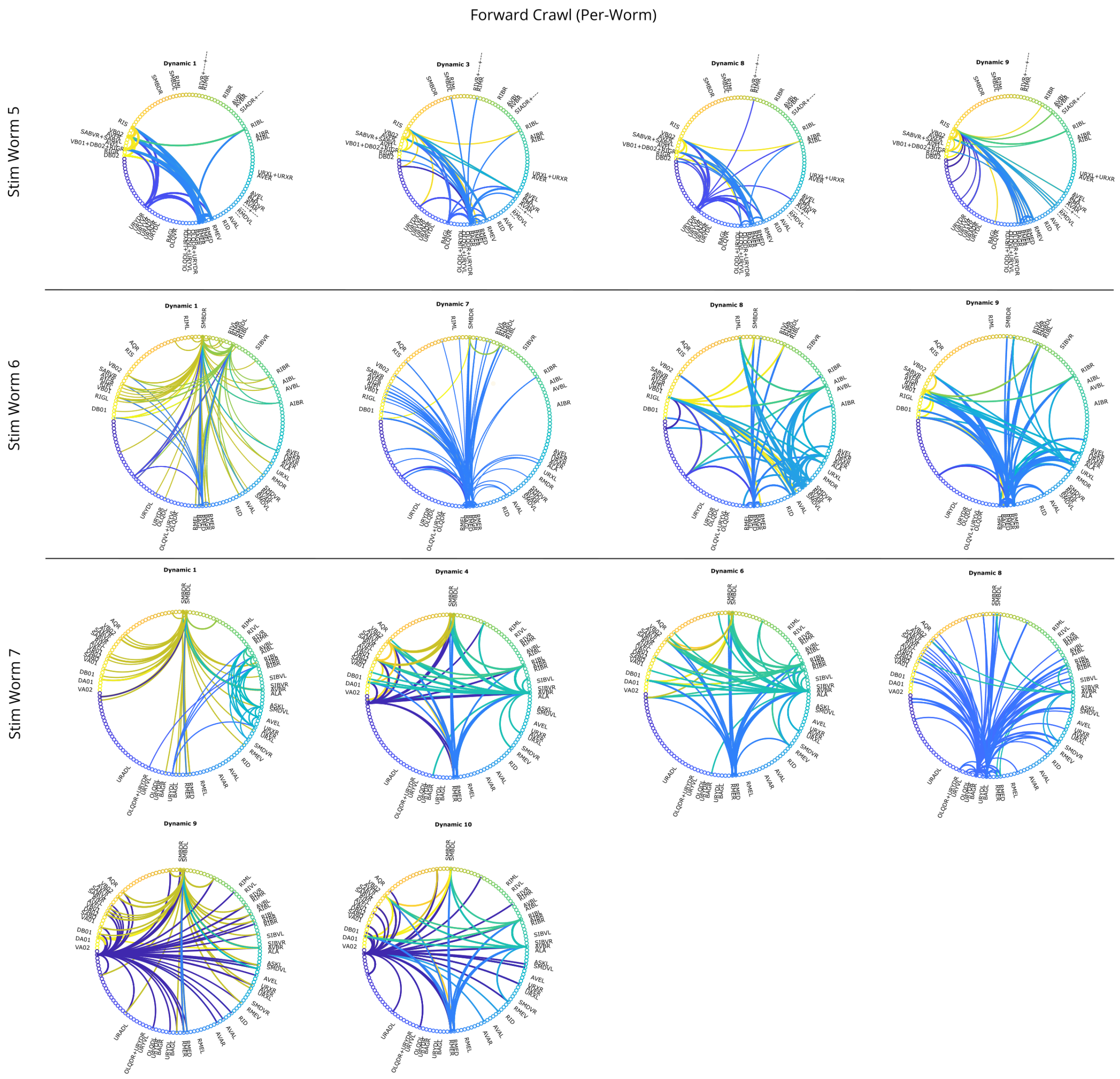


Supplementary Figure 26: **Connectivity maps by per-worm dynamics operator for each Stim worm during Forward Crawl (State 1), part 2.** Strength of connection (absolute value) shown by line width; source of line indicated by color.

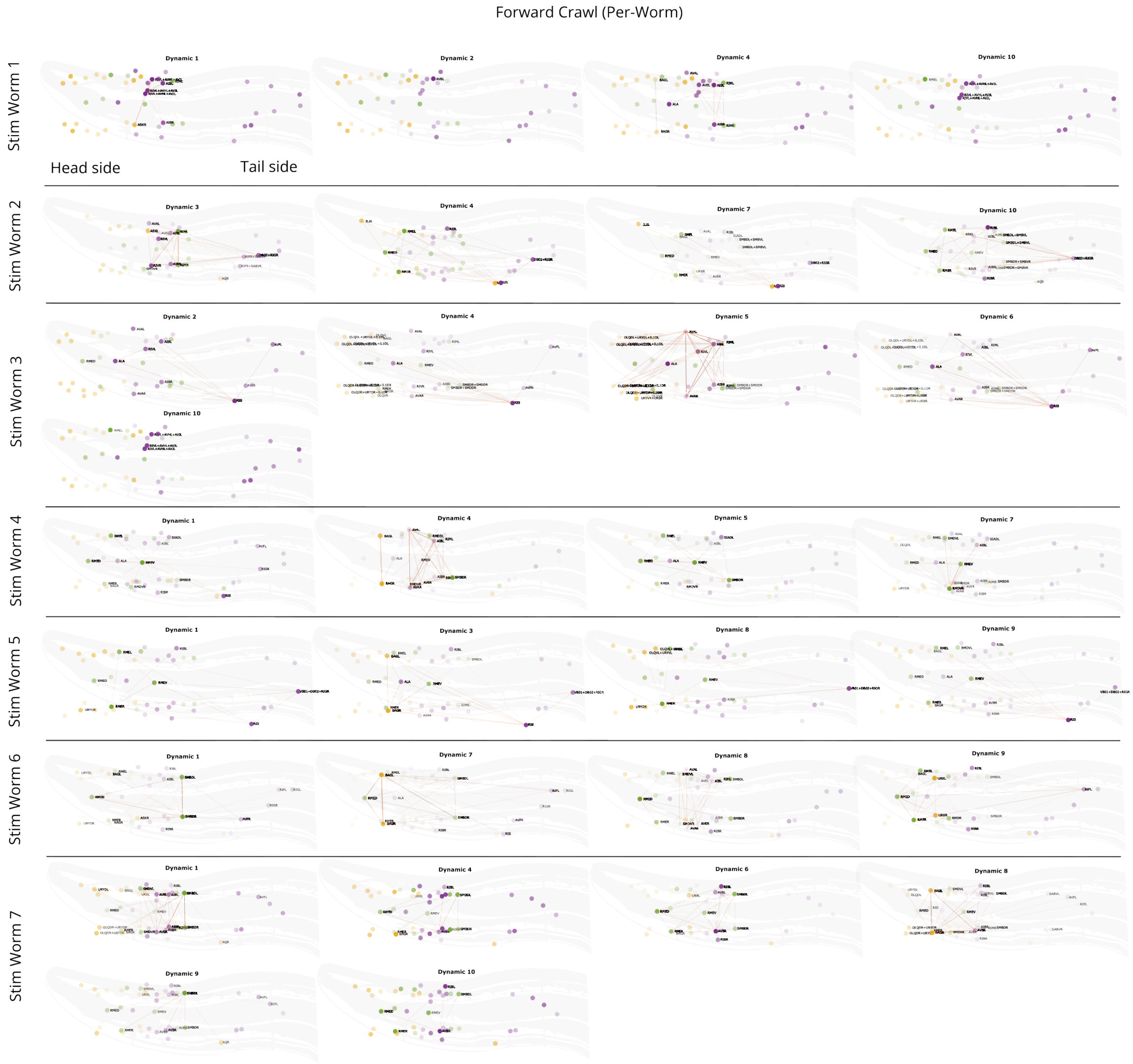


Supplementary Figure 27: **Connectivity maps by per-worm dynamics operator for each Stim worm during Forward Crawl (State 1), time-averaged.** Strength of connection (absolute value) shown by orange line width. Neuron opacity corresponds to reconstructed activity. Neuron color corresponds to class: yellow sensory, green interneuron, purple motor.

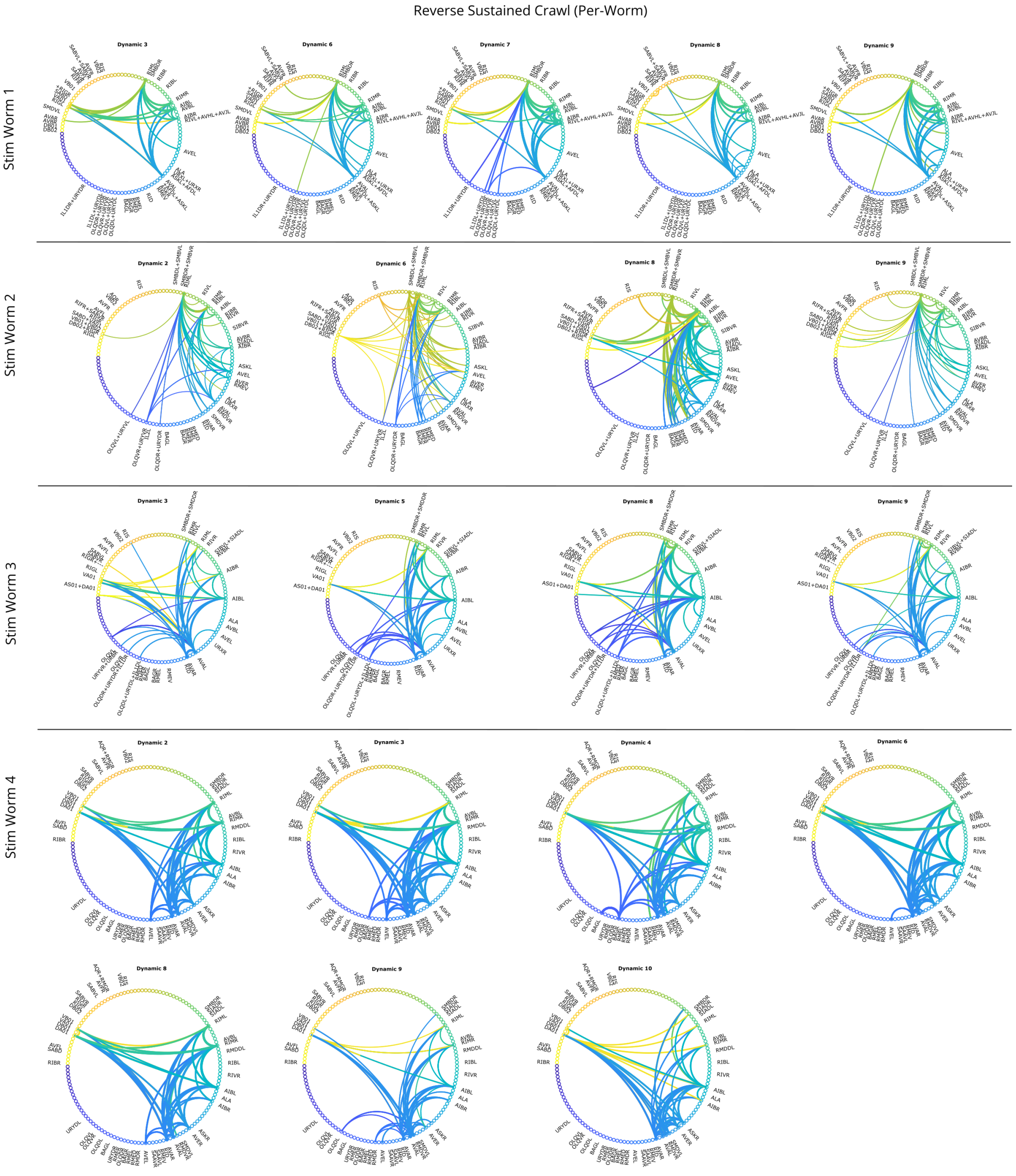


Supplementary Figure 28: **Connectivity maps by per-worm dynamics operator for each Stim worm during Reverse Sustained Crawl (State 3), part 1.** Strength of connection (absolute value) shown by line width; source of line indicated by color.

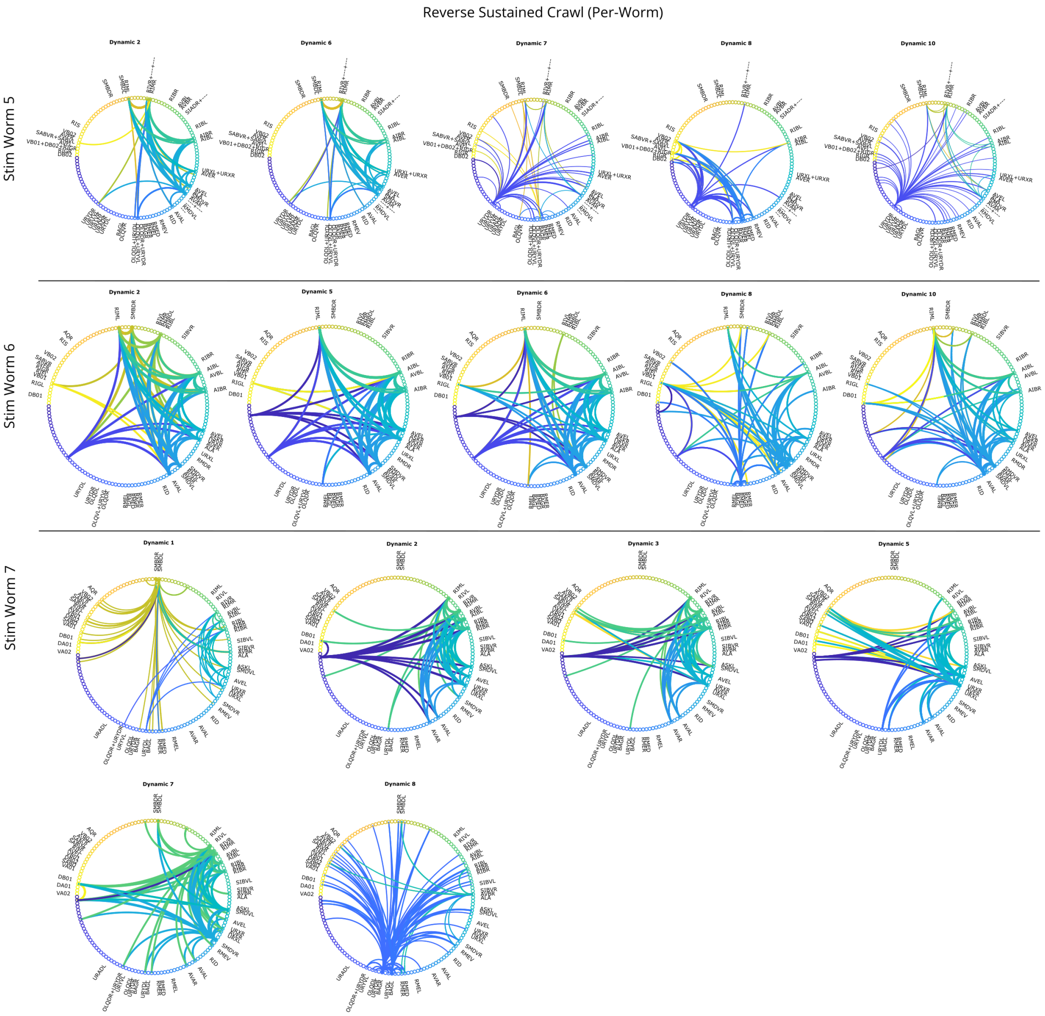


Supplementary Figure 29: **Connectivity maps by per-worm dynamics operator for each Stim worm during Reverse Sustained Crawl (State 3), part 2.** Strength of connection (absolute value) shown by line width; source of line indicated by color.

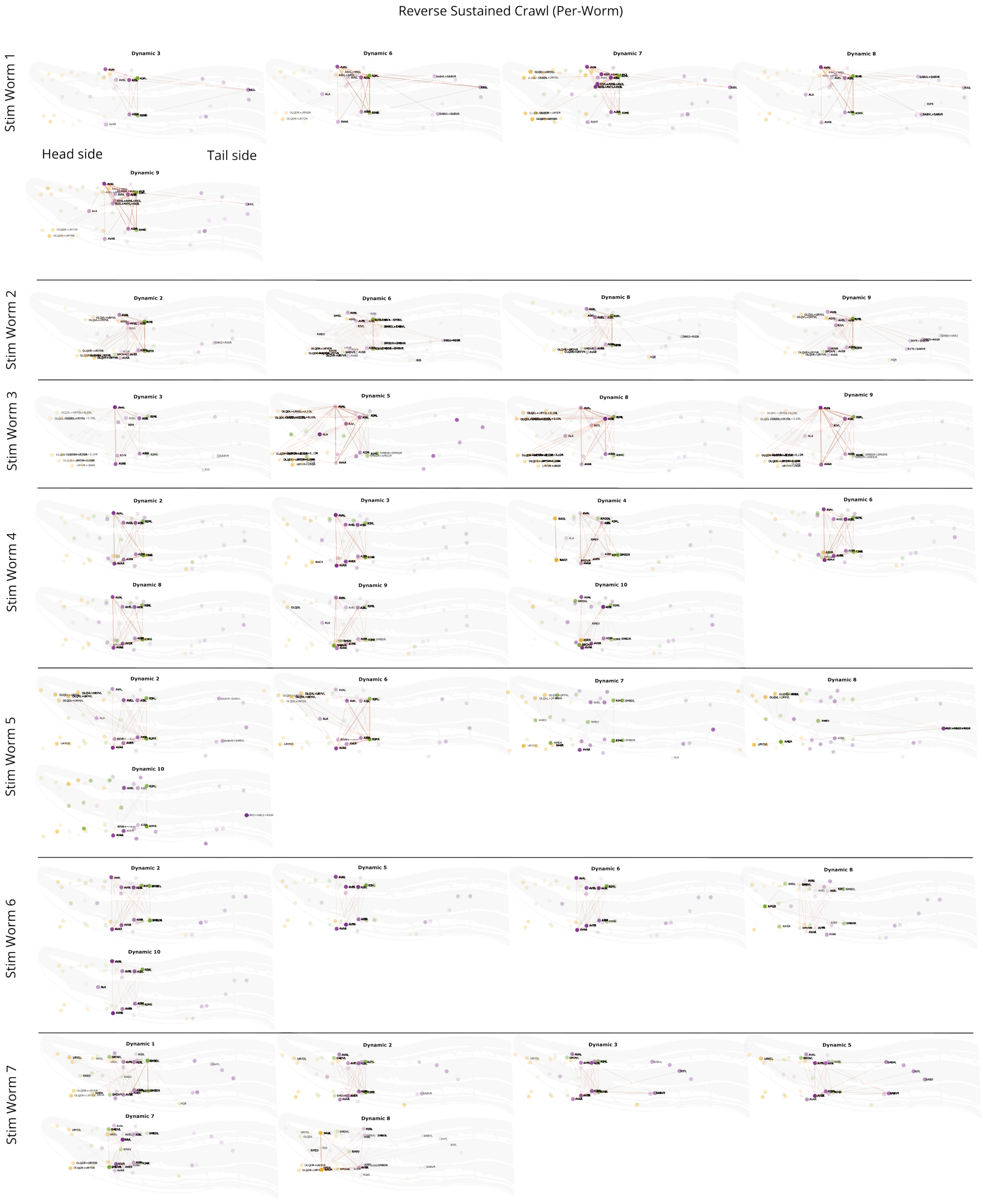


Supplementary Figure 30: **Connectivity maps by per-worm dynamics operator for each Stim worm during Reverse Sustained Crawl (State 3), time-averaged.** Strength of connection (absolute value) shown by orange line width. Neuron opacity corresponds to reconstructed activity. Neuron color corresponds to class: yellow sensory, green interneuron, purple motor.

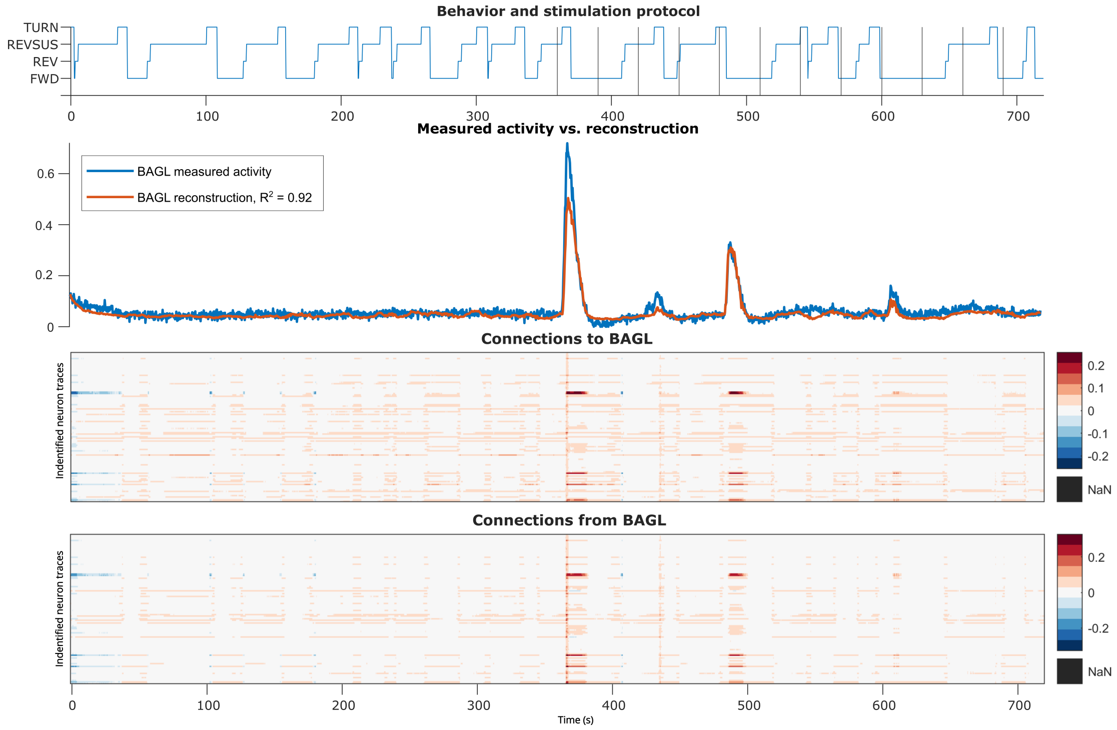


Supplementary Figure 31: **Stim Worm 1 BAGL sensory neuron reconstructed activity and connectivity.**

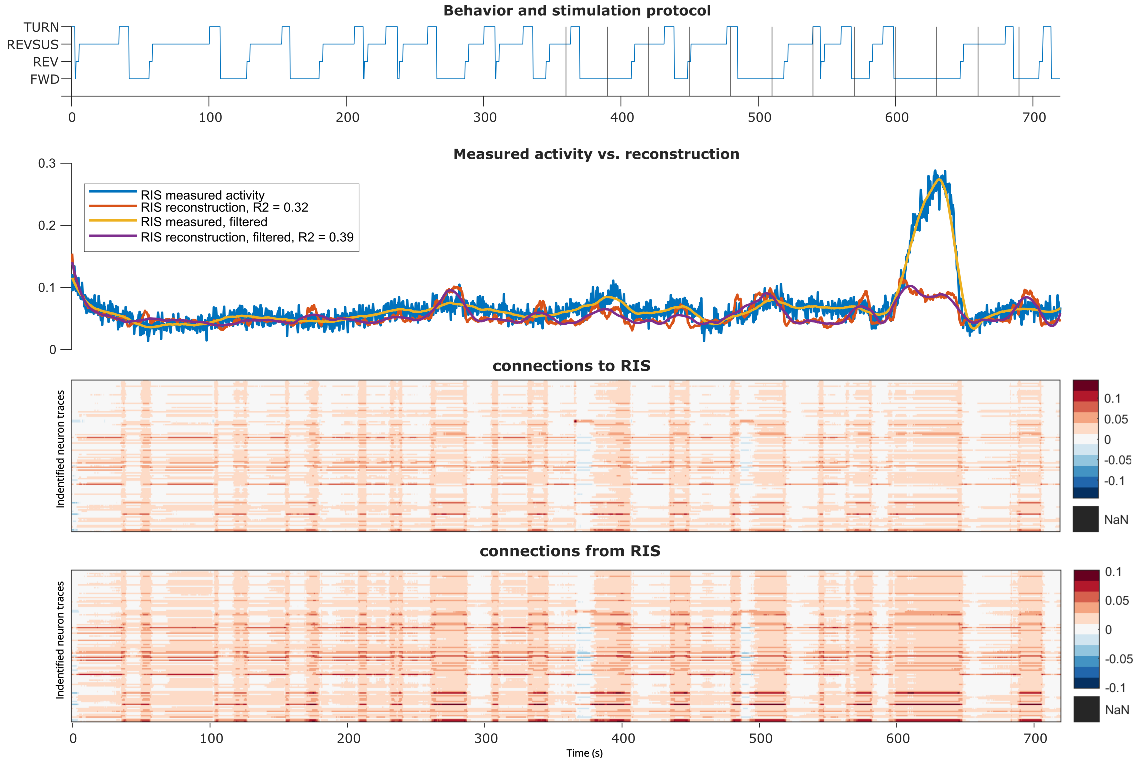


Supplementary Figure 32: **Stim Worm 1 RIS interneuron reconstructed activity and connectivity.**

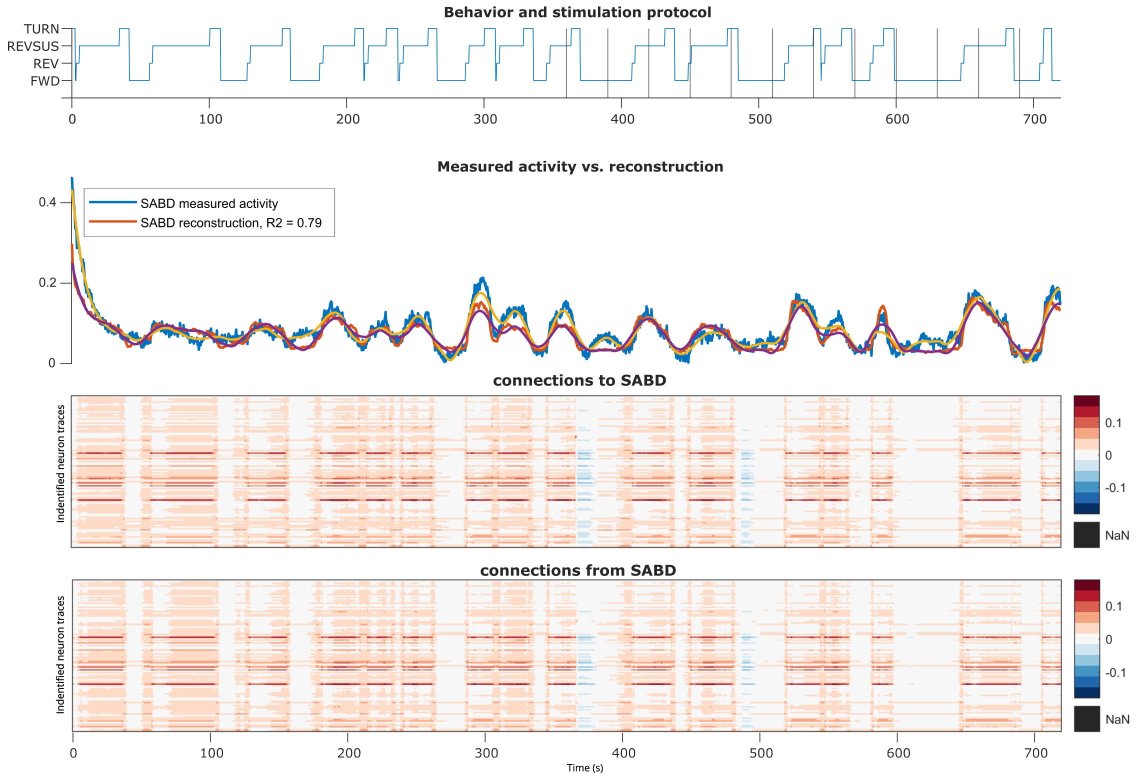


Supplementary Figure 33: **Stim Worm 1 SABD inter-/motor neuron reconstructed activity and connectivity.**

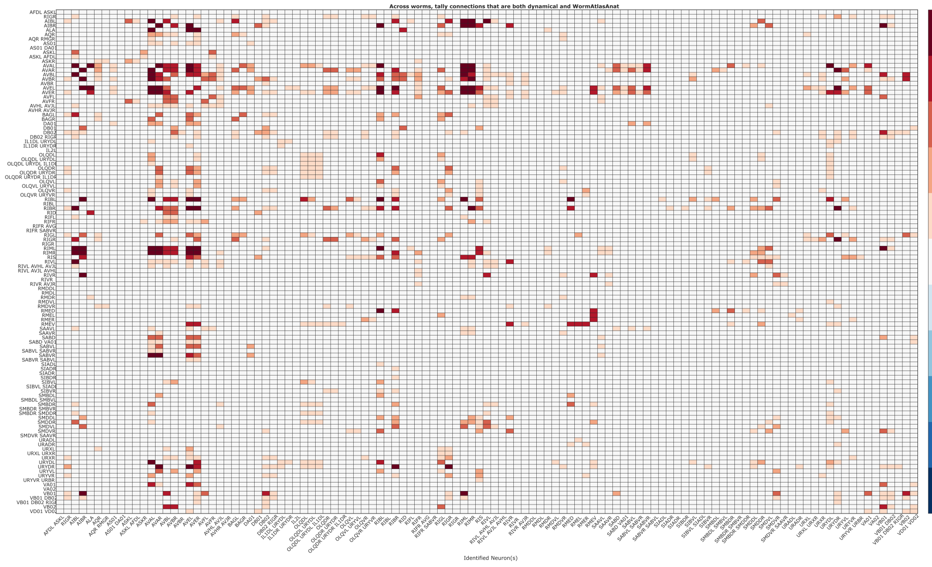


Supplementary Figure 34: **Number of connections recognized by both dLDS and the Worm Atlas [6].** Tallied strong connections (absolute value above a threshold) across all 12 Stim and NoStim worms.

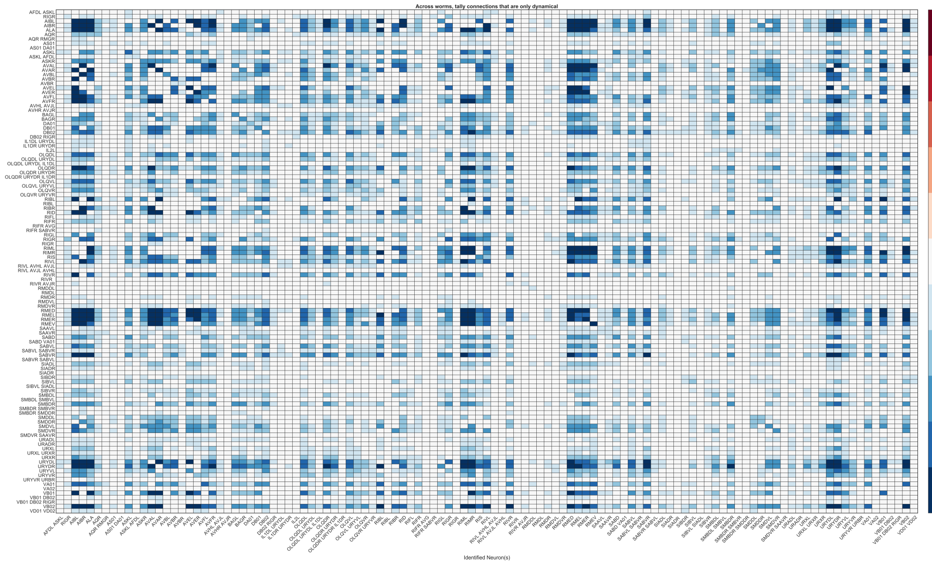


Supplementary Figure 35: **Number of connections recognized by only dLDS, not the Worm Atlas [6].** Tallied strong connections (absolute value above a threshold) across all 12 Stim and NoStim worms.

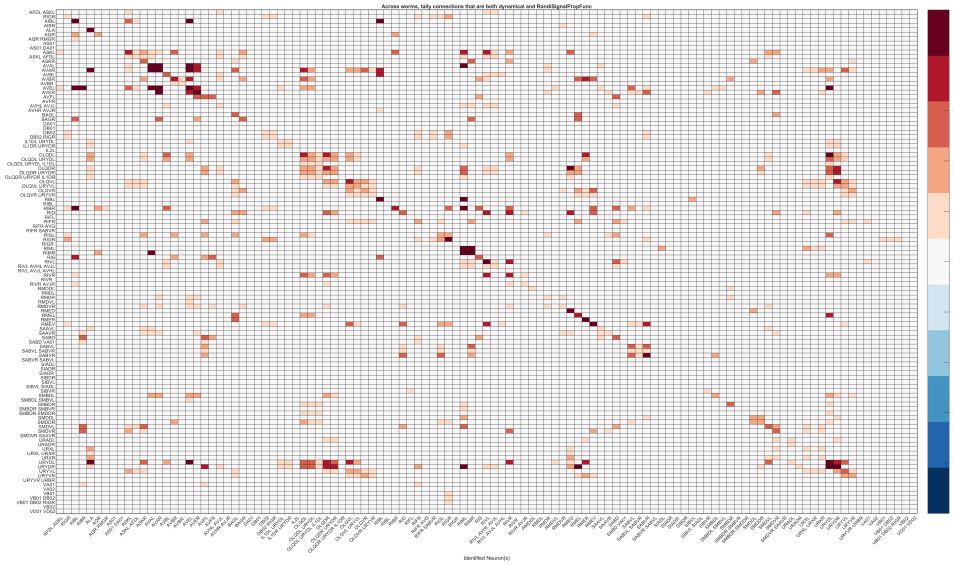


Supplementary Figure 36: **Number of connections recognized by both dLDS and the Randi *et al.* functional connectivity atlas.** Tallied strong connections (absolute value above a threshold) across all 12 Stim and NoStim worms.

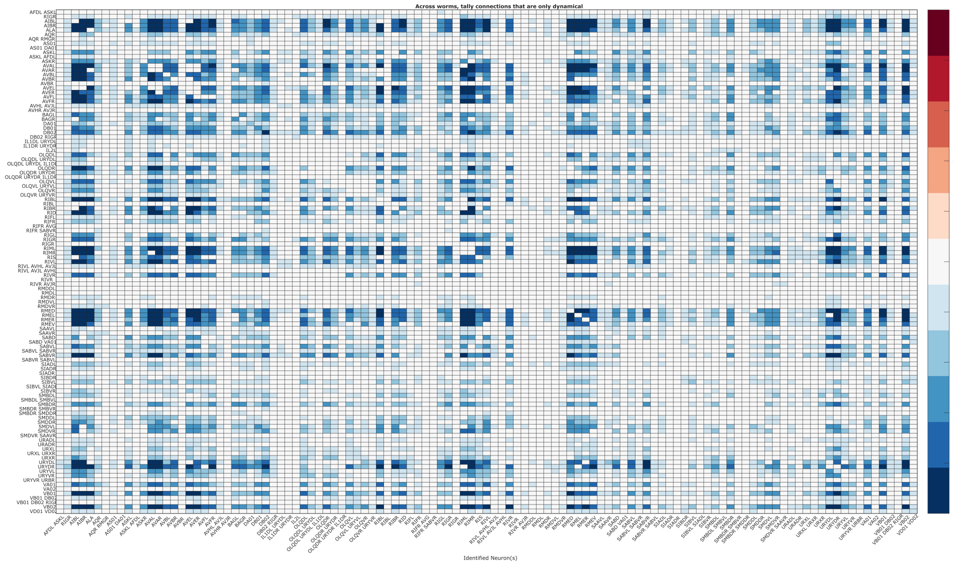


Supplementary Figure 37: **Number of connections recognized by only dLDS, not the Randi *et al.* functional connectivity atlas.** Tallied strong connections (absolute value above a threshold) across all 12 Stim and NoStim worms.

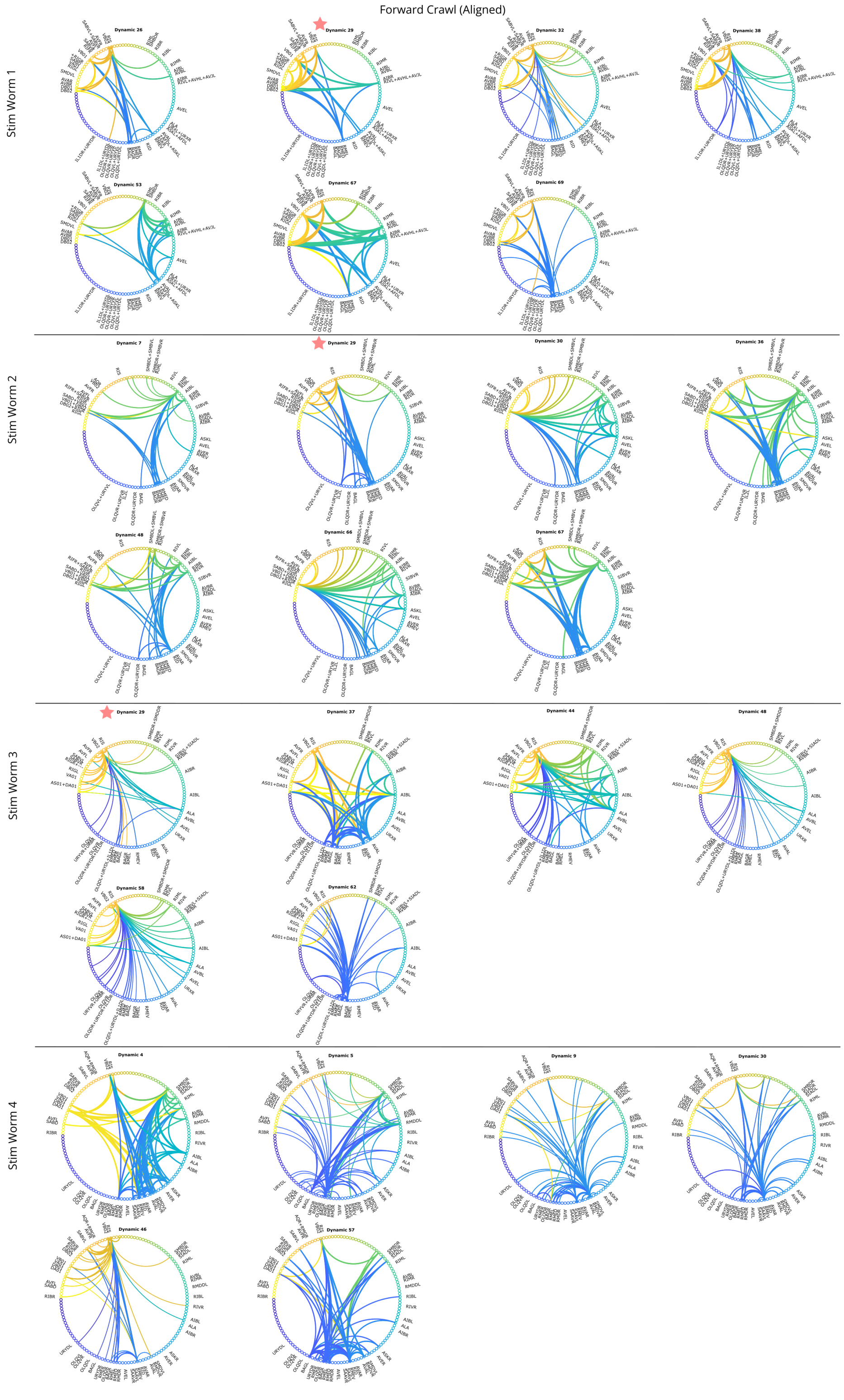


Supplementary Figure 38: **Connectivity maps by aligned dynamics operator for each Stim worm during Forward Crawl (State 1), part 1.** Strength of connection (absolute value) shown by line width; source of line indicated by color.

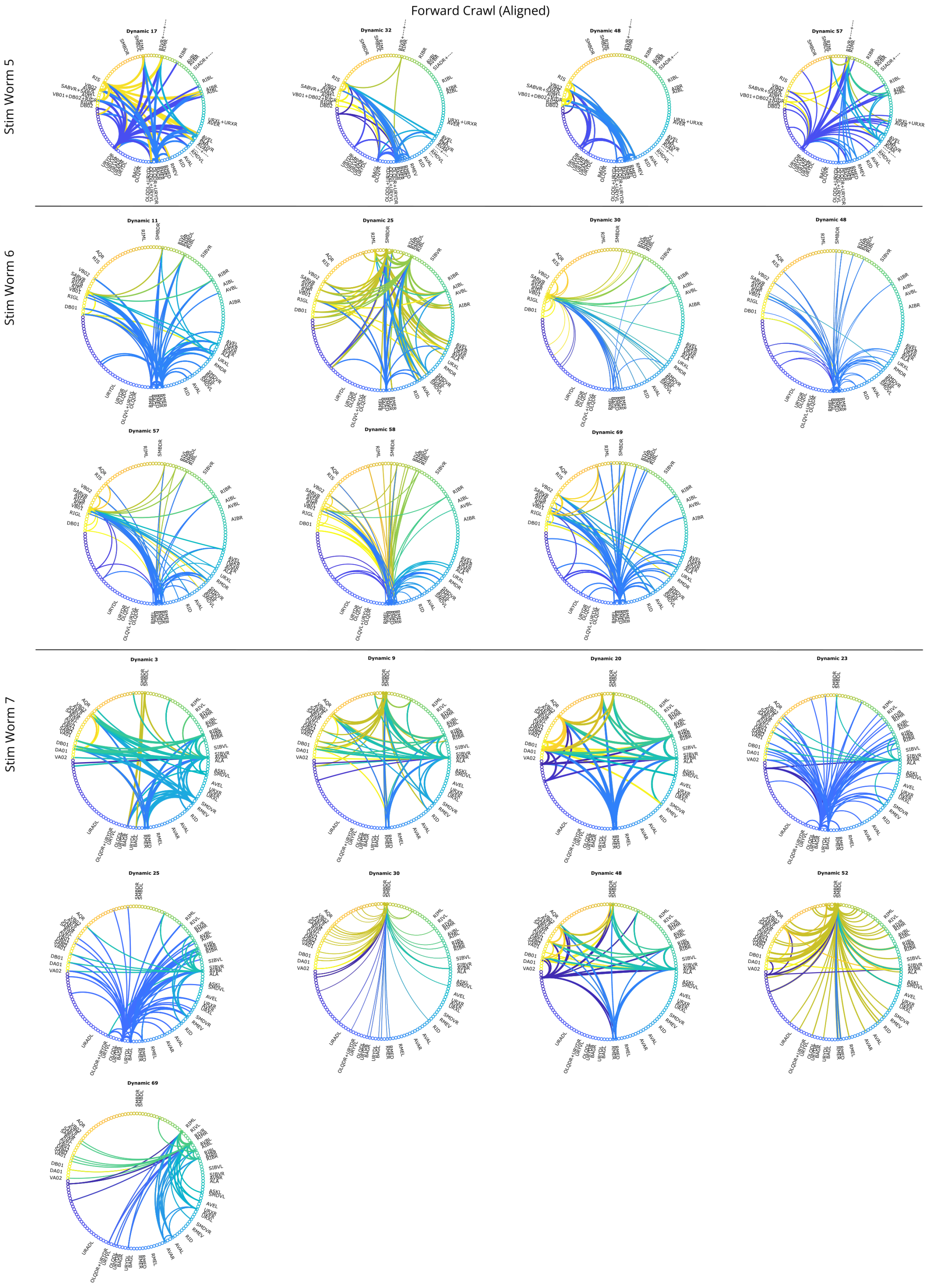


Supplementary Figure 39: **Connectivity maps by aligned dynamics operator for each Stim worm during Forward Crawl (State 1), part 2.** Strength of connection (absolute value) shown by line width; source of line indicated by color.

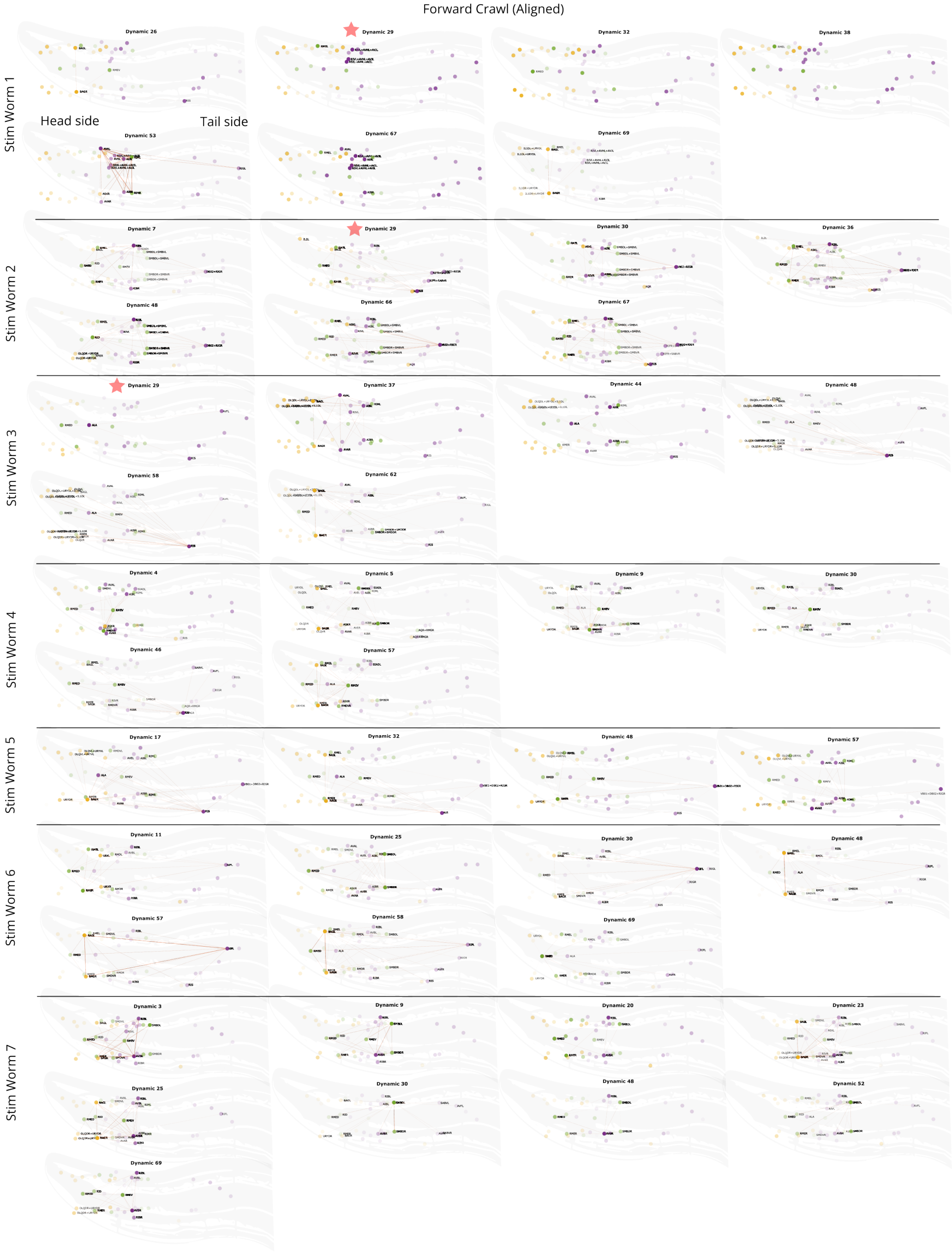


Supplementary Figure 40: **Connectivity maps by aligned dynamics operator for each Stim worm during Forward Crawl (State 1), time-averaged.** Strength of connection (absolute value) shown by orange line width. Neuron opacity corresponds to reconstructed activity. Neuron color corresponds to class: yellow sensory, green interneuron, purple motor.

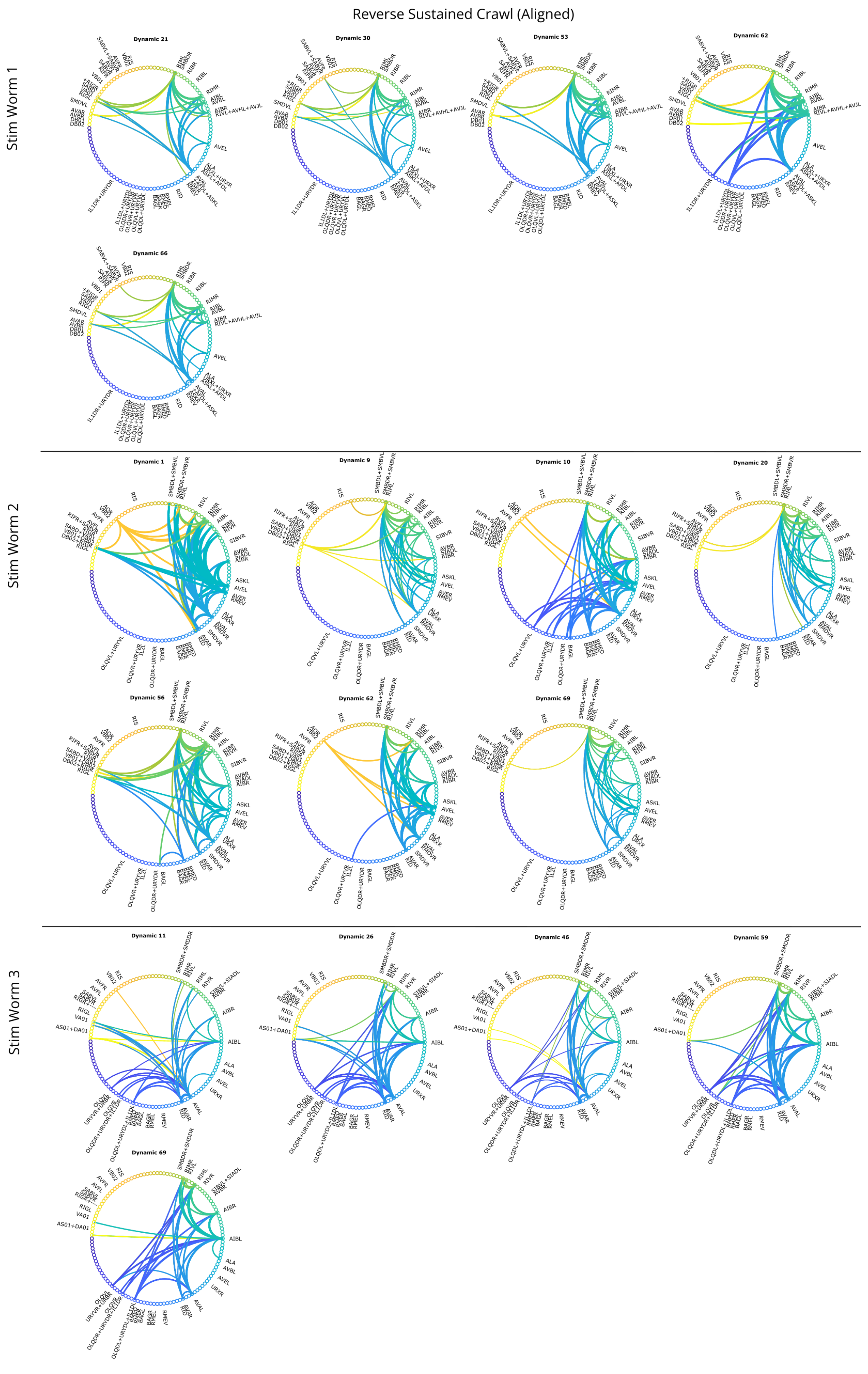


Supplementary Figure 41: **Connectivity maps by aligned dynamics operator for each Stim worm during Reverse Sustained Crawl (State 3), part 1.** Strength of connection (absolute value) shown by line width; source of line indicated by color.

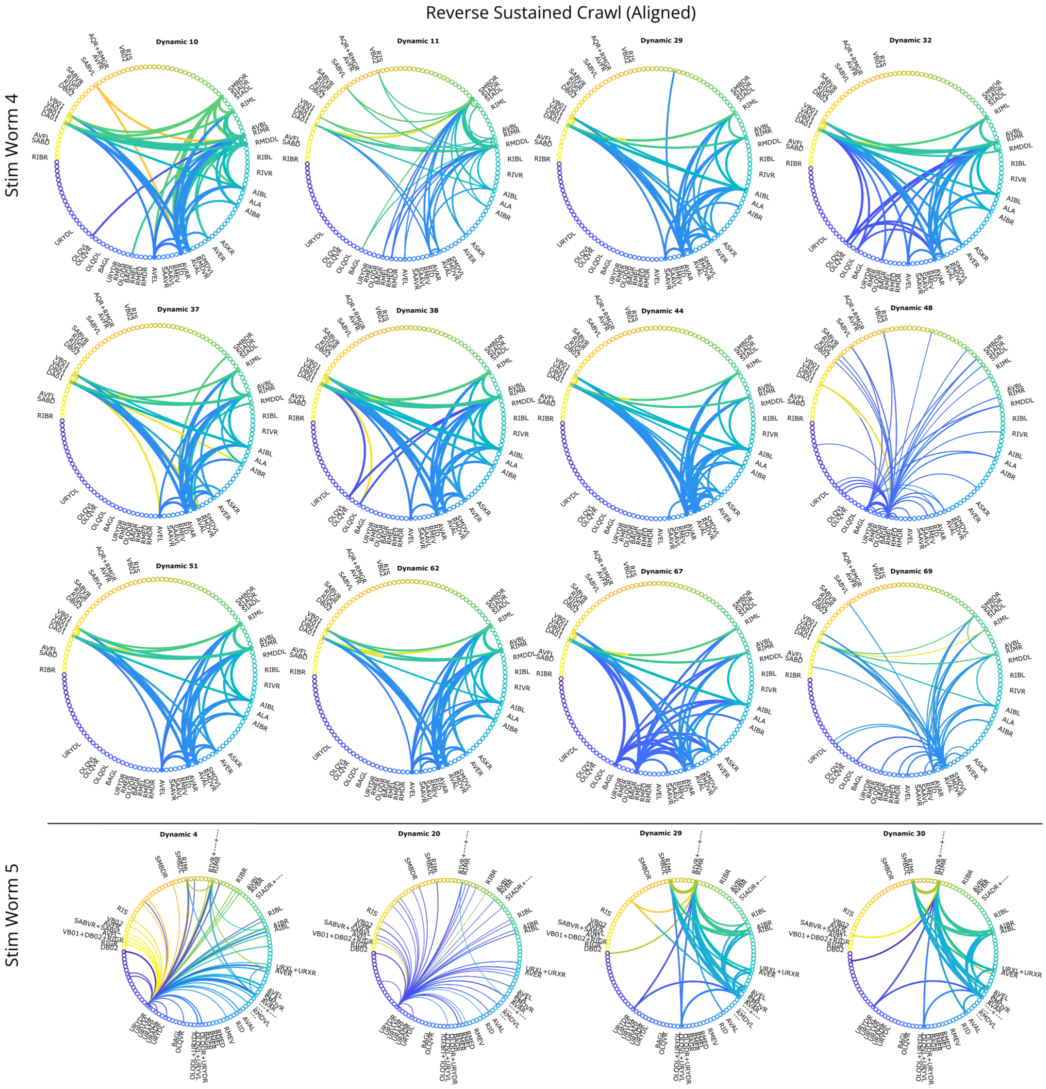


Supplementary Figure 42: **Connectivity maps by aligned dynamics operator for each Stim worm during Reverse Sustained Crawl (State 3), part 2.** Strength of connection (absolute value) shown by line width; source of line indicated by color.

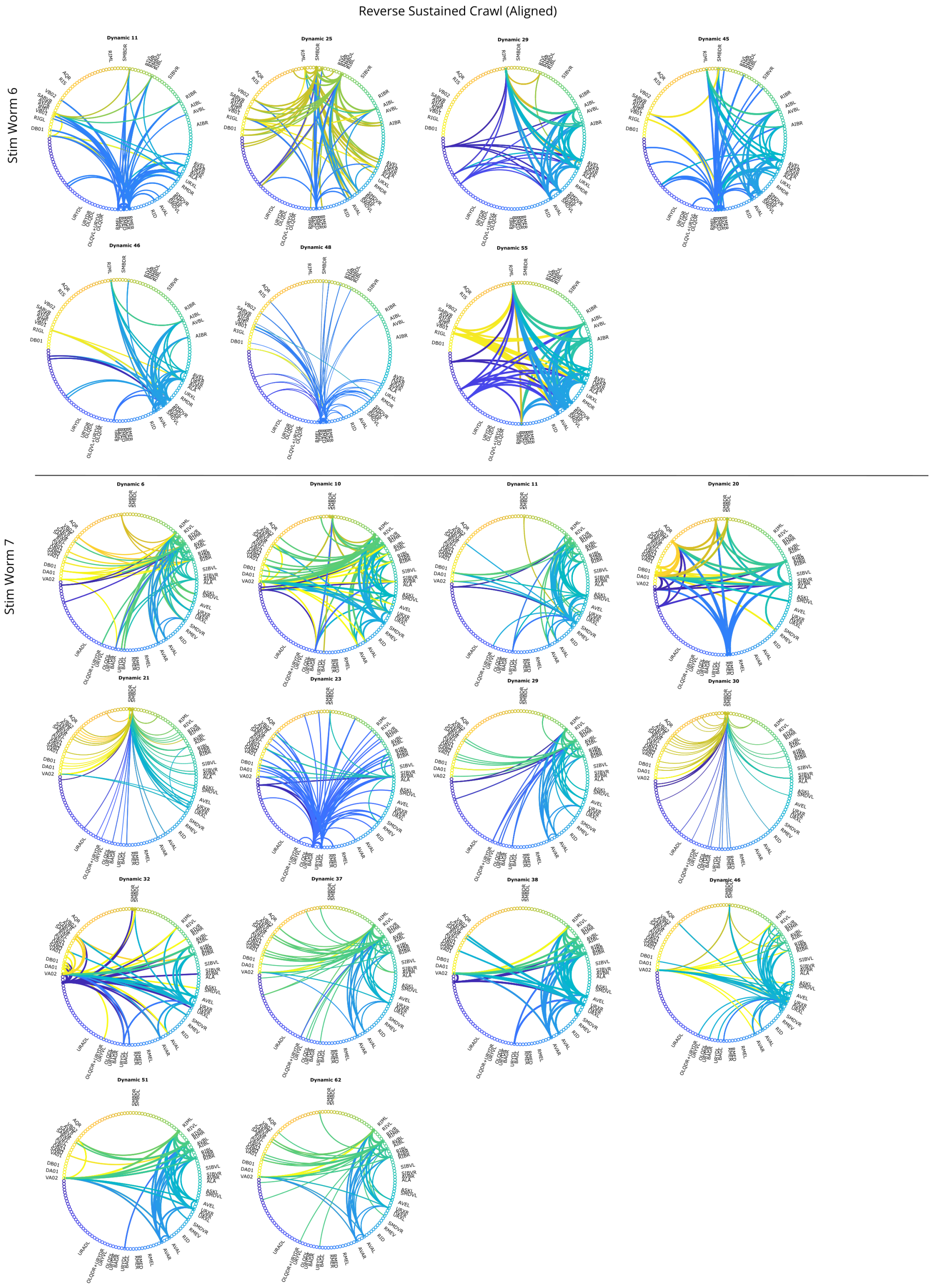


Supplementary Figure 43: **Connectivity maps by aligned dynamics operator for each Stim worm during Reverse Sustained Crawl (State 3), part 3.** Strength of connection (absolute value) shown by line width; source of line indicated by color.

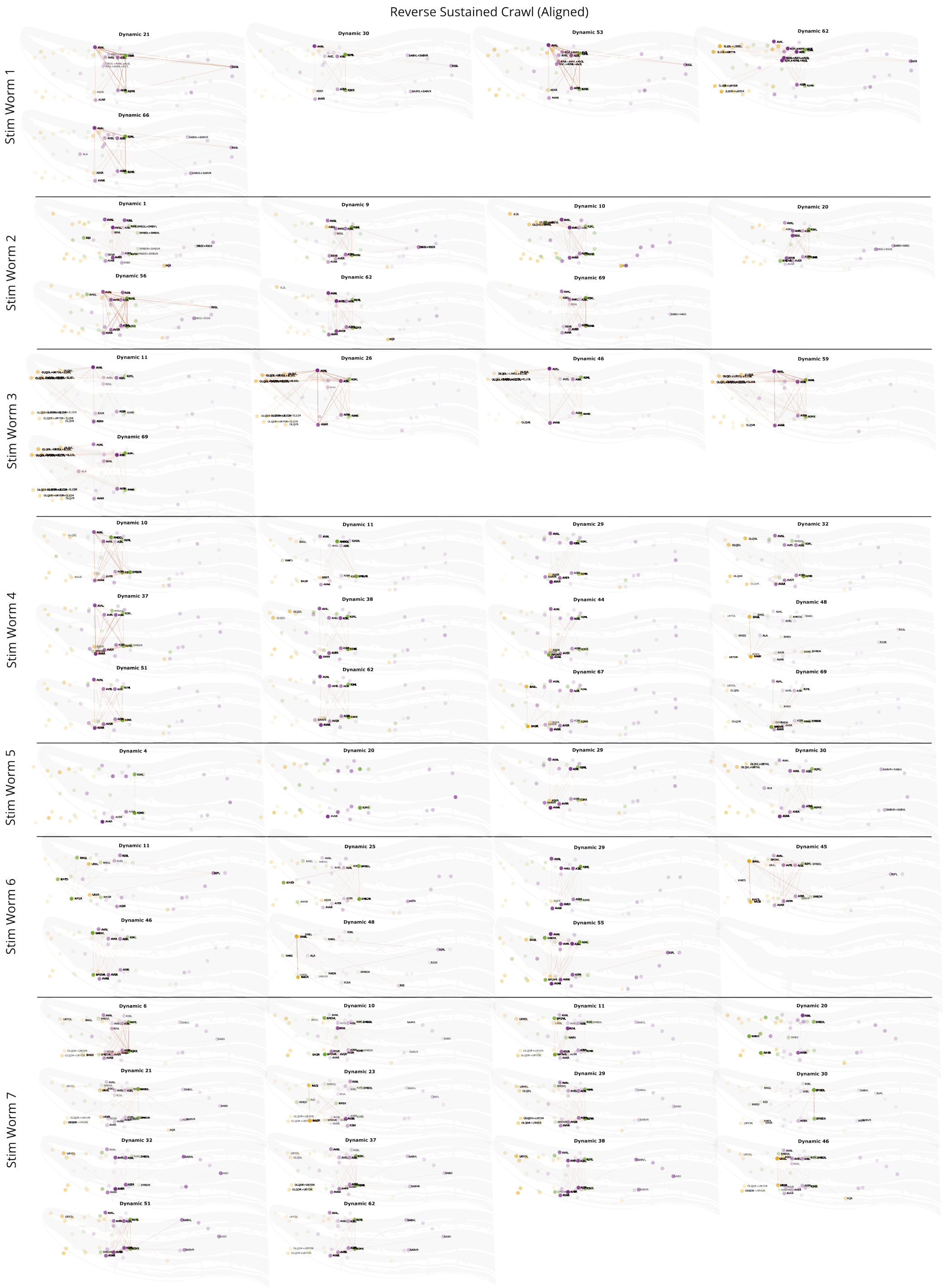


Supplementary Figure 44: **Connectivity maps by aligned dynamics operator for each Stim worm during Reverse Sustained Crawl (State 3), time-averaged.** Strength of connection (absolute value) shown by orange line width. Neuron opacity corresponds to reconstructed activity. Neuron color corresponds to class: yellow sensory, green interneuron, purple motor.

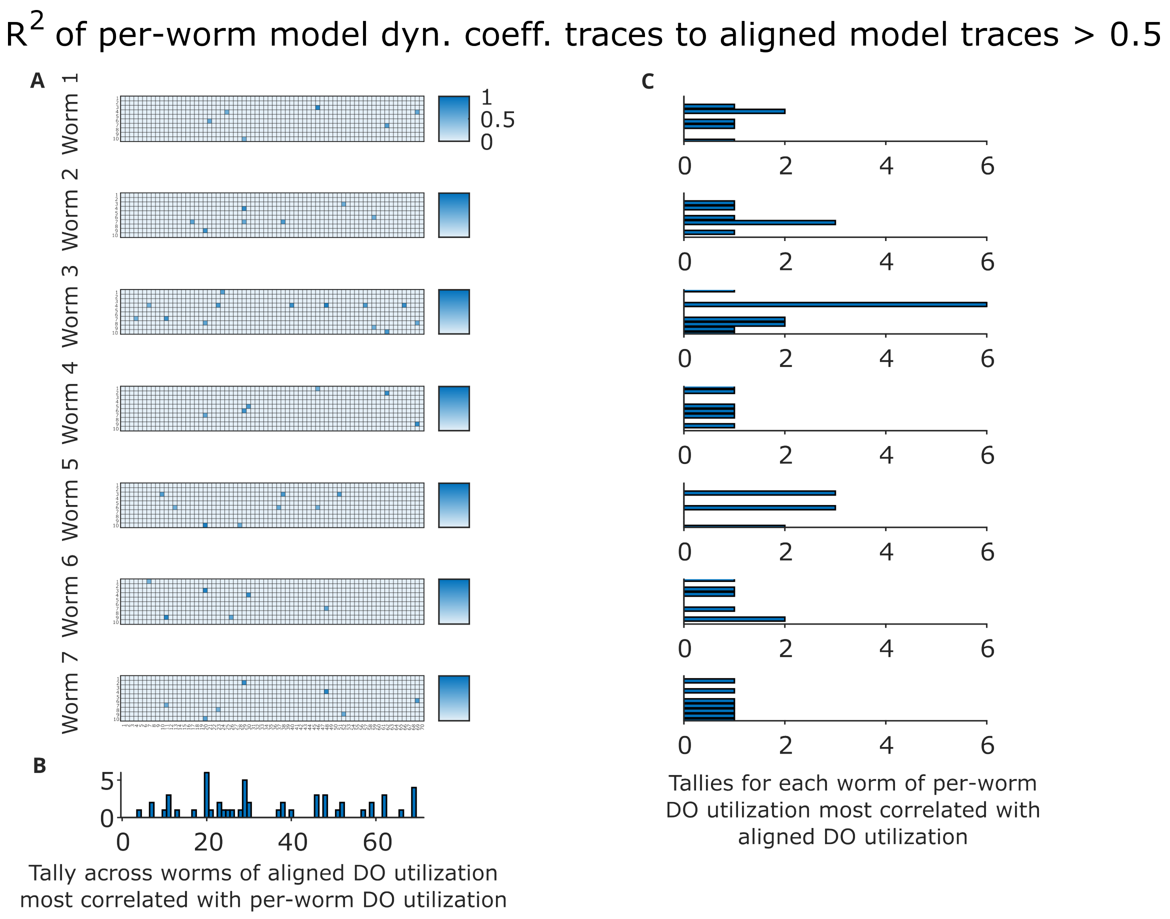


Supplementary Figure 45: **Correlation between dynamics coefficient time traces in each Stim worm's per-worm vs. aligned models. A:** Only $R^2$ values above 0.5 are shown. **B:** a tally of the number of times each aligned model DO was strongly correlated in its utilization with any per-worm model DOs. **C:** a tally for each worm of the number of times each per-worm model DO was strongly correlated in its utilization with any aligned model DOs. Note: aligned model DO utilization does not strictly coincide with aligned model DO composition, as seen in Supplementary Figure 46.

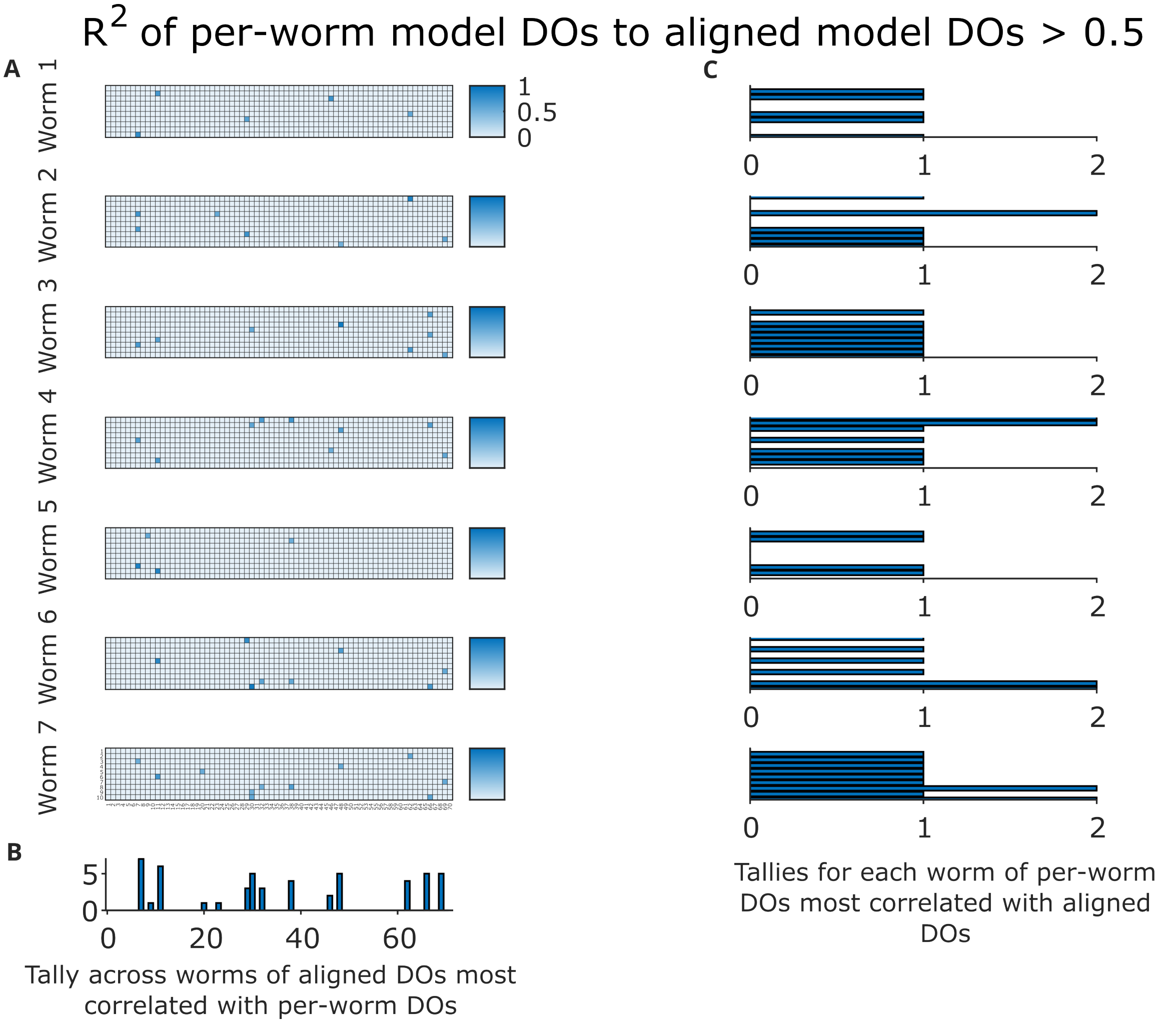


Supplementary Figure 46: **Correlation between dynamics operators (matrices) in each Stim worm's per-worm vs. aligned models.** Only $R^2$ values above 0.5 are shown. **Right column:** a tally for each worm of the number of times each per-worm model DO was strongly correlated with any aligned model DOs. **Left column, bottom:** a tally of the number of times each aligned model DO was strongly correlated with any per-worm model DOs.

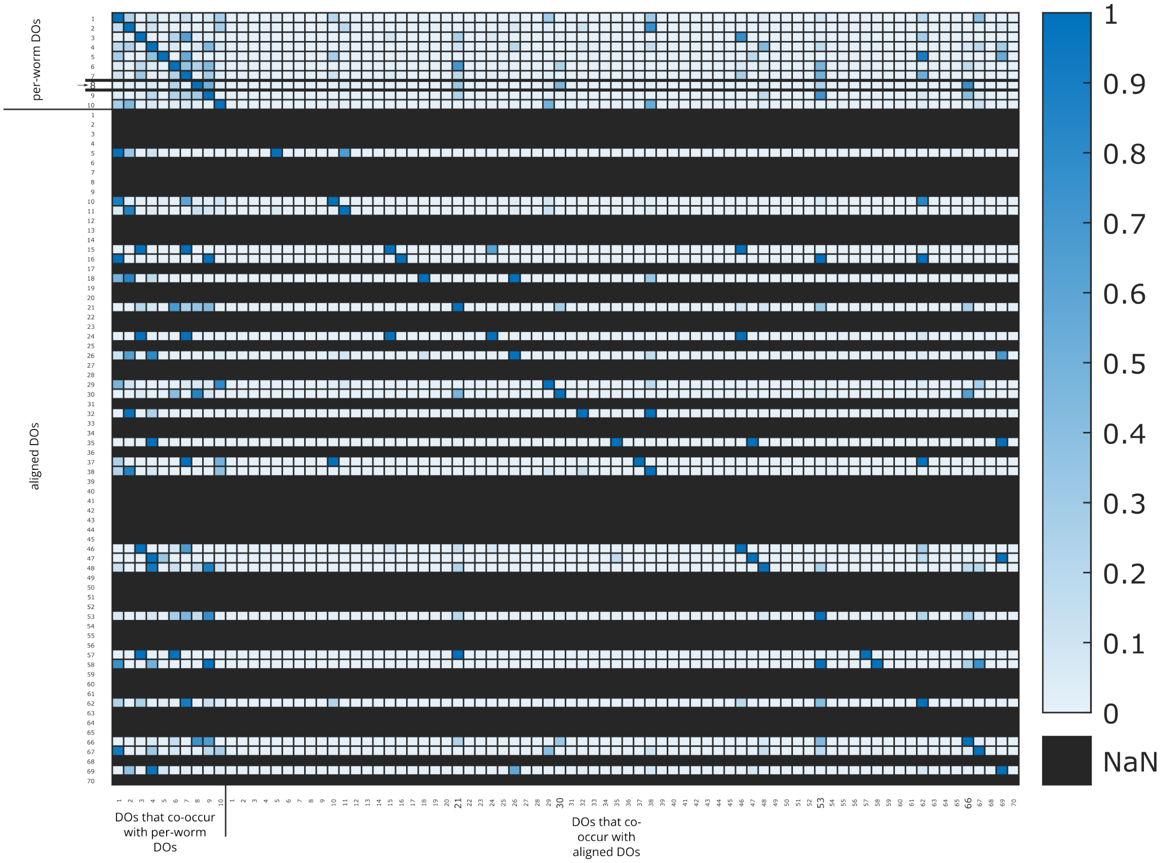


Supplementary Figure 47: **Co-utilization in time of per-worm and aligned DOs, Stim Worm 1.** Per-worm operator 8 is highlighted. A number of aligned operators were not utilized. There is a marked asymmetry, in that both the per-worm DOs and aligned DOs usually co-occur with multiple per-worm DOs densely, but the per-worm and aligned DOs co-occur with a sparser, more distributed set of aligned DOs.
